# Genomic divergence shaped the genetic regulation of meiotic homologous recombination in *Brassica* allopolyploids

**DOI:** 10.1101/2024.12.10.627763

**Authors:** Alexandre Pelé, Matthieu Falque, Maryse Lodé-Taburel, Virginie Huteau, Jérôme Morice, Olivier Coriton, Olivier C. Martin, Anne-Marie Chèvre, Mathieu Rousseau-Gueutin

## Abstract

The tight regulation of meiotic recombination between homologs is disrupted in *Brassica* AAC allotriploids, a genomic configuration that may have facilitated the formation of rapeseed (*Brassica napus* L.) ∼7,500 years ago. Indeed, the presence of the haploid C genome induces supernumerary crossovers between homologous A chromosomes with dramatically reshaped distribution. However, the genetic mechanisms driving this phenomenon and their divergence between nascent and established lineages remain unclear. To address these concerns, we generated hybrids carrying additional C chromosomes derived either from an established lineage of the allotetraploid *B. napus* or from its diploid progenitor *B. oleracea*. We then assessed recombination variation across twelve populations by mapping male meiotic crossovers using Single Nucleotide Polymorphism markers evenly distributed across the sequenced A genome. Our findings reveal that the C09 chromosome of *B. oleracea* is responsible for the formation of additional crossovers near pericentromeric regions. Interestingly, its counterpart from an established lineage of *B. napus* shows no significant effect on its own, despite having a similar content of meiotic genes. However, we showed that the *B. napus* C09 chromosome influences crossover formation through inter-chromosomal epistatic interactions with other specific C chromosomes. These results provide new insights into the genetic regulation of homologous recombination in *Brassica* and emphasize the role of genomic divergence since the formation of the allopolyploid *B. napus*.

## Introduction

Meiotic recombination is a major driver of genetic diversity, contributing to plant evolution, diversification and adaptation (Otto and Lenormand, 2002; Rice, 2002). Indeed, homologous crossovers result in reciprocal exchanges of DNA between non-sister chromatids, therefore creating novel allelic combinations. However, homologous recombination is tightly regulated (for review see Mercier *et al*., 2015). While early Prophase I of meiosis typically leads to hundreds of DNA Double Strand Breaks (DSBs) (*e.g.*, 150-250 in *Arabidopsis thaliana*), only a small fraction is repaired as crossovers (*e.g.*, ∼10 in *A. thaliana*) (Chelysheva *et al*., 2007; Sanchez-Moran *et al*., 2007; Ferdous *et al*., 2012). Most eukaryotes experience one obligatory crossover per chromosome pair and per meiosis to ensure their faithful segregation, but rarely more than three (Mercier *et al*., 2015). Such limitation stems from diverse proteins promoting the repair of DSBs into non-crossovers, such as FANCM, RECQ4, FIGL1 or FANCC in *A. thaliana* (Crismani *et al*., 2012; Girard *et al*., 2015; Séguéla-Arnaud *et al*., 2015; Singh *et al*., 2023), as well as from the so-called phenomenon of interference that reduces the likelihood of two close-by crossovers (Muller, 1916; Jones, 2006; von Diezmann and Rog, 2021). Among the two parallel pathways giving rise to crossovers in plants, only the Class I is subject to significant interference and relies on ZMM complexes (for ZIP1-4, MER3 and MSH4/5 in yeast). A minority of crossovers can also form by a MUS81-dependent pathway, referred to as Class II (∼1 crossover per meiosis in *A. thaliana*) (Mézard *et al*., 2007; Osman *et al*., 2011). Furthermore, crossovers are unevenly distributed along chromosomes (Mercier *et al*., 2015; Mézard *et al*., 2015; Dłużewska *et al*., 2018). Typically, crossover rate tends to gradually increase toward chromosomes extremities while centromeres are totally deprived of any recombination event. This pattern is particularly marked in crops like bread wheat, rapeseed, maize, and potato for which crossover suppression may extend over dozens of Mega base pairs around centromeres (Darrier *et al*., 2017; Marand *et al*., 2017; Kianian *et al*., 2018; Boideau *et al*., 2021). Fine-resolution studies conducted in *A. thaliana* and some crops highlighted the influence of genomic and epigenetic features on crossover formation. Crossovers are prevalent at distal regions that are enriched in genes and in H3K4me3 histone marks, and which present a low nucleosome density and methylation level (Choi *et al*., 2013, 2015; Yelina *et al*., 2015; Marand *et al*., 2017; Kianian *et al*., 2018). On the contrary, crossovers are almost totally absent from the pericentromeric regions that are rich in transposable elements and in repressive H3K9me2 histone marks, dense in nucleosomes and heavily methylated (Sequeira-Mendes *et al*., 2014, Yelina *et al*., 2015, Underwood *et al*., 2018; Fernandes *et al*., 2019; Rowan *et al*., 2019; Fernandes *et al*., 2023).

Despite this strict regulation, the limits of homologous recombination can be largely and naturally overcome in plants by modifying the ploidy level. Early observations were made in *Brassica* allotriploids (2*n*=3x=29, AAC) derived from the cross between the allotetraploid *B. napus* (2*n*=4x=38, AACC) and its diploid parental species *B. rapa* (2*n*=2x=20, AA). Compared to its diploid and allotetraploid counterparts, the allotriploid hybrids showed a genome-wide crossover rate 1.7 to 3.4-fold higher between homologous A chromosomes during male and female meiosis, respectively (Leflon *et al*., 2010; Pelé *et al*., 2017a; Boideau *et al*., 2021). Moreover, this unprecedented boost in crossover numbers was accompanied by reduced interference and dramatic changes in the shape of recombination landscapes. The presence of the haploid C genome led to the formation of crossovers in the pericentromeric regions of A chromosomes, which are entirely devoid of recombination events in diploids and allotetraploids (Pelé *et al*., 2017a; Boideau *et al*., 2021; Boideau *et al*., 2024). Recently, similar observations were also made in *Triticum*, with 3 to 4-fold higher crossover rates and pericentromeric recombination between homologous A and homologous B genomes of pentaploid (AABBD, 2*n*=5x=35) *versus* hexaploid hybrids (AABBDD, 2*n*=6x=42) (Yang *et al*., 2022). As for *Brassica* allotriploids, higher rates of homologous recombination were observed when the haploid genome derived from established allopolyploid lineages rather than from diploid progenitors (Pelé *et al*., 2017a; Yang *et al*., 2022). Such a crossover reshaping may have significantly contributed to the speciation success of *B. napus* (7,500 years ago) whose origin likely required a triploid bridge (Husband, 2004; Chalhoub *et al*., 2014; Mason *et al*., 2011; Pelé *et al*., 2018; Cao *et al*., 2023). Note that AAC plants are fertile and may produce allotetraploid individuals by self-fertilization (Leflon *et al*., 2006; Pelé *et al*., 2018). However, little is known about this phenomenon that can also be used as an efficient tool to reduce linkage disequilibrium in plant breeding programs (Blary and Jenczewski, 2019; Boideau *et al*., 2021; Tourrette *et al*., 2021; Capilla-Pérez *et al*., 2024).

In a previous study, we tested the hypothesis of whether the increased crossover rate in AAC allotriploids results from the nature or the number of additional *B. oleracea* C chromosomes by investigating aneuploid hybrids (Suay *et al.,* 2014). The most noticeable result of that work emerged from a hybrid carrying a complete diploid A genome and the single haploid C09 chromosome. This hybrid showed intermediate crossover rate, for the linkage group studied (A07), when compared to diploid AA and allotriploid AAC plants. On the contrary, hybrids carrying either the single C06 or the chromosomes C01, C02 and C03 together exhibited a similar crossover level to the diploid AA plant. Taken together, and by analyzing other aneuploid hybrids, we pointed out that the *B. oleracea* C09 chromosome may act as a major determinant in the variation of crossover frequency in AAC allotriploids, and hypothesized that C04 and C08 chromosomes may have a minor contribution. However, our previous study was only performed on the A07 chromosome using a dozen markers, so this work should be extended to the entire A genome. Importantly, it remains unknown whether the C09, or another C chromosome, is responsible for the modified distribution of crossovers and the formation of new recombining regions closer to centromeres. Finally, only the effects of additional C chromosomes derived from *B. oleracea* have been tested so far (Suay *et al*., 2014), despite the higher crossover rates found in AAC allotriploids harboring *B. napus* rather than *B. oleracea* C genome (Pelé *et al*., 2017a). Therefore, whether the genetic regulation of homologous recombination may have diverged following long-term subgenome coevolution remains to be elucidated. To determine the effect of additional C chromosomes deriving from an established lineage of *B. napus*, we took advantage of the previous extraction of *B. napus* A subgenome from the reference cultivar ‘Darmor’ (Pelé *et al*., 2017b).

In the present work we investigate more deeply the deregulation of homologous recombination in *Brassica* AAC allotriploids. To that end, we created a unique plant material enabling us to assess the impact of the C09 chromosome of *B. oleracea* or *B. napus* on male meiosis. We generated and then analyzed twelve segregating populations through Single Nucleotide Polymorphism markers covering homogeneously the whole A genome. Our results confirm that the C09 chromosome of *B. oleracea* acts as a major determinant in the crossover rate variations arising in allotriploids, all the while pointing out its key role in crossover distribution and interference changes. Strikingly, we revealed for the first time that this genetic regulation differs when this chromosome originates from an established lineage of *B. napus*, despite its similar content for meiotic genes with the C09 chromosome of *B. oleracea*. Indeed, we pointed out that crossover variation relies on inter-chromosomal epistatic interactions between *B. napus* C chromosomes, which suggests a role of genomic divergence.

## Materials and methods

### Plant materials

Two types of hybrid construction were used to study the impact of the *B. oleracea* or *B. napus* C09 chromosome on homologous recombination in *Brassica* AAC allotriploids (Pelé *et al*., 2017a). The production of these hybrids, using *B. rapa*, *B. oleracea* and *B. napus* seeds from the Genetic Resource Center BrACySol (UMR IGEPP, Ploudaniel, France), is illustrated in Fig. 1a and b, with the corresponding chromosomal composition shown in Fig. 1c.

**Fig. 1.**
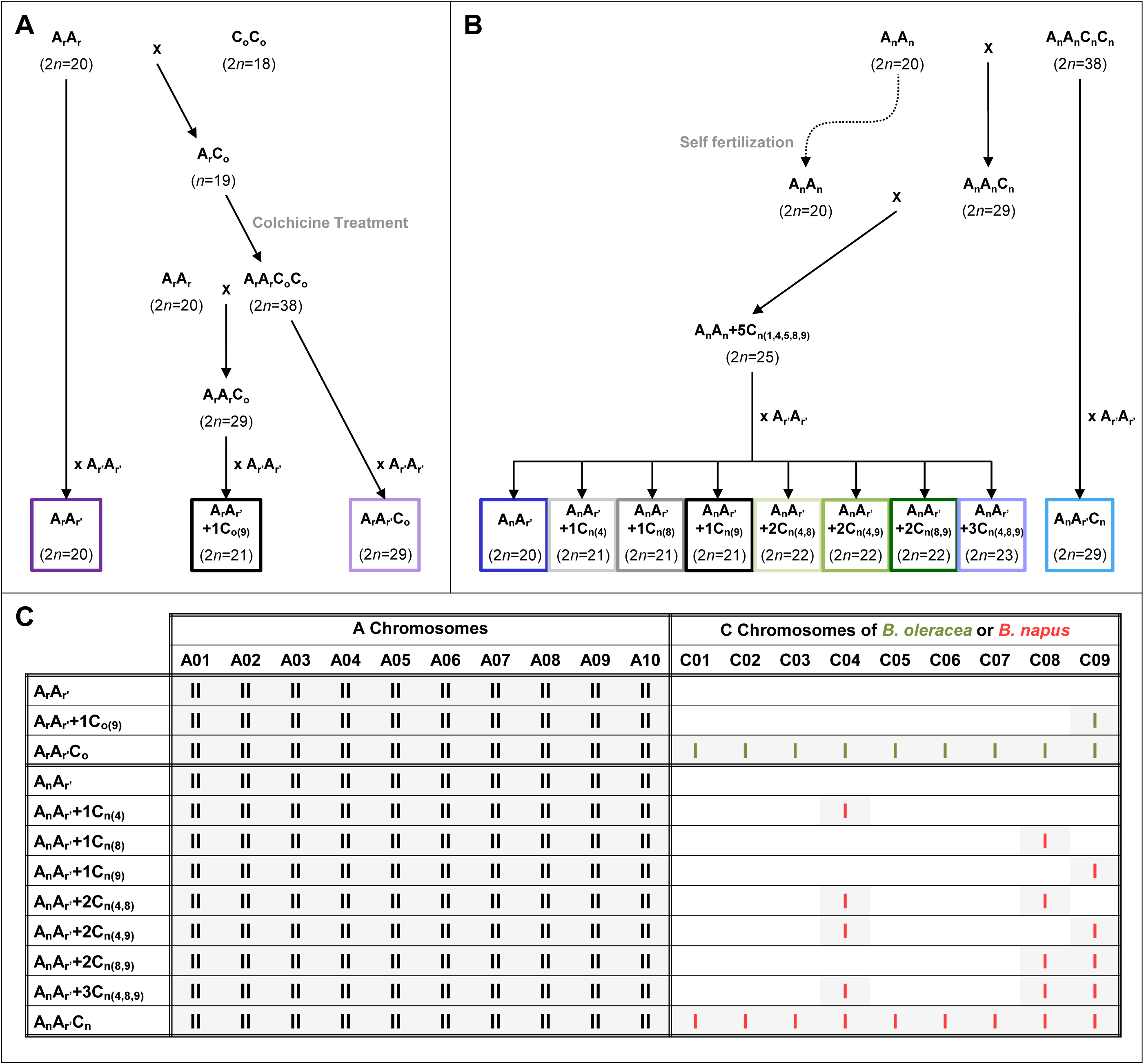
Figure detailing the production of the two sets of hybrids sharing an identical A genome with additional C chromosomes deriving either from (A) *B. oleracea* or (B) *B. napus*, along with their (C) chromosomal composition. A_r_A_r_ and A_r’_A_r’_ represent the *B. rapa* cultivars ‘C1.3’ and ‘Chiifu-401’, respectively. C_o_C_o_ designates the *B. oleracea* cv. ‘RC34’, and A_n_A_n_C_n_C_n_ corresponds to the allotetraploid *B. napus* cv. ‘Darmor’. The A_n_A_n_ plant represents the diploid AA component extracted from the allotetraploid *B. napus* cv. ‘Darmor’ (Pelé *et al*. 2017b). I and II indicate univalent and bivalent chromosomes, respectively. Green and red bars refer to *B. oleracea* or *B. napus* C chromosomes, respectively.

For the first type of hybrids, a sequence of crosses was designed to assess the genome-wide effect of the *B. oleracea* C09 chromosome (Fig. 1a). For that purpose, we used the previously generated F1 hybrids including the diploid A_r_A_r’_ (2*n*=2x=20), the allotriploid A_r_A_r’_C_o_ (2*n*=3x=29) and the A_r_A_r’_+1C_o(9)_ hybrid (2*n*=21) (Suay *et al*., 2014; Pelé *et al*., 2017a), that exhibit identical A genotypes and display either no C_o_ chromosome in A_r_A_r’_, the single C09 in A_r_A_r’_+1C_o(9)_ or the full set (C01 to C09) in A_r_A_r’_C_o_ (Fig. 1c).

For the second type of hybrids, the sequence of crosses was intended to assess the genome-wide effect of the *B. napus* C09 chromosome (Fig. 1b). As controls, we exploited the diploid A_n_A_r’_ (2*n*=2x=20) and the allotriploid A_n_A_r’_C_n_ (2*n*=3x=29) hybrids, generated by Pelé *et al*. (2017a). Moreover, we produced an A_n_A_r’_+1C_n(9)_ hybrid with very similar A genotype and displaying the single additional *B. napus* C09 chromosome versus none in A_n_A_r’_ and the full haploid set of C_n_ chromosomes in A_n_A_r’_C_n_ (Fig. 1c). To that end, we took advantage of the extraction of the diploid A_n_A_n_ component (2*n*=2x=20) (Pelé *et al*., 2017b) from the allotetraploid *B. napus* cv. ‘Darmor’ (A_n_A_n_C_n_C_n_, 2*n*=4x=38). The diploid A_n_A_n_ plant was first crossed as female with (1) *B. napus* cv. ‘Darmor’ to obtain an allotriploid A_n_A_n_C_n_ (2*n*=3x=29), which was then crossed to the selfed diploid A_n_A_n_ plant. The 84 progenies were screened with SSR markers specific to each C_n_ chromosome (Table S1). A plant displaying the C01, C04, C05, C08 and C09 chromosomes (referred to as A_n_A_n_+5C_n(1,4,5,8,9)_), was selected and crossed with *B. rapa* ‘Chiifu-401’. Among the 282 progenies, we identified an A_n_A_r’_+1C_n(9)_ hybrid. Furthermore, we also selected hybrids that carry an A_n_A_r’_ genome and the C04, C08 and C09 chromosomes (either alone or in combination), hereafter referred to as A_n_A_r’_+1C_n(4)_, A_n_A_r’_+1C_n(8)_, A_n_A_r’_+2C_n(4+8)_, A_n_A_r’_+1C_n(4+9)_, A_n_A_r’_+1C_n(8+9)_ and A_n_A_r’_+3C_n(4+8+9)_.

For each of the final F1 hybrids, segregating populations were produced and genotyped to assess the recombination during male meiosis between the 10 pairs of homologous A chromosomes (Table 1). Indeed, due to the low vigor of plants, progenies were generated by crossing F1 hybrids as male to *B. napus* cv. ‘Darmor’ for the first type of hybrids, and to the Korean spring rapeseed line *B. napus* cv. ‘Yudal’ for the second type of hybrids. All progenies were obtained by manual pollination in the same environmental conditions.

**Table 1.**
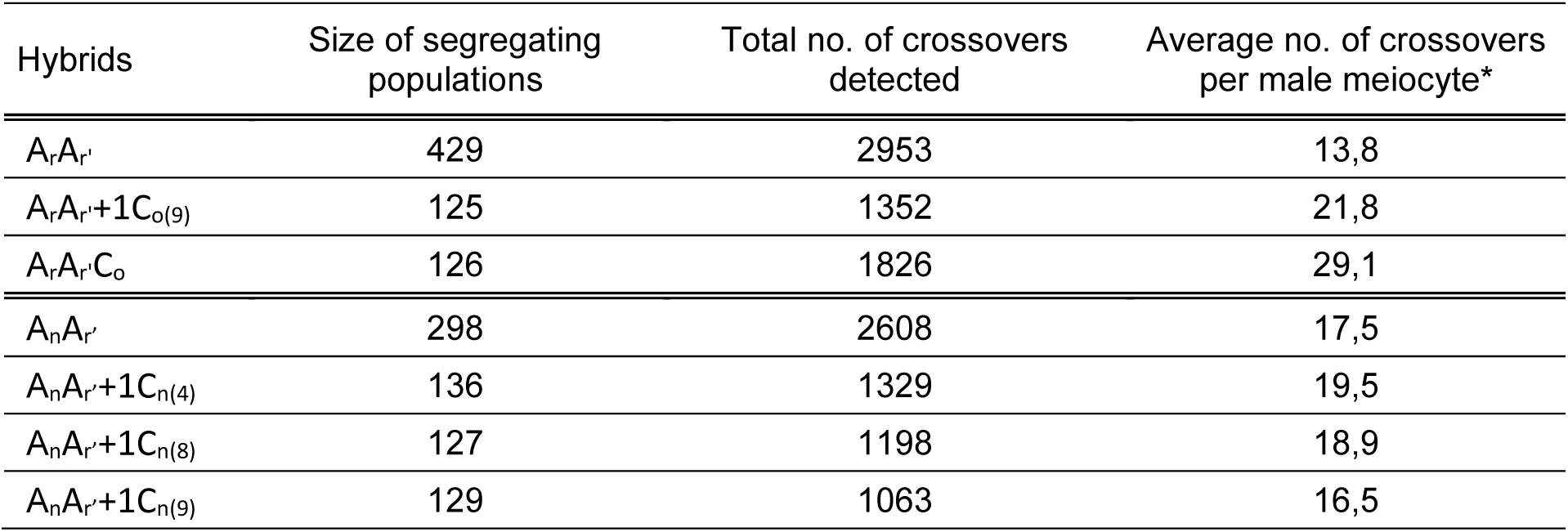
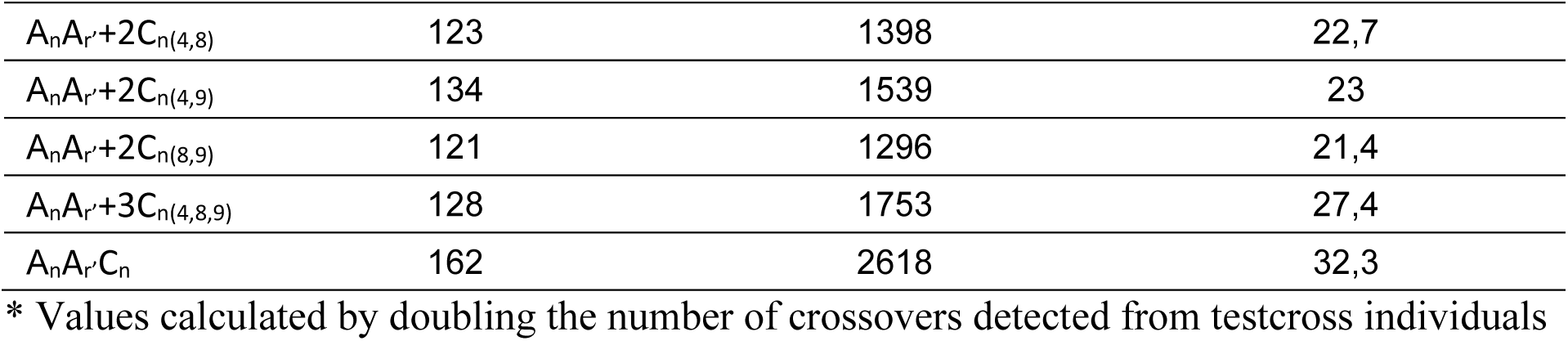
Size of segregating populations analyzed per hybrid and the number of male crossovers detected between the homologous A genomes.

### Cytogenetic characterization

Briefly, young floral buds of all hybrids were harvested to assess the meiotic behavior from 20 Pollen Mother Cells (PMCs) by scoring Metaphase I pairing following the protocol of Suay *et al*. (2014). BAC-FISH experiments were then performed using the *B. oleracea* Bob014O06 BAC clone in a genomic in situ hybridization (GISH)-like approach, which allows for distinguishing the C chromosomes (Howell *et al*., 2002), as well as the *B. rapa* KBrH033J07, KBrB043F18 and KBrH080A08 BAC clones that specifically hybridize to the A05/C04, A09/C08, A10/C09 chromosomes, respectively (Xiong *et al*., 2011). BAC clones were labelled by random priming, with either Alexa 488-5-dUTP or biotin-14-dUTP (Invitrogen, life technologies).

### DNA preparation and plant selection

Genomic DNA from lyophilized young leaves of each sample was extracted with the sbeadex^TM^ maxi plant kit (LGC Genomics, Teddington Middlesex, UK) on the oKtopure^TM^ robot at the GENTYANE platform (INRAE, Clermont-Ferrand, France).

The nature of additional C_n_ chromosomes in hybrids was examined using SSR markers described by Suay *et al*. (2014) (Table S1). Polymerase Chain Reaction assays were performed on a 16 Capillary ABI Prism 3130xl, as described by Esselink *et al*. (2004). The size of PCR products was visualized using GeneMapper^TM^ v3.7 (Applied Biosystems).

### Genomic characterization

To ensure the occurrence of the C09 chromosome in the A_n_A_r’_+1C_n(9)_ hybrid and validate the absence of homoeologous exchanges while producing this material, we genotyped this hybrid using the 60K Illumina^®^ SNP array as well as 3 technical replicates of the lines used for its creation: *B. rapa* cv. ‘Chiifu-401’ and ‘C1.3’, and *B. napus* cv. ‘Darmor’. The physical positions of these SNPs were retrieved from *B. napus* cv. ‘Darmor’ v10 (Rousseau-Gueutin *et al*. 2020). The genotyping data obtained were visualized using Genome Studio V2011.1 software (Illumina, Inc., San Diego, CA, USA) and processed with a manually adapted cluster file. To ensure that the SNPs were specific to the C chromosomes and did not amplify on both homoeologous chromosomes of *B. napus,* we first retrieved the SNPs amplifying in *B. napus* cv. ‘Darmor’ but not in *B. rapa* cv. ‘C1.3’ and ‘Chiifu-401’. The physical locations of these SNPs were then compared with their genetic positions obtained from a *B. napus* Darmor-Yudal genetic map, generated using the same 60K SNP array (Bilgrami *et al*., 2023). Finally, the amplification of the 4,807 SNPs having a concordant physical and genetic location on the same C chromosome were considered in the A_n_A_r’_+1C_n(9)_ hybrid.

### Recombination assessment between pairs of A chromosomes

Evaluation of crossover rate between pairs of homologous A chromosomes was conducted from a genotyping approach per segregating population. To that end, we used 169 KASP markers, 163 of which were common between hybrids and six specific to each type, as employed Pelé *et al*. (2017a) to compare the same diploid and allotriploid hybrids. The physical positions of these markers (Table S2) were obtained using *B. rapa* cv. ‘Chiifu-401’ v1.5 genome assembly (Wang *et al*., 2011), therefore enabling comparisons with our previous study (Pelé *et al*., 2017a). Following the synthesis of SNP primers (LGC Genomics, Teddington Middlesex, UK), the genotyping was conducted by the GENTYANE platform (INRAE, Clermont-Ferrand, France) using the Biomark^TM^ HD system (Fluidigm technology) and KASPar^TM^ chemistry (Wang *et al*., 2009). Hybridizations were run according to the platform procedures using 96.96 Dynamic Array™ IFC components. Technical replicates of each hybrid and of parental lines were always included. The obtained genotyping data were visualized using Fluidigm SNP Genotyping Analysis V4.1.2 software (Wang *et al*., 2009) and processed manually.

From these processed genotyping data, we first discarded the few samples with missing data, giving rise to 125 to 429 individuals analyzed per segregating population (Table 1). Then, after validating the polymorphism and Mendelian segregation of all SNP markers, the linkage analyzes were conducted using CarthaGene software version 1.3 (De Givry *et al*., 2005). From each segregating population, the linkage groups were established separately at a Logarithm of Odds Score (LOD) threshold of 4.0. The order of SNP markers was then estimated by using the multiple 2-point maximum likelihood method at a LOD threshold of 3.0 and a maximum recombination frequency of 0.4. Finally, the Kosambi function was applied to evaluate the genetic distances in centimorgan (cM) between linked SNP markers (Kosambi, 1943).

### Statistical analysis

Mendelian segregation was verified for each SNP marker from Chi-squared tests at a significance threshold of 5%.

Heterogeneity of crossover rates among progenies was assessed for every interval between adjacent SNP markers using a 2-by-2 Chi-squared analysis and considering a significance threshold of 5%. A conservative Bonferroni-corrected threshold of 5% was applied for assessing the heterogeneity of crossover rates among progenies at chromosome and genome scales, using the number of intervals between adjacent SNP markers per A chromosome and for the A genome-wide, respectively (Rice, 1989).

The following relationships were studied by regression analyses conducted after the visual validation of residuals’ normality: (i) the average number of crossovers formed per A chromosome in a meiosis *vs* their physical length covered by SNP markers (in Mbp); (ii) the relative recombination rates normalized per A chromosome (%) *vs* their relative distance from the centromeres (%), using the centromeres’ locations determined by Mason *et al*. (2016). For each relationship tested, p-values were provided by Fisher-Tests.

### Comparisons of crossover landscapes and interference using the Kullback-Leibler divergence

Given discrete classes (i=1,…,C) and two associated probability distributions ({P_i_, i=1,…,C} and {Q_i_, i=1,…,C}, the Kullback-Liebler divergence (hereafter denoted KL) between P and Q is defined as KL (P, Q) = Σ_i_ P_i_ log(P_i_/Q_i_). When the two distributions P and Q are identical, the divergence vanishes, otherwise it is strictly positive. We use KL to provide a quantitative measure of the difference between P and Q when considering recombination landscapes or the distributions of inter-crossover distances. However, since those distributions relate to continuous variables, we have to introduce binnings before being able to use the KL framework.

#### [I] The KL measure of divergence between a genotype’s recombination landscape and the flat landscape

For the binning, we first segmented each chromosome into 10 bins of identical physical size.

##### (1) Value of the KL divergence

We used all Q_i_ = 1/10 (because there are 10 bins) as the flat landscape, and we set P_i_ to the number of crossovers in the i’th bin followed by normalization to obtain a probability distribution. Those 20 values allowed us to calculate the KL divergence KL (P, Q) between the genotype’s landscape and the flat one, for each chromosome. (We used the KL.plugin(P,Q) function of the R package « entropy ».) The case of pooled chromosomes was obtained simply by pooling together all the crossovers when constructing the P_i_.

##### (2) p-value when rejecting the hypothesis H0 that the data is statistically compatible with a flat landscape

We generated the distribution of the KL divergence under H0 by simulation as follows. For each chromosome, we produced 10,000 randomizations of the population’s crossovers, repositioning the crossovers randomly according to a flat distribution. For each randomization, we calculated its KL divergence to the flat landscape as previously. That generated the expected (under H0) distribution for KL values (for each chromosome, but the same method was used for the pooled chromosomes as explained above). Then the p-value associated with rejecting H0 was the proportion of the 10,000 simulated KL values that were at least as large as the KL value found when using the actual (non-randomized) dataset.

##### (3) Comparing these KL values across different genotypes

The KL values of different genotypes quantify their divergence to the flat landscape Q_i_=1/10 and provide an ordering therein for the degree of flatness of their landscapes. To determine if the orderings were statistically significant, we performed the following calculations. (i) We first determined the statistical distribution of KL for each genotype by the bootstrap method. Specifically, given the population of N individuals associated with a given genotype, draw N individuals from that population with replacement. For each such sampling, compute the KL divergence to the flat landscape (Q_i_ =1/10) when using those bootstrap individuals. Doing that 10,000 times, we obtained the distribution of the KL measure of divergence to flatness for that genotype. (ii) For any given pair of genotypes, we then counted for these simulated KL values the number k of occurrences where the ordering was opposite from the one in the experimental dataset. The p-value associated with the hypothesis H0 that the ordering in the experimental dataset for that pair is incorrect was then k/10,000. Such an approach can be used to produce a p-value when comparing any pair, but when comparing all pairs we applied the Bonferroni correction. The associated (adjusted) p-values (obtained by multiplying by the number of comparisons) were used to define groups for the different genotypes. These groups are then labeled a, b, c etc. containing members having high to low KL values, two separate groups having an adjusted p-value less than 5% for their comparison.

#### [II] The KL measure of divergence between the recombination landscapes of two genotypes

##### (1) Value of that KL divergence

The procedure is similar to that used above, the key difference being that both distributions in the expression KL (P, Q) are derived from the data: P is the histogram from the first genotype’s landscape and Q is that from the second.

##### (2) Confidence intervals for each KL value

To obtain the confidence interval of a KL value (with two fixed genotypes), we performed bootstrapping on each of the associated crossover datasets, leading to 10,000 simulated crossover datasets for the first genotype and the same for the second. We then computed the 10,000 associated KL values. The confidence interval was then obtained from that empirical KL distribution via quantiles at 0.025 and 0.975.

#### [III] Using the KL measure of divergence to probe interference strengths

To produce the data to be analyzed for each chromosome, we focused on the genetic distances between adjacent crossovers. The genetic length of the chromosome of interest was discretized into 10 equal-sized bins to produce a distribution of these inter-crossover distances. In the case of our analysis for all chromosomes, the data from all chromosomes were pooled before applying the binning.

##### (1) Value of the KL divergence to the "no interference" case

The H0 hypothesis here corresponds to having no interference. The objective is to measure KL (P, Q) where P is the distribution of (normalized) distances between adjacent crossovers while Q is the corresponding distribution in the absence of interference. Even though Q is not known *a priori*, it corresponds to having crossovers arise independently. To simulate that situation, we removed all correlations between crossovers by shuffling them across different individuals in the population. We did this 10,000 times to generate our reference "no interference" distribution of crossover positions. Note that the shuffling maintains the recombination landscape but removes any dependency (interference) between crossovers. The dataset of these shuffled crossovers was then used to construct Q which was then fed into the formula of KL (P, Q).

##### (2) p-value when rejecting the hypothesis H0 that the data is statistically compatible with no interference

Given the experimental distribution of crossovers, the use of one random shuffle "S" allows one to produce one instance P_S_ of a histogram of (normalized) distances between adjacent crossovers under H0 and thus one value KL (P_S_, Q). Repeating this for 10,000 different shuffles, we obtained the empirical distribution of KL under H0. The p-value when rejecting H0 was then just the fraction of cases where these KL (P_S_, Q) were at least as large as KL (P, Q).

##### (3) Comparing these KL values across different genotypes

In direct analogy with the case of landscapes, the KL divergences to the "no interference" situation can be used to order different genotypes according to their interference strength. We determined the statistical significance of that ordering in the same way as was done for the landscapes, that is we used the bootstrap (resampling) approach to obtain the distribution of the KL divergence for each genotype. Those distributions were then used to compute the p-values for all pair-wise comparisons (again we applied the Bonferroni correction) from which we identified groups of genotypes according to their KL level, groups labeled a, b, c etc.

### Identification of the meiotic genes content present on either B. oleracea cv. ‘RC34 or B. napus cv. ‘Darmor’ C chromosomes

To identify the meiotic genes present on the *B. oleracea* cv. ‘RC34’ and *B. napus* cv. ‘Darmor’ C chromosomes, we first retrieved a list of Arabidopsis meiotic genes derived from Lloyd *et al*. (2014) and Higgins *et al*. (2021) and then recovered *A. thaliana* (TAIR10: https://www.arabidopsis.org), *B. oleracea* cv. ‘RC34’ (Maillet *et al*., unpublished data) and *B. napus* cv. ‘Darmor’ v10 (Rousseau-Gueutin *et al*., 2021) amino-acid gene sequences to identify their orthologous relationships using Orthofinder (Emms and Kelly, 2019). Additionally, for the genes present on the C09 chromosome, we validated these analyses by performing molecular phylogeny (Maximum Parsimony, bootstrap 1000) using Mega X (Kumar *et al*., 2018). More precisely, the amino-acid gene sequences of Arabidopsis meiotic genes were self-blasted and blasted against the amino-acid gene sequences of several *B. napus* (‘Darmor’ v10: Rousseau-Gueutin *et al*., 2020; ‘RCC-S0’: Maillet *et al*., unpublished data; ‘Gangan’, ‘No2127’, ‘Zs11’, ‘Zheyou’, ‘Westar’, ‘Quinta’: Song *et al*., 2020); *B. oleracea* (‘D134’, ‘HDEM’: Belser *et al*., 2018; ‘KORSO’: Guo *et al*., 2021; ‘D134’: Lv *et al*., 2020). We then performed alignments between the orthologous *B. napus* cv. ‘Darmor’ and *B. oleracea* cv. ‘RC34’ to compare their lengths and sequence identity. For the few genes absent on the C09 chromosome of *B. oleracea* cv. ‘RC34’, Blastn analyses were performed to validate that their absence was not due to missing annotation.

## Results

### The B. oleracea C09 chromosome is responsible for the formation of additional crossovers closer to centromeres in Brassica allotriploids

To characterize the genome-wide contribution of the *B. oleracea* C09 chromosome on the crossover reshaping observed in AAC allotriploids, we conducted a genotyping approach on segregating populations generated from each hybrid presented on Fig. 1a, including the diploid A_r_A_r’_ (2*n*=20), the A_r_A_r’_+1C_o(9)_ hybrid (2*n*=21) and the allotriploid A_r_A_r’_C_o_ (2*n*=29). These hybrids carry an identical A genotype, and only differ by their content in additional *B. oleracea* C_o_ chromosomes (Fig. 1c), which remain unpaired at Metaphase I of meiosis in contrast to the A homologs that form 10 bivalents (Suay *et al*., 2014; Pelé *et al*., 2017a). Crossover detection during male meiosis was performed from the genotyping of segregating populations using 169 SNP markers distributed along A chromosomes. These markers, always showing the expected Mendelian segregation and having concordant genetic and physical positions, covered 93.6% of the sequenced A genome of *B. rapa* cv. ‘Chiifu-401’ (Wang *et al*., 2011) – used to obtain all hybrids (Fig. 1) – with on average one SNP every 1.5 Mbp (SE=0.07, n=159; Table S2). In total, these SNPs were used to genotype 125 to 429 progenies per hybrid, giving rise to a number of crossovers analyzed ranging from 1,352 to 2,953 per segregation population (Table 1).

We extended our previous work to the whole A genome scale, determining the effect of the additional *B. oleracea* C09 chromosome on crossover rates for each A chromosome. We found the following ordering between the total genetic lengths of the hybrids: A_r_A_r’_ < A_r_A_r’_+1C_o(9)_ < A_r_A_r’_C_o_ (Corrected Chi-squared test, *p*<2.2E-16; Fig. 2a), with 13.8 (SE=0.20, n=429), 21.8 (SE=0.46, n=125) and 29.1 (SE=0.56, n=126) crossovers detected on average per meiocyte, respectively. Compared to the diploid A_r_A_r’_, the average number of crossovers formed in A_r_A_r’_+1C_o(9)_ was 1.6-fold higher but remained 1.3-fold lower than in A_r_A_r’_C_o_. With regards to the total variation in crossover frequency arising from the complete C_o_ genome, about half (52.7%) was explained by the single additional *B. oleracea* C09 chromosome. Furthermore, this trend also occurred at the level of each A chromosome (Fig. S1a), which generally exhibit a greater frequency of multiple crossovers when hybrids carried either the C09 or the full set of nine C_o_ chromosomes (Fig. S2). Indeed, the average number of crossovers was always higher in the A_r_A_r’_+1C_o(9)_ hybrid than in the diploid (with factors from 1.1 to 2.0) and lower compared to the allotriploid (with factors from 1.2 to 1.7), the A01 chromosome excepted. Statistically, these observations are also verified, with a few exceptions (Fig. S1a), which may result from the differences in the sizes of A chromosomes. For each hybrid, we identified significant positive linear regressions between chromosome size (in Mbp) and their average number of crossovers (Fisher test, *p*<0.05), with R² values ranging from 0.52 to 0.77 (Fig. 2b). Thus, while the smallest A chromosomes exhibited one to two crossovers on average per meiosis for each hybrid, greater differences were found for the largest chromosomes with on average up to two crossovers in A_r_A_r’_, against three in A_r_A_r’_+1C_o(9)_ and four in A_r_A_r’_C_o_.

**Fig. 2.**
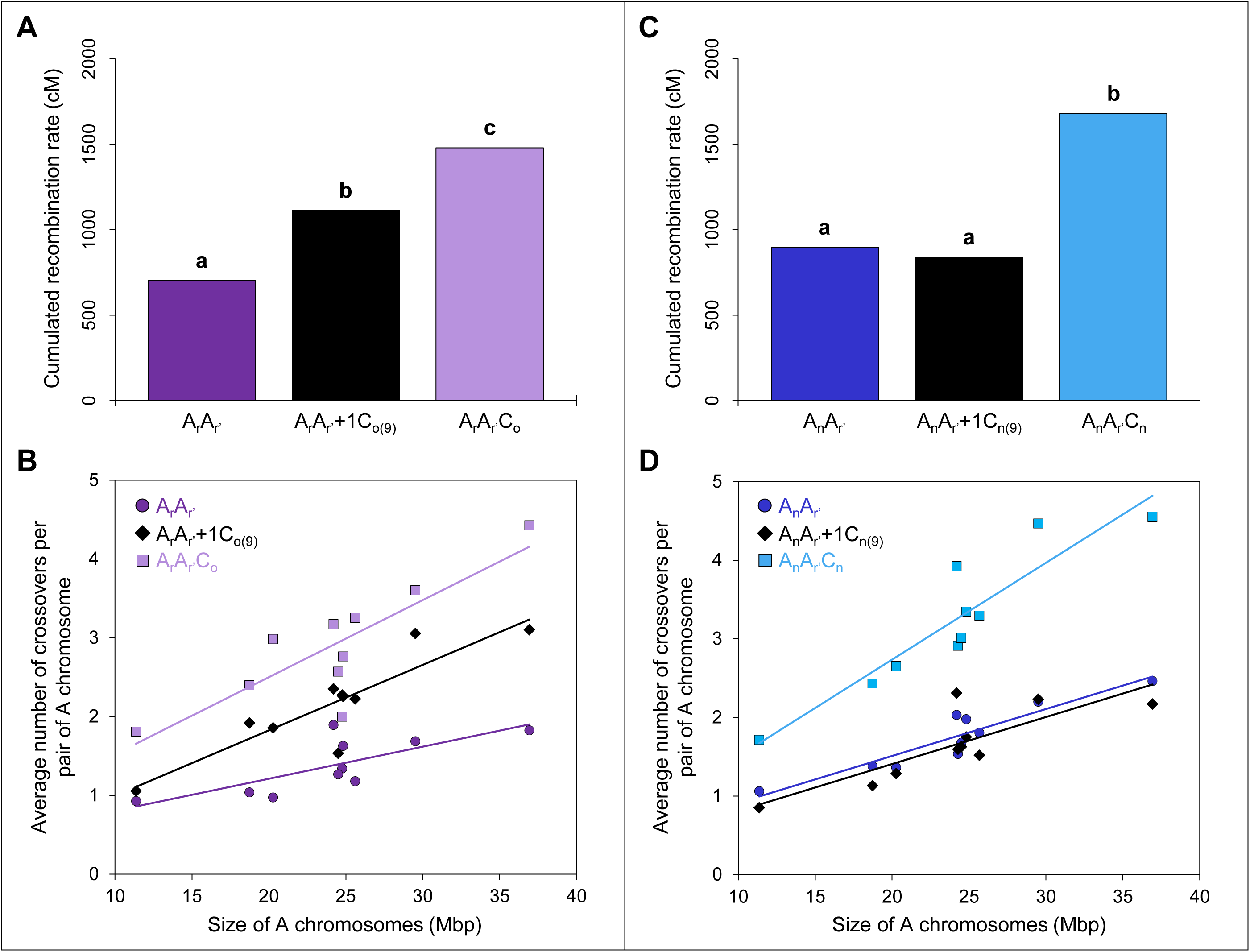
Differential contributions of the C09 chromosome from (A-B) *B. oleracea* and (C-D) *B. napus* to crossover rate variation between homologous A chromosomes in *Brassica* AAC allotriploids. **A,C.** Recombination rates in Centimorgan (cM) for the cumulated A chromosomes in hybrids carrying (**A**) *B. oleracea* C_o_ and (**B**) *B. napus* C_n_ chromosomes. Statistical differences based on a Bonferroni corrected Chi-squared test (5% threshold) are indicated with letters a to c. **B,D.** Relationship between the average numbers of crossovers per homologous A chromosome pair and their physical size (Mbp) covered by SNP markers in hybrids carrying (**C**) *B. oleracea* C_o_ and (**D**) *B. napus* C_n_ chromosomes. A_r_A_r’_ (purple circles): y=0.0407x+0.3968; R²=0.52. A_r_A_r’_+1C_o(9)_ (black diamonds): y=0.0831x+0.1633; R²=0.77. A_r_A_r’_C_o_ (light purple squares): y=0.0978x+0.5446; R²=0.68. A_n_A_r’_ (blue circles): y=0.0597x+0.315; R²=0.85. A_n_A_r’_+1C_n(9)_ (black diamonds): y=0.0598x+0.2118; R²=0.64. A_n_A_r’_C_n_ (light blue squares): y=0.1232x+0.2732; R²=0.83.

Having found that half of the extra crossovers formed between the homologous A genomes in *Brassica* allotriploids can be justified from the single additional B. oleracea C09 chromosome, we also investigated its putative role in the reshaping of crossover distribution. To do so, we followed the approach of Pelé *et al*. (2017a) that used the so-called Kullback-Leibler (KL) divergence to quantitatively compare two distributions. The KL divergence vanishes when the two distributions are identical, and it rises as they become more different. As shown qualitatively in Fig. 3a when pooling the 10 A chromosomes, the recombination landscape of A_r_A_r’_+1C_o(9)_ was closer to that of A_r_A_r’_C_o_ than to that of A_r_A_r’_. This is confirmed quantitatively by the KL values of divergence to the flat landscape, with 0.248, 0.075 and 0.054 for A_r_A_r’_, A_r_A_r’_+1C_o(9)_ and A_r_A_r’_C_o_, respectively (see Methods for details). To further support this claim, Fig. 3c shows that the divergence between landscapes is far smaller when comparing A_r_A_r’_+1C_o(9)_ to A_r_A_r’_C_o_ than when comparing it to A_r_A_r’_. Statistically, when ordering these three genotypes, we confirmed that the A_r_A_r’_+1C_o(9)_ and A_r_A_r’_C_o_ hybrids belong to the same group, and both differ significantly from the diploid A_r_A_r’_ (Fig. 3d). These trends also arise at the individual chromosome level, though the statistical tests are less powerful there; the corresponding plots are shown in Figs. S3-4, and the data are presented in Table S3-S4. Together, these results suggest that the single additional C09 chromosome of *B. oleracea* modifies the global crossover distribution along A chromosomes in a similar way as the complete haploid C_o_ genome does.

**Fig. 3.**
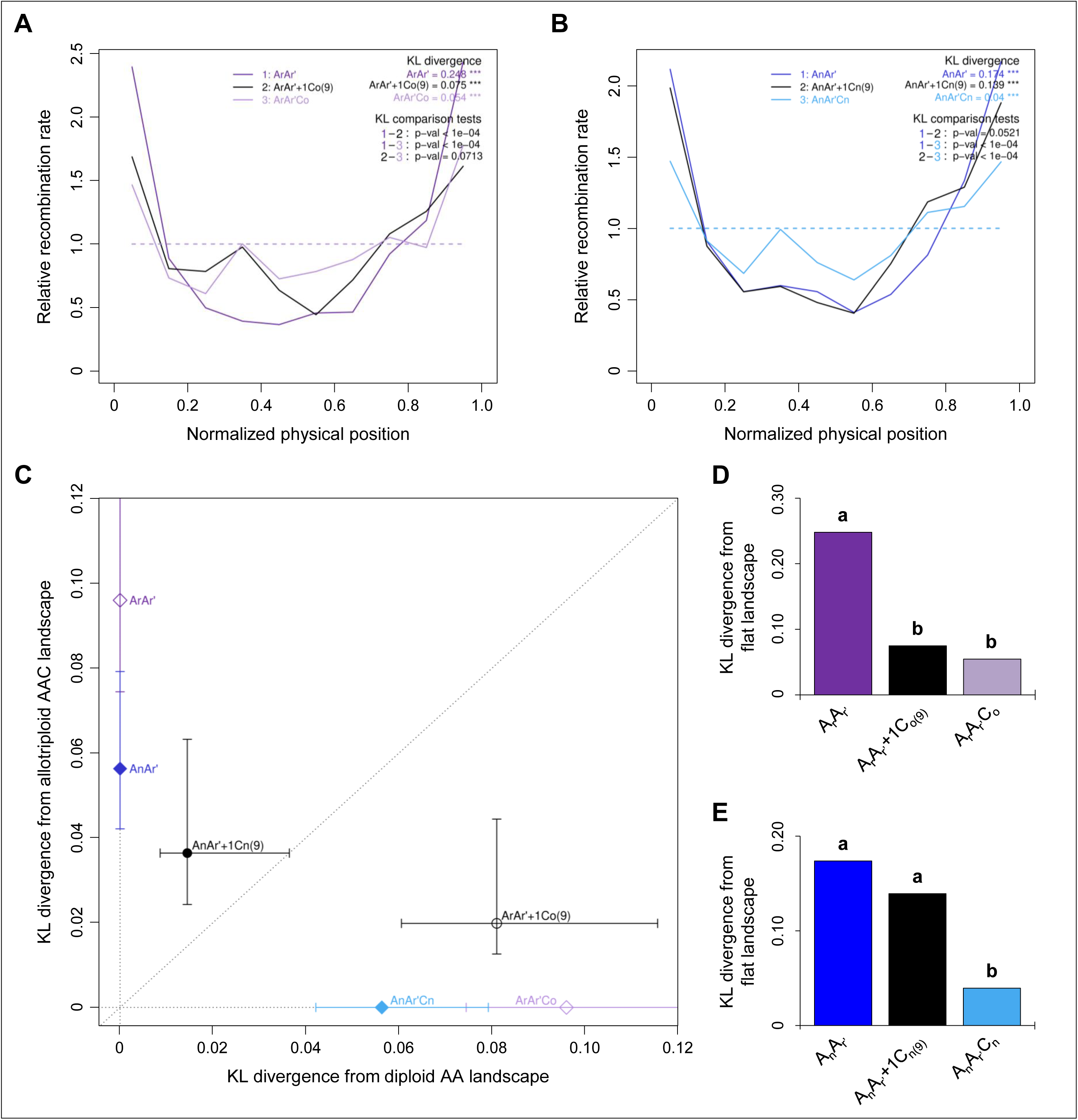
Differential contributions of the C09 chromosome from (A,C,D) *B. oleracea* and (B,C,E) *B. napus* to the reshaping of crossover landscapes between homologous A chromosomes in *Brassica* AAC allotriploids. The data represent the 10 A chromosomes pooled. **A-B**. Relative recombination rate across 10 bins along A chromosomes in hybrids carrying (**A**) *B. oleracea* C_o_ and (**B**) *B. napus* C_n_ chromosomes. KL divergence: divergence between the observed crossover landscape and the theoretical flat landscape (dashed line). *** denotes p-values<10^-3^ for testing H0: ‘flat landscape’ (10^4^ bootstraps). KL comparison tests: pairwise comparisons between above-mentioned KL values. p-values for testing H0: ‘KL values are identical’ (10^4^ bootstraps). **C**. Comparison of crossover landscape divergence between hybrids carrying the C09 chromosome and their respective diploid and allotriploid controls. Error bars represent 10^4^ bootstrap estimates. **D-E**. Values of divergence from the theoretical flat crossover landscape for the hybrids carrying (**D**) *B. oleracea* C_o_ and (**E**) *B. napus* C_n_ chromosomes. Statistical differences are indicated by letters (a, b), based on a Bonferroni-corrected test (10^4^ bootstraps, 5% threshold).

As established by Pelé *et al*. (2017a), patterns for the recombination landscapes between AA diploids and AAC allotriploids mostly differ regarding their correlation with the A genome architecture (genes and transposable elements densities, and the location of centromeres). Specifically, for the diploid A_r_A_r’_, the highest recombination rates arose mostly in the distal parts of the A chromosomes, enriched in genes and depleted in transposable elements, while the lowest always colocalized around the centromeric regions that show a reversed pattern for genes and transposable elements densities (Fig. 4). In contrast, crossover rates were more homogenous along every A chromosome in the allotriploid A_r_A_r’_C_o_ (Fig. 4). Statistically, these observations are verified by regression analyses (see Methods) showing that crossover rates tend to increase gradually from centromeres to chromosome extremities in the diploid A_r_A_r’_ (Fisher test, *p*<2.2E-16, R²=0.55; Fig. S5a) but not in the allotriploid A_r_A_r’_C_o_ (*p*=0.17). In the case of the A_r_A_r’_+1C_o(9)_ hybrid, this relationship was strongly reduced (*p*<2.2E-16, R²=0.19), indicating that crossover rates were less related to the A genome architecture, as observed for the allotriploid. At a local scale, the most marked differences with the diploid A_r_A_r’_ arose outside distal genomic regions, both in the A_r_A_r’_C_o_ and A_r_A_r’_+1C_o(9)_ hybrids (Fig. 4). By comparing the proportion of crossovers arising in the 159 given intervals sprinkling the A genome, 82 significant differences were detected between A_r_A_r’_ and A_r_A_r’_C_o_, and 53 between A_r_A_r’_ and A_r_A_r’_+1C_o(9)_ (Chi-squared test, *p*<0.05; Table S5), with 41 intervals in common between the two comparisons. In all cases, crossover frequencies were higher than in the diploid and strikingly, the genomic regions surrounding centromeres were concerned by these variations. Indeed, while the pericentromeric regions were totally deprived of crossovers in the diploid A_r_A_r’_ (∼10.6% of the A genome), this was not the case when hybrids carried either the additional *B. oleracea* C09 or all 9 C_o_ chromosomes (Fig. 4, Table S6). Specifically, only 2.9% and 0.3% of the A genome did not show any crossover formed from the segregating populations of the A_r_A_r’_+1C_o(9)_ and A_r_A_r’_C_o_ hybrids, respectively. Thus, we concluded that the modification of the distribution and formation of new recombining regions closer to the centromeres in AAC allotriploids mostly relies on the *B. oleracea* C09 chromosome.

**Fig. 4.**
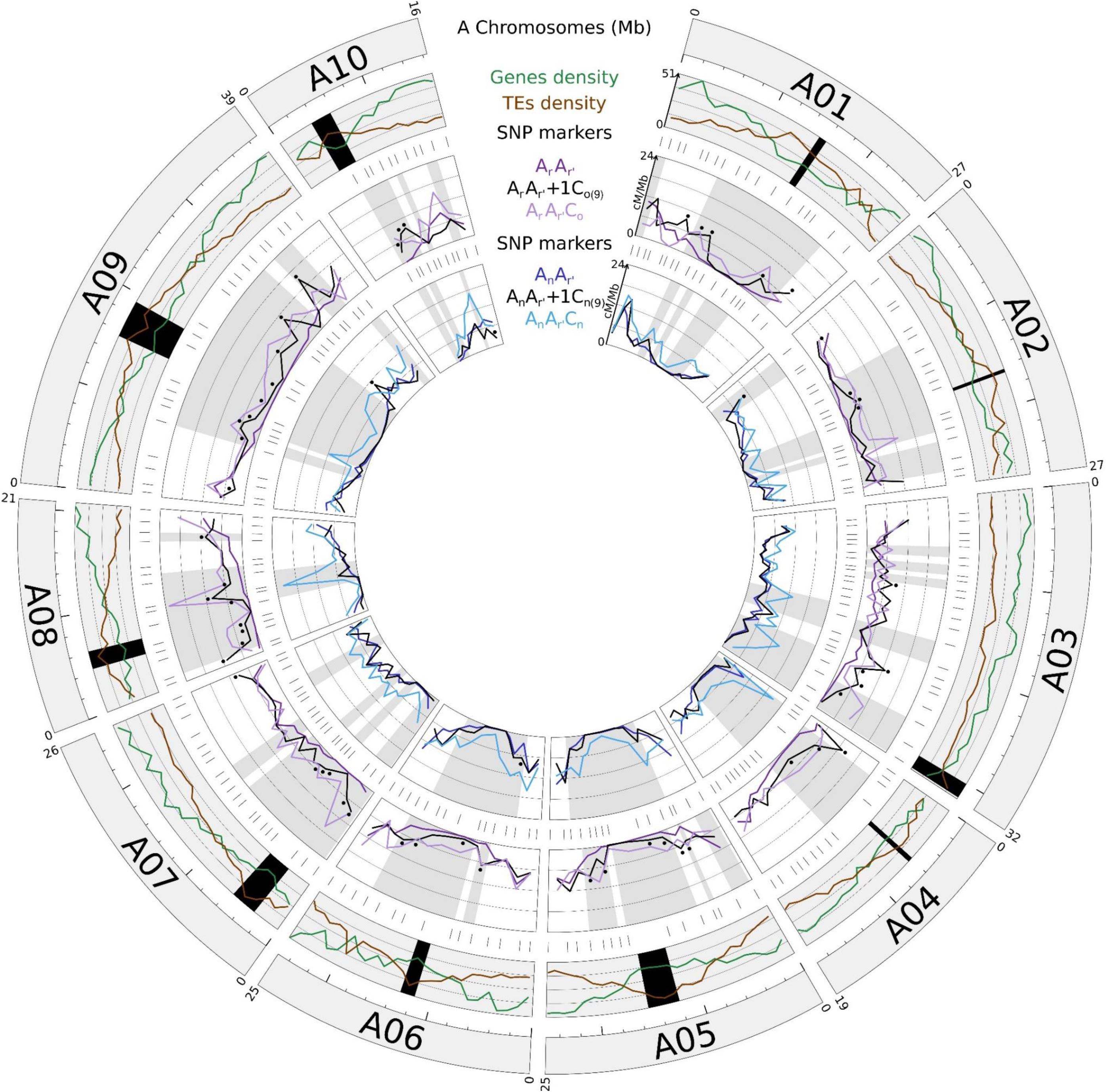
Circos diagram depicting the recombination rates along the 10 A chromosomes in cM per Mbp between the hybrids carrying *B. oleracea* C_o_ or *B. napus* C_n_ chromosomes. In the first outer circle are represented the 10 A chromosomes of the *B. rapa* cv. ‘Chiifu-401’ genome sequence (Wang *et al*., 2011). Their sizes are indicated by the values in megabase pairs above each chromosome, and a ruler is drawn underneath each chromosome, with larger and smaller tick marks every 10 and 2 Mbp, respectively. In the second outer circle is detailed the structure of each A chromosome: genes and transposable elements (TEs) densities from *B. rapa* cv. ‘Chiifu-401’ genome assembly (Wang *et al*., 2011). The active centromeres are delimited in black using the positions established by Mason *et al*. (2016). In the third and fifth outer circles are indicated the positions of the 169 SNP markers used for genotyping the progenies deriving from the hybrids carrying *B. oleracea* C_o_ or *B. napus* C_n_ chromosomes, respectively. In the fourth and sixth outer circles are represented the pairwise comparisons of the recombination landscapes (in cM per Mb) for progenies deriving from the hybrids carrying *B. oleracea* C_o_ or *B. napus* C_n_ chromosomes, respectively. For each interval between adjacent SNP markers, the heterogeneity of crossover rates was assessed using Chi-squared tests and significant differences at a threshold of 5% were indicated in grey for each pairwise comparison between diploids and allotriploids, and by the black stars when comparing diploids and hybrids carrying the additional C09 chromosome of either *B. oleracea* (4^th^ outer circle) or *B. napus* (6^th^ outer circle).

Theoretically, one may expect that increased crossover rates and even reshaped recombination landscapes follow from reduced interference. Thus, we quantified the degree to which the *B. oleracea* C09 chromosome contributed to the reduced crossover interference observed in AAC allotriploids (Pelé *et al*., 2017a). For that, we focused on the distribution of distances between adjacent crossovers and applied the KL measure of divergence between these distributions. We also compared to the non-interference case which is simulated by shuffling crossovers between individuals to remove any inter-crossover correlations (see Methods for details). As displayed in Fig. 5a with all A chromosomes pooled, we found that the distribution of distances between adjacent crossovers in the A_r_A_r’_+1C_o(9)_ showed more interference than the A_r_A_r’_C_o_ but less than the A_r_A_r’_. This is confirmed by the quantitative KL values of divergence to the non-interference distributions (KL of 0.233, 0.154 and 0.081 for A_r_A_r’_, A_r_A_r’_+1C_o(9)_ and A_r_A_r’_C_o_, respectively) and by the statistical ordering of these three genotypes (Fig. 5b). While the A_r_A_r’_ and A_r_A_r’_C_o_ hybrids belong to two separate groups, the A_r_A_r’_+1C_o(9)_ hybrid was not significantly different from either. Similar trends are observed per individual A chromosome, with less statistical significance (Figs. S6-S7 and Table S7). These results suggest that the addition of the *B. oleracea* C09 chromosome alone accounts for about half of the reduction in interference seen in the allotriploid.

**Fig. 5.**
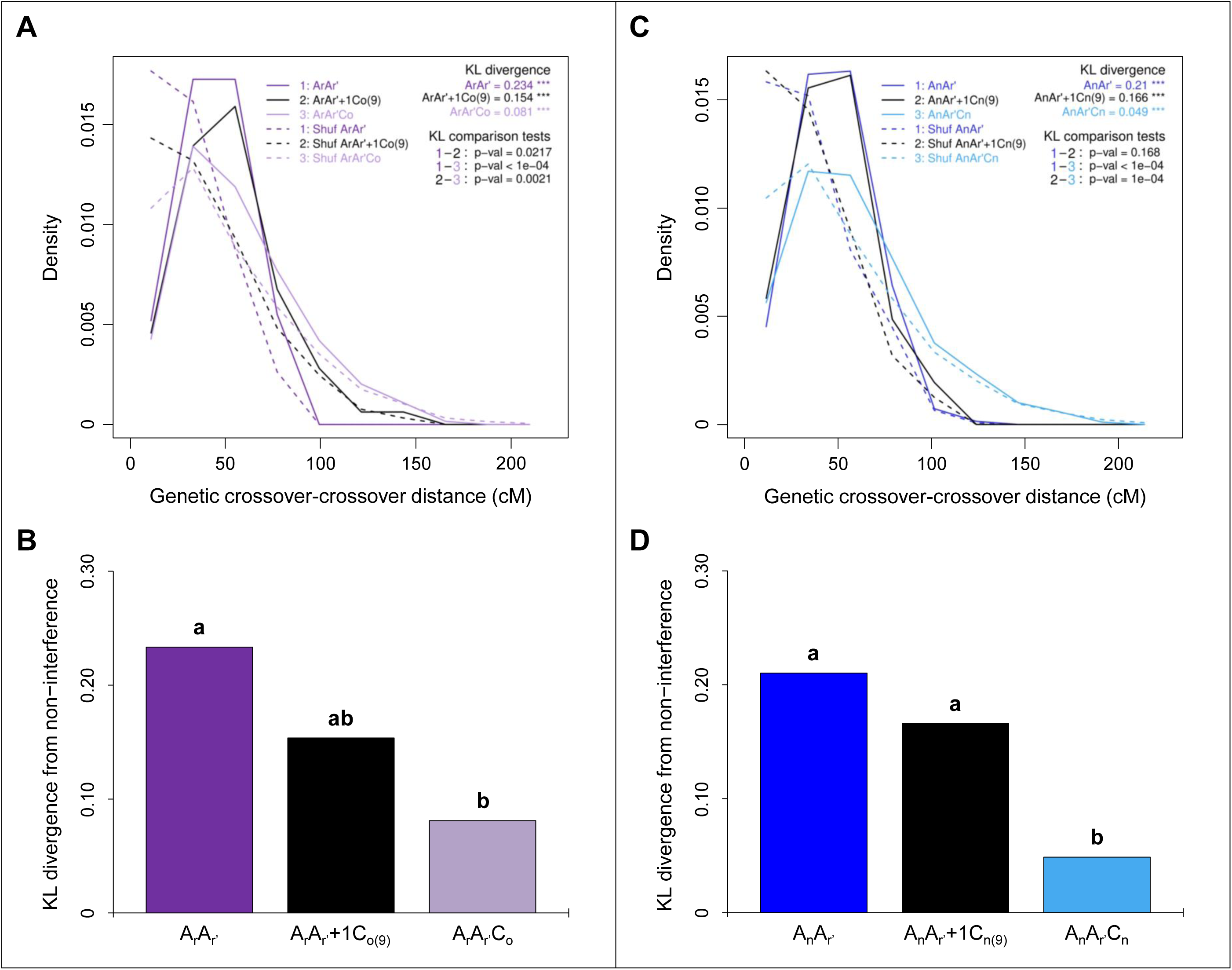
Differential contributions of the C09 chromosome from (A-B) *B. oleracea* and (C-D) *B. napus* to the reduction of crossover interference between homologous A chromosomes in *Brassica* AAC allotriploids. The data represented the 10 A chromosomes pooled. **A,C.** Density of distances between successive crossovers across 10 bins (relative to chromosome length) in hybrids carrying (**A**) *B. oleracea* C_o_ and (**C**) *B. napus* C_n_ chromosomes. KL divergence: divergence between the experimental distribution and the expected one in the absence of interference (obtained by shuffling crossovers across different individuals in the population; dashed line). *** indicates p-values<10^-3^ for testing H0: ‘no interference’ (10^4^ bootstraps). KL comparison tests: pairwise comparisons between above-mentioned KL values. p-values for testing H0: ‘KL values are identical’ (10^4^ bootstraps). **B,D.** Values of divergence from the non-interference scenario in hybrids carrying (**B**) *B. oleracea* C_o_ and (**D**) *B. napus* C_n_ chromosomes. Statistical differences are indicated with letters (a, b) based on a Bonferroni-corrected test (10^4^ bootstraps, 5% threshold).

### The B. napus C09 chromosome does not on its own contribute to the crossover variation in Brassica allotriploids

Given the higher crossover rate observed in AAC allotriploids harboring *B. napus* rather than *B. oleracea* C genome (Pelé *et al*., 2017a), we expect that the C09 chromosomes of *B. napus* contributes more significantly than its *B. oleracea* counterpart to the reshaping of crossovers in allotriploids. To test this hypothesis, we conducted a genotyping approach on segregating populations from the hybrids presented on Fig. 1b. In addition to the diploid A_n_A_r’_ (2*n*=20) and the allotriploid A_n_A_r’_C_n_ (2*n*=29) obtained by Pelé *et al*. (2017a), we generated multiple hybrids carrying combinations of C_n_ chromosomes including an A_n_A_r’_+1C_n(9)_ hybrid by exploiting the diploid A_n_A_n_ component extracted from *B. napus* cv. ‘Darmor’. These hybrids mostly differ by their content in additional C_n_ chromosomes, with the single C09 in A_n_A_r’_+1C_n(9)_ and the full haploid set (C01 to C09) in A_n_A_r’_C_n_ (Fig. 1c). In both cases, the additional C_n_ chromosomes always remain unpaired at Metaphase I in contrast to the A homologs that form 10 bivalents (see Fig. S8, Table S8 and Pelé *et al*. (2017a) for the diploid and allotriploid controls). From high throughput genotyping using the *Brassica* 60K Illumina^®^ Infinium SNP array, we determined that only the SNPs specific of the C09 chromosome amplified in the A_n_A_r’_+1C_n(9)_ hybrid (see Methods). This confirmed that the A_n_A_r’_+1C_n(9)_ plant displays the entire C09 chromosome of *B. napus* without any introgressions from other C_n_ chromosomes that could have originated from the crossing scheme. Crossover detection during male meiosis was performed from the genotyping of segregating populations using the same set of 169 SNPs previously described (excepted for six SNPs non polymorph for this set of hybrids that were substituted; Table S2). In total, 129 to 298 individuals obtained from each hybrid were genotyped, giving rise to a number of crossovers analyzed ranging from 1,063 to 2,618 per segregation population (Table 1).

In striking contrast to our observations using the hybrids with *B. oleracea* C_o_ chromosomes, we found that the crossover rate between homologous A chromosomes in AAC allotriploids is not at all explained by the presence of the C09 chromosome when it originates from *B. napus*. Indeed, the A_n_A_r’_ and A_n_A_r’_+1C_n(9)_ hybrids showed similar number of crossovers formed on average per meiosis, with 17.5 (SE=0.26, n=298) and 16.5 (SE=0.39, n=129) crossovers detected, respectively (Table 1). However, a 1.8-fold increase arose between homologous A genomes in the presence of the nine C_n_ chromosomes in the allotriploid A_n_A_r’_C_n_, which showed 32.3 crossovers on average (SE=0.54, n=162). Consistently, significant differences were only found for the A_n_A_r’_-A_n_A_r’_C_n_ and A_n_A_r’_+1C_n(9)_-A_n_A_r’_C_n_ comparisons when considering the total genetic lengths (Corrected Chi-squared test, *p*<2.2E-16; Fig. 2c). The same result holds at the scale of each A chromosome: we found no significant differences for the A_n_A_r’_-A_n_A_r’_+1C_n(9)_ comparisons (*p*<0.05; Fig. S1b) but significantly higher crossover rates in the allotriploid (*p*>0.05). On average, the number of crossovers was 1.6 to 2.2-fold higher in the allotriploid; the size of the A chromosomes driving these differences as deciphered from positive linear regressions (Fisher test, *p*<0.05), with R² values ranging from 0.64 to 0.85 depending on the hybrid (Fig. 2d). Moreover, greater frequencies of multiple crossovers formed per A chromosome were only detected in the allotriploid (Fig. S9).

Similarly, we showed that the single additional *B. napus* C09 chromosome does not even partly explain the changes of recombination landscapes observed between the A_n_A_r’_ and A_n_A_r’_C_n_ hybrids for the whole A genome nor for individual A chromosomes. Indeed, when comparing two distributions using the Kullback-Leibler (KL) divergence, we found that the recombination landscape of A_n_A_r’_+1C_n(9)_ was closer to that of A_n_A_r’_ than to that of A_n_A_r’_C_n,_ as shown qualitatively in Fig. 3b, when pooling the 10 A chromosomes. This is confirmed by the quantitative KL values of divergence to the flat landscape with 0.174, 0.139 and 0.040 for the A_n_A_r’_, A_n_A_r’_+1C_n(9)_ and A_n_A_r’_C_n_ hybrids, respectively. To further support this claim, Fig. 3c shows that the difference between landscapes is far smaller when comparing A_n_A_r’_+1C_n(9)_ to A_n_A_r’_ than when comparing it to A_n_A_r’_C_n_. Moreover, this conclusion is statistically supported when ordering these three genotypes with the A_n_A_r’_ and A_n_A_r’_+1C_n(9)_ hybrids belonging to the same group and both significantly different from the A_n_A_r’_C_n_ allotriploid (Fig. 3e). At the individual A chromosome level, similar trends were also found, although the statistical tests were less powerful (Figs. S3-4, Tables S3-S4).

As previously described, crossover landscape differences between diploids and allotriploids can be put in correspondence with the A genome architecture (Fig. 4). Indeed, while regression analyses revealed that crossover rates tend to increase gradually from centromeres to chromosome extremities in all hybrids (Fisher test, *p*<0.05; Fig. S5b), R² values vary extensively, with 0.54 for the diploid A_n_A_r’_ and 0.11 for the allotriploid A_n_A_r’_C_n_. In the case of the A_n_A_r’_+1C_n(9)_ hybrid, we found a R² value of 0.34 showing that the crossover distribution is less homogenous along the A chromosomes than in the allotriploids. Consistently, when comparing the proportion of crossovers arising in the 159 intervals sprinkling the A genome, the most marked differences between the diploid A_n_A_r’_ and the allotriploid A_n_A_r’_C_n_ were observed in pericentromeres and surrounding regions, with 58 significant differences (Chi-squared test, *p*<0.05; Fig. 4, Table S5-6). However, among these 58 intervals, only 2 (3.4%) showed significant differences between the A_n_A_r’_ and A_n_A_r’_+1C_n(9)_ hybrids, and both were located on chromosome extremities. Together, these findings show that the *B. napus* C09 chromosome alone cannot be held responsible for much of the modifications arising in AAC allotriploids, at the level of crossover landscapes or for the generation of new recombining regions located near the centromeres.

To test whether the *B. napus* C09 chromosome might be responsible for part of the reduced crossover interference observed in AAC allotriploids (Pelé *et al*., 2017a), we performed the same tests as were used for assessing the effect of the *B. oleracea* C09 chromosome. Specifically, we analyzed the distributions of the inter-crossover distances using the KL measure of divergence. As for the landscapes, the results for the pooled A chromosomes displays on Fig. 5c showed that the A_n_A_r’_ and A_n_A_r’_+1C_n(9)_ hybrids have very similar distributions whereas the A_n_A_r’_C_n_ allotriploid has a strongly reduced interference. This is confirmed by the quantitative KL values of divergence to the no-interference distributions (KL of 0.210, 0.166 and 0.049 for A_n_A_r’_, A_n_A_r’_+1C_n(9)_ and A_n_A_r’_C_n_ respectively) and with the statistical ordering of these three genotypes (Fig. 5d). Indeed, the A_n_A_r’_ and A_n_A_r’_+1C_n(9)_ hybrids belong to the same group while both differ significantly to the A_n_A_r’_C_n_ allotriploid. The same trends hold chromosome by chromosome and when comparing directly the KL divergences between these distributions of inter-crossover distances (Fig. S6-S7, Table S7). Together, these results suggest that the addition of the *B. napus* C09 chromosome alone does not contribute to the reduction in interference seen in the allotriploid.

To better understand the contrasted effect observed between the C09 chromosome of *B. oleracea* and *B. napus*, we investigated their content in meiotic genes. According to the list provided in Lloyd *et al*. (2014) and Higgins *et al*. (2021), we could identify 16 meiotic genes (Table S9). We did not find any difference between *B. oleracea* (cv. ‘RC34’) and *B. napus* (cv. ‘Darmor’) C09 chromosomes apart from the *RECQ4B* and *NPS1* gene copies that are missing in ‘RC34’. For these two latter genes, we extended our analysis to currently available assembled C genomes and observed that these gene copies were present in other *B. oleracea* genomes.

Given these results, we hypothesized that inter-chromosomal epistatic interactions arose at the time where C_n_ chromosomes originate from *B. napus*. This is consistent with our previous work in which we suggested that the C04 and C08 chromosomes of *B. oleracea* could also be involved in the crossover rate variations observed in AAC allotriploids (Suay *et al*., 2014). In addition to the A_n_A_r’_+1C_n(9)_ hybrid, we generated six other hybrids carrying a complete A_n_A_r’_ genome with the following additional C_n_ chromosomes of *B. napus* alone or in combinations: C04, C08 and C09 (Fig. 1b and 1c). After confirming the karyotype of each hybrid through BAC-FISH experiments (Fig. S8), and observing that C_n_ chromosomes always remain unpaired at Metaphase I of meiosis (Table S8), we detected male crossovers from the genotyping of segregating populations as previously described (Table 1). Briefly, our analysis revealed that, as for the A_n_A_r’_+1C_n(9)_ hybrid, the single addition of either the C04 chromosome in A_n_A_r’_+1C_n(4)_ or the C08 in A_n_A_r’_+1C_n(8)_ did not result in any significant change for crossover variation (Fig. 6). On the other hand, the occurrence of at least two of the C_n_ chromosomes tested always resulted in significant differences, with the most dramatic changes found when the C04, C08 and C09 chromosomes of *B. napus* are combined. These claims are particularly obvious when comparing the total genetic length for the whole A genome (Fig. 6a). Indeed, a significant increase in the crossover rate was only detected in hybrids carrying at least two additional C_n_ chromosomes, compared to the diploid A_n_A_r’_ control (Corrected Chi-squared test, *p*<0.05). The largest increase was found from the A_n_A_r’_+3C_n(4,8,9)_ hybrid that exhibits significantly more crossovers than any other hybrid tested, the allotriploid A_n_A_r’_C_n_ excepted, with 27.4 crossovers on average per meiocyte (SE=0.59, n=128; Table 1, Fig. 6a). Consistently with the hypothesis of inter-chromosomal epistatic interactions, the combination of the C04, C08 and C09 chromosomes of *B. napus* explained two thirds (66.7%) of the whole crossover rate variation between the diploid and allotriploid, which is significantly more than expected in case of additive effect of C_n_ chromosomes (Chi-squared test, *p*<0.05). Similarly, only the hybrids carrying at least two additional C_n_ chromosomes exhibited a recombination landscape much closer to that of the allotriploid A_n_A_r’_C_n_ than to the diploid A_n_A_r’_. This is illustrated in Fig 6b for the pooled 10 A chromosomes and is further supported quantitatively by the KL values of divergence to the flat landscape (Fig. S3, Table S3) and statistically through genotype ordering (Fig. S4). Interestingly, we still observed a significant difference between the A_n_A_r’_+2C_n(4,9)_ hybrid and the allotriploid control, suggesting that the inter-chromosomal epistatic interactions between the *B. napus* C04 and C09 chromosomes plays a less pivotal role in the variation of the recombination landscapes. Furthermore, when comparing the proportion of crossovers arising in the 159 intervals sprinkling the A genome, the number of intervals showing significant differences with the allotriploid was reduced when hybrids carried at least two additional C_n_ chromosomes (Table S5-S6). For instance, only six intervals showed significant differences between the A_n_A_r’_+3C_n(4,8,9)_ hybrid and the allotriploid, and none of them was located in or near pericentromeric regions (Chi-squared test, *p*<0.05). Finally, the intensity of interference followed similar trends when measured from the pooled distribution of distances between adjacent crossovers. Indeed, only the hybrids carrying at least two additional C_n_ chromosomes showed an interference level closer to that of the allotriploid A_n_A_r’_C_n_, rather than to that of the diploid A_n_A_r’_ (Fig. 6c). This is confirmed by the quantitative KL values of divergence to the non-interference distributions (Fig. S6, Table S7) and with the statistical ordering of the genotypes (Fig. 6c). However, the level of interference did not differ significantly between the A_n_A_r’_+2C_n(4,8)_ hybrid and the diploid control, suggesting that the inter-chromosomal epistatic interactions between the *B. napus* C04 and C08 chromosomes plays a less decisive role in this variation. Despite less powerful analysis, we found similar trends per individual A chromosome for crossover rate, distribution and interference variations, as can be seen in Figs. S3-4, S6-7, S10 and Tables S3, S4 and S7. Together, these results suggest that variation in crossover number and patterning arising in *Brassica* allotriploids mostly results from inter-chromosomal epistatic interactions between the C04, C08 and C09 chromosomes when they derive from *B. napus*.

**Fig. 6.**
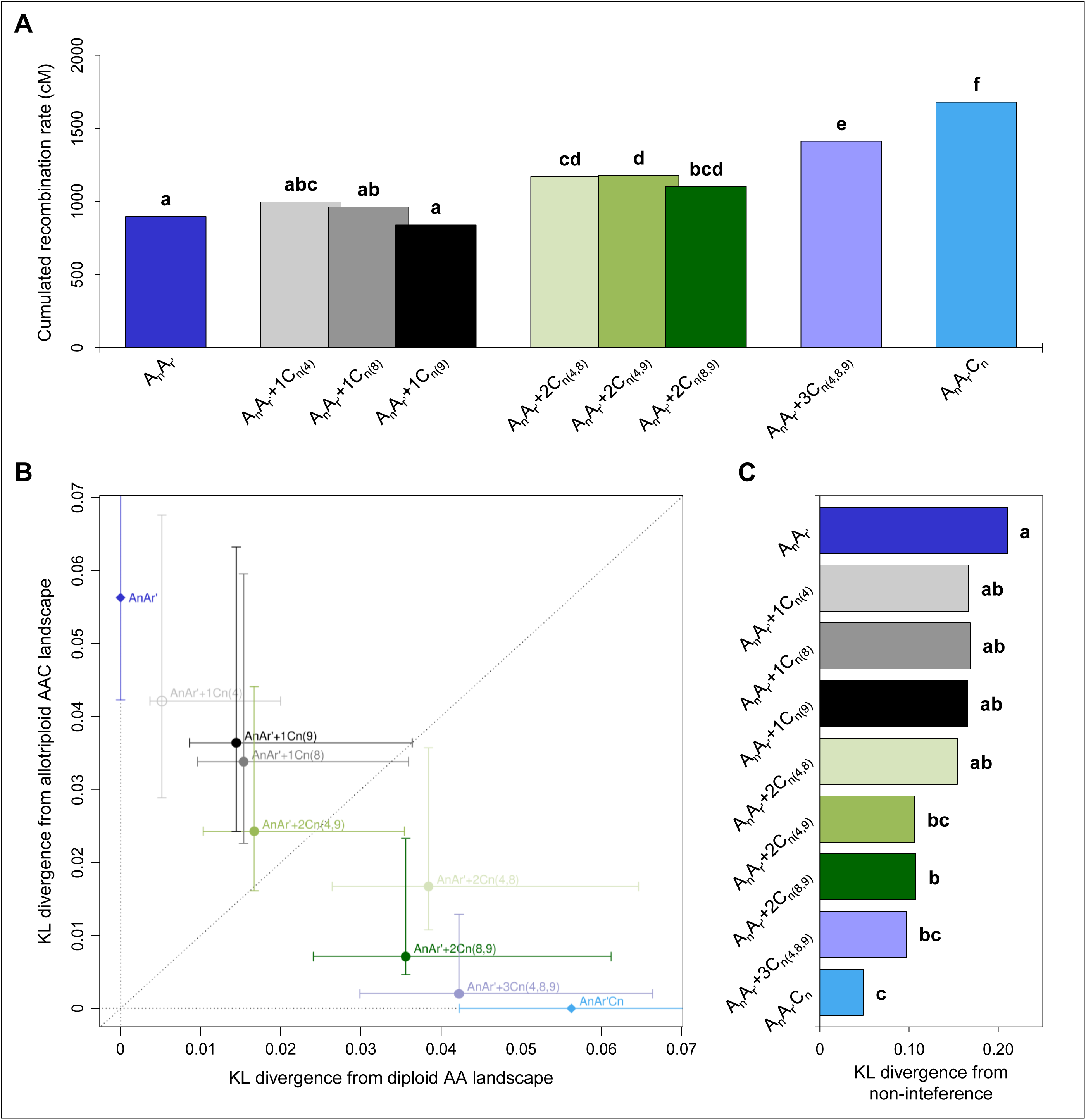
Individual and combined effects of the C04, C08 and C09 chromosomes from *B. napus* on the crossover variations between homologous A chromosomes in *Brassica* AAC allotriploids. The data represent the 10 A chromosomes pooled. **A.** Recombination rates in Centimorgan (cM) for the cumulated A chromosomes. Statistical differences based on a Bonferroni-corrected Chi-squared test (5% threshold) are indicated with letters (a–f). **B.** Comparison of crossover landscape divergence between hybrids carrying additional C_n_ chromosomes from *B. napus* with their diploid and allotriploid controls. Error bars represent 10^4^ bootstraps estimates. **C.** Values of divergence from the non-interference scenario. Statistical differences based on a Bonferroni-corrected test (10^4^ bootstraps, 5% threshold) are indicated with letters (a–c).

To gain a deeper understanding of these potential inter-chromosomal epistatic interactions, we also examined the meiotic gene content of the *B. napus* cv. ‘Darmor’ C04 and C08 chromosomes and identified 13 and 10 meiotic genes, respectively (Table S10). Interestingly, we found copies of meiotic genes known as anti-crossover factors, along with their associated co-factors, which naturally limit Class II crossovers in *A. thaliana* (Crismani *et al*., 2012; Girard *et al*., 2014; 2015; Séguéla-Arnaud *et al*., 2015; 2017; Fernandes *et al*., 2018a; Singh *et al*., 2023). Notably, we detected a copy of *FLIP* on chromosome C08 and *RMI1* on chromosome C09, both of which are co-factors of the anti-crossover factors *FIGL1* and *RECQ4A/B*, respectively (Séguéla-Arnaud *et al*., 2017; Fernandes *et al*., 2018a). Furthermore, copies of *RECQ4A* and *RECQ4B* were found on the *B. napus* C08 and C09 chromosomes, respectively.

## Discussion

### The formation of supernumerary crossovers closer to the centromeres in Brassica allotriploids is genetically regulated

In this study, we demonstrated that the crossover variations occurring between homologous A genomes in *Brassica* AAC allotriploids relate to specific additional C chromosomes. Notably, we identified that the C09 chromosome of *B. oleracea* acts as a major determinant in the reshaping of crossover landscapes. Indeed, the occurrence of this single haploid chromosome is sufficient to promote the formation of crossovers in normally cold recombining regions just as occurs in allotriploids. However, this single additional chromosome does not fully explain the changes in crossover rate and interference (∼50%), suggesting the involvement of other C chromosomes. This hypothesis is supported by our analysis of hybrids carrying the C04, C08 and C09 chromosomes of *B. napus* alone or in combinations. Indeed, while no effect on homologous recombination was observed when a single C_n_ chromosome was present, significant changes always arose in hybrids that combine at least two of these chromosomes. Even more striking, the A_n_A_r’_+3C_n(4,8,9)_ hybrid showed very similar crossover variations that what occurs in the allotriploids, suggesting inter-chromosomal epistatic interactions between the C04, C08 and C09 chromosomes of *B. napus*. Therefore, our results confirm the consequences of specific C chromosomes, either for their direct effect when deriving from *B. oleracea* or possible indirect effects in the case of *B. napus*. This strongly suggests that crossover variations occurring in *Brassica* allotriploids are genetically controlled, most presumably by dosage sensitive genes as no change in homologous recombination is observed between diploid AA and allotetraploid AACC hybrids (Boideau *et al*., 2021).

To our knowledge, only four meiotic genes have been found to date to influence crossover formation in a dosage-sensitive manner in plants: *HEI10* and *ASY1* in *A. thaliana* (Ziolkowski *et al*., 2017; Lambing *et al*., 2019), and *MSH4* and *ASY3* in *B. napus* (Gonzalo *et al*., 2019; Chu *et al*., 2024). While null mutations of these genes always result in a dramatic decrease in crossover numbers, the removal and/or addition of copies leads to intriguing variations regarding our observations in *Brassica* AAC allotriploids. For example, the *HEI10* gene (*i.e*., *ZIP3* in yeast), which encodes a conserved ubiquitin E3 ligase belonging to the ZMM complex, has been shown to cause reduced recombination frequency in heterozygous mutants and an elevated crossover rate in Arabidopsis plants carrying additional copies (Chelysheva *et al*., 2012; Serra *et al*., 2017; Ziolkowski *et al*., 2017; Durand *et al*., 2022). In contrast, heterozygous mutants for the axial element *ASY1* exhibited a redistribution of crossovers toward the chromosome extremities (Lambing *et al*., 2019). In *B. napus*, when a single copy of the other axial element *ASY3* remained functional (out of the four copies present in the allotetraploid), a significant increase in crossover number was detected, potentially associated with reduced interference intensity (Chu *et al*., 2024). Finally, in *Brassica* AC amphihaploids, the knockout of one functional copy of *MSH4*, another gene encoding a key protein of the ZMM complex, resulted in a strong reduction of meiotic crossovers between non-homologous chromosomes (Higgins *et al*., 2004; Gonzalo *et al*., 2019). However, we did not detect any copies of these meiotic genes on the *B. oleracea* C09 chromosome, which drives most of the crossover changes in allotriploids (Table S9). Interestingly, our analysis identified one copy of *ASY3* on the *B. napus* C04 chromosome (Table S10). Nevertheless, it is highly unlikely that this gene contributes directly to the crossover variations observed in *Brassica* allotriploid, as the single addition of the *B. napus* C04 chromosome in the A_n_A_r’_+1C_n(4)_ hybrid was not associated with any changes in crossover rate, distribution or interference. Except for these cases, our knowledge remains limited regarding dosage-sensitive genes influencing crossovers formation. The development of polyploids carrying mutations in meiotic genes, as recently achieved in autotetraploids of Arabidopsis (Parra-Nunez *et al*., 2024), will facilitate in the future the identification of candidate causal genes in *Brassica* allotriploids.

As previously mentioned, we highlighted potential inter-chromosomal epistatic interactions between the C04, C08, and C09 chromosomes of *B. napus* in the crossover changes occurring in *Brassica* allotriploids. To our knowledge, a few number of epistatic interactions between meiotic genes influencing crossover variation have been well-characterized in *A. thaliana*. For instance, the heat shock protein HSBP and the co-chaperone J3, encoded respectively by *HCR2* and *HCR3* genes, were both identified though a forward genetic screens as co-factors of the *HEI10* ubiquitin E3 ligase in *A. thaliana* (Kim *et al*., 2022; 2024). While HSBP represses *HEI10* expression (Kim *et al*., 2022), the J3 co-chaperone promotes the proteasomal degradation of HEI10 (Kim *et al*., 2024). On the other hand, multiple proteins have been shown to limit Class II crossover formation in *A. thaliana*, in conjunction with the anti-crossover factors *FANCM*, *RECQ4*, *FIGL1* and *FANCC* (Crismani *et al*., 2012; Girard *et al*., 2014; 2015; Séguéla-Arnaud *et al*., 2015; 2017; Fernandes *et al*., 2018a; Singh *et al*., 2023). Notably, we can mention the complexes FANCM-MHF1-MHF2 (Girard *et al*., 2014), RECQ4-RMI1-TOP3a (Séguéla-Arnaud *et al*., 2017), FIGL1-FLIP (Fernandes *et al*., 2018a) and lately FANCC-FANCE-FANCF (Singh *et al*., 2023). In our analysis, we detected a total of 39 meiotic genes located on the C04, C08 and C09 *B. napus* chromosomes (Table S9-10). Interestingly, we identified a copy of *FLIP* and *RECQ4A* on chromosome C08 as well as *RMI1* and *RECQ4B* on chromosome C09. Whether epistatic interactions between these genes could account for the inter-chromosomal epistatic interactions observed in this study remains unknown. However, this seems unlikely, as *FLIG1* and *RECQ4* have been primarily linked to Class II crossover variations, whereas the supernumerary crossovers observed during the male meiosis of *Brassica* AAC allotriploids were shown to originate from the Class I crossover pathway (Leflon *et al*., 2010; Pelé *et al*., 2017a). Furthermore, none of the genes discussed thus far have been associated with an increased crossover rate closer to the centromeres, as found in *Brassica* allotriploids (Jing *et al*. unpublished data).

As for the causal genes, the mechanisms involved in the modifications of homologous recombination in AAC allotriploids are not yet known, especially regarding the changes in crossover landscapes. Given that A genotypes were identical or very similar between the hybrids of each type, a reasonable assumption is that specific additional C chromosomes could affect the epigenetic features of homologous A chromosomes, as extensively discussed by Pelé *et al*. (2017a). Briefly, this hypothesis was motivated by two elements. First, following a polyploidization event, modifications for DNA methylation have been observed in resynthesized allopolyploids (Edger *et al*., 2017), including *B. napus* (Lukens *et al*., 2006; Gaeta *et al*., 2007; Książczyk *et al*., 2011; Martinez Palacios *et al*., 2019). Second, several publications pointed-out that the epigenetic features of DNA regulate the formation of crossovers, notably in pericentromeric regions, as well as interference (Colomé-Tatché *et al*., 2012; Melamed-Bessudo & Levy, 2012; Mirouze *et al*., 2012; Yelina *et al*., 2012; Choi *et al*., 2013; Jahns *et al*., 2014; Shilo *et al*., 2015; Yelina *et al*., 2015a,b; Choi *et al*., 2018; Naish *et al*., 2021; Fernandes *et al*., 2024). This hypothesis became strongly realistic with the works of Underwood *et al*. (2018). Indeed, these authors showed that disruption of H3K9me2 and non-CG DNA methylation pathways increased pericentromeric crossovers in *A. thaliana* based on the mutation of *KYP*, *SUVH5* and *SUVH6* genes. However, no copy of these genes was detected on the C09 chromosome of *B. oleracea* cv. ‘RC34’ and *B. napus* cv. ‘Darmor’.

### Regulation of homologous recombination in Brassica allopolyploids may have changed following genomic divergence

The most striking result of this study stems from the comparison of the direct effect of the C09 chromosome deriving from *B. oleracea* and *B. napus*. Indeed, while the single addition of the C09 chromosome of *B. oleracea* explains most of the modifications taking place between pairs of A homologs in *Brassica* AAC allotriploids, no variation was detected when it derives from an established lineage of *B. napus*. To explain such differences, one attractive hypothesis could relate to the genomic divergence undergone by the C genome of *B. napus*, compared to its *B. oleracea* diploid progenitor, following the polyploidization event that occurred ∼7,500 years ago (Chalhoub *et al*., 2014). Indeed, allopolyploidy is well recognized as a major driving force in plants speciation, evolution, and adaptation (Comai, 2005; Leitch and Leitch, 2008; Soltis and Soltis, 2012; Wendel, 2015; Alix *et al*., 2017; Qiu *et al*., 2020). Allopolyploid formation is generally accompanied by a ‘genome shock’ and “transcriptome shock” resulting in genomic restructuring and novel opportunities to generate functional diversification between related genes. Duplicated genes may thus be subjected to physical loss (non-reciprocal exchanges, genic conversion, and fractionation), functional loss (pseudogenization), and/or functional diversification (sub- and neofunctionalization) (Ohno, 1970; Wendel, 2000; Adams, 2007; Woodhouse *et al*., 2010; Sankoff *et al*., 2012; Tayale and Parisod, 2013; Yoo *et al*., 2014). Consistently, such modifications were extensively reported from the study of resynthesized and naturally occurring allotetraploids of *B. napus* (Song *et al*., 1995; Albertin *et al*., 2006; Gaeta *et al*., 2007; Xu *et al*., 2009; Gaeta & Pires, 2010; Marmagne *et al*., 2010; Xiong *et al*., 2011; Rousseau-Gueutin *et al*., 2016; Ferreira de Carvalho *et al*., 2021). Furthermore, meiotic genes were found to return to a single copy more rapidly than the genome-wide average in angiosperms after polyploidization events (De Smet *et al*., 2013; Lloyd *et al*., 2014). Last but not least, the role of genomic divergence in regulating homeologous recombination in allopolyploids is well established. Indeed, while associations between homeologous chromosomes are commonly observed in newly formed allopolyploids, they occur infrequently in established lineages, thereby facilitating the production of balanced gametes (Lloyd and Bomblies, 2016; Pelé *et al*., 2018). This is primarily attributed to the set-up of genetic controls preventing homeologous crossovers in established lineages (Jenczewski and Alix, 2004). The most studied example corresponds to the *Ph1* locus in the allohexaploid bread wheat, a complex cluster of defective cyclin dependent kinases-like (CDK) and methyl transferase genes, which includes an inserted paralog of the *ZIP4* gene (Sears, 1976; Griffiths et al., 2006; Knight et al., 2010; Greer et al., 2012; Martín et al., 2014, Rey *et al*., 2017). Although the causal gene remains unknown, a major quantitative trait locus, *BnaPh1*, controlling homoeologous recombination in *Brassica napus*, has recently been mapped to chromosome A09 (Higgins *et al*., 2021). In this study, we did not detect any physical loss of meiotic genes located on the C09 chromosome in *B. napus* ‘Darmor’ when compared to *B. oleracea* (Table S9). Interestingly, we noticed that in *B. oleracea* cv. ‘RC34’, the copies of *RECQ4B* and *MPSI* genes on the C09 chromosome seem pseudogenized compared to *B. napus* and other *B. oleracea* cultivars. To our knowledge, only the mutation of *RECQ4* has been associated with crossover rate variations in Arabidopsis and some crops, but exclusively on chromosome extremities (Séguéla-Arnaud *et al*., 2015; Fernandes *et al*., 2018b; Mieulet *et al*., 2018; de Maagd *et al*., 2020). Together, this suggests that the contrasted effect of the C09 chromosome on crossover variations may result from functional loss or diversification of meiotic genes rather than physical loss in *B. napus*.

Nevertheless, we cannot totally exclude that the contrasted effect of the C09 chromosome is due to allelic variations in loci regulating crossover formation. Indeed, due to the complexity of obtaining our plant material (Pelé *et al*., 2017b), we only used one cultivar of *B. oleracea* and *B. napus*. However, extensive variations are observed for crossover rate and landscapes between and within species (Coop and Przeworski, 2007; Smukowski and Noor, 2011; Mercier *et al*., 2015; Stapley *et al*., 2017; Haenel *et al*., 2018; Brazier and Glémin, 2022), as exemplified between different combinations of crosses in *B. oleracea* (Cai *et al*., 2023) or between homeologous chromosomes in resynthesized *B. napus* generated from different diploid cultivars (Ferreira de Carvalho *et al*., 2021; Katche *et al*., 2023). Recently, genome-wide crossover differences between *A. thaliana* accessions were associated with allelic variation in three genes: *HEI10*, *TAF4B*, and *SNI1* (Ziółkowski *et al*., 2017; Lawrence *et al*., 2019; Zhu *et al*., 2021). However, the most dramatic changes were always found in subtelomeric regions. In *Brassica napus*, a similar locus has been identified on the C09 chromosome (Jenczewski *et al*., 2003; Liu *et al*., 2006). Specifically, *PrBn* regulates non-homologous crossover variation between amphihaploid AC plants obtained from different cultivars of *B. napus*. Moreover, it was suggested that this genetic control might be responsible for the recombination rate differences observed between homologous A chromosomes of AAC hybrids produced from different *B. napus* varieties (Nicolas *et al*., 2009). Whether allelic variation in *PrBn* is responsible of the contrasted effect of the C09 chromosome of *B. oleracea* ‘RC34’ and *B. napus* ‘Darmor’ is not yet known, but this locus was not associated with changes in crossover distribution (Nicolas *et al*., 2009; Nicolas *et al*., 2012). In our analysis, we did not identify copies of *HEI10*, *TAF4B*, and *SNI1* on the C09 chromosomes of *B. oleracea* and *B. napus*. Furthermore, when comparing the orthologous copies of meiotic genes between the cultivars ‘Darmor’ and ‘RC34’ on C09, we observed that they were very conserved, both in length and sequence identity (ranging from 95.3% to 100%; Table S9). To definitively rule out the possibility of causal allelic variation, important efforts would be necessary to develop the same plant material as presented in this study, but with C chromosomes originating from other *B. oleracea* and *B. napus* cultivars.

In conclusion, this study highlights the role of specific chromosomes and genomic divergence in modifying the tightly regulated homologous recombination pattern in *Brassica* allopolyploids. Although drawing conclusions about the candidate causal genes is challenging at this stage, the identification of a contrasted effect between the *B. oleracea* and *B. napus* C09 chromosome paves the way for developing segregating populations. This will enable the mapping of involved loci and, ultimately, the identification of causal genes, as currently being done in our group. This study will be crucial in the future for enhancing the genetic diversity in *Brassica* breeding programs. Moreover, it may help extend this knowledge to other crop species, such as bread wheat, which has shown similar homologous recombination changes in pentaploids (Yang *et al*., 2022).

## Acknowledgements

We acknowledge the Genetic Resource Center (BrACySol, UMR IGEPP, Ploudaniel, France) for providing seeds and the staff for their technical assistance in greenhouses (especially L. Charlon, P. Rolland, J.P. Constantin, J.M. Lucas and F. Letertre). We also thank the UMR INRA 1095 GENTYANE platform (Clermont-Ferrand, France, http://gentyane.clermont.inra.fr/) and the UMR 8199 genotyping service (Lille, France, http://wwwgood.ibl.fr/index.fr/index.php/services-prestations) for the generation of genotyping data. We warmly thank the plant molecular cytogenetic platform (INRAE, Biogenouest, Le Rheu, France, https://www6.rennes.inrae.fr/igepp_eng/About-IGEPP/Platforms/Molecular-Cytogenetics-Platform-PCMV) for helping and participating in the cytogenetic experiments of this study. Alexandre Pelé was supported by a fellowship from BAP INRA and Brittany region. This work was funded by BAP INRA department and ANR CROC: Project ANR-14-CE19-0004. We also thank Dr. Franz Boideau and Prof. Piotr A. Ziolkowski, both from the Laboratory of Genome Biology of Adam Mickiewicz University (Poznan, Poland), for their critical reading of the manuscript and advice, respectively.

## Author contributions

This study was conducted by A.P., M.R.-G., A.-M.C. for the design of the research, A.P., M.L-T. for the plant production, A.P., O.C., V.H. for the cytogenetic analyses, A.P., M.L-T., M.R.-G. for the molecular analyses, A.P., M.F., O.C.M., J.M., M.R.-G., A.-M.C. for data analyses. A.P. wrote the first draft and A.P., M.F., O.C.M., M.R.-G., A.-M.C. reviewed and edited the manuscript.

## Supporting Information

**Fig. S1 Recombination rates in Centimorgan (cM) for each of the 10 homologous A chromosomes in hybrids carrying (A) *B. oleracea* C_o_ or (B) *B. napus* C_n_ chromosomes.** Statistical differences, determined by a Bonferroni-corrected Chi-squared test (5% threshold), are indicated with letters (a–c).

**Fig. S2 Frequency of crossovers per chromatid for the progenies derived from hybrids carrying *B. oleracea* C_o_ chromosomes.**

**Fig. S3 Relative recombination rate across 10 bins along each individual A chromosome and for the 10 A chromosomes pooled in each hybrid.** KL divergence: divergence between the observed crossover landscape and the theoretical flat landscape (dashed line). *** indicates p-values<10^-3^ for testing H0: ‘flat landscape’ (10^4^ bootstraps). KL comparison tests: pairwise comparisons between above-mentioned KL values. p-values for testing H0: ‘KL values are identical’ are obtained by 10^4^ bootstraps.

**Fig. S4 Landscape flatness for each individual A chromosome and for the 10 A chromosomes pooled between the hybrids carrying *B. oleracea* C_o_ or *B. napus* C_n_ chromosomes.** Flatness is proxied by the KL divergence between the observed crossover landscape (distribution across 10 bins) and the corresponding theoretical flat distribution.

**Fig. S5 Relationship between the relative recombination rates (normalized per A chromosome, %) and their relative distance from the centromeres (%) in hybrids carrying (A) *B. oleracea* C_o_ or (B) *B. napus* C_n_ chromosomes.** A_r_A_r’_ (purple circles): y=0,1637x-1,4544; R²=0.55. A_r_A_r’_+1C_o(9)_ (black diamonds): y=0,0630x+3,3110; R²=0.19. A_r_A_r’_C_o_ (light purple squares): y=0,01348x+5,6515; R²=n.s.. A_n_A_r’_ (blue circles): y=0,1387x-0,2705; R²=0.54. A_n_A_r’_+1C_n(9)_ (black diamonds): y=0,1086x+1,1521; R²=0.34. A_n_A_r’_C_n_ (light blue squares): y=0,0446x+4,1816; R²=0.11.

**Fig. S6 Density of distances between successive crossovers across 10 bins (relative to chromosome length) for each individual A chromosome and for the 10 A chromosomes pooled in each hybrid.** KL divergence: divergence between the experimental distribution and the expected one in the absence of interference (obtained by shuffling crossovers across different individuals in the population; dashed line). *** indicates p-values<10^-3^ for testing H0: ‘no interference’ (10^4^ bootstraps). KL comparison tests: pairwise comparisons between above-mentioned KL values. p-values for testing H0: ‘KL values are identical’ are obtained by 10^4^ bootstraps.

**Fig. S7 Divergence from the non-interference situation for each individual A chromosome and for the 10 A chromosomes pooled between hybrids carrying *B. oleracea* C_o_ or *B. napus* C_n_ chromosomes.** Values of KL divergence between the observed distribution of inter-crossover distances across 10 bins and the corresponding distribution without interference (obtained by shuffling the data). Statistical differences, determined by a Bonferroni-corrected test using 10^4^ bootstraps (5% threshold), are indicated with letters (a, b, c).

**Fig. S8 Meiotic observations of hybrids carrying a diploid A_n_A_r’_ genome and the additional chromosomes C04, C08 and C09 of *B. napus* single or in combination.** (**1-7.A**) Pollen Mother Cells showing ten bivalents as well as one (**1-3.A**), two (**4-6.A**) or three (**7.A**) univalents indicated by the red stars. FISH analyses carried out with the BAC Bob014O06, revealing the C chromosomes unpaired (**1-7.B**, in green), the BAC KBrH033J07 showing the A05 and C04 homoeologous pairs (**1-7.C**, in red), the BAC KBrB043F18 showing the A09 and C08 homoeologous pairs (**1-7.D**, in purple), and the BAC KBrH080A08 showing the A10 and C09 homoeologous pairs (**1-7.E**, in orange). Bars, 5 µm.

**Fig. S9 Frequency of crossovers per chromatid for the progenies derived from hybrids carrying *B. napus* C_n_ chromosomes.**

**Fig. S10 Recombination rates in Centimorgan (cM) for each of the 10 homologous A chromosomes in hybrids carrying additional *B. napus* C_n_ chromosomes.**

**Table S1. Characteristics of SSR markers used to study the segregation of *B. napus* C_n_ chromosomes.**

**Table S2. Characteristics of SNP markers genetically mapped on the 10 homologous A chromosomes of the *B. rapa* cv. ‘Chiifu-401’ genome sequence (Wang *et al*., 2011), used for hybrids carrying *B. oleracea* C_o_ and *B. napus* C_n_ chromosomes.**

**Table S3. Landscape flatness analysis.** Detail of the KL values and p-values corresponding to the results presented in Figures 3 and Supplementary Figures S3 and S4. The numbers 1, 2, 3 indicate the hybrids of the columns Pop_1, Pop_2 and Pop_3, respectively. The line "Pool" in the "Chromosome" column corresponds to the pooled analysis of the 10 A chromosomes together. Column details: ‘Pop_n’ refers to the name of the hybrid n, ‘KL_n’ gives the KL divergence value between the observed distribution of crossovers for the hybrid n, and the corresponding flat histogram (H0 hypothesis), ‘pvalH0_n’ corresponds to that H0 hypothesis, and ‘pvalDiffKL_n-p’ refers to the p-value of the hypothesis “KL_n - KL_p=0”.

**Table S4. Landscapes comparison analysis.** Detail of the KL values and p-values corresponding to the results presented in Figures 3C and 6B. The numbers 1, 2, 3 indicate the hybrids of the columns Pop_1, Pop_2 and Pop_3, respectively. The line "Pool" in the "Chromosome" column corresponds to the pooled analysis of the 10 A chromosomes together. Column details: ‘Pop_n’ refers to the name of the hybrid n. ‘KL_n-p’ gives the values of the KL divergences between observed distributions of crossovers in hybrids of Pop_n and Pop_p. ‘Inf_n-p’ and ‘Sup_n-p’ indicate the corresponding confidence intervals at α=5% (2.5% on each side, from 10^4^ bootstraps).

**Table S5. Chi-squared comparisons between hybrids carrying *B. oleracea* C_o_ or *B. napus* C_n_ chromosomes for crossover rate heterogeneity per interval between adjacent linked SNP markers.**

**Table S6. Genetic distances in centimorgan (cM) between linked SNP markers for each hybrid.** Intervals including pericentromeric regions are indicated in grey.

**Table S7. Interference analysis.** Detail of the KL values and p-values corresponding to the results presented in Figure 5 and Supplementary Figures S6 and S7. The numbers 1, 2, 3 indicate the hybrids of the columns Pop_1, Pop_2 and Pop_3, respectively. The line "Pool" in the "Chromosome" column corresponds to the pooled analysis of the 10 A chromosomes together. Column details: ‘Pop_n’ refers to the name of the hybrid, ‘KL_n’ gives the value of the KL divergence between the observed distribution of distances between successive crossovers for the hybrid n, and the corresponding distribution without interference obtained by shuffling the data (H0 hypothesis), ‘pvalH0_n’ corresponds to that H0 hypothesis, and ‘pvalDiffKL_n-p’ refers to the p-value of the hypothesis “KL_n – KL_p=0”.

**Table S8. Meiotic behavior observed in Pollen Mother Cells at Metaphase I of hybrids carrying combinations of C_n_ chromosomes from *B. napus*.** % Cells: percentage of cells with the expected behavior. I and II represent univalents and bivalents, respectively.

**Table S9. Comparison of the meiotic gene content on the C09 chromosome between *B. napus* cv. ‘Darmor’ and *B. oleracea* cv. ‘RC34’.**

**Table S10. List of the meiotic gene content on the C04 and C08 chromosomes of *B. napus* cv. ‘Darmor’ and their orthologous genes in *A. thaliana*.**

