## Supplementary figures and images for "Genomic divergence shaped the genetic regulation of meiotic homologous recombination in *Brassica* allopolyploids"

### Supplemental Figure 1

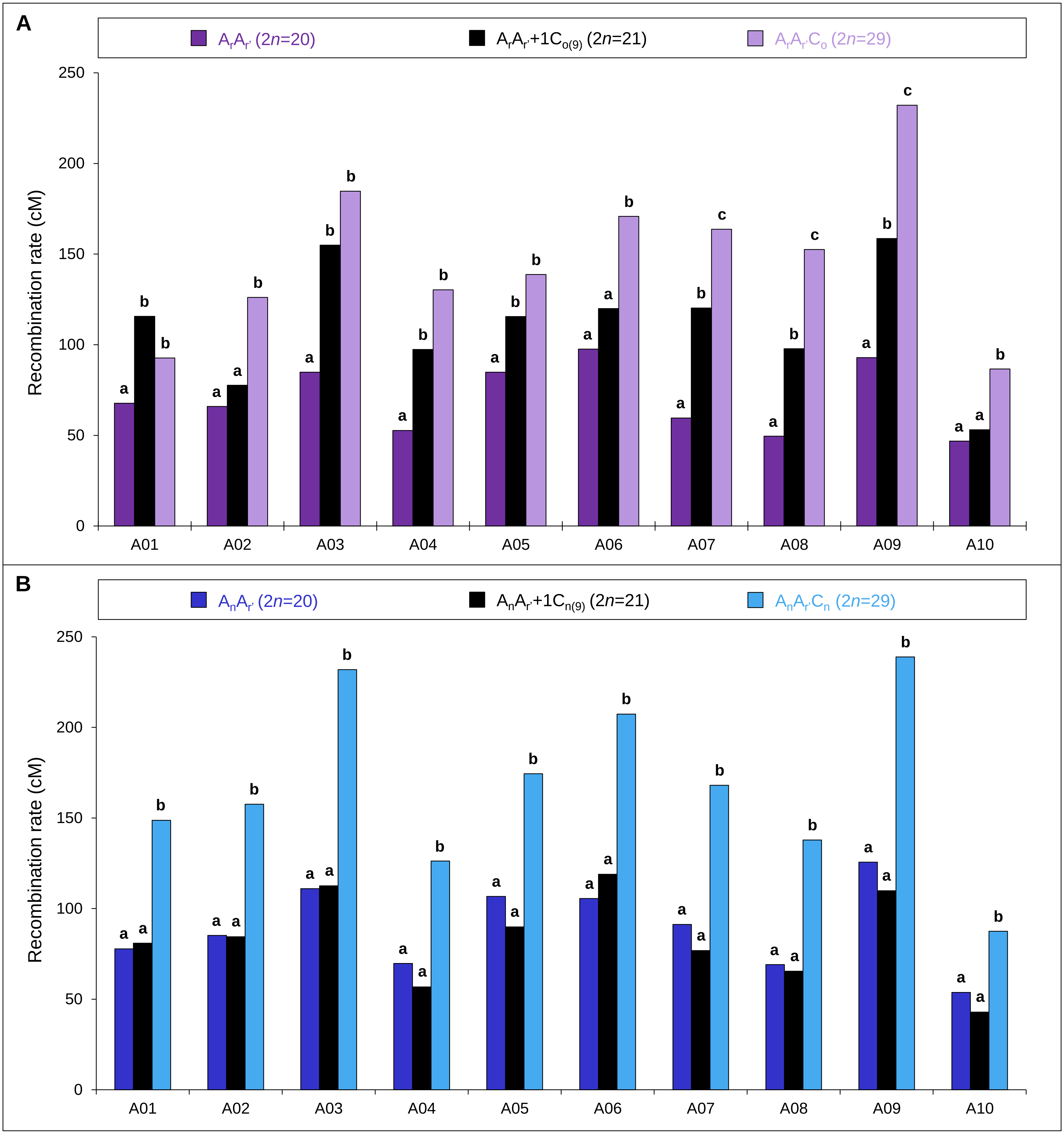

### Supplemental Figure 2

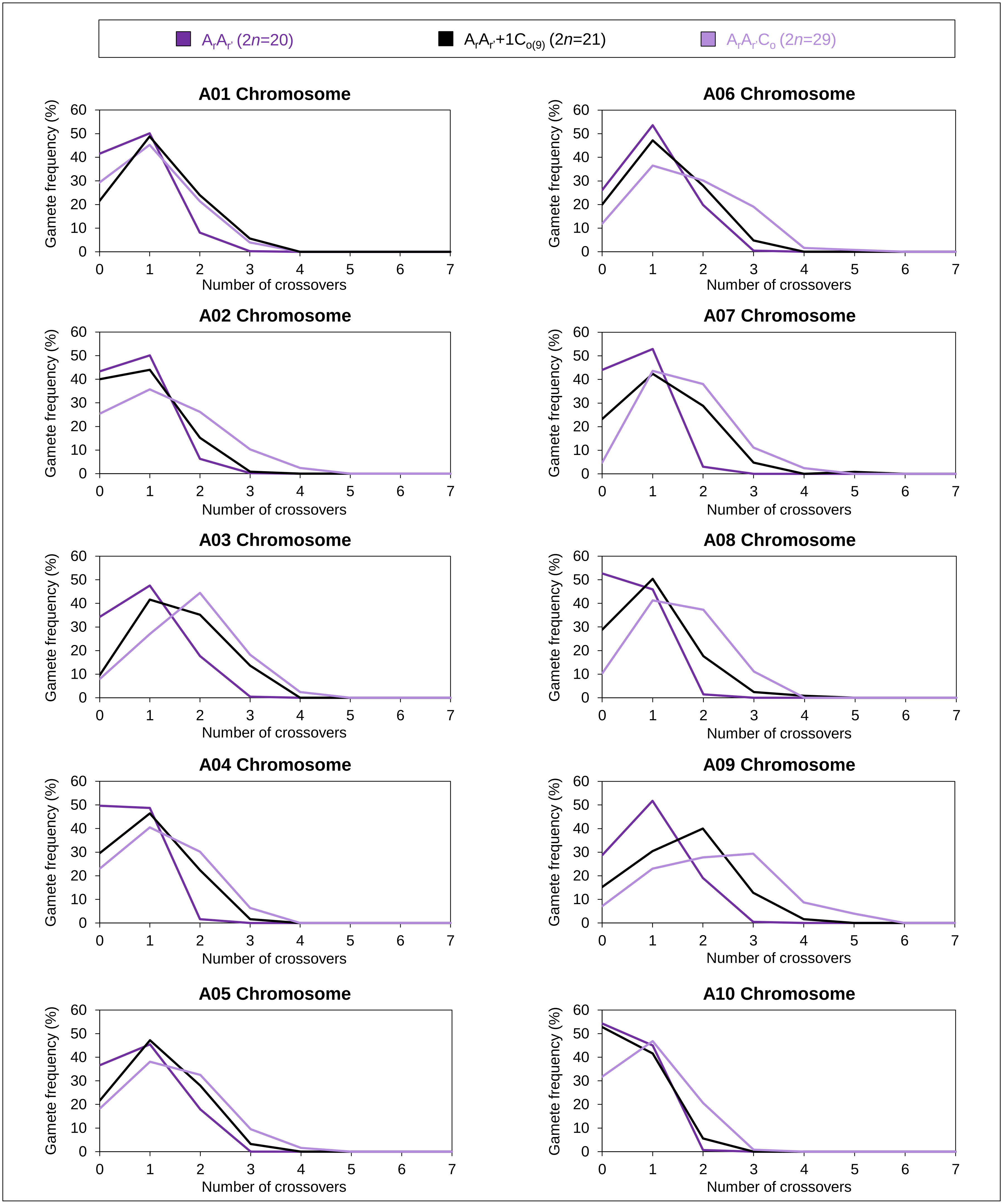

### Supplemental Figure 5

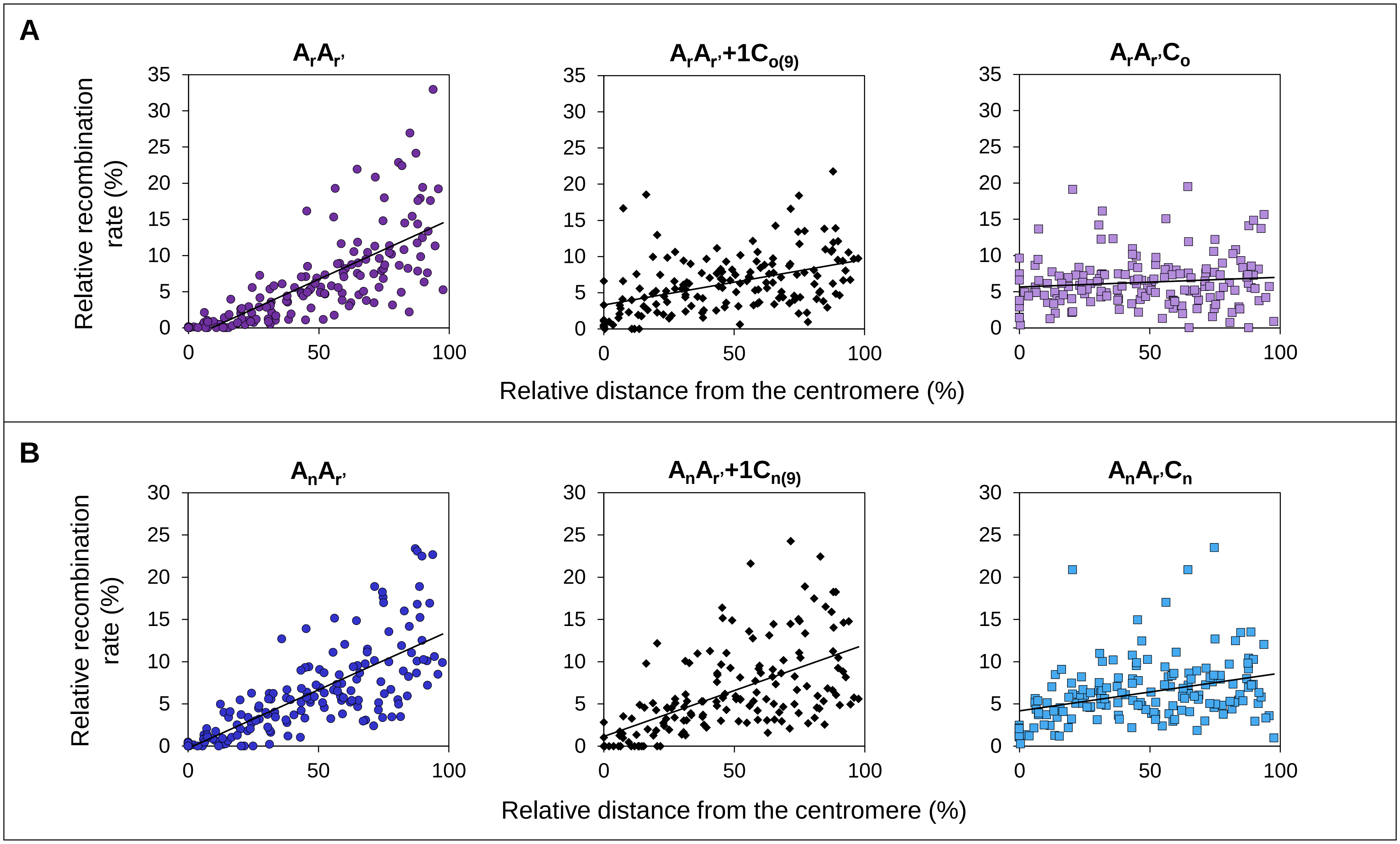

### Supplemental Figure 8

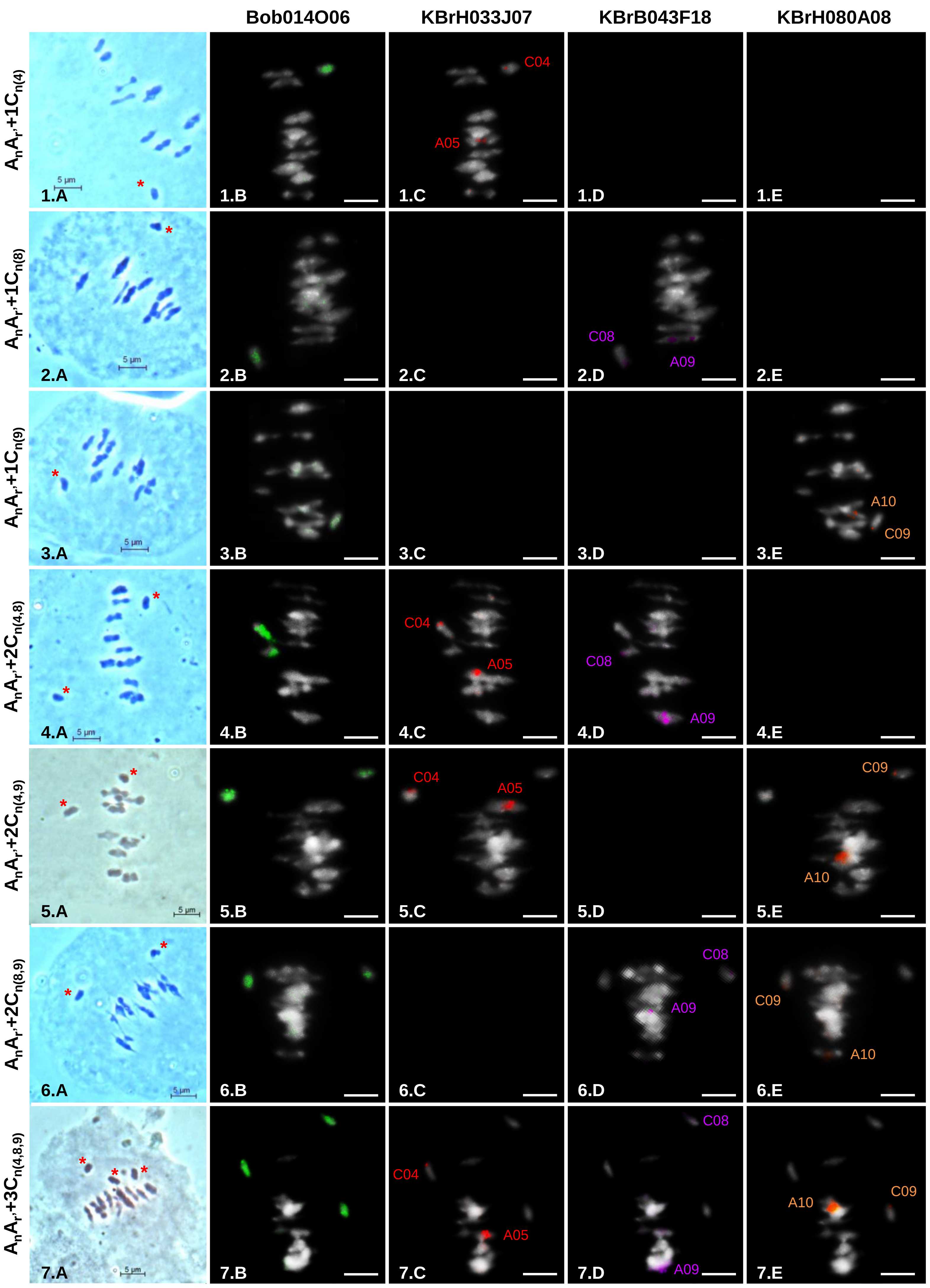

### Supplemental Figure 9

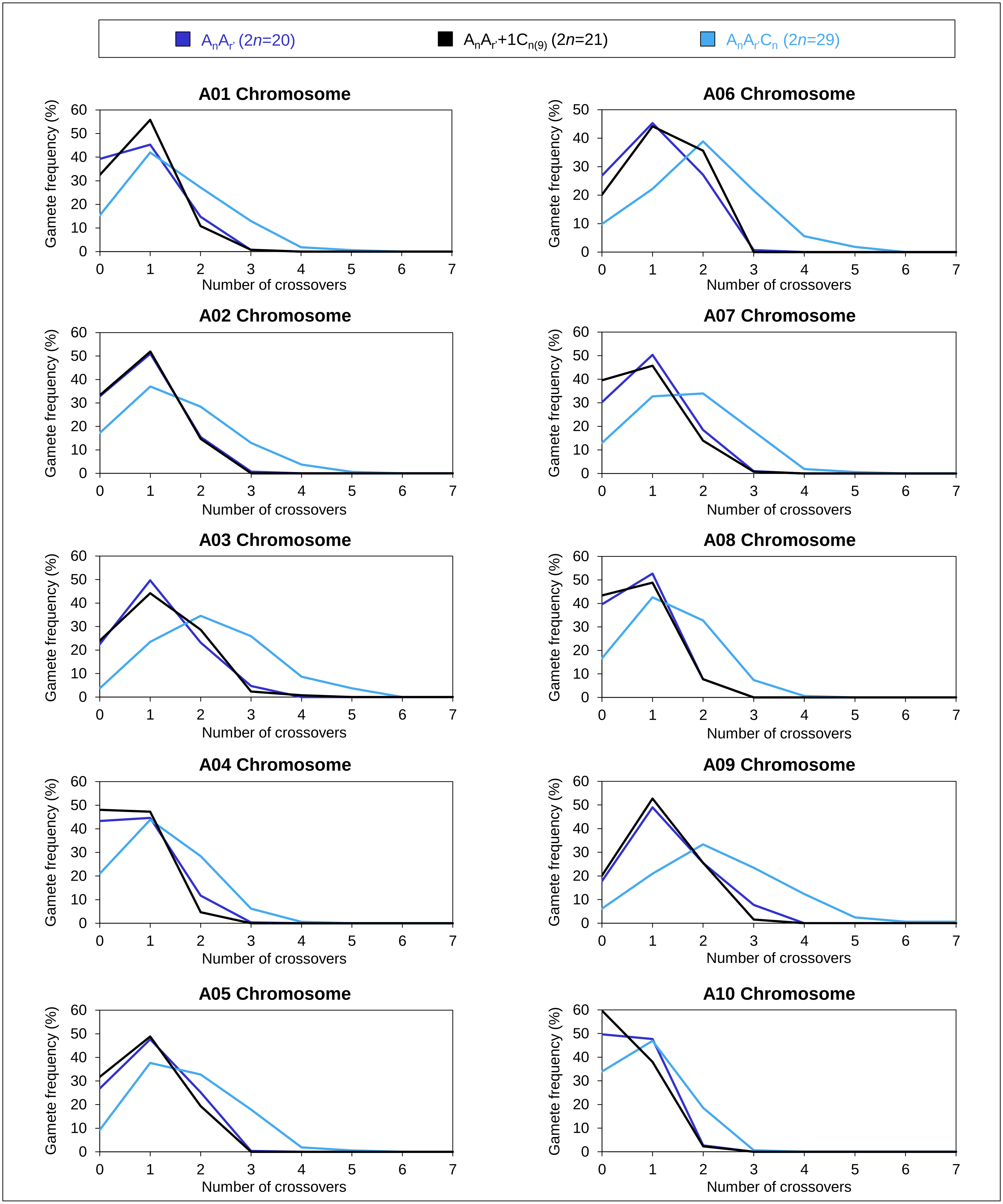

### Supplemental Figure 10

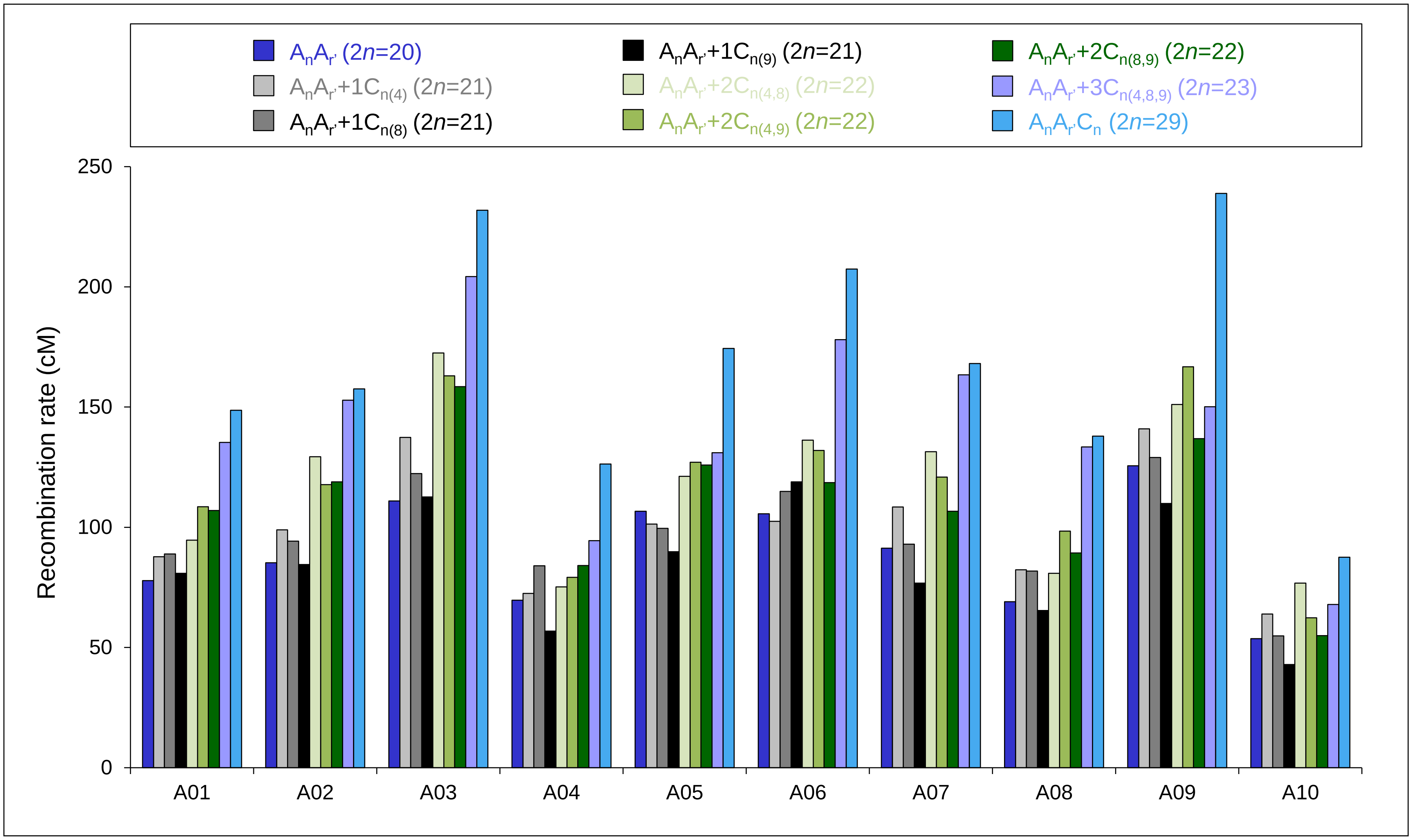
