## Supplemental Figure 3 for "Genomic divergence shaped the genetic regulation of meiotic homologous recombination in *Brassica* allopolyploids"

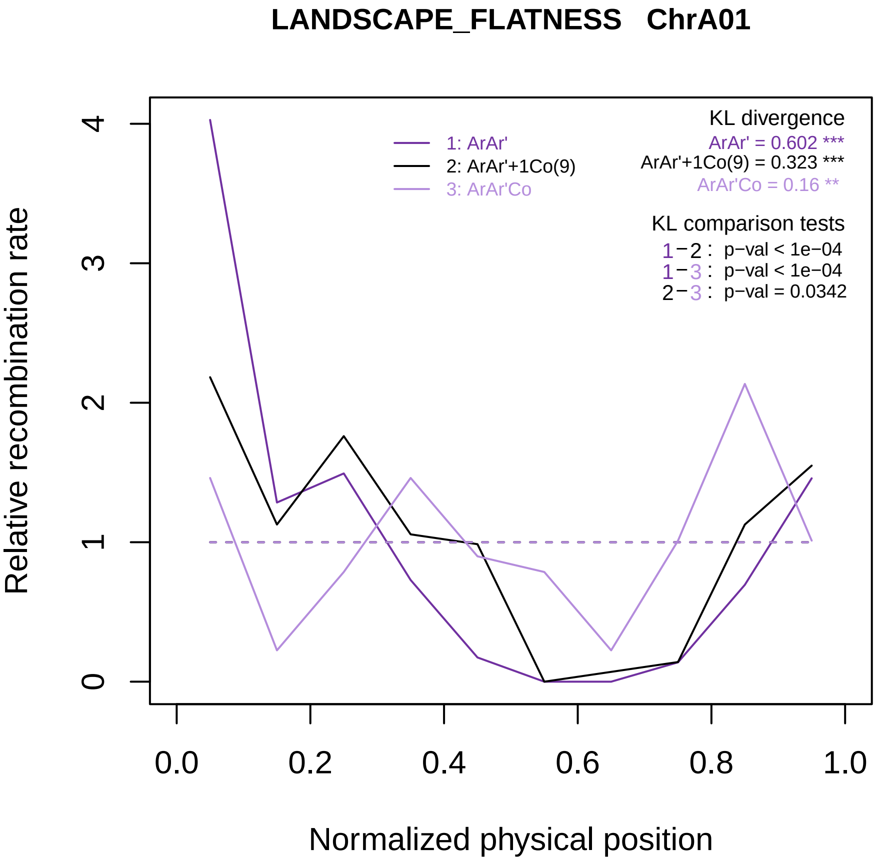

LANDSCAPE_FLATNESS   ChrA01
KL divergence
ArAr' = 0.602 ***
ArAr'+1Co(9) = 0.323 ***
ArAr'Co = 0.16 **
1: ArAr'
2: ArAr'+1Co(9)
3: ArAr'Co
KL comparison tests
1−2 : p−val < 1e−04
1−3 : p−val < 1e−04
2−3 : p−val = 0.0342
Relative recombination rate
0
1
2
3
4
0.0
0.2
0.4
0.6
0.8
1.0
Normalized physical position

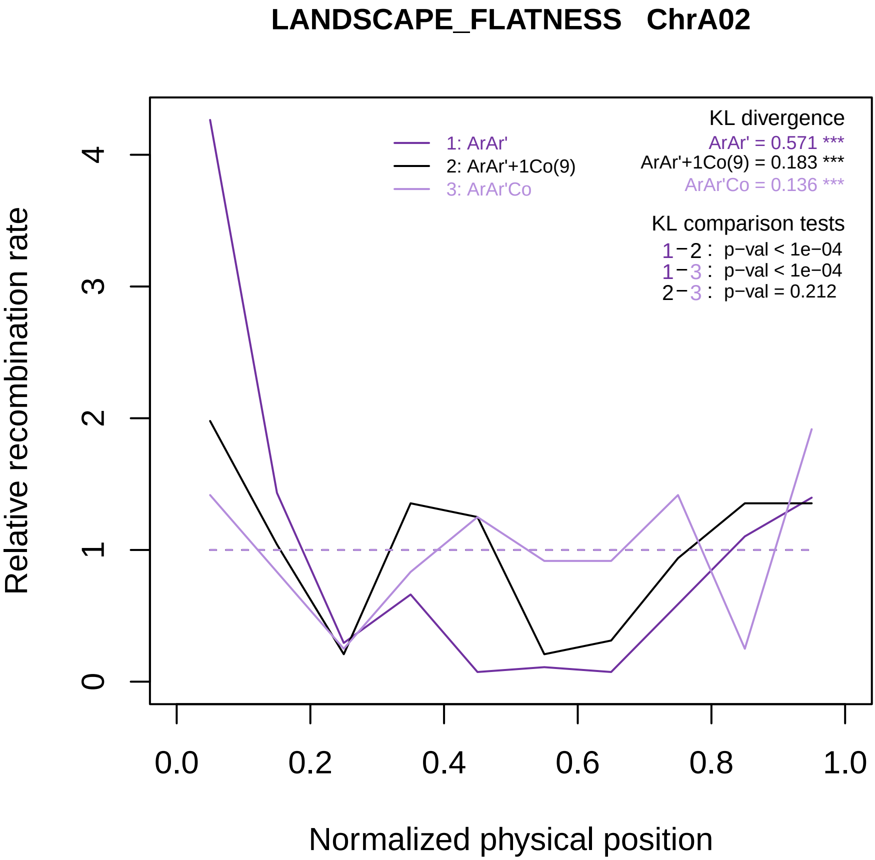
LANDSCAPE_FLATNESS ChrA02
KL divergence
ArAr' = 0.571 ***
ArAr'+1Co(9) = 0.183 ***
ArAr'Co = 0.136 ***
1: ArAr'
2: ArAr'+1Co(9)
3: ArAr'Co
KL comparison tests
1−2 : p−val < 1e−04
1−3 : p−val < 1e−04
2−3 : p−val = 0.212
Relative recombination rate
0
1
2
3
4
0.0
0.2
0.4
0.6
0.8
1.0
Normalized physical position

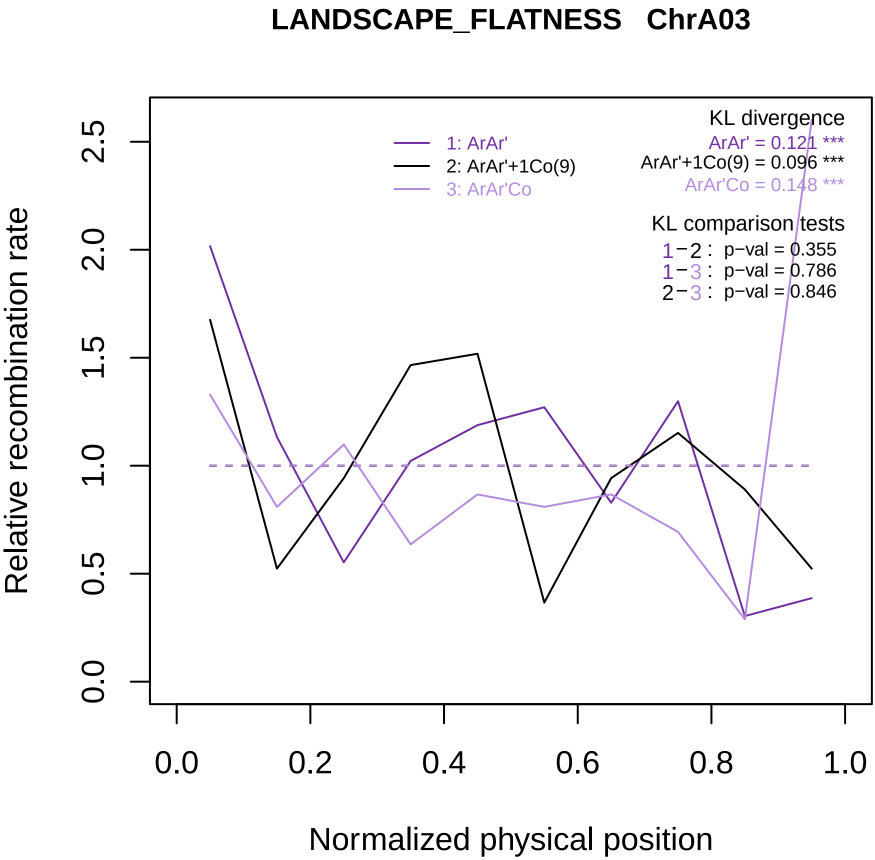

LANDSCAPE_FLATNESS ChrA03
KL divergence
ArAr' = 0.121 ***
ArAr'+1Co(9) = 0.096 ***
ArAr'Co = 0.148 ***
1: ArAr'
2: ArAr'+1Co(9)
3: ArAr'Co
KL comparison tests
1−2 : p−val = 0.355
1−3 : p−val = 0.786
2−3 : p−val = 0.846
Relative recombination rate
0.0
0.5
1.0
1.5
2.0
2.5
0.0
0.2
0.4
0.6
0.8
1.0
Normalized physical position

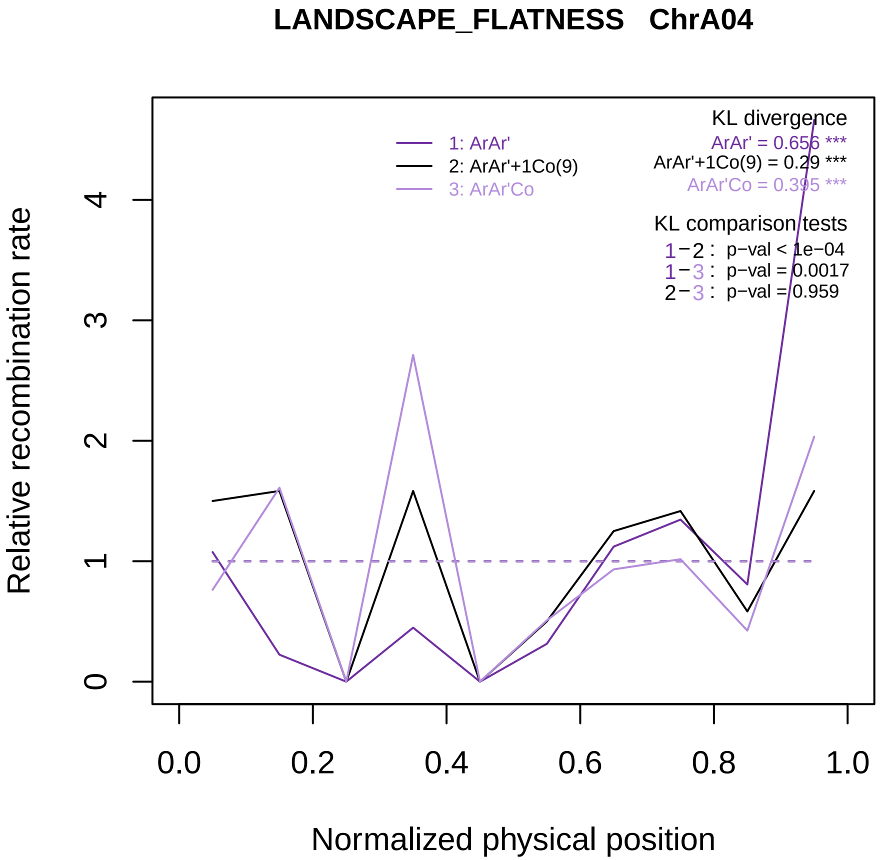

LANDSCAPE_FLATNESS ChrA04
KL divergence
ArAr' = 0.656 ***
ArAr'+1Co(9) = 0.29 ***
ArAr'Co = 0.395 ***
1: ArAr'
2: ArAr'+1Co(9)
3: ArAr'Co
KL comparison tests
1−2 : p−val < 1e−04
1−3 : p−val = 0.0017
2−3 : p−val = 0.959
Relative recombination rate
0
1
2
3
4
0.0
0.2
0.4
0.6
0.8
1.0
Normalized physical position

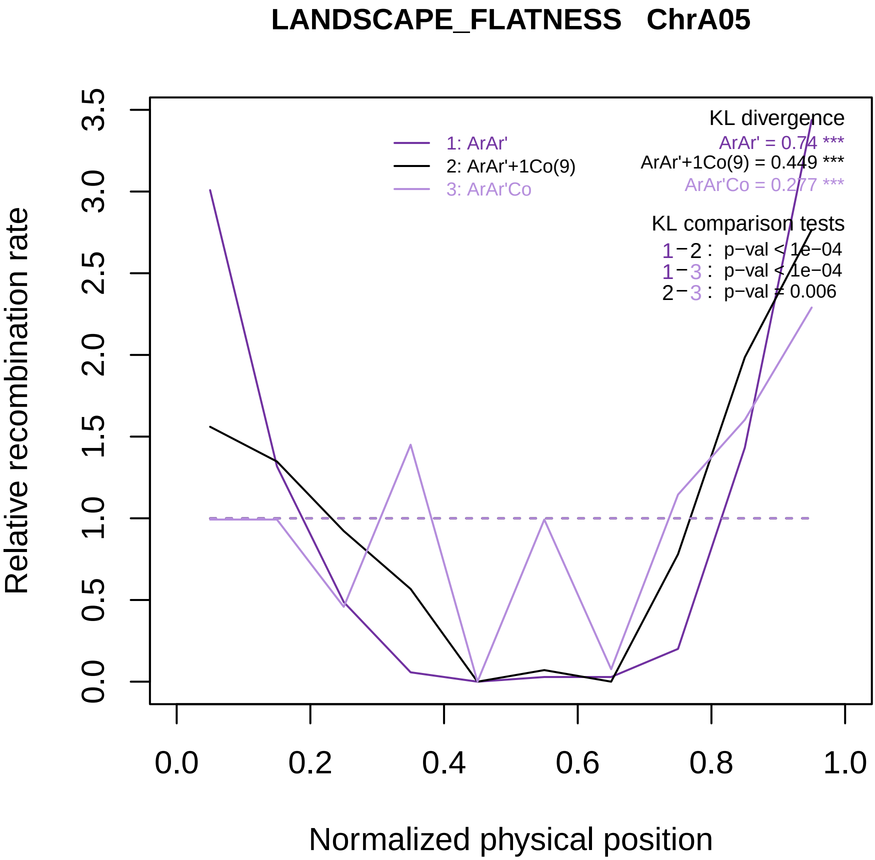
LANDSCAPE_FLATNESS ChrA05
Relative recombination rate
Normalized physical position
0.0
0.5
1.0
1.5
2.0
2.5
3.0
3.5
0.0
0.2
0.4
0.6
0.8
1.0
1: ArAr'
2: ArAr'+1Co(9)
3: ArAr'Co
KL divergence
ArAr' = 0.74 ***
ArAr'+1Co(9) = 0.449 ***
ArAr'Co = 0.277 ***
KL comparison tests
1−2 : p−val < 1e−04
1−3 : p−val < 1e−04
2−3 : p−val = 0.006

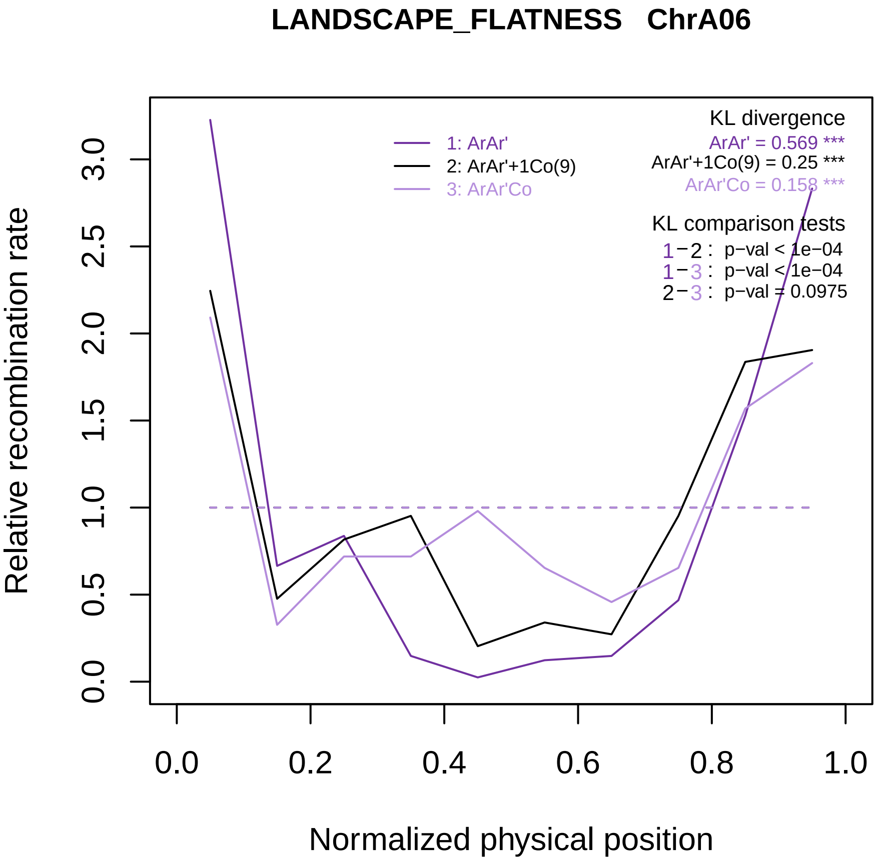

LANDSCAPE_FLATNESS ChrA06
KL divergence
ArAr' = 0.569 ***
ArAr'+1Co(9) = 0.25 ***
ArAr'Co = 0.158 ***
1: ArAr'
2: ArAr'+1Co(9)
3: ArAr'Co
KL comparison tests
1−2 : p−val < 1e−04
1−3 : p−val < 1e−04
2−3 : p−val = 0.0975
Relative recombination rate
0.0
0.5
1.0
1.5
2.0
2.5
3.0
0.0
0.2
0.4
0.6
0.8
1.0
Normalized physical position

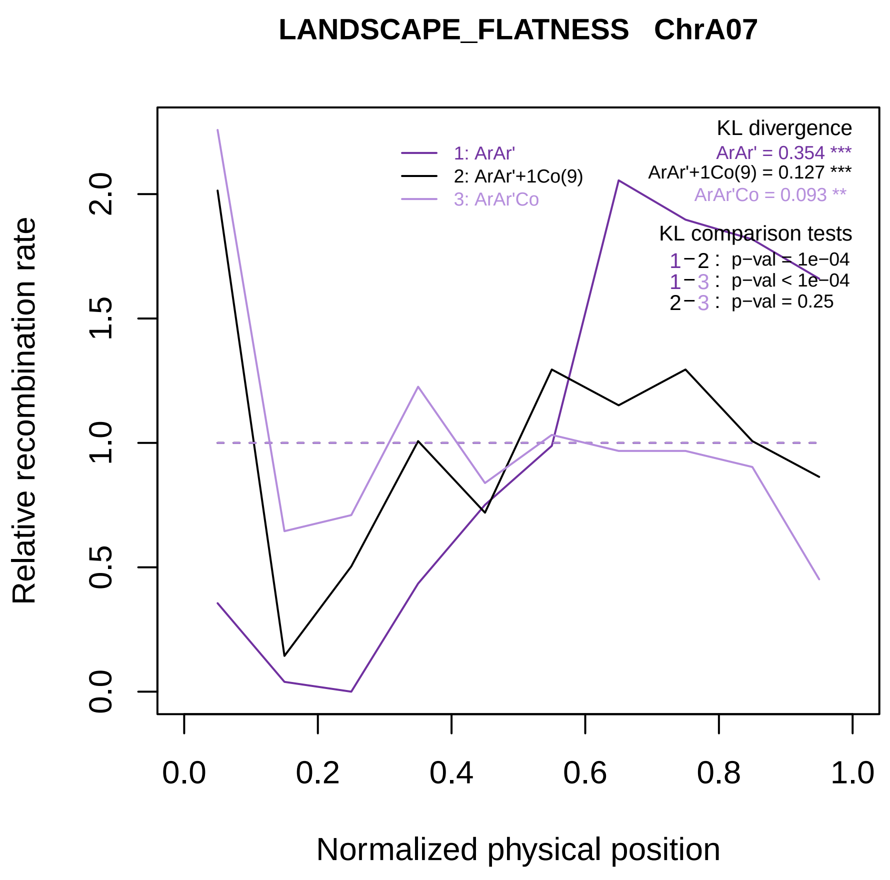
LANDSCAPE_FLATNESS ChrA07
KL divergence
ArAr' = 0.354 ***
ArAr'+1Co(9) = 0.127 ***
ArAr'Co = 0.093 **
1: ArAr'
2: ArAr'+1Co(9)
3: ArAr'Co
KL comparison tests
1−2 : p−val = 1e−04
1−3 : p−val < 1e−04
2−3 : p−val = 0.25
Relative recombination rate
0.0
0.5
1.0
1.5
2.0
0.0
0.2
0.4
0.6
0.8
1.0
Normalized physical position

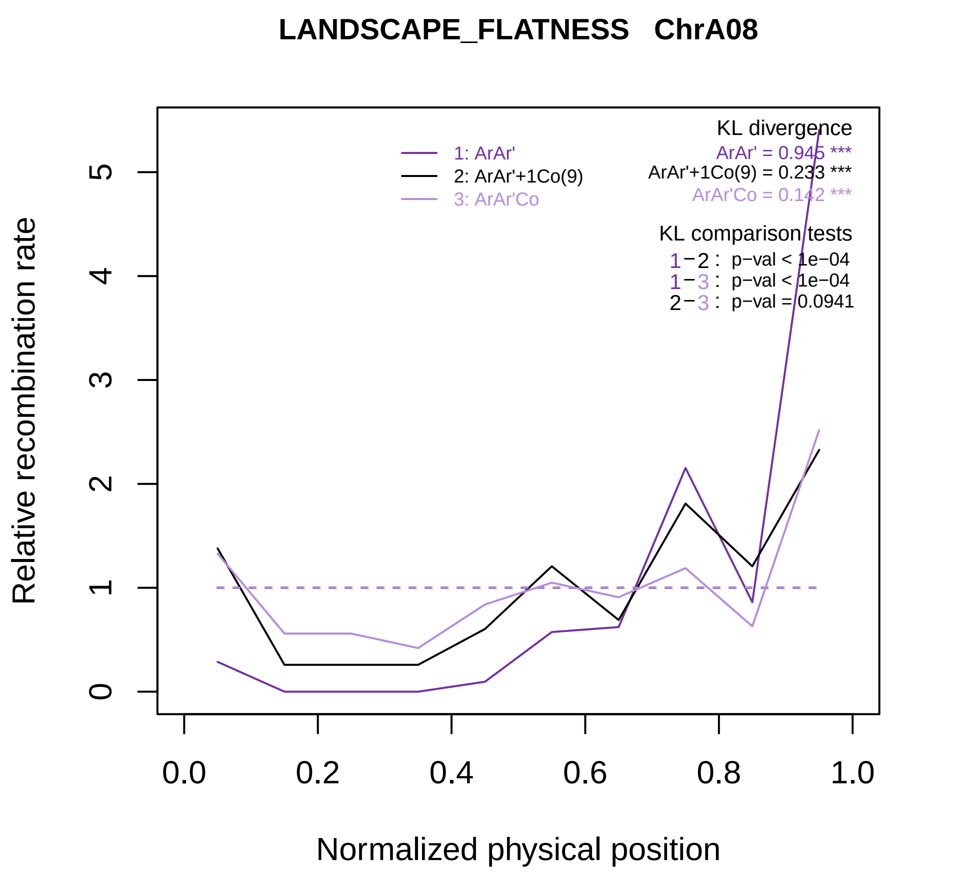

LANDSCAPE_FLATNESS ChrA08
Relative recombination rate
Normalized physical position
0
1
2
3
4
5
0.0
0.2
0.4
0.6
0.8
1.0
1: ArAr'
2: ArAr'+1Co(9)
3: ArAr'Co
KL divergence
ArAr' = 0.945 ***
ArAr'+1Co(9) = 0.233 ***
ArAr'Co = 0.142 ***
KL comparison tests
1−2 : p−val < 1e−04
1−3 : p−val < 1e−04
2−3 : p−val = 0.0941

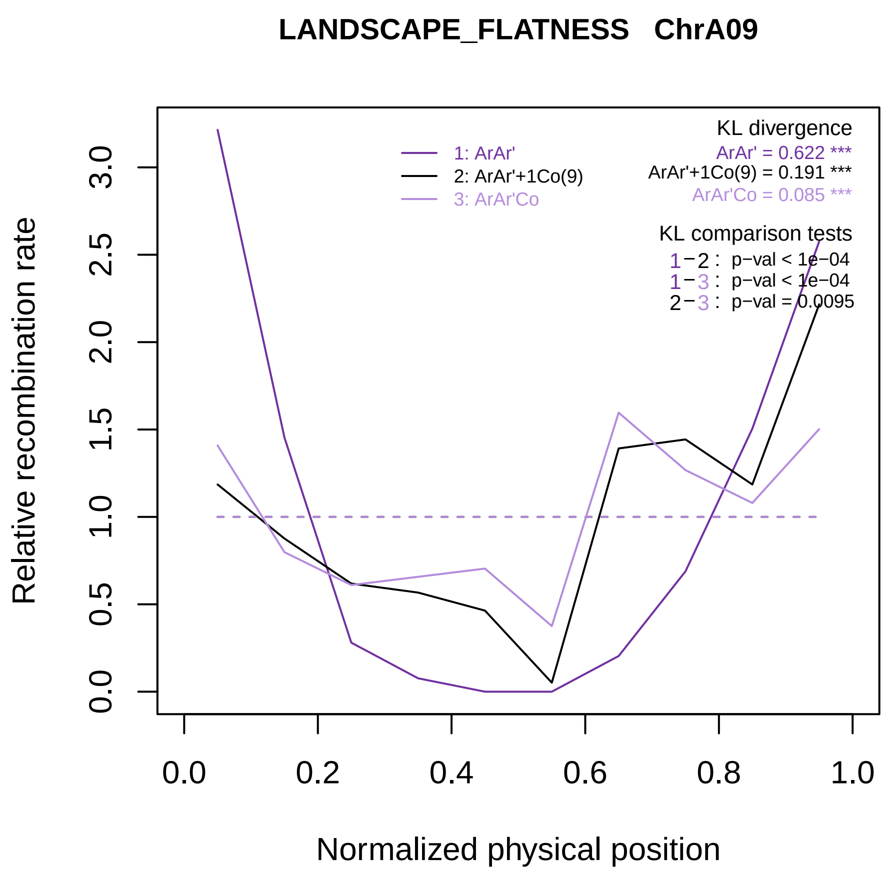

LANDSCAPE_FLATNESS ChrA09
Relative recombination rate
Normalized physical position
0.0
0.5
1.0
1.5
2.0
2.5
3.0
0.0
0.2
0.4
0.6
0.8
1.0
1: ArAr'
2: ArAr'+1Co(9)
3: ArAr'Co
KL divergence
ArAr' = 0.622 ***
ArAr'+1Co(9) = 0.191 ***
ArAr'Co = 0.085 ***
KL comparison tests
1−2 : p−val < 1e−04
1−3 : p−val < 1e−04
2−3 : p−val = 0.0095

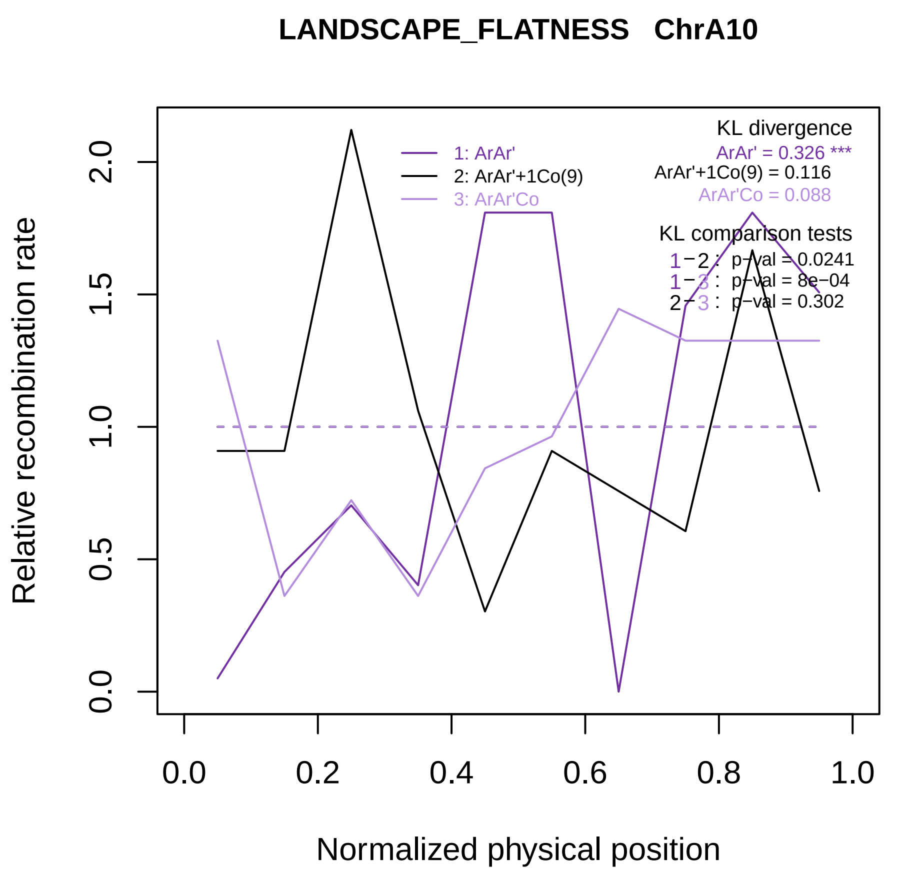

LANDSCAPE_FLATNESS ChrA10
KL divergence
ArAr' = 0.326 ***
ArAr'+1Co(9) = 0.116
ArAr'Co = 0.088
1: ArAr'
2: ArAr'+1Co(9)
3: ArAr'Co
KL comparison tests
1−2 : p−val = 0.0241
1−3 : p−val = 8e−04
2−3 : p−val = 0.302
Relative recombination rate
0.0
0.5
1.0
1.5
2.0
0.0
0.2
0.4
0.6
0.8
1.0
Normalized physical position

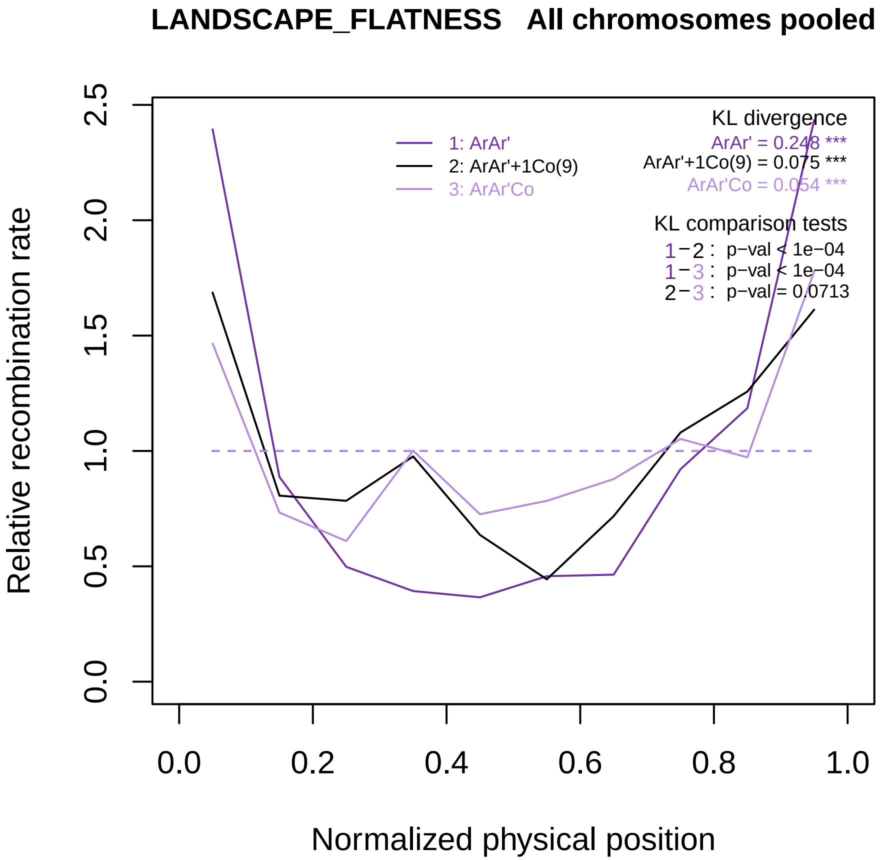

LANDSCAPE_FLATNESS All chromosomes pooled
Relative recombination rate
Normalized physical position
0.0
0.5
1.0
1.5
2.0
2.5
0.0
0.2
0.4
0.6
0.8
1.0
1: ArAr'
2: ArAr'+1Co(9)
3: ArAr'Co
KL divergence
ArAr' = 0.248 ***
ArAr'+1Co(9) = 0.075 ***
ArAr'Co = 0.054 ***
KL comparison tests
1−2 : p−val < 1e−04
1−3 : p−val < 1e−04
2−3 : p−val = 0.0713

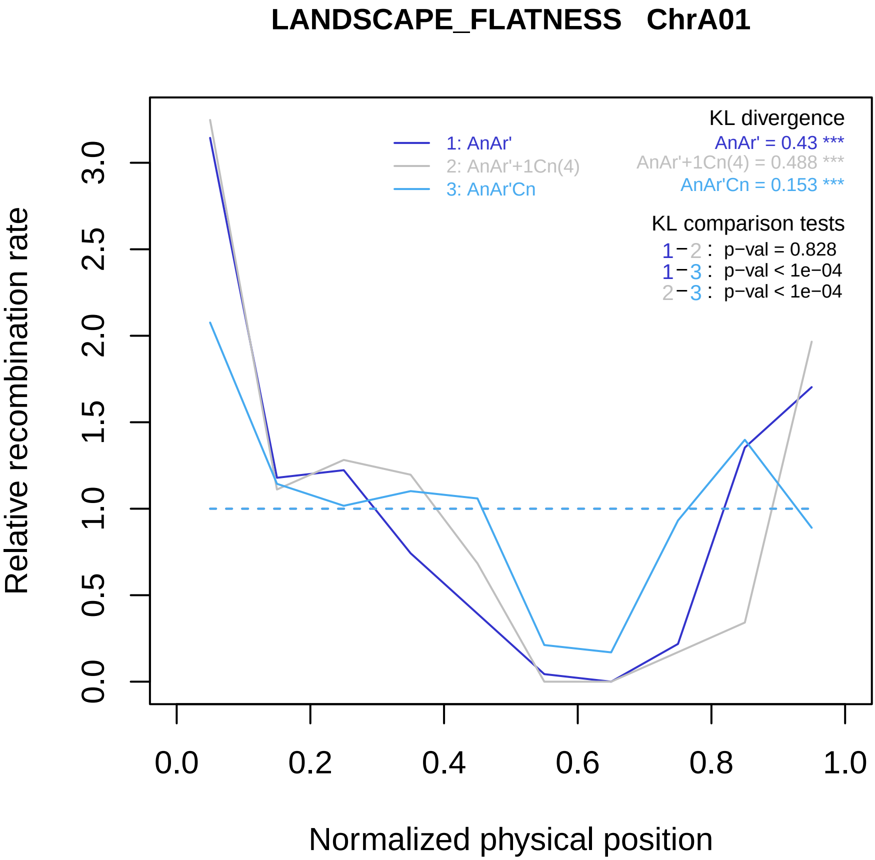
LANDSCAPE_FLATNESS ChrA01
Relative recombination rate
0.0
0.5
1.0
1.5
2.0
2.5
3.0
Normalized physical position
0.0
0.2
0.4
0.6
0.8
1.0
1: AnAr'
2: AnAr'+1Cn(4)
3: AnAr'Cn
KL divergence
AnAr' = 0.43 ***
AnAr'+1Cn(4) = 0.488 ***
AnAr'Cn = 0.153 ***
KL comparison tests
1−2 : p−val = 0.828
1−3 : p−val < 1e−04
2−3 : p−val < 1e−04

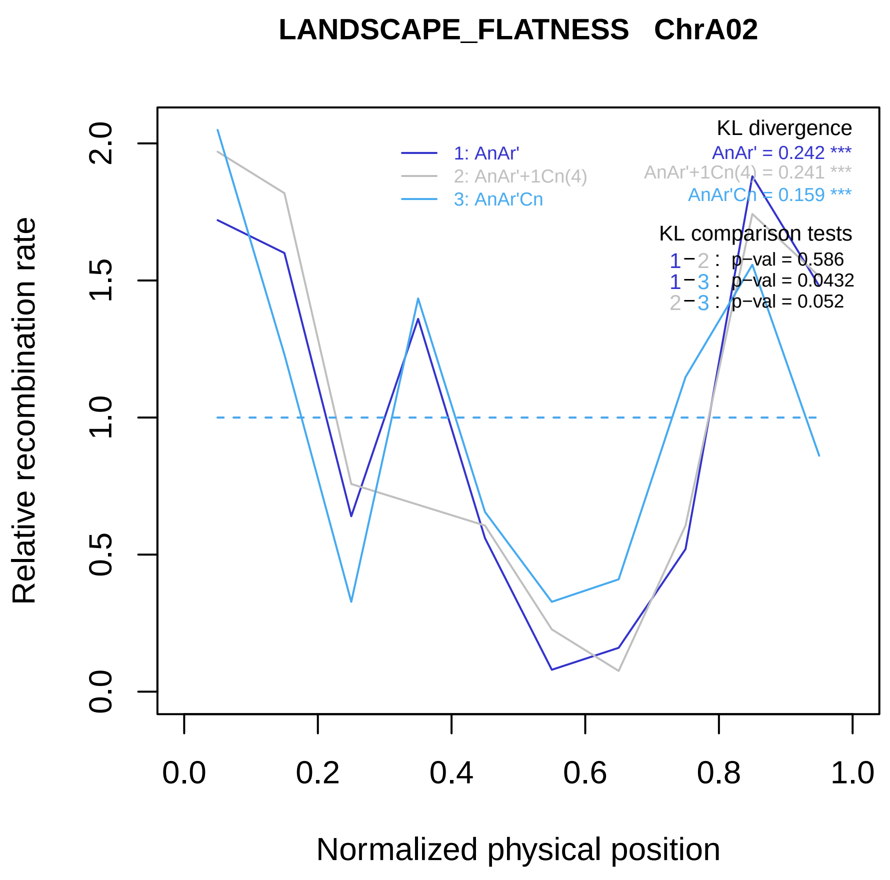

LANDSCAPE_FLATNESS ChrA02
Relative recombination rate
Normalized physical position
0.0
0.5
1.0
1.5
2.0
0.0
0.2
0.4
0.6
0.8
1.0
1: AnAr'
2: AnAr'+1Cn(4)
3: AnAr'Cn
KL divergence
AnAr' = 0.242 ***
AnAr'+1Cn(4) = 0.241 ***
AnAr'Cn = 0.159 ***
KL comparison tests
1−2 : p−val = 0.586
1−3 : p−val = 0.0432
2−3 : p−val = 0.052

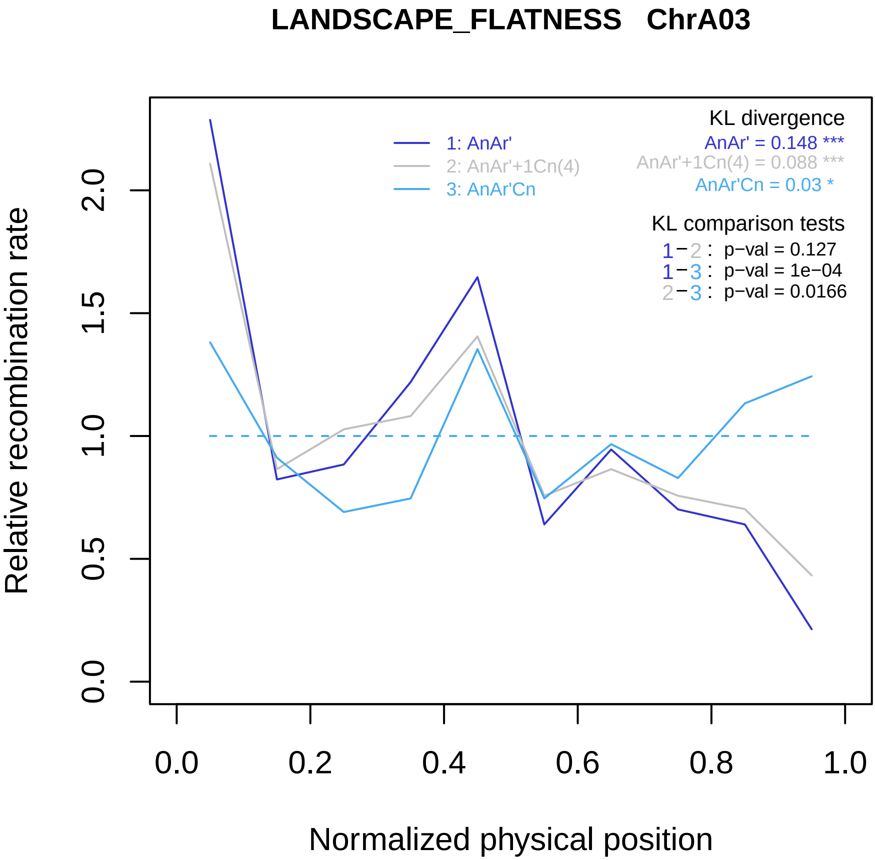

LANDSCAPE_FLATNESS ChrA03
Relative recombination rate
Normalized physical position
0.0
0.5
1.0
1.5
2.0
0.0
0.2
0.4
0.6
0.8
1.0
1: AnAr'
2: AnAr'+1Cn(4)
3: AnAr'Cn
KL divergence
AnAr' = 0.148 ***
AnAr'+1Cn(4) = 0.088 ***
AnAr'Cn = 0.03 *
KL comparison tests
1−2 : p−val = 0.127
1−3 : p−val = 1e−04
2−3 : p−val = 0.0166

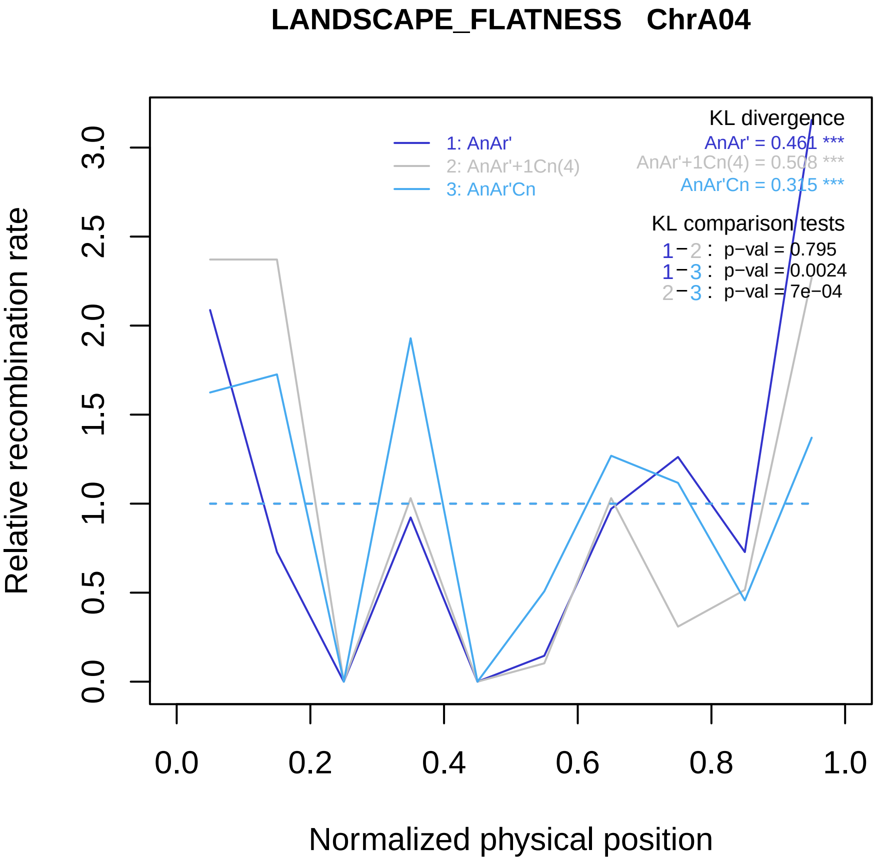

LANDSCAPE_FLATNESS ChrA04
Relative recombination rate
0.0
0.5
1.0
1.5
2.0
2.5
3.0
Normalized physical position
0.0
0.2
0.4
0.6
0.8
1.0
1: AnAr'
2: AnAr'+1Cn(4)
3: AnAr'Cn
KL divergence
AnAr' = 0.461 ***
AnAr'+1Cn(4) = 0.508 ***
AnAr'Cn = 0.315 ***
KL comparison tests
1−2 : p−val = 0.795
1−3 : p−val = 0.0024
2−3 : p−val = 7e−04

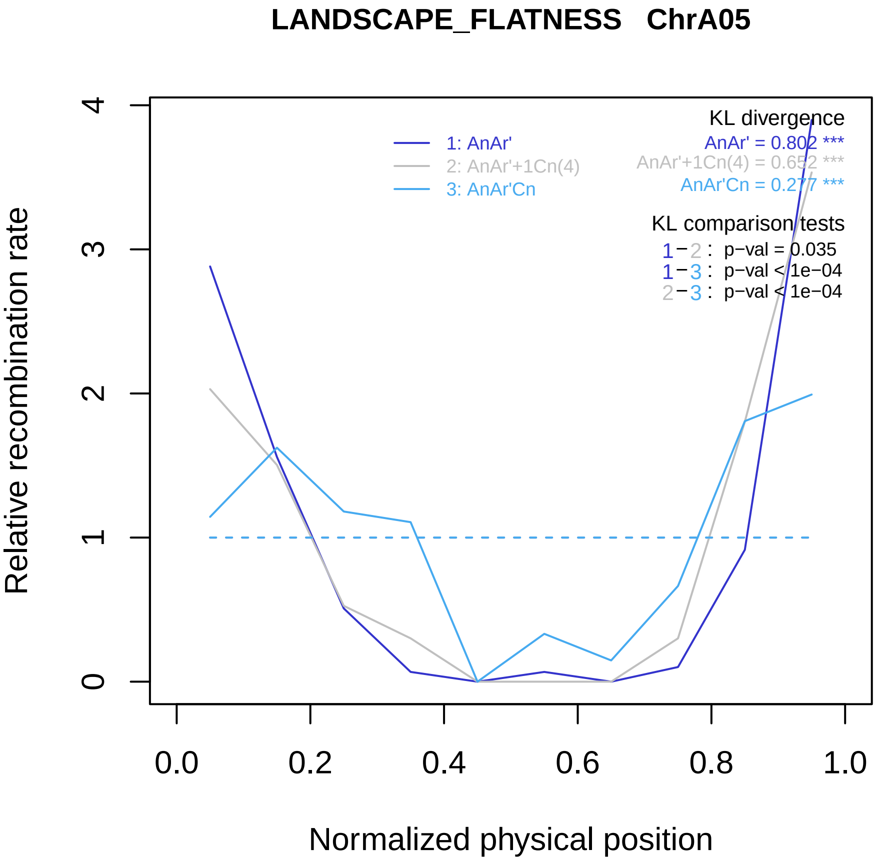

LANDSCAPE_FLATNESS ChrA05
Relative recombination rate
Normalized physical position
0
1
2
3
4
0.0
0.2
0.4
0.6
0.8
1.0
1: AnAr'
2: AnAr'+1Cn(4)
3: AnAr'Cn
KL divergence
AnAr' = 0.802 ***
AnAr'+1Cn(4) = 0.652 ***
AnAr'Cn = 0.277 ***
KL comparison tests
1−2 : p−val = 0.035
1−3 : p−val < 1e−04
2−3 : p−val < 1e−04

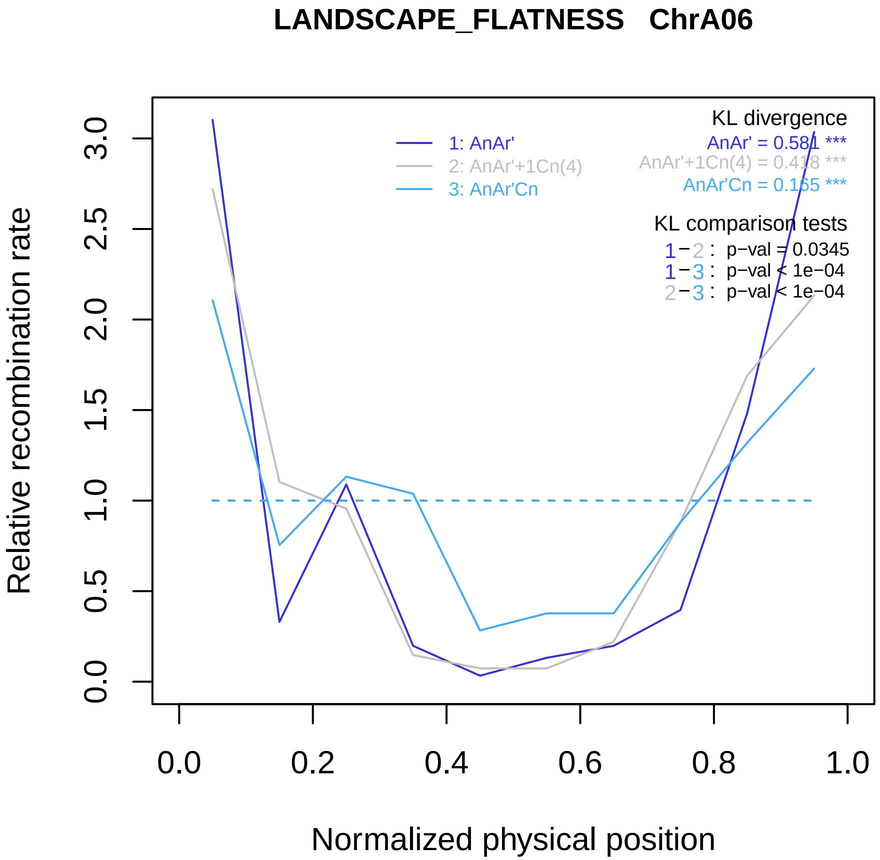
LANDSCAPE_FLATNESS ChrA06
Relative recombination rate
Normalized physical position
0.0
0.5
1.0
1.5
2.0
2.5
3.0
0.0
0.2
0.4
0.6
0.8
1.0
1: AnAr'
2: AnAr'+1Cn(4)
3: AnAr'Cn
KL divergence
AnAr' = 0.581 ***
AnAr'+1Cn(4) = 0.418 ***
AnAr'Cn = 0.165 ***
KL comparison tests
1−2 : p−val = 0.0345
1−3 : p−val < 1e−04
2−3 : p−val < 1e−04

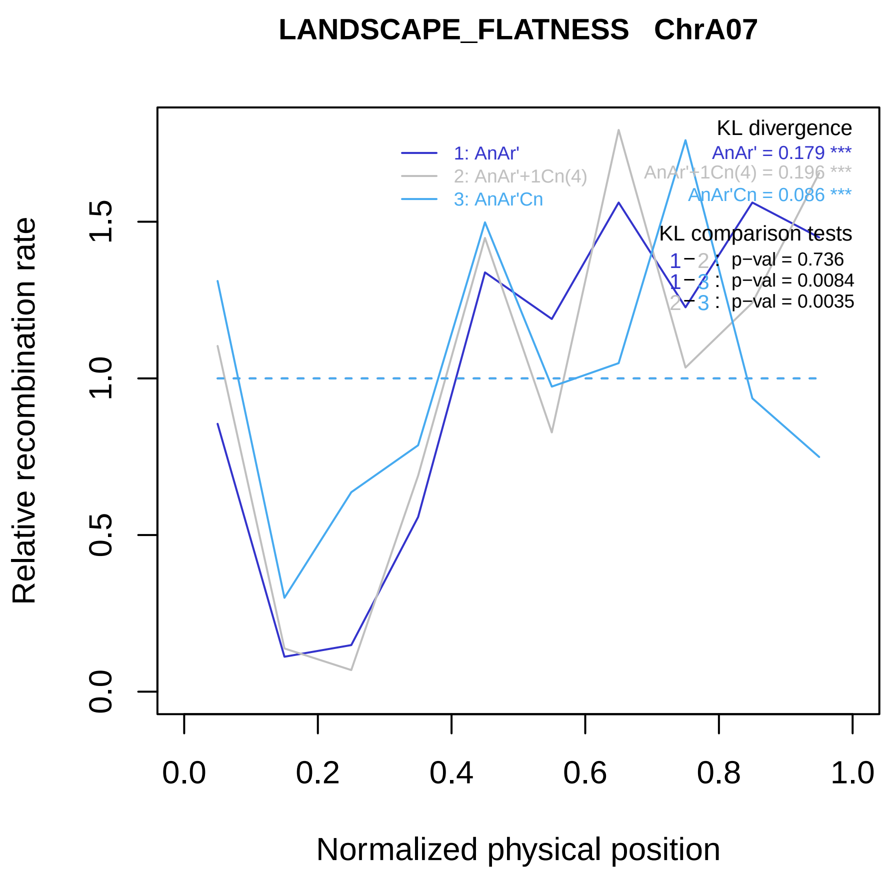

LANDSCAPE_FLATNESS   ChrA07
Relative recombination rate
Normalized physical position
0.0
0.5
1.0
1.5
0.0
0.2
0.4
0.6
0.8
1.0
1: AnAr'
2: AnAr'+1Cn(4)
3: AnAr'Cn
KL divergence
AnAr' = 0.179 ***
AnAr'+1Cn(4) = 0.196 ***
AnAr'Cn = 0.086 ***
KL comparison tests
1 − 2 : p−val = 0.736
1 − 3 : p−val = 0.0084
2 − 3 : p−val = 0.0035

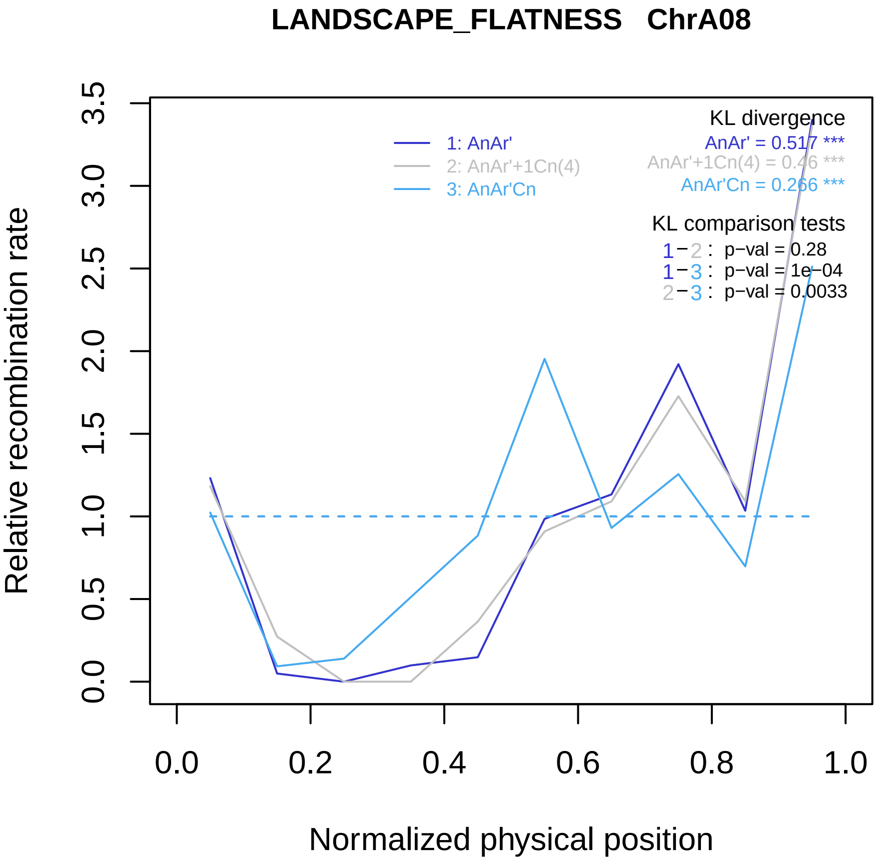
LANDSCAPE_FLATNESS ChrA08
Relative recombination rate
0.0
0.5
1.0
1.5
2.0
2.5
3.0
3.5
Normalized physical position
0.0
0.2
0.4
0.6
0.8
1.0
1: AnAr'
2: AnAr'+1Cn(4)
3: AnAr'Cn
KL divergence
AnAr' = 0.517 ***
AnAr'+1Cn(4) = 0.46 ***
AnAr'Cn = 0.266 ***
KL comparison tests
1−2 : p−val = 0.28
1−3 : p−val = 1e−04
2−3 : p−val = 0.0033

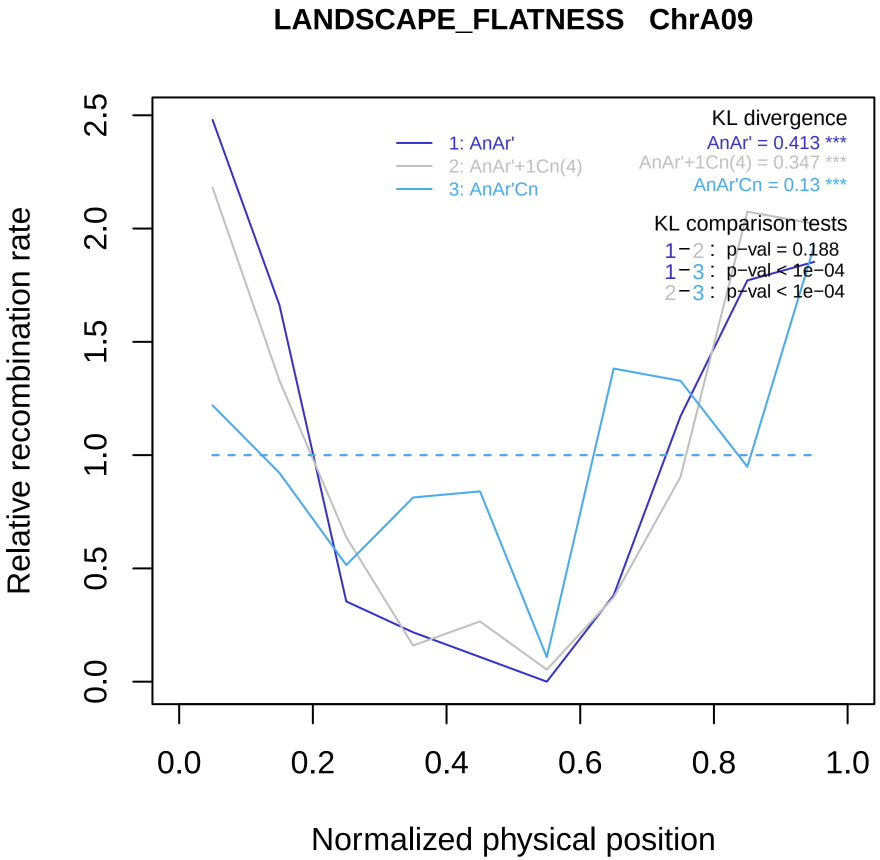

LANDSCAPE_FLATNESS ChrA09
Relative recombination rate
Normalized physical position
0.0
0.5
1.0
1.5
2.0
2.5
0.0
0.2
0.4
0.6
0.8
1.0
1: AnAr'
2: AnAr'+1Cn(4)
3: AnAr'Cn
KL divergence
AnAr' = 0.413 ***
AnAr'+1Cn(4) = 0.347 ***
AnAr'Cn = 0.13 ***
KL comparison tests
1−2 : p−val = 0.188
1−3 : p−val < 1e−04
2−3 : p−val < 1e−04

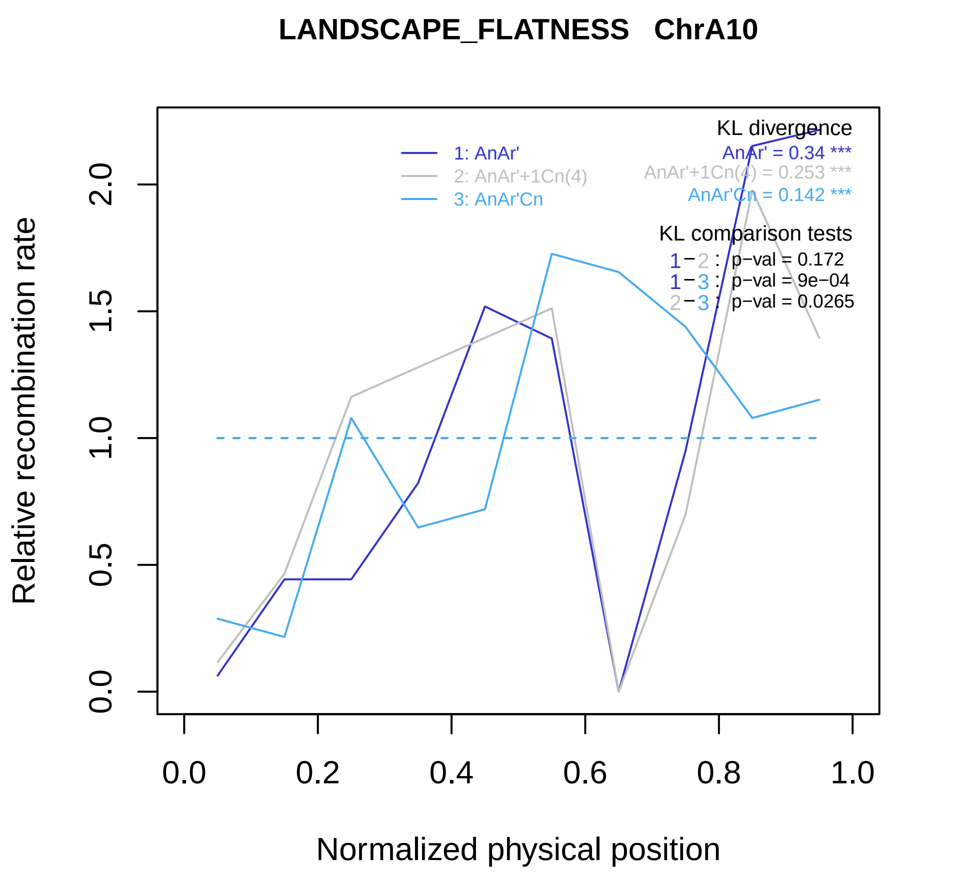

LANDSCAPE_FLATNESS ChrA10
Relative recombination rate
Normalized physical position
0.0
0.5
1.0
1.5
2.0
0.0
0.2
0.4
0.6
0.8
1.0
1: AnAr'
2: AnAr'+1Cn(4)
3: AnAr'Cn
KL divergence
AnAr' = 0.34 ***
AnAr'+1Cn(4) = 0.253 ***
AnAr'Cn = 0.142 ***
KL comparison tests
1−2 : p−val = 0.172
1−3 : p−val = 9e−04
2−3 : p−val = 0.0265

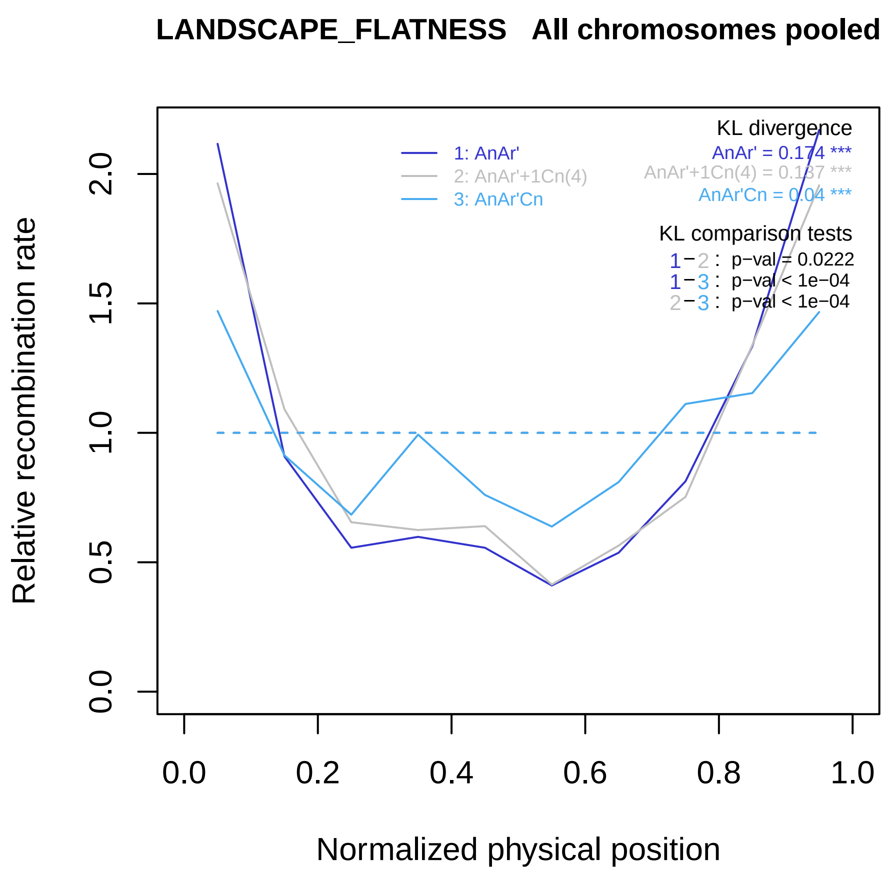
LANDSCAPE_FLATNESS All chromosomes pooled
KL divergence
AnAr' = 0.174 ***
AnAr'+1Cn(4) = 0.137 ***
AnAr'Cn = 0.04 ***
1: AnAr'
2: AnAr'+1Cn(4)
3: AnAr'Cn
KL comparison tests
1−2 : p−val = 0.0222
1−3 : p−val < 1e−04
2−3 : p−val < 1e−04
Relative recombination rate
0.0
0.5
1.0
1.5
2.0
0.0
0.2
0.4
0.6
0.8
1.0
Normalized physical position

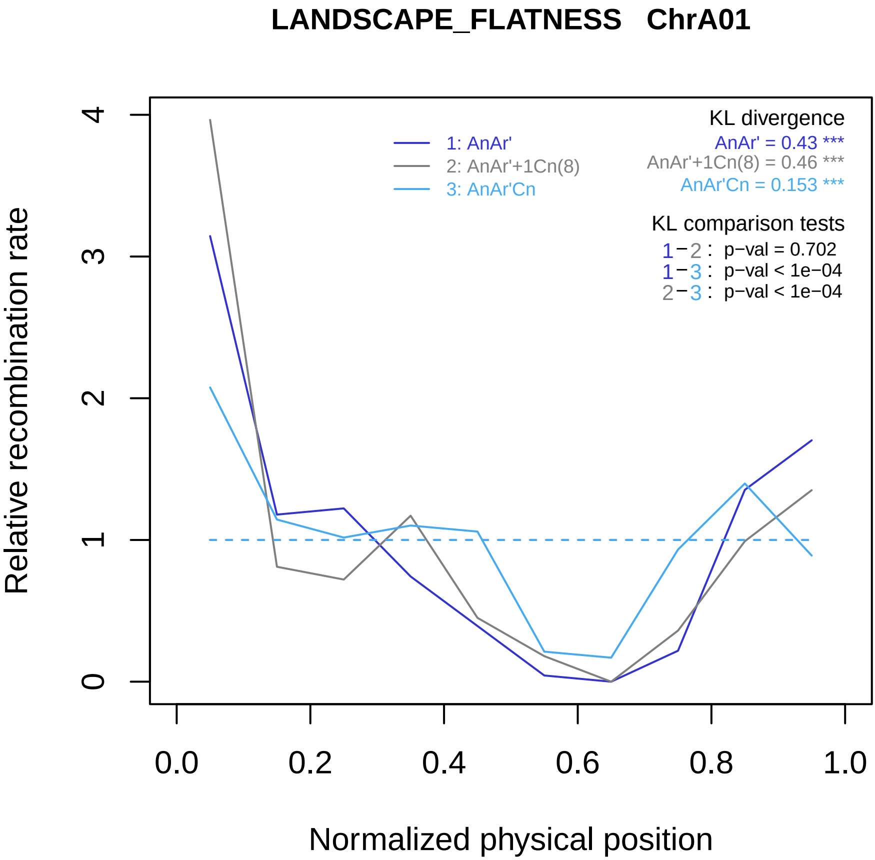

LANDSCAPE_FLATNESS ChrA01
KL divergence
AnAr' = 0.43 ***
AnAr'+1Cn(8) = 0.46 ***
AnAr'Cn = 0.153 ***
1: AnAr'
2: AnAr'+1Cn(8)
3: AnAr'Cn
KL comparison tests
1−2 : p−val = 0.702
1−3 : p−val < 1e−04
2−3 : p−val < 1e−04
Relative recombination rate
0
1
2
3
4
0.0
0.2
0.4
0.6
0.8
1.0
Normalized physical position

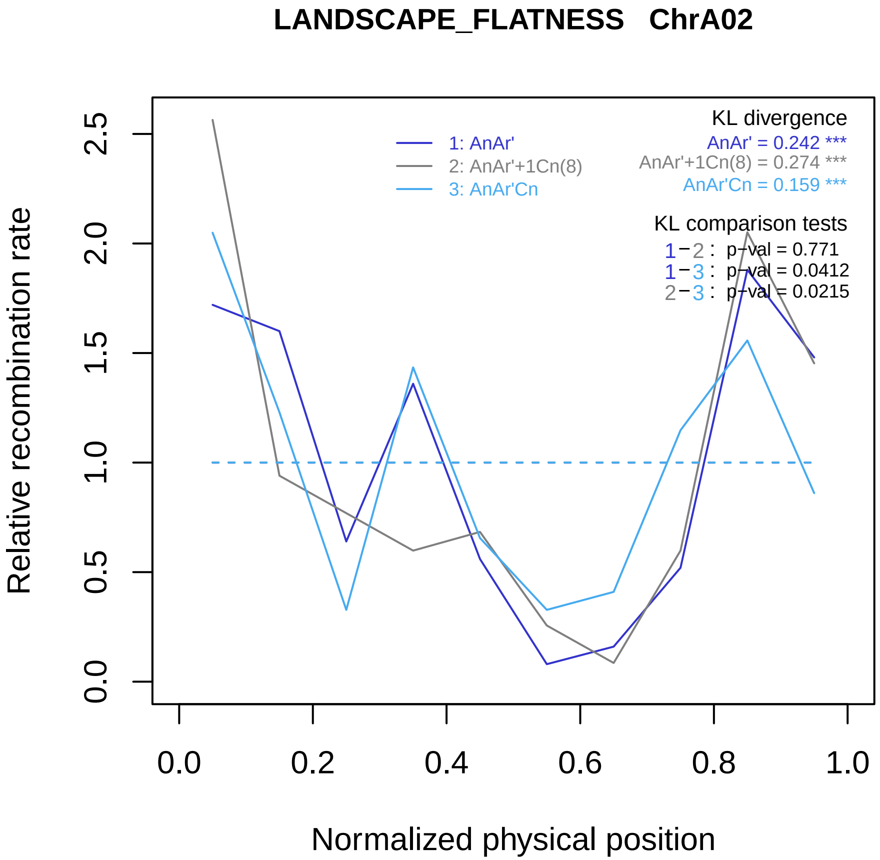

LANDSCAPE_FLATNESS ChrA02
KL divergence
AnAr' = 0.242 ***
AnAr'+1Cn(8) = 0.274 ***
AnAr'Cn = 0.159 ***
1: AnAr'
2: AnAr'+1Cn(8)
3: AnAr'Cn
KL comparison tests
1−2 : p−val = 0.771
1−3 : p−val = 0.0412
2−3 : p−val = 0.0215
Relative recombination rate
0.0
0.5
1.0
1.5
2.0
2.5
0.0
0.2
0.4
0.6
0.8
1.0
Normalized physical position

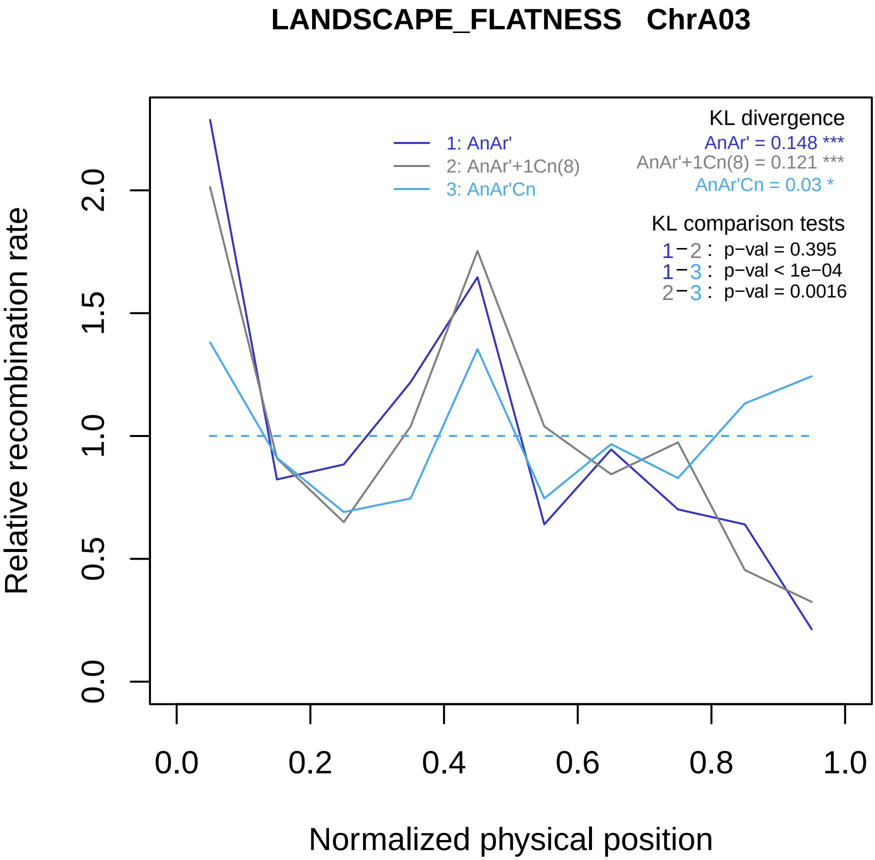

LANDSCAPE_FLATNESS ChrA03
KL divergence
AnAr' = 0.148 ***
AnAr'+1Cn(8) = 0.121 ***
AnAr'Cn = 0.03 *
1: AnAr'
2: AnAr'+1Cn(8)
3: AnAr'Cn
KL comparison tests
1−2 : p−val = 0.395
1−3 : p−val < 1e−04
2−3 : p−val = 0.0016
Relative recombination rate
0.0
0.5
1.0
1.5
2.0
0.0
0.2
0.4
0.6
0.8
1.0
Normalized physical position

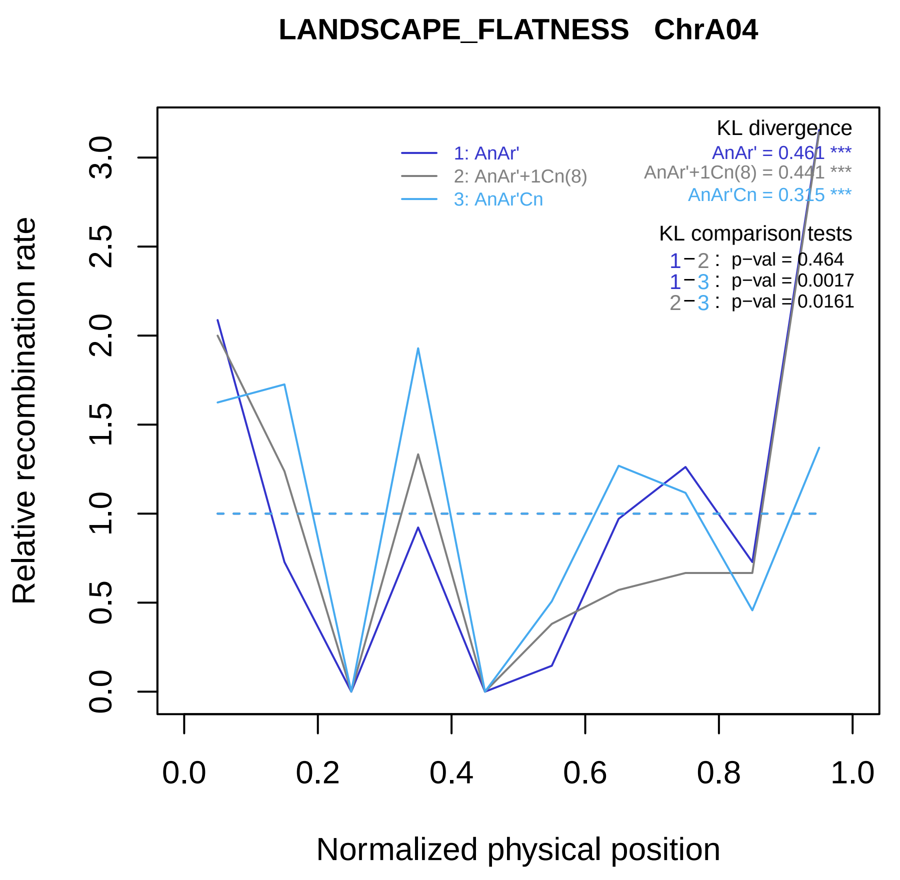
LANDSCAPE_FLATNESS ChrA04
KL divergence
AnAr' = 0.461 ***
AnAr'+1Cn(8) = 0.441 ***
AnAr'Cn = 0.315 ***
1: AnAr'
2: AnAr'+1Cn(8)
3: AnAr'Cn
KL comparison tests
1−2 : p−val = 0.464
1−3 : p−val = 0.0017
2−3 : p−val = 0.0161
Relative recombination rate
0.0
0.5
1.0
1.5
2.0
2.5
3.0
0.0
0.2
0.4
0.6
0.8
1.0
Normalized physical position

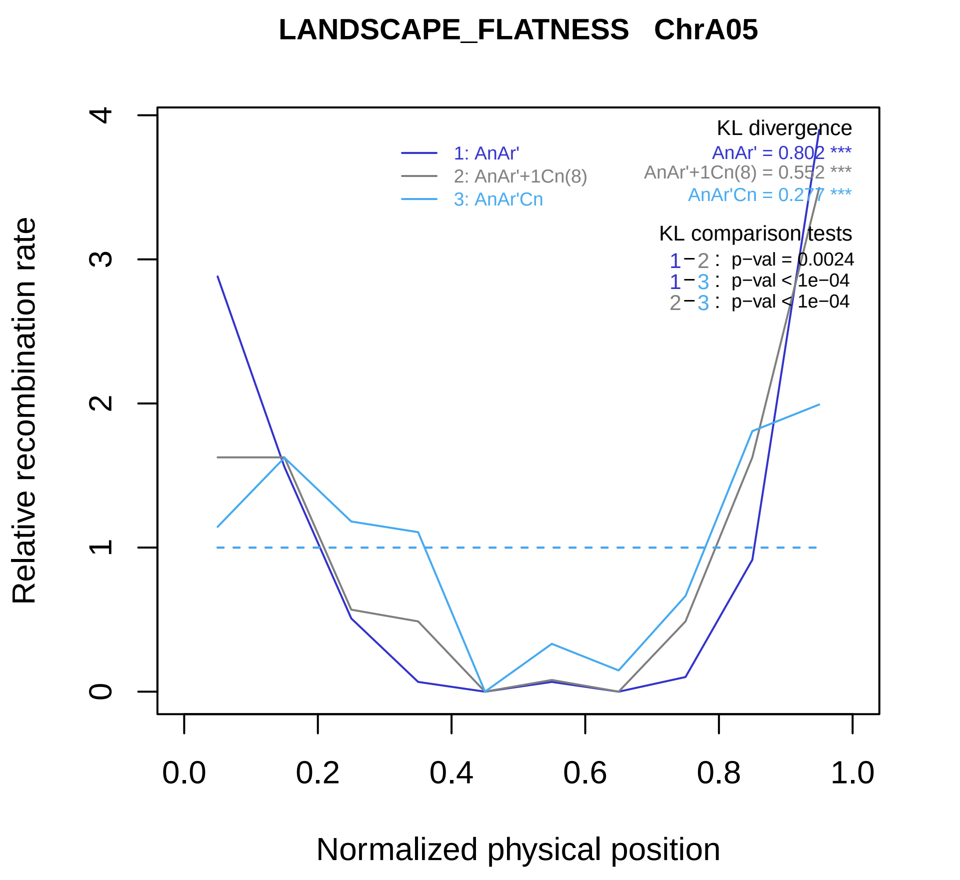
LANDSCAPE_FLATNESS ChrA05
Relative recombination rate
Normalized physical position
0
1
2
3
4
0.0
0.2
0.4
0.6
0.8
1.0
1: AnAr'
2: AnAr'+1Cn(8)
3: AnAr'Cn
KL divergence
AnAr' = 0.802 ***
AnAr'+1Cn(8) = 0.552 ***
AnAr'Cn = 0.277 ***
KL comparison tests
1−2 : p−val = 0.0024
1−3 : p−val < 1e−04
2−3 : p−val < 1e−04

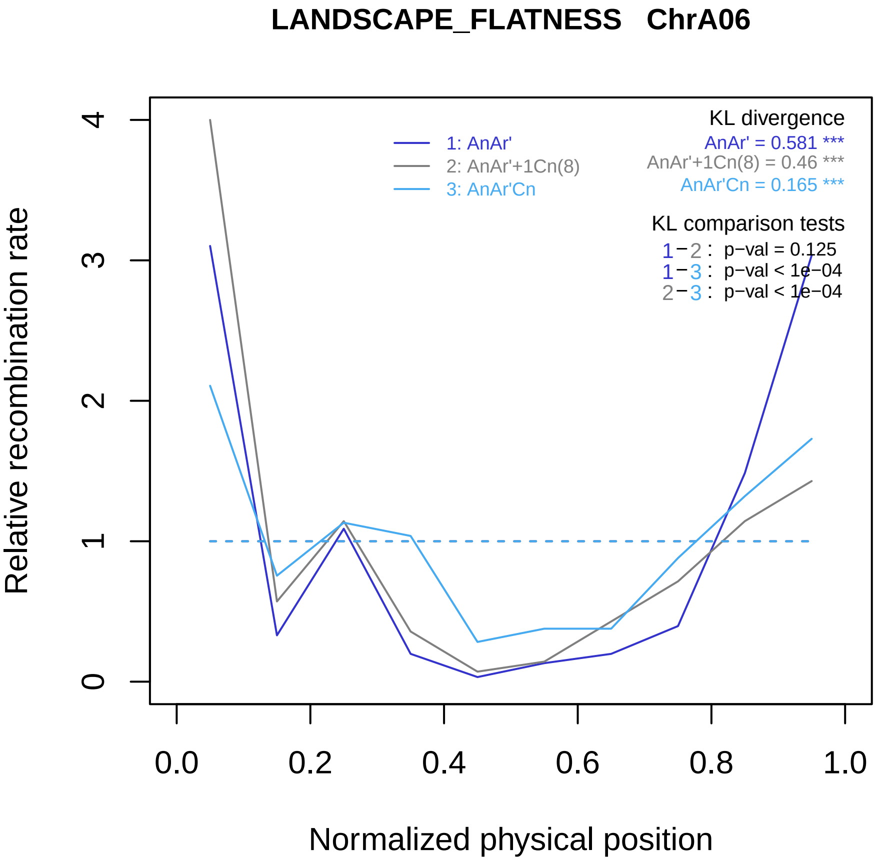
LANDSCAPE_FLATNESS ChrA06
KL divergence
AnAr' = 0.581 ***
AnAr'+1Cn(8) = 0.46 ***
AnAr'Cn = 0.165 ***
1: AnAr'
2: AnAr'+1Cn(8)
3: AnAr'Cn
KL comparison tests
1−2 : p−val = 0.125
1−3 : p−val < 1e−04
2−3 : p−val < 1e−04
Relative recombination rate
0
1
2
3
4
0.0
0.2
0.4
0.6
0.8
1.0
Normalized physical position

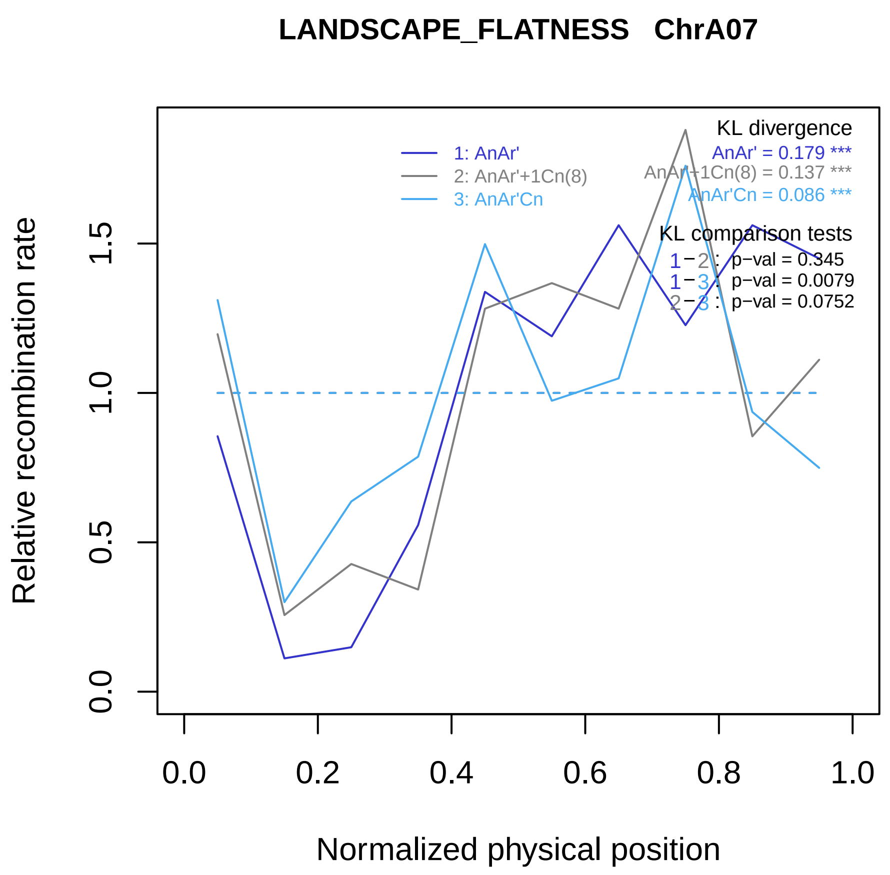

LANDSCAPE_FLATNESS ChrA07
KL divergence
AnAr' = 0.179 ***
AnAr'+1Cn(8) = 0.137 ***
AnAr'Cn = 0.086 ***
1: AnAr'
2: AnAr'+1Cn(8)
3: AnAr'Cn
KL comparison tests
1−2 : p−val = 0.345
1−3 : p−val = 0.0079
2−3 : p−val = 0.0752
Relative recombination rate
0.0
0.5
1.0
1.5
0.0
0.2
0.4
0.6
0.8
1.0
Normalized physical position

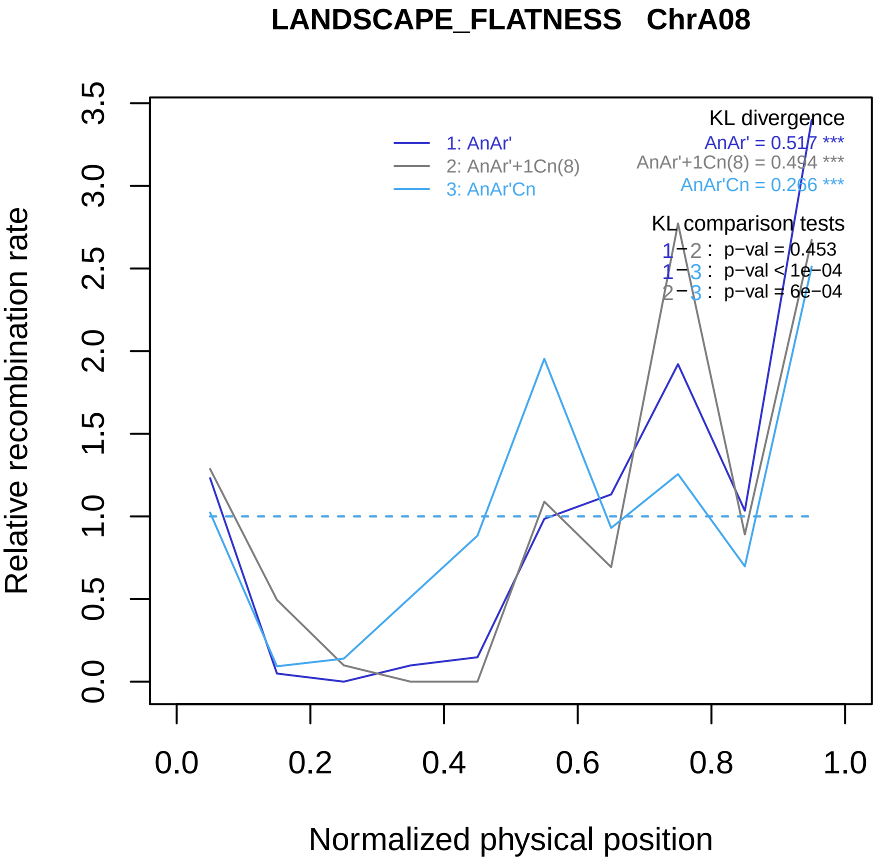

LANDSCAPE_FLATNESS ChrA08
KL divergence
AnAr' = 0.517 ***
AnAr'+1Cn(8) = 0.494 ***
AnAr'Cn = 0.266 ***
1: AnAr'
2: AnAr'+1Cn(8)
3: AnAr'Cn
KL comparison tests
1 − 2 : p−val = 0.453
1 − 3 : p−val < 1e−04
2 − 3 : p−val = 6e−04
Relative recombination rate
0.0
0.5
1.0
1.5
2.0
2.5
3.0
3.5
0.0
0.2
0.4
0.6
0.8
1.0
Normalized physical position

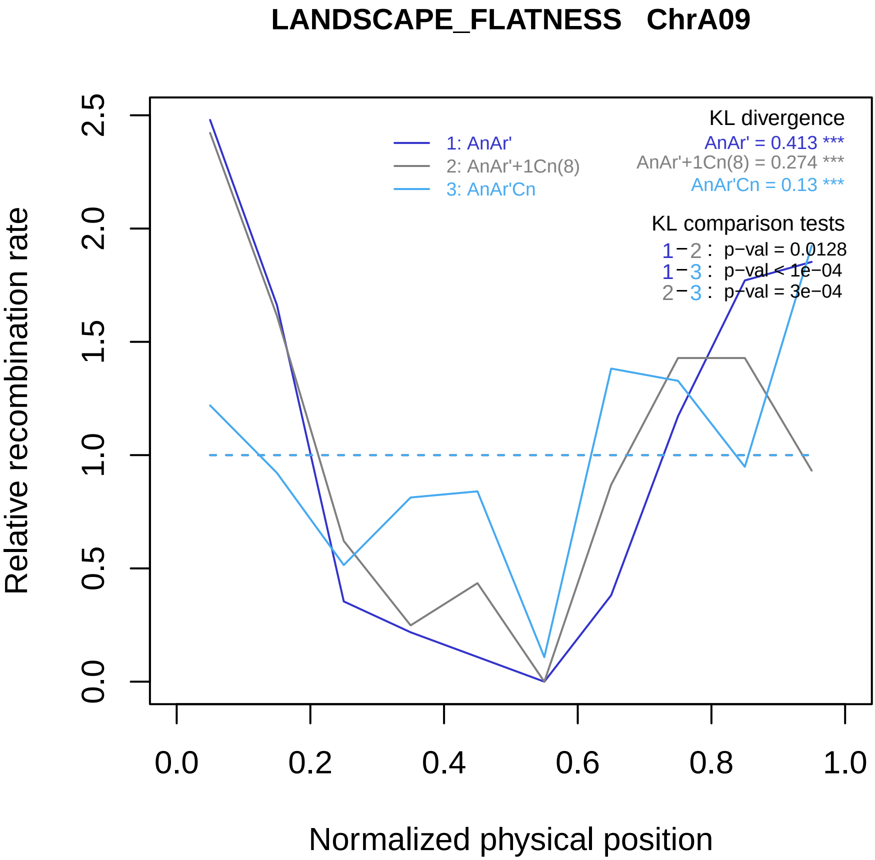

LANDSCAPE_FLATNESS ChrA09
KL divergence
AnAr' = 0.413 ***
AnAr'+1Cn(8) = 0.274 ***
AnAr'Cn = 0.13 ***
1: AnAr'
2: AnAr'+1Cn(8)
3: AnAr'Cn
KL comparison tests
1−2 : p−val = 0.0128
1−3 : p−val < 1e−04
2−3 : p−val = 3e−04
Relative recombination rate
0.0
0.5
1.0
1.5
2.0
2.5
0.0
0.2
0.4
0.6
0.8
1.0
Normalized physical position

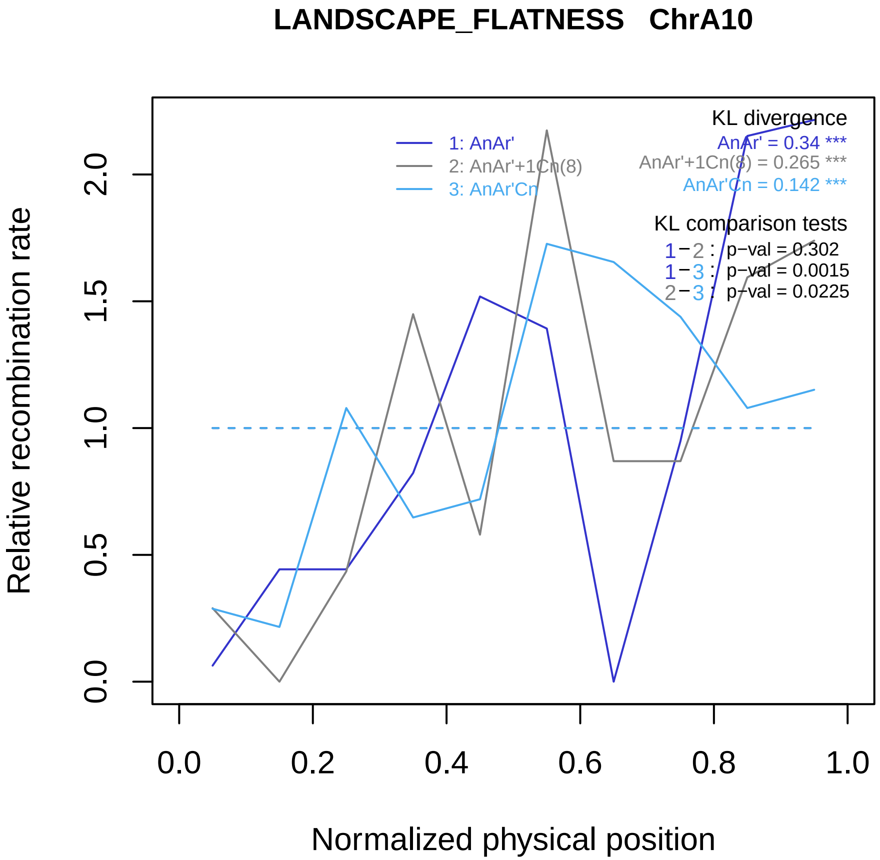
LANDSCAPE_FLATNESS ChrA10
Relative recombination rate
0.0
0.5
1.0
1.5
2.0
Normalized physical position
0.0
0.2
0.4
0.6
0.8
1.0
1: AnAr'
2: AnAr'+1Cn(8)
3: AnAr'Cn
KL divergence
AnAr' = 0.34 ***
AnAr'+1Cn(8) = 0.265 ***
AnAr'Cn = 0.142 ***
KL comparison tests
1−2 : p−val = 0.302
1−3 : p−val = 0.0015
2−3 : p−val = 0.0225

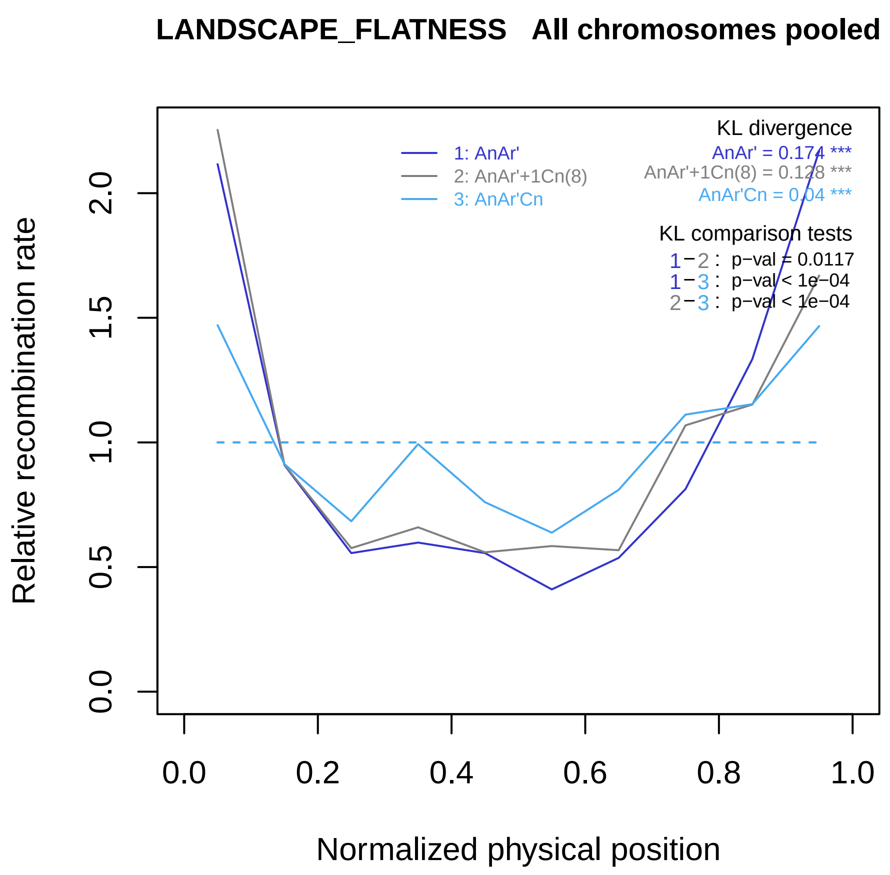

LANDSCAPE_FLATNESS   All chromosomes pooled
KL divergence
AnAr' = 0.174 ***
AnAr'+1Cn(8) = 0.128 ***
AnAr'Cn = 0.04 ***
1: AnAr'
2: AnAr'+1Cn(8)
3: AnAr'Cn
KL comparison tests
1−2 : p−val = 0.0117
1−3 : p−val < 1e−04
2−3 : p−val < 1e−04
Relative recombination rate
0.0
0.5
1.0
1.5
2.0
0.0
0.2
0.4
0.6
0.8
1.0
Normalized physical position

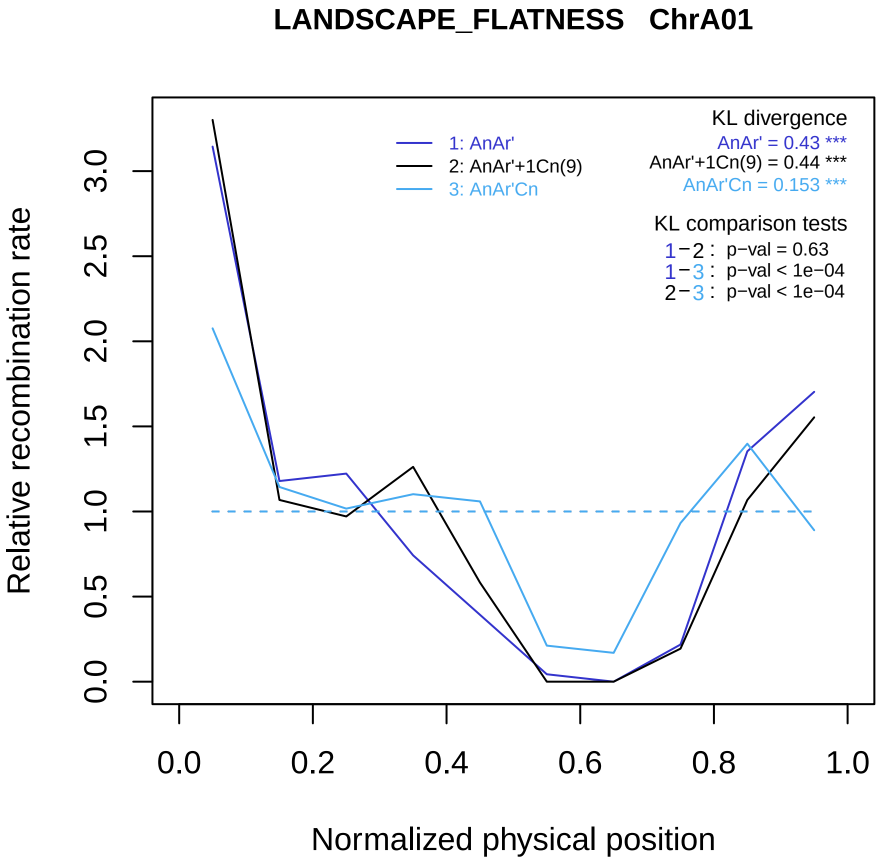

LANDSCAPE_FLATNESS ChrA01
KL divergence
AnAr' = 0.43 ***
AnAr'+1Cn(9) = 0.44 ***
AnAr'Cn = 0.153 ***
1: AnAr'
2: AnAr'+1Cn(9)
3: AnAr'Cn
KL comparison tests
1−2 : p−val = 0.63
1−3 : p−val < 1e−04
2−3 : p−val < 1e−04
Relative recombination rate
0.0
0.5
1.0
1.5
2.0
2.5
3.0
0.0
0.2
0.4
0.6
0.8
1.0
Normalized physical position

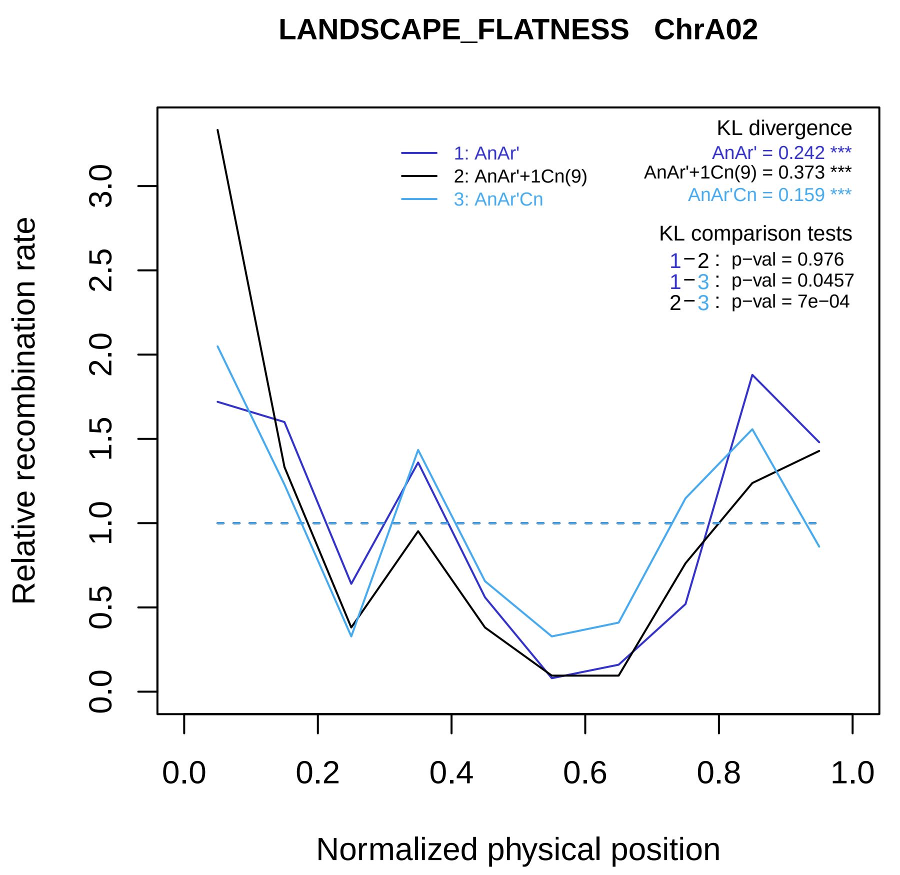

LANDSCAPE_FLATNESS ChrA02
KL divergence
AnAr' = 0.242 ***
AnAr'+1Cn(9) = 0.373 ***
AnAr'Cn = 0.159 ***
1: AnAr'
2: AnAr'+1Cn(9)
3: AnAr'Cn
KL comparison tests
1−2 : p−val = 0.976
1−3 : p−val = 0.0457
2−3 : p−val = 7e−04
Relative recombination rate
0.0
0.5
1.0
1.5
2.0
2.5
3.0
0.0
0.2
0.4
0.6
0.8
1.0
Normalized physical position

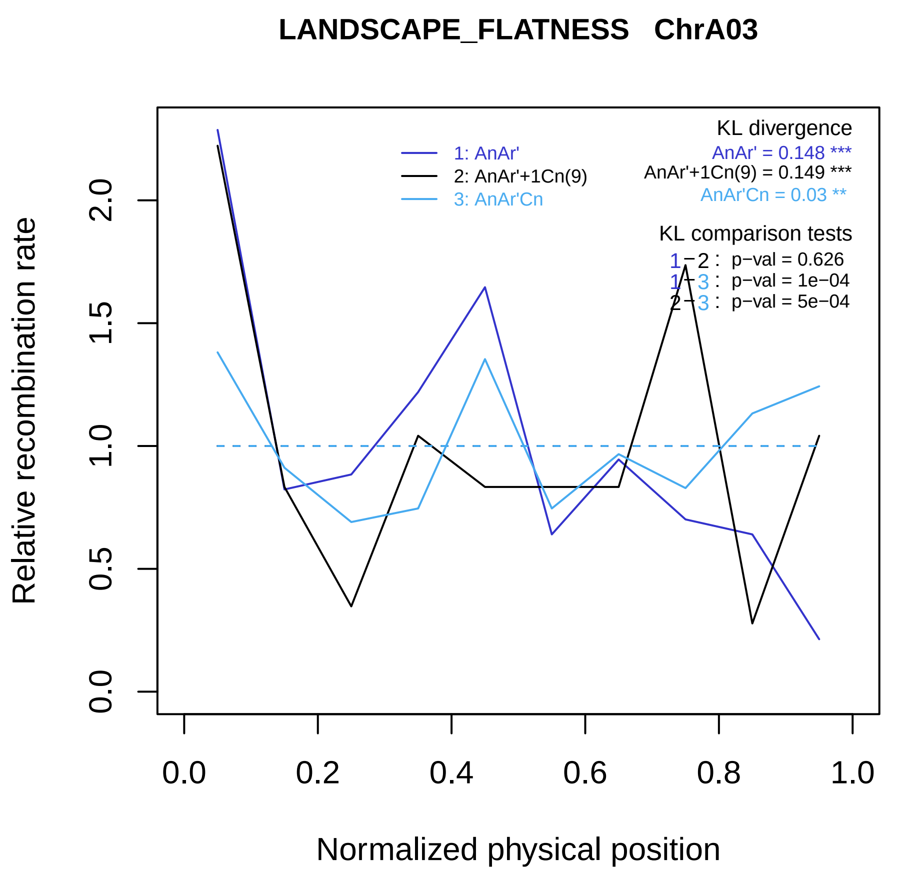
LANDSCAPE_FLATNESS ChrA03
1: AnAr'
2: AnAr'+1Cn(9)
3: AnAr'Cn
KL divergence
AnAr' = 0.148 ***
AnAr'+1Cn(9) = 0.149 ***
AnAr'Cn = 0.03 **
KL comparison tests
1−2 : p−val = 0.626
1−3 : p−val = 1e−04
2−3 : p−val = 5e−04
Relative recombination rate
0.0
0.5
1.0
1.5
2.0
0.0
0.2
0.4
0.6
0.8
1.0
Normalized physical position

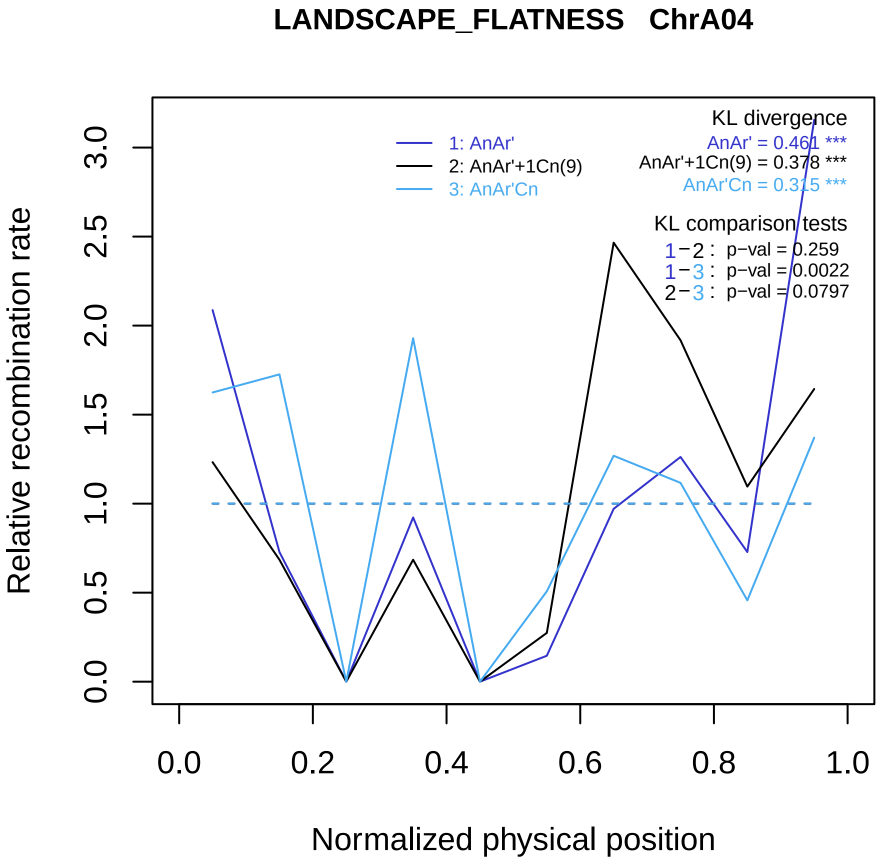
LANDSCAPE_FLATNESS ChrA04
KL divergence
AnAr' = 0.461 ***
AnAr'+1Cn(9) = 0.378 ***
AnAr'Cn = 0.315 ***
1: AnAr'
2: AnAr'+1Cn(9)
3: AnAr'Cn
KL comparison tests
1−2 : p−val = 0.259
1−3 : p−val = 0.0022
2−3 : p−val = 0.0797
Relative recombination rate
0.0
0.5
1.0
1.5
2.0
2.5
3.0
0.0
0.2
0.4
0.6
0.8
1.0
Normalized physical position

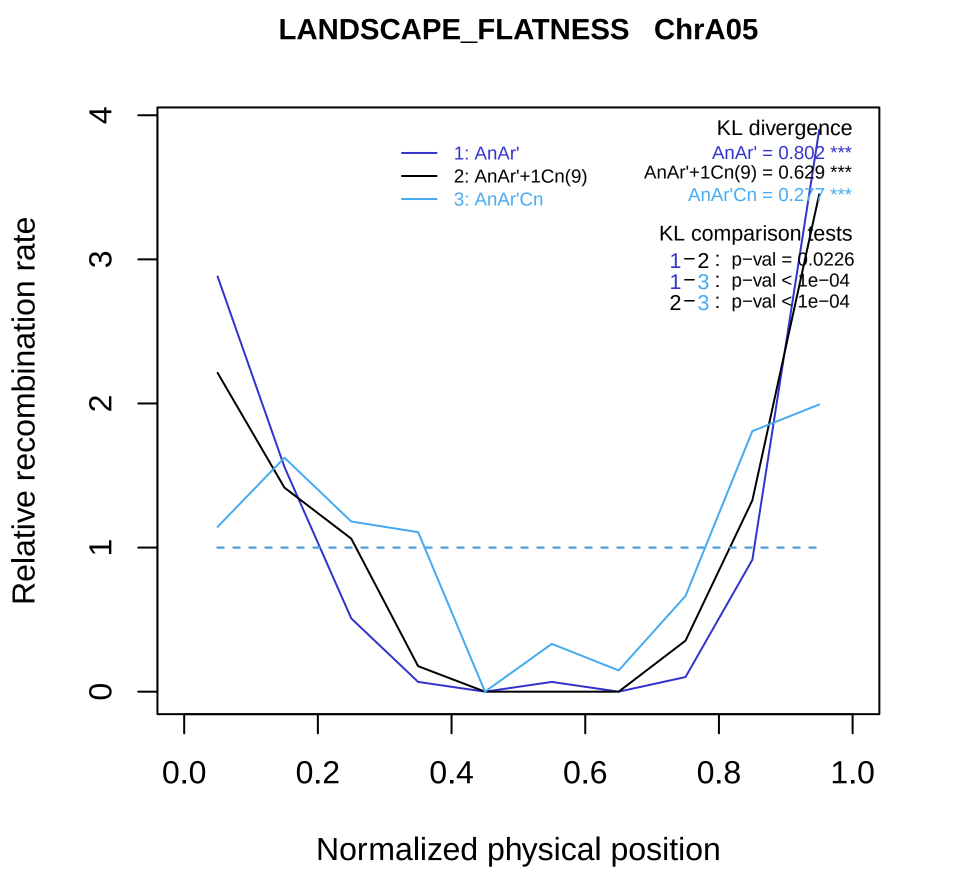

LANDSCAPE_FLATNESS ChrA05
Relative recombination rate
Normalized physical position
0
1
2
3
4
0.0
0.2
0.4
0.6
0.8
1.0
1: AnAr'
2: AnAr'+1Cn(9)
3: AnAr'Cn
KL divergence
AnAr' = 0.802 ***
AnAr'+1Cn(9) = 0.629 ***
AnAr'Cn = 0.277 ***
KL comparison tests
1−2 : p−val = 0.0226
1−3 : p−val < 1e−04
2−3 : p−val < 1e−04

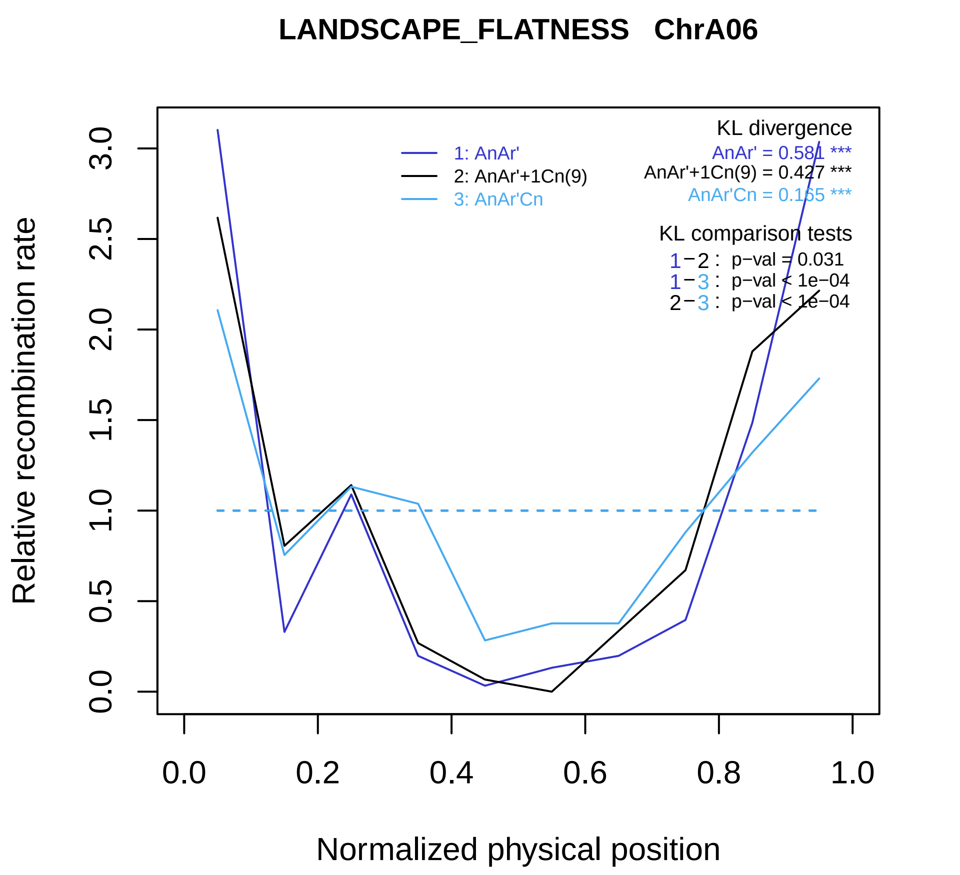
LANDSCAPE_FLATNESS ChrA06
Relative recombination rate
Normalized physical position
0.0
0.5
1.0
1.5
2.0
2.5
3.0
0.0
0.2
0.4
0.6
0.8
1.0
1: AnAr'
2: AnAr'+1Cn(9)
3: AnAr'Cn
KL divergence
AnAr' = 0.581 ***
AnAr'+1Cn(9) = 0.427 ***
AnAr'Cn = 0.165 ***
KL comparison tests
1−2 : p−val = 0.031
1−3 : p−val < 1e−04
2−3 : p−val < 1e−04

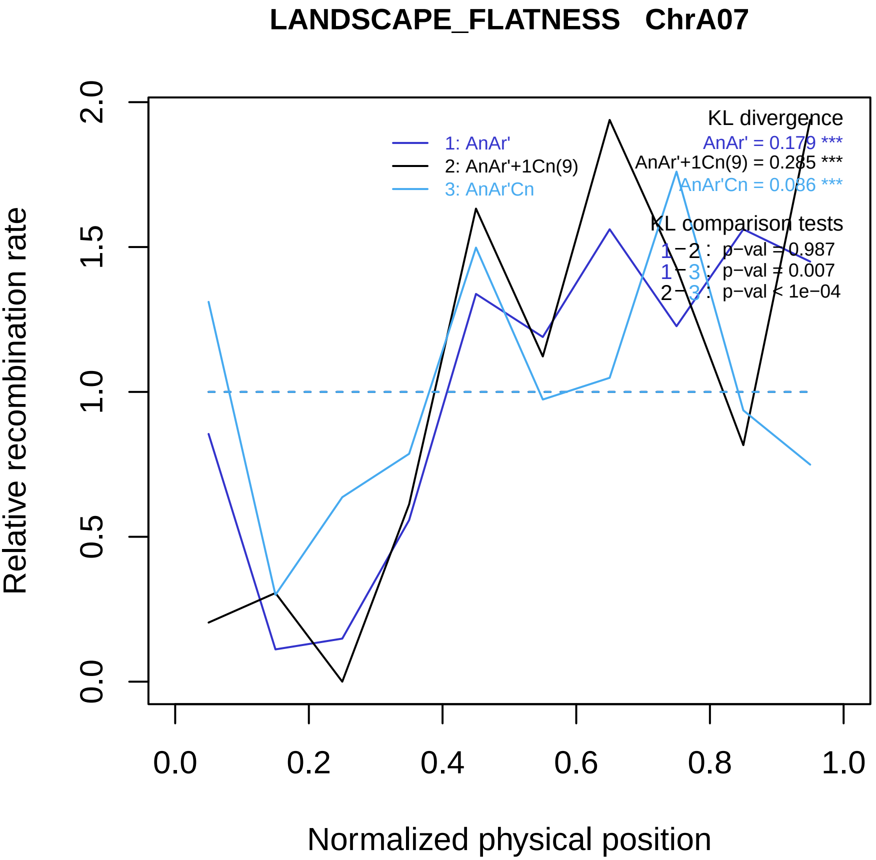
LANDSCAPE_FLATNESS ChrA07
KL divergence
AnAr' = 0.179 ***
AnAr'+1Cn(9) = 0.285 ***
AnAr'Cn = 0.086 ***
1: AnAr'
2: AnAr'+1Cn(9)
3: AnAr'Cn
KL comparison tests
1−2 : p−val = 0.987
1−3 : p−val = 0.007
2−3 : p−val < 1e−04
Relative recombination rate
0.0
0.5
1.0
1.5
2.0
0.0
0.2
0.4
0.6
0.8
1.0
Normalized physical position

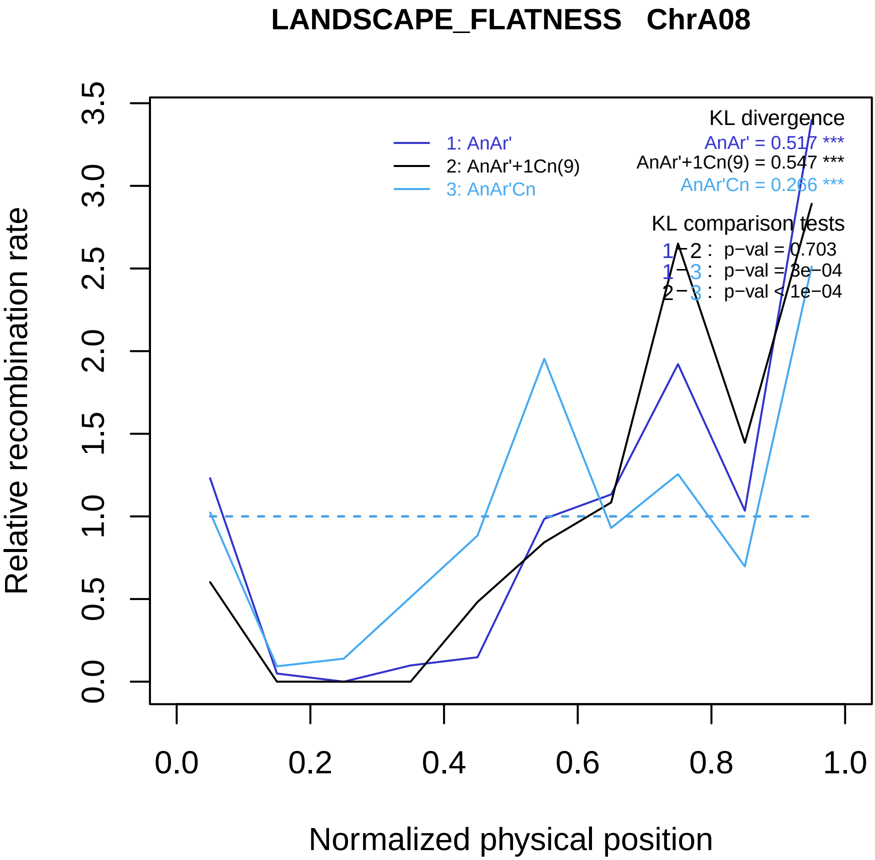

LANDSCAPE_FLATNESS ChrA08
KL divergence
AnAr' = 0.517 ***
AnAr'+1Cn(9) = 0.547 ***
AnAr'Cn = 0.266 ***
1: AnAr'
2: AnAr'+1Cn(9)
3: AnAr'Cn
KL comparison tests
1−2 : p−val = 0.703
1−3 : p−val = 3e−04
2−3 : p−val < 1e−04
Relative recombination rate
0.0
0.5
1.0
1.5
2.0
2.5
3.0
3.5
0.0
0.2
0.4
0.6
0.8
1.0
Normalized physical position

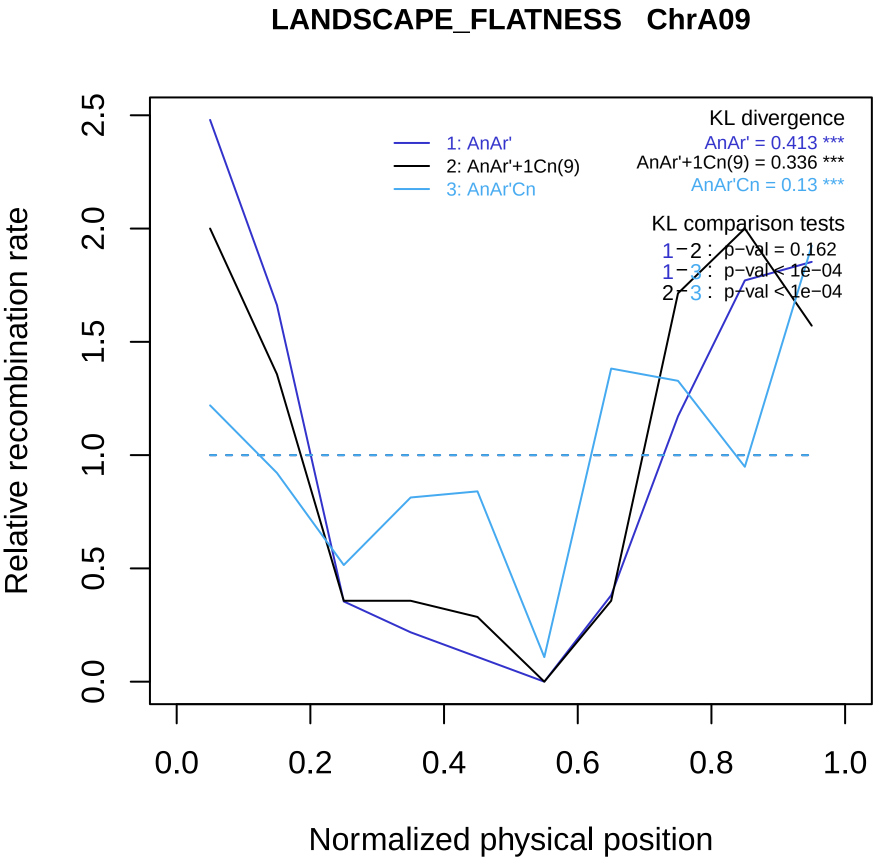

LANDSCAPE_FLATNESS ChrA09
Relative recombination rate
Normalized physical position
0.0
0.5
1.0
1.5
2.0
2.5
0.0
0.2
0.4
0.6
0.8
1.0
1: AnAr'
2: AnAr'+1Cn(9)
3: AnAr'Cn
KL divergence
AnAr' = 0.413 ***
AnAr'+1Cn(9) = 0.336 ***
AnAr'Cn = 0.13 ***
KL comparison tests
1−2 : p−val = 0.162
1−3 : p−val < 1e−04
2−3 : p−val < 1e−04

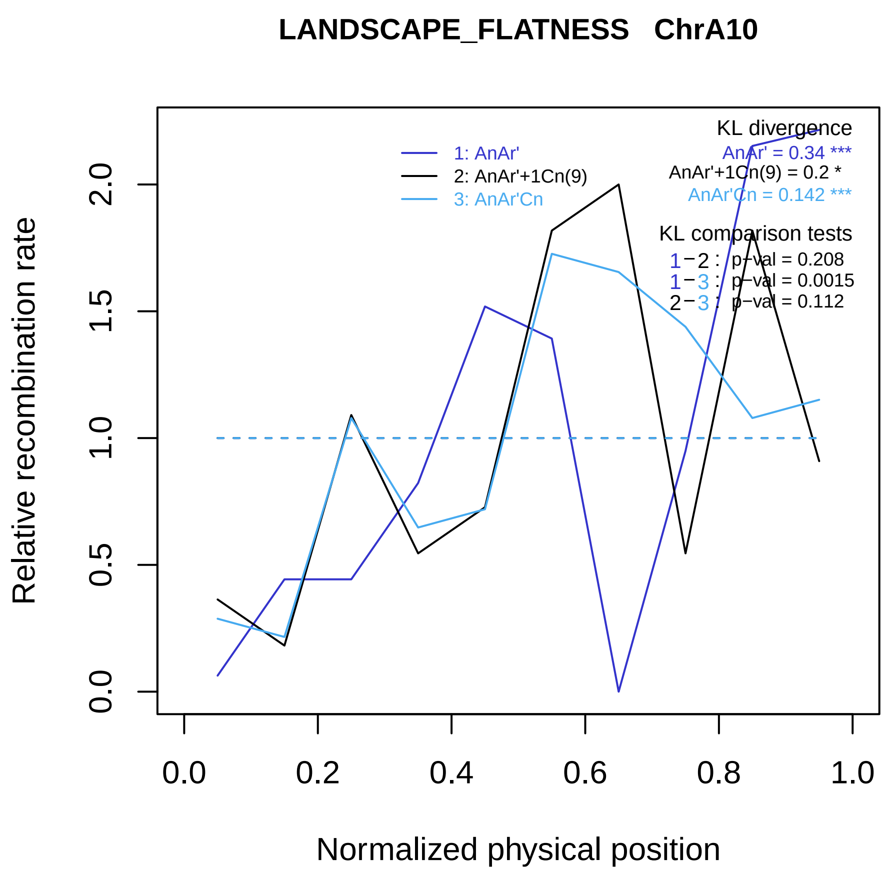

LANDSCAPE_FLATNESS   ChrA10
1: AnAr'
2: AnAr'+1Cn(9)
3: AnAr'Cn
KL divergence
AnAr' = 0.34 ***
AnAr'+1Cn(9) = 0.2 *
AnAr'Cn = 0.142 ***
KL comparison tests
1−2 : p−val = 0.208
1−3 : p−val = 0.0015
2−3 : p−val = 0.112
Relative recombination rate
0.0
0.5
1.0
1.5
2.0
0.0
0.2
0.4
0.6
0.8
1.0
Normalized physical position

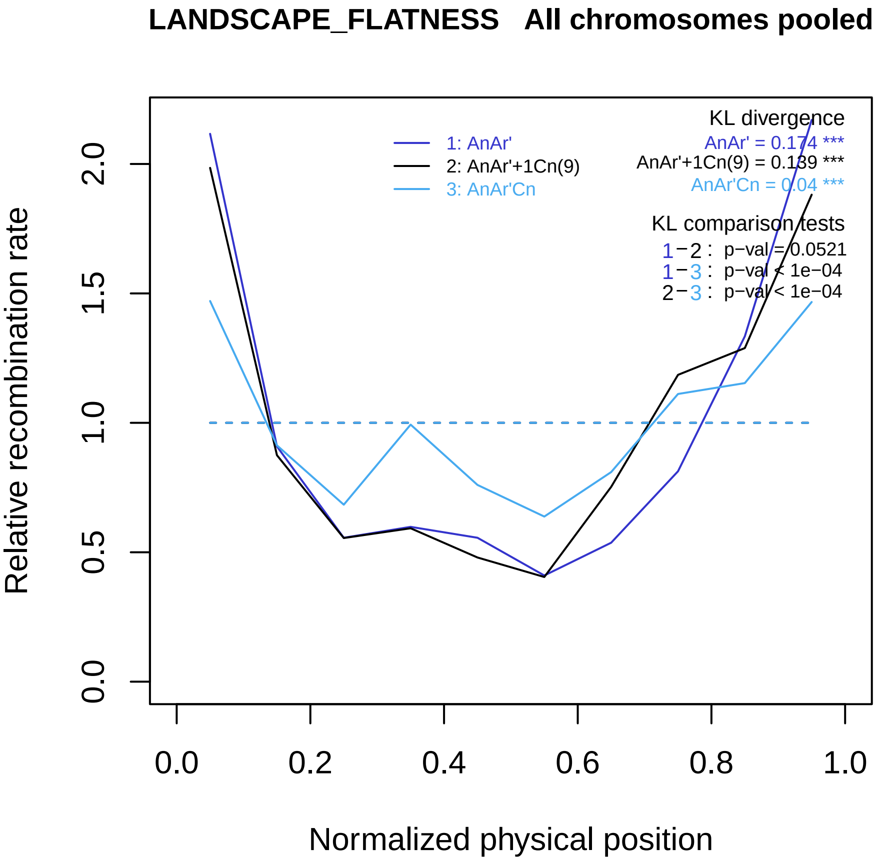

LANDSCAPE_FLATNESS All chromosomes pooled
Relative recombination rate
Normalized physical position
0.0
0.5
1.0
1.5
2.0
0.0
0.2
0.4
0.6
0.8
1.0
1: AnAr'
2: AnAr'+1Cn(9)
3: AnAr'Cn
KL divergence
AnAr' = 0.174 ***
AnAr'+1Cn(9) = 0.139 ***
AnAr'Cn = 0.04 ***
KL comparison tests
1−2 : p−val = 0.0521
1−3 : p−val < 1e−04
2−3 : p−val < 1e−04

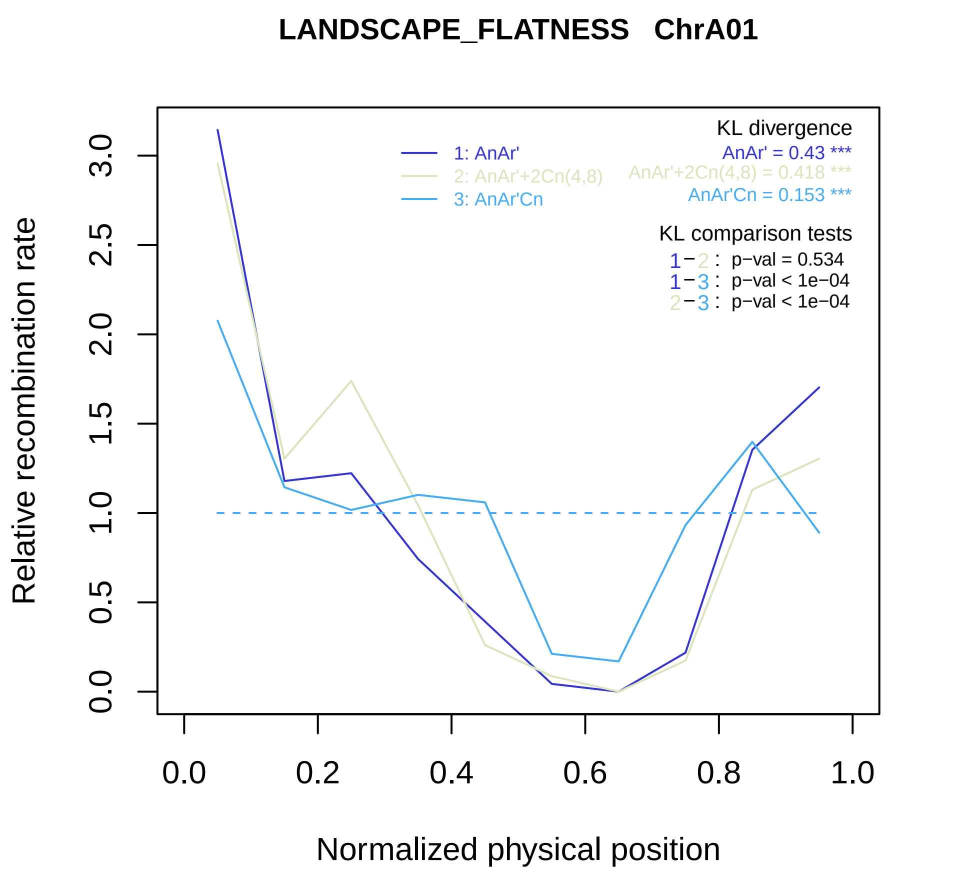
LANDSCAPE_FLATNESS ChrA01
KL divergence
AnAr' = 0.43 ***
AnAr'+2Cn(4,8) = 0.418 ***
AnAr'Cn = 0.153 ***
1: AnAr'
2: AnAr'+2Cn(4,8)
3: AnAr'Cn
KL comparison tests
1−2 : p−val = 0.534
1−3 : p−val < 1e−04
2−3 : p−val < 1e−04
Relative recombination rate
Normalized physical position
0.0
0.5
1.0
1.5
2.0
2.5
3.0
0.0
0.2
0.4
0.6
0.8
1.0

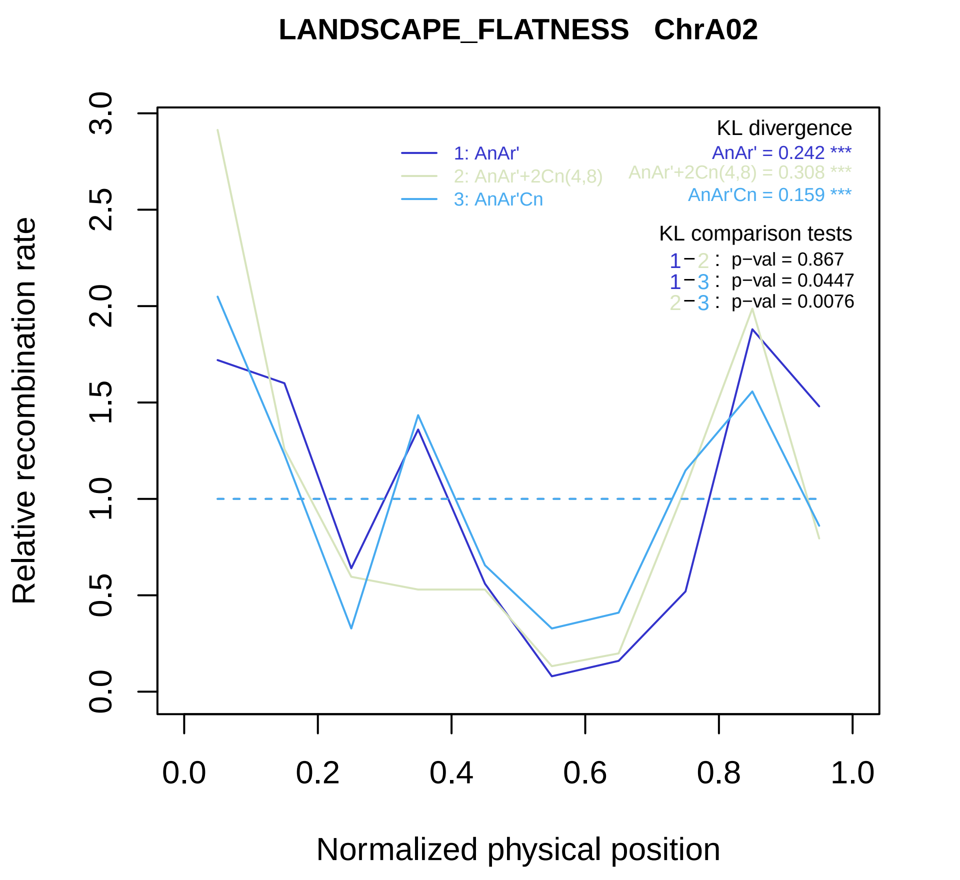

LANDSCAPE_FLATNESS ChrA02
KL divergence
1: AnAr'
2: AnAr'+2Cn(4,8)
3: AnAr'Cn
AnAr' = 0.242 ***
AnAr'+2Cn(4,8) = 0.308 ***
AnAr'Cn = 0.159 ***
KL comparison tests
1−2 : p−val = 0.867
1−3 : p−val = 0.0447
2−3 : p−val = 0.0076
Relative recombination rate
0.0
0.5
1.0
1.5
2.0
2.5
3.0
0.0
0.2
0.4
0.6
0.8
1.0
Normalized physical position

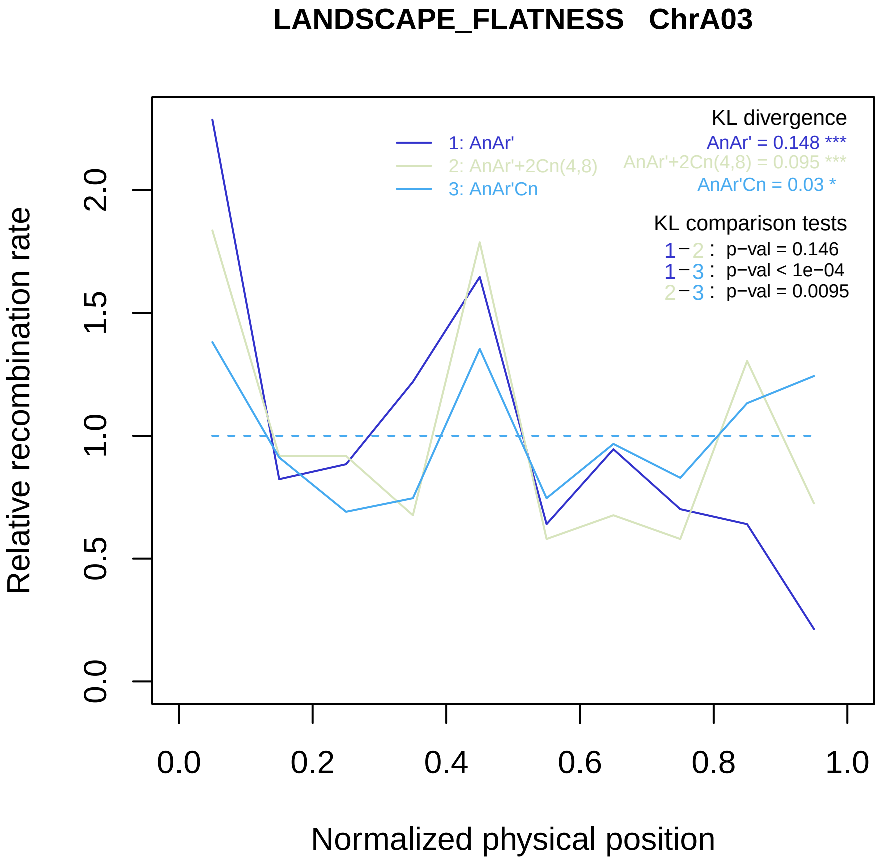

LANDSCAPE_FLATNESS ChrA03
KL divergence
AnAr' = 0.148 ***
AnAr'+2Cn(4,8) = 0.095 ***
AnAr'Cn = 0.03 *
1: AnAr'
2: AnAr'+2Cn(4,8)
3: AnAr'Cn
KL comparison tests
1 − 2 : p−val = 0.146
1 − 3 : p−val < 1e−04
2 − 3 : p−val = 0.0095
Relative recombination rate
0.0
0.5
1.0
1.5
2.0
0.0
0.2
0.4
0.6
0.8
1.0
Normalized physical position

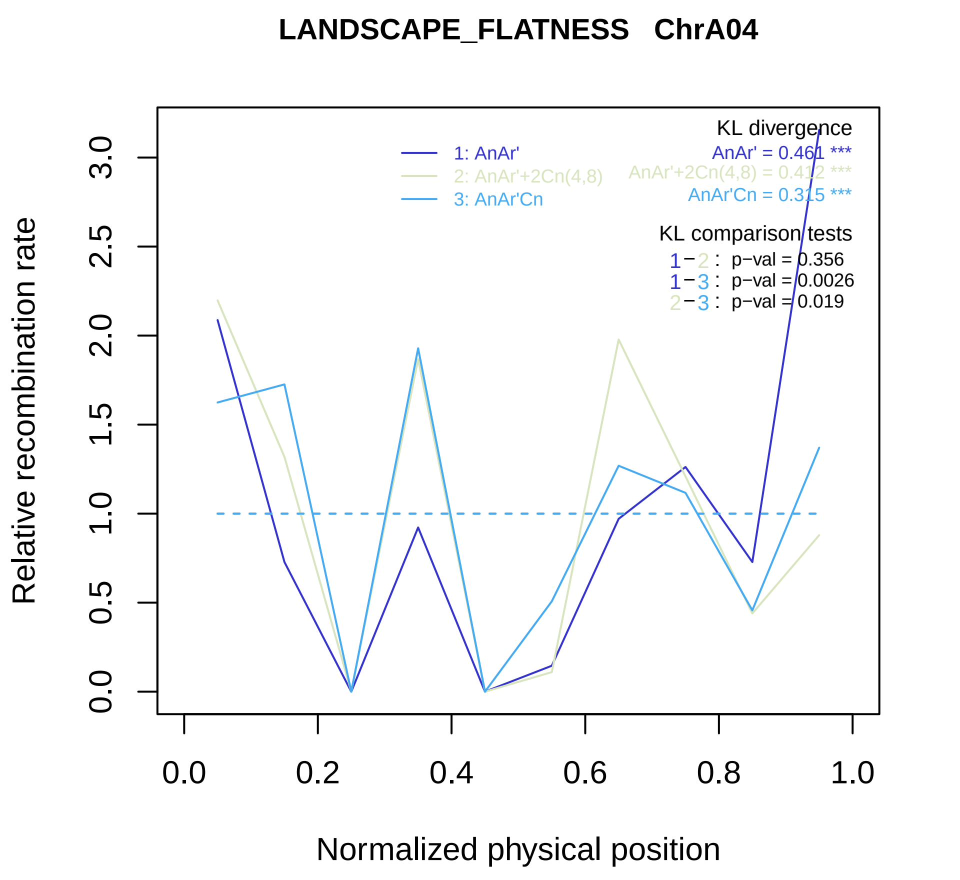

LANDSCAPE_FLATNESS ChrA04
Relative recombination rate
Normalized physical position
0.0
0.5
1.0
1.5
2.0
2.5
3.0
0.0
0.2
0.4
0.6
0.8
1.0
1: AnAr'
2: AnAr'+2Cn(4,8)
3: AnAr'Cn
KL divergence
AnAr' = 0.461 ***
AnAr'+2Cn(4,8) = 0.412 ***
AnAr'Cn = 0.315 ***
KL comparison tests
1−2 : p−val = 0.356
1−3 : p−val = 0.0026
2−3 : p−val = 0.019

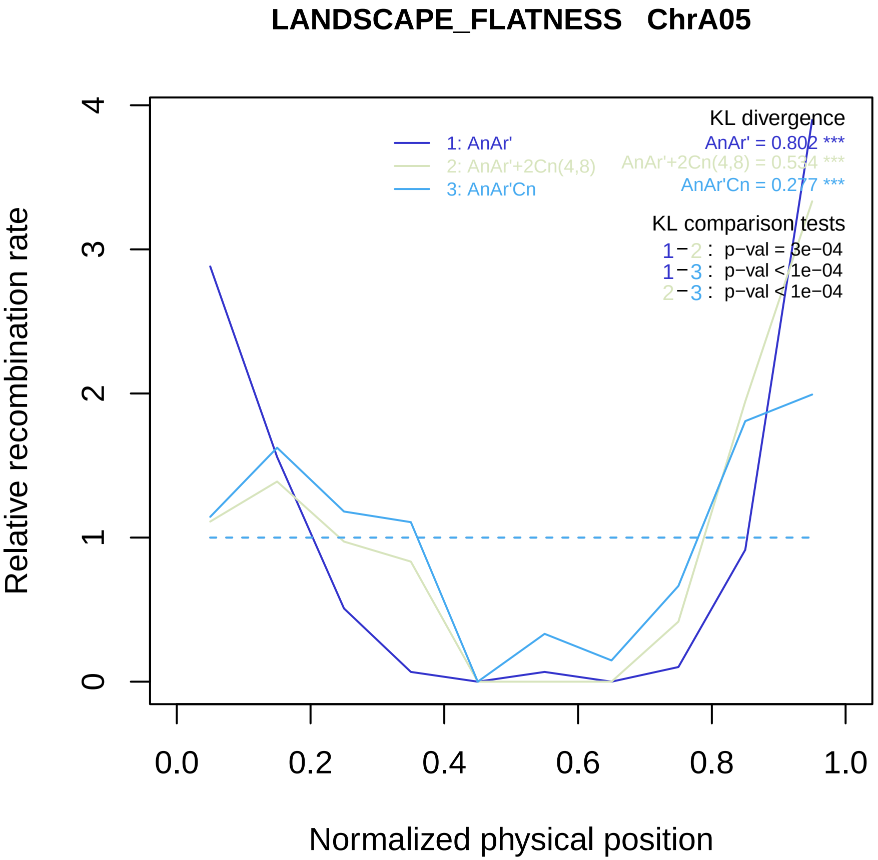
LANDSCAPE_FLATNESS ChrA05
Relative recombination rate
Normalized physical position
0
1
2
3
4
0.0
0.2
0.4
0.6
0.8
1.0
1: AnAr'
2: AnAr'+2Cn(4,8)
3: AnAr'Cn
KL divergence
AnAr' = 0.802 ***
AnAr'+2Cn(4,8) = 0.534 ***
AnAr'Cn = 0.277 ***
KL comparison tests
1−2 : p−val = 3e−04
1−3 : p−val < 1e−04
2−3 : p−val < 1e−04

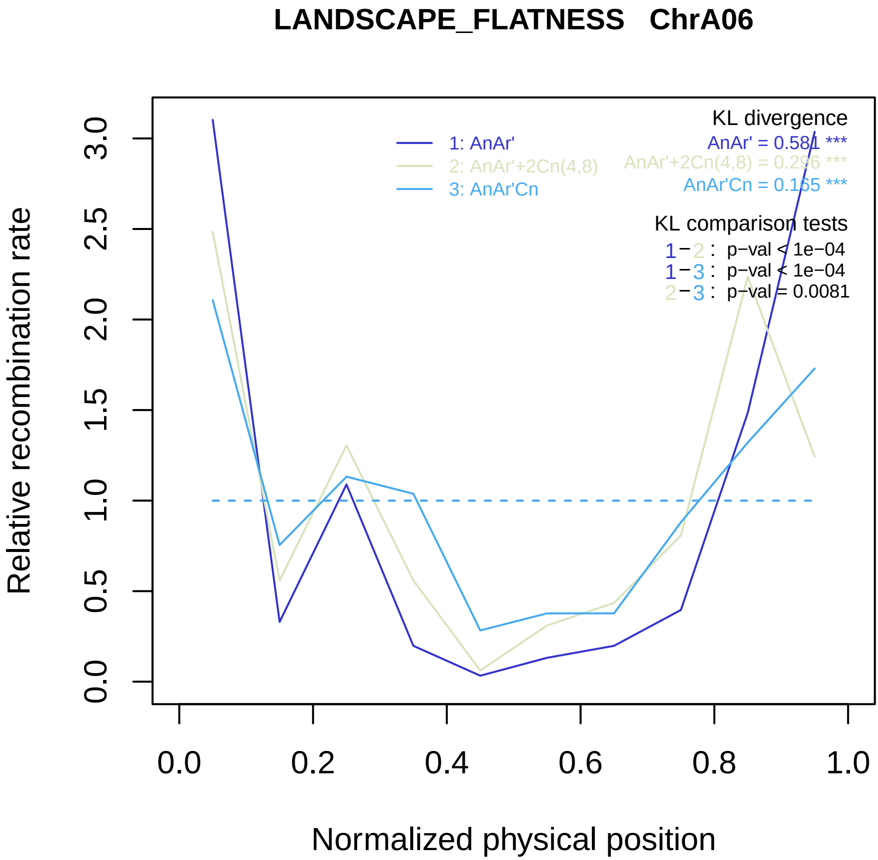
LANDSCAPE_FLATNESS ChrA06
KL divergence
AnAr' = 0.581 ***
AnAr'+2Cn(4,8) = 0.296 ***
AnAr'Cn = 0.165 ***
1: AnAr'
2: AnAr'+2Cn(4,8)
3: AnAr'Cn
KL comparison tests
1−2 : p−val < 1e−04
1−3 : p−val < 1e−04
2−3 : p−val = 0.0081
Relative recombination rate
0.0
0.5
1.0
1.5
2.0
2.5
3.0
0.0
0.2
0.4
0.6
0.8
1.0
Normalized physical position

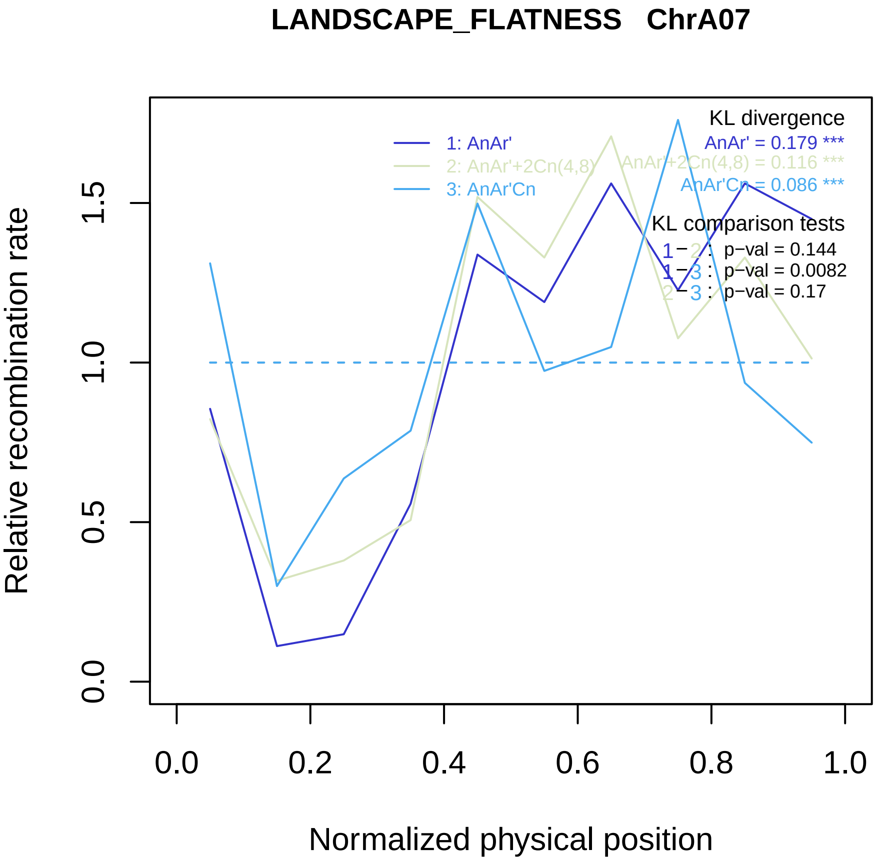

LANDSCAPE_FLATNESS ChrA07
Relative recombination rate
0.0
0.5
1.0
1.5
Normalized physical position
0.0
0.2
0.4
0.6
0.8
1.0
1: AnAr'
2: AnAr'+2Cn(4,8)
3: AnAr'Cn
KL divergence
AnAr' = 0.179 ***
AnAr'+2Cn(4,8) = 0.116 ***
AnAr'Cn = 0.086 ***
KL comparison tests
1 − 2 : p−val = 0.144
1 − 3 : p−val = 0.0082
2 − 3 : p−val = 0.17

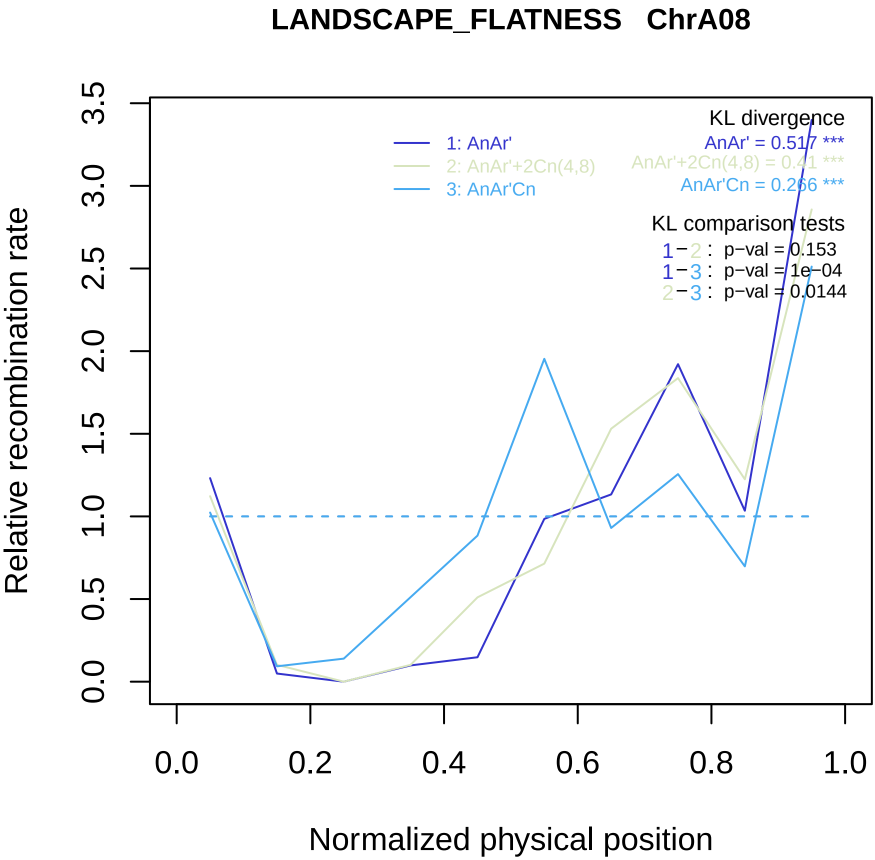

LANDSCAPE_FLATNESS ChrA08
KL divergence
AnAr' = 0.517 ***
AnAr'+2Cn(4,8) = 0.41 ***
AnAr'Cn = 0.266 ***
1: AnAr'
2: AnAr'+2Cn(4,8)
3: AnAr'Cn
KL comparison tests
1−2 : p−val = 0.153
1−3 : p−val = 1e−04
2−3 : p−val = 0.0144
Relative recombination rate
0.0
0.5
1.0
1.5
2.0
2.5
3.0
3.5
0.0
0.2
0.4
0.6
0.8
1.0
Normalized physical position

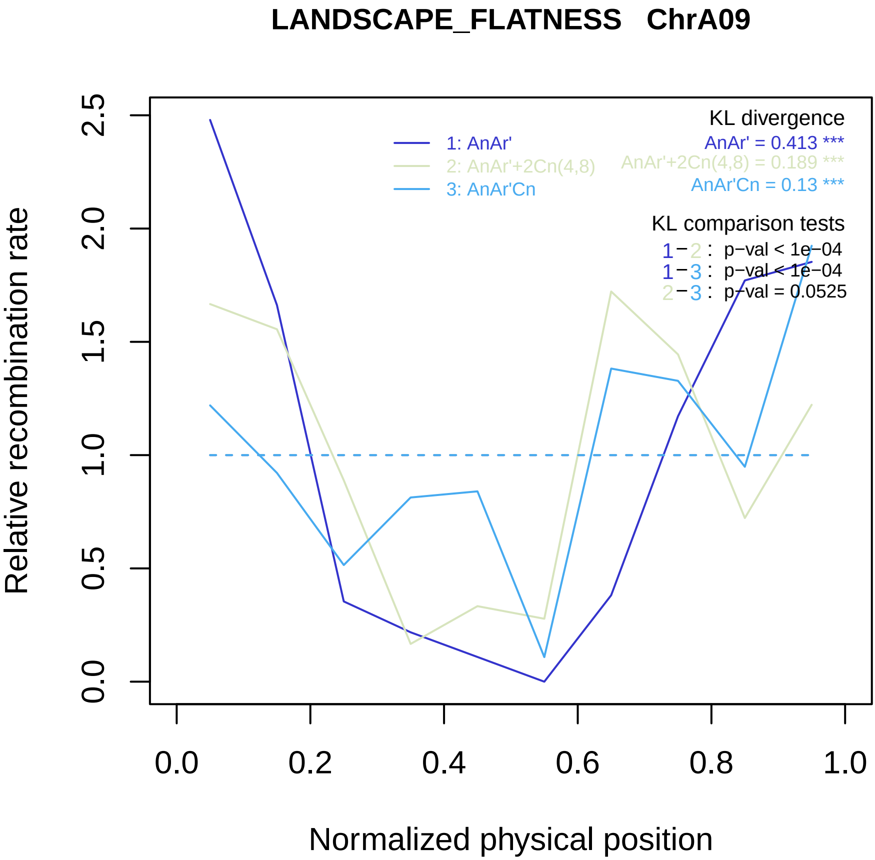
LANDSCAPE_FLATNESS ChrA09
KL divergence
AnAr' = 0.413 ***
AnAr'+2Cn(4,8) = 0.189 ***
AnAr'Cn = 0.13 ***
1: AnAr'
2: AnAr'+2Cn(4,8)
3: AnAr'Cn
KL comparison tests
1−2 : p−val < 1e−04
1−3 : p−val < 1e−04
2−3 : p−val = 0.0525
Relative recombination rate
0.0
0.5
1.0
1.5
2.0
2.5
0.0
0.2
0.4
0.6
0.8
1.0
Normalized physical position

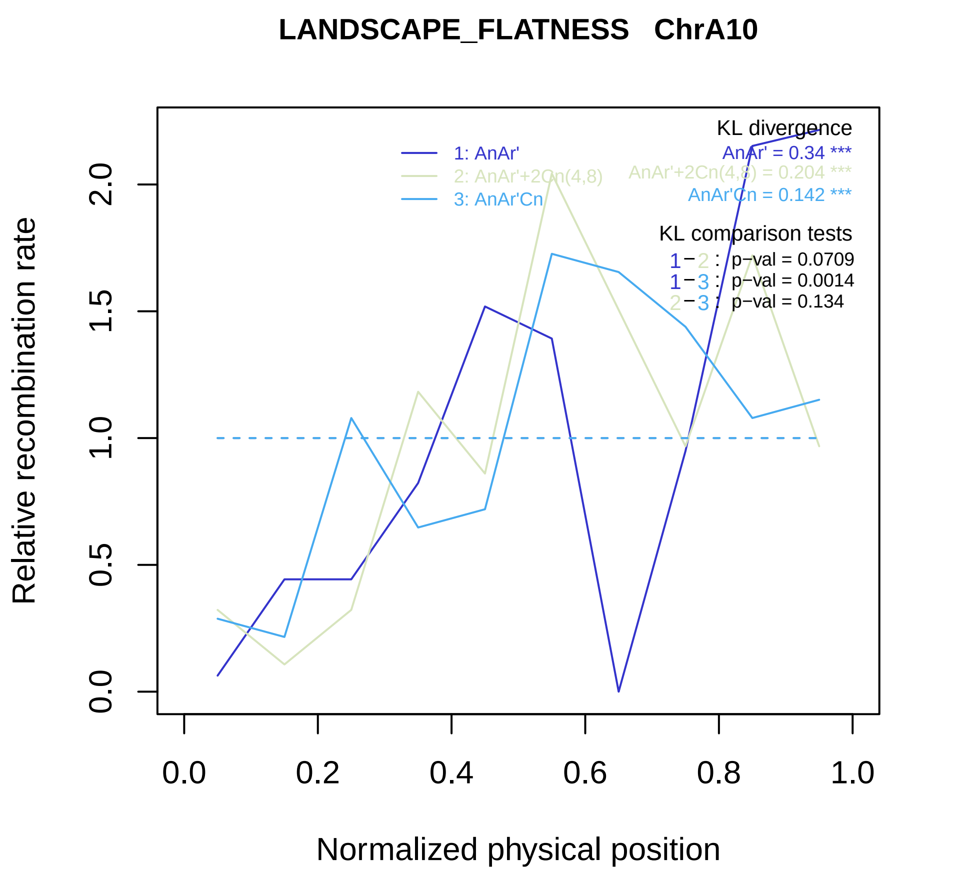
LANDSCAPE_FLATNESS ChrA10
Relative recombination rate
Normalized physical position
0.0
0.5
1.0
1.5
2.0
0.0
0.2
0.4
0.6
0.8
1.0
1: AnAr'
2: AnAr'+2Cn(4,8)
3: AnAr'Cn
KL divergence
AnAr' = 0.34 ***
AnAr'+2Cn(4,8) = 0.204 ***
AnAr'Cn = 0.142 ***
KL comparison tests
1−2 : p−val = 0.0709
1−3 : p−val = 0.0014
2−3 : p−val = 0.134

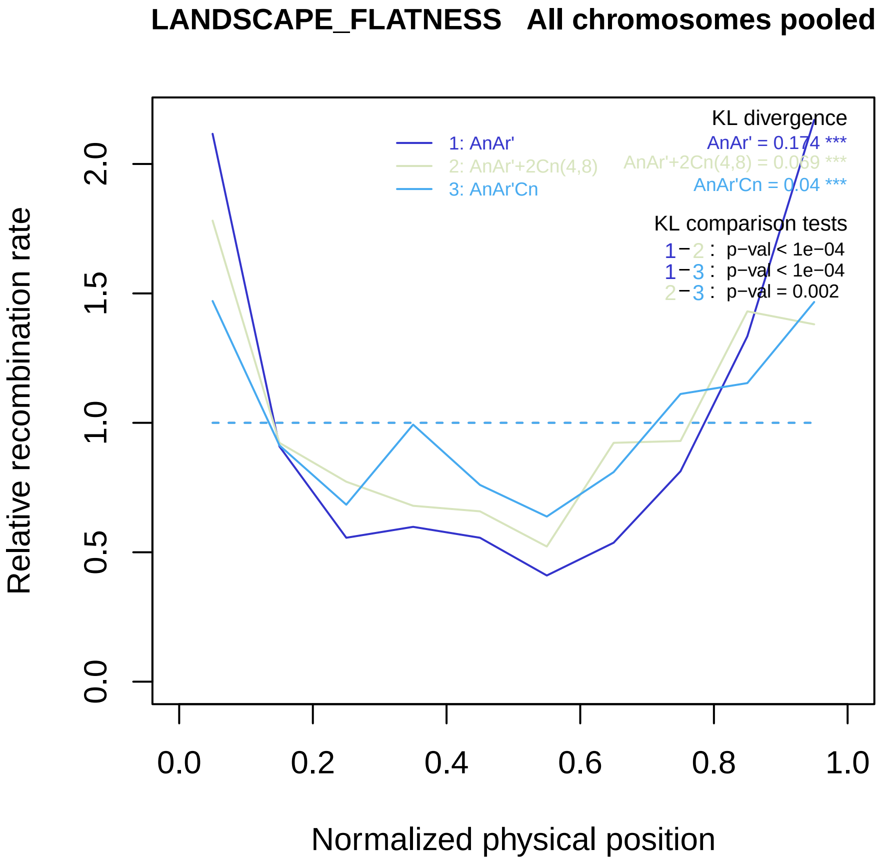
LANDSCAPE_FLATNESS All chromosomes pooled
KL divergence
1: AnAr'
2: AnAr'+2Cn(4,8)
3: AnAr'Cn
AnAr' = 0.174 ***
AnAr'+2Cn(4,8) = 0.069 ***
AnAr'Cn = 0.04 ***
KL comparison tests
1−2 : p−val < 1e−04
1−3 : p−val < 1e−04
2−3 : p−val = 0.002
Relative recombination rate
0.0
0.5
1.0
1.5
2.0
0.0
0.2
0.4
0.6
0.8
1.0
Normalized physical position

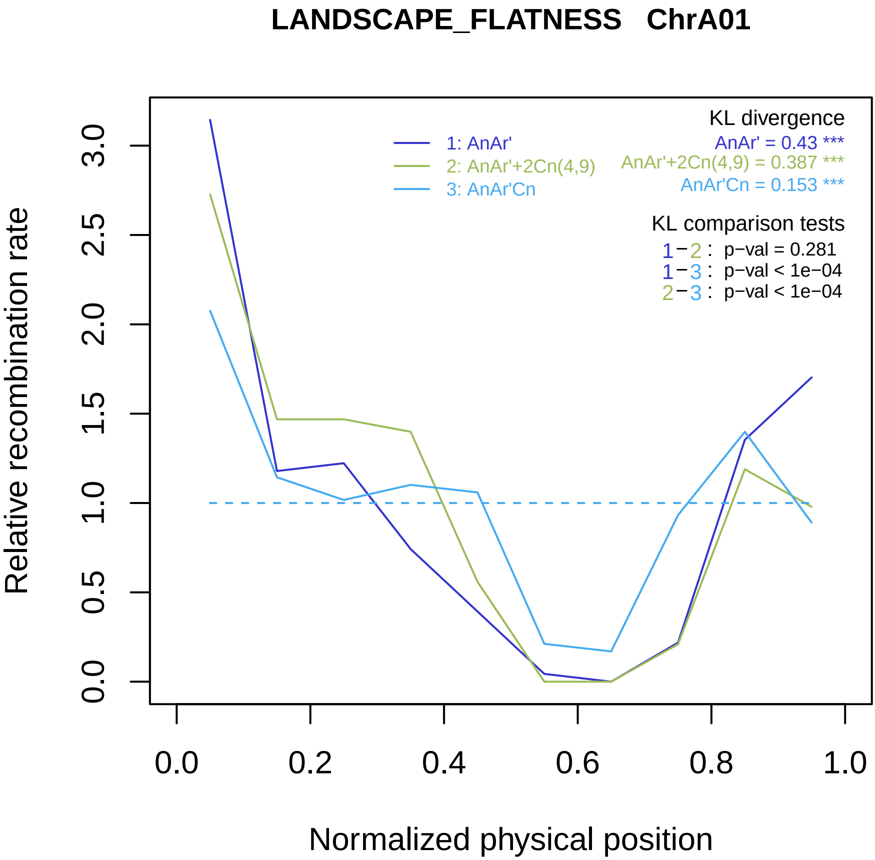

LANDSCAPE_FLATNESS ChrA01
KL divergence
1: AnAr'
2: AnAr'+2Cn(4,9)
3: AnAr'Cn
AnAr' = 0.43 ***
AnAr'+2Cn(4,9) = 0.387 ***
AnAr'Cn = 0.153 ***
KL comparison tests
1−2 : p−val = 0.281
1−3 : p−val < 1e−04
2−3 : p−val < 1e−04
Relative recombination rate
0.0
0.5
1.0
1.5
2.0
2.5
3.0
0.0
0.2
0.4
0.6
0.8
1.0
Normalized physical position

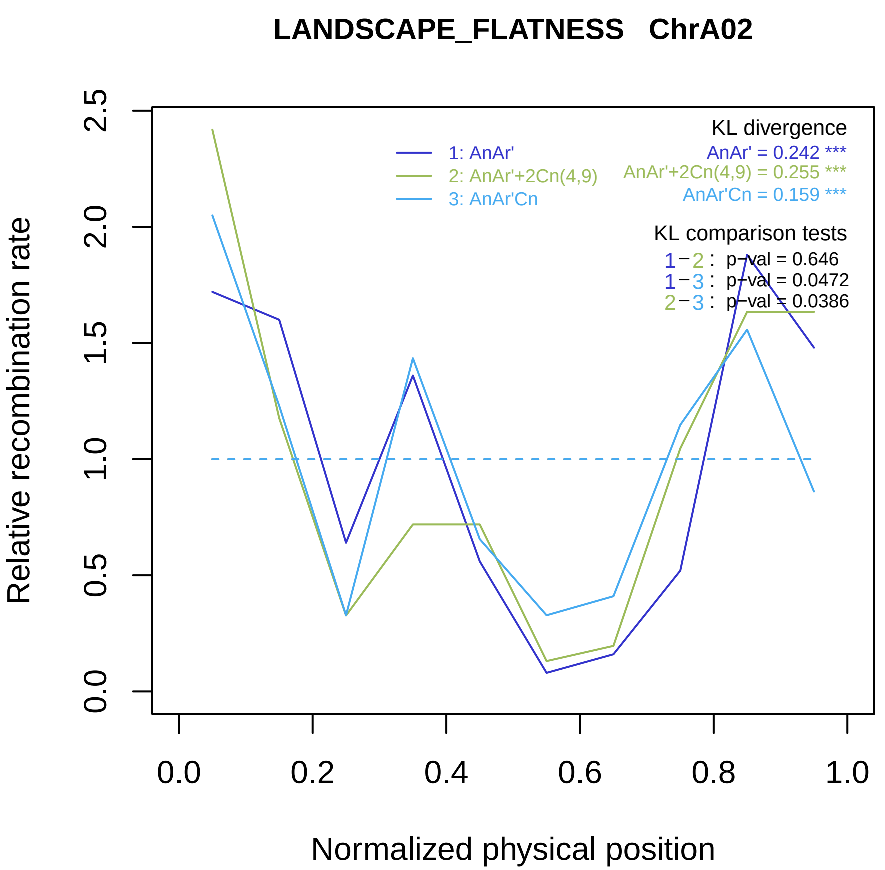

LANDSCAPE_FLATNESS ChrA02
Relative recombination rate
Normalized physical position
0.0
0.5
1.0
1.5
2.0
2.5
0.0
0.2
0.4
0.6
0.8
1.0
1: AnAr'
2: AnAr'+2Cn(4,9)
3: AnAr'Cn
KL divergence
AnAr' = 0.242 ***
AnAr'+2Cn(4,9) = 0.255 ***
AnAr'Cn = 0.159 ***
KL comparison tests
1−2 : p−val = 0.646
1−3 : p−val = 0.0472
2−3 : p−val = 0.0386

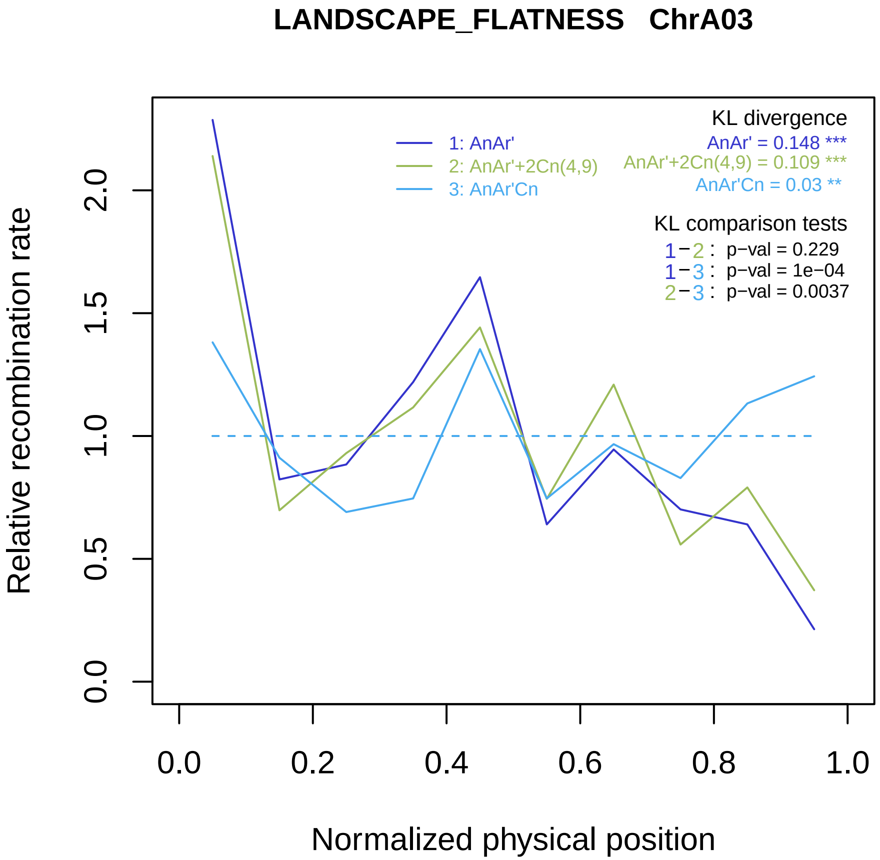

LANDSCAPE_FLATNESS ChrA03
KL divergence
1: AnAr'
2: AnAr'+2Cn(4,9)
3: AnAr'Cn
AnAr' = 0.148 ***
AnAr'+2Cn(4,9) = 0.109 ***
AnAr'Cn = 0.03 **
KL comparison tests
1−2 : p−val = 0.229
1−3 : p−val = 1e−04
2−3 : p−val = 0.0037
Relative recombination rate
0.0
0.5
1.0
1.5
2.0
0.0
0.2
0.4
0.6
0.8
1.0
Normalized physical position

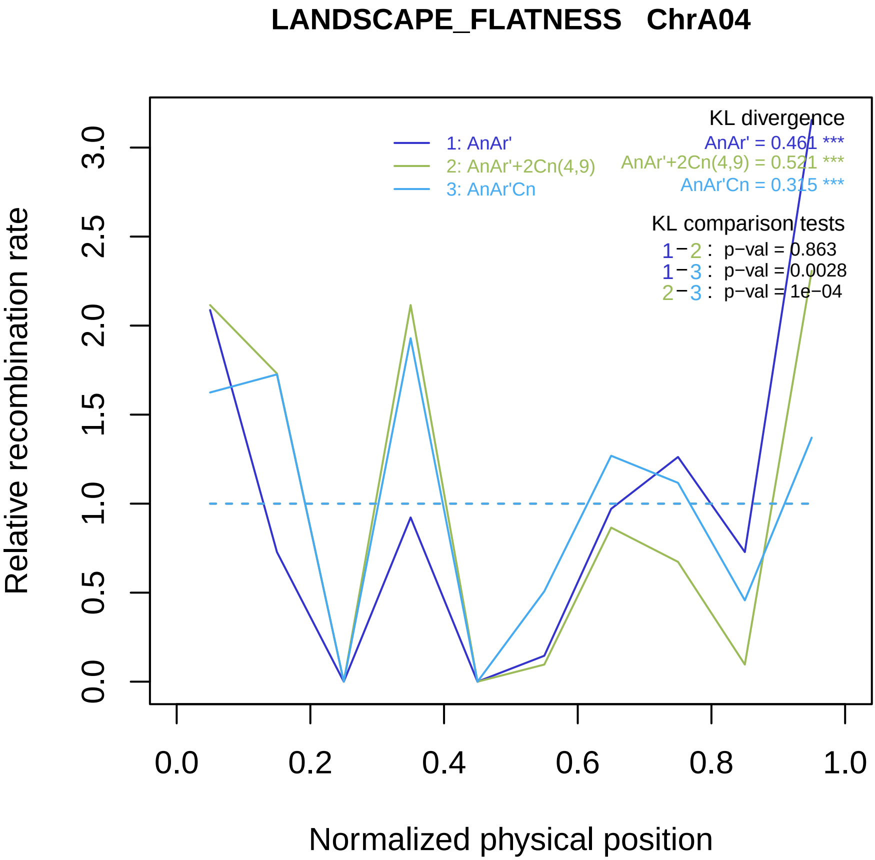
LANDSCAPE_FLATNESS ChrA04
KL divergence
AnAr' = 0.461 ***
AnAr'+2Cn(4,9) = 0.521 ***
AnAr'Cn = 0.315 ***
1: AnAr'
2: AnAr'+2Cn(4,9)
3: AnAr'Cn
KL comparison tests
1−2 : p−val = 0.863
1−3 : p−val = 0.0028
2−3 : p−val = 1e−04
Relative recombination rate
0.0
0.5
1.0
1.5
2.0
2.5
3.0
0.0
0.2
0.4
0.6
0.8
1.0
Normalized physical position

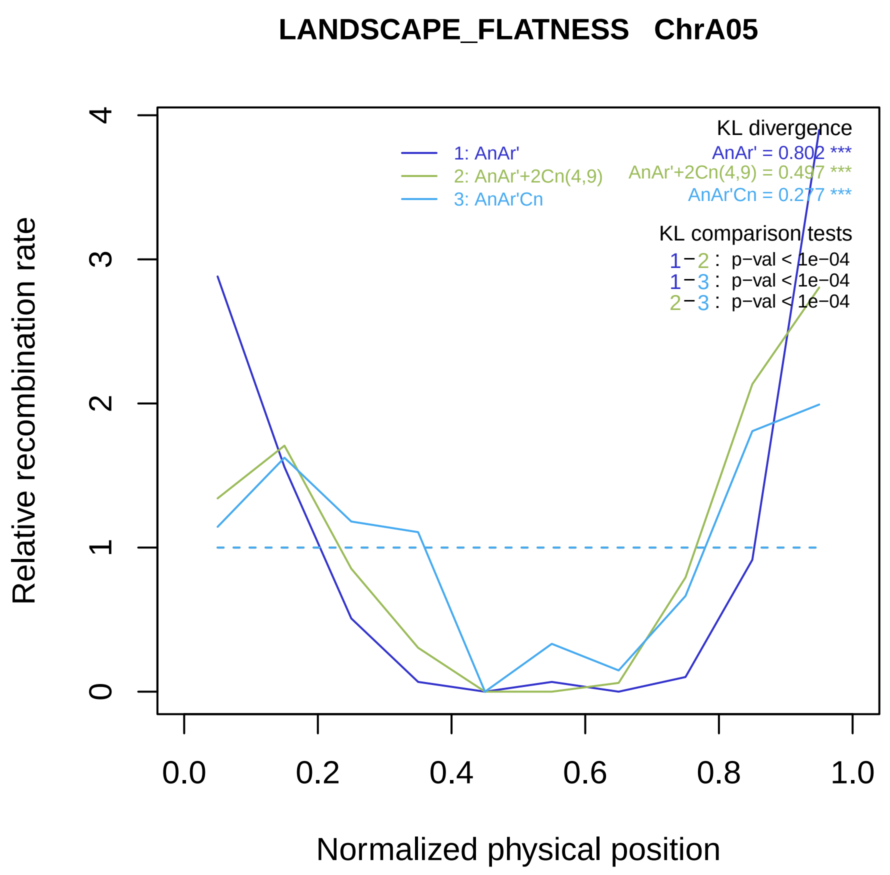
LANDSCAPE_FLATNESS ChrA05
Relative recombination rate
Normalized physical position
0
1
2
3
4
0.0
0.2
0.4
0.6
0.8
1.0
1: AnAr'
2: AnAr'+2Cn(4,9)
3: AnAr'Cn
KL divergence
AnAr' = 0.802 ***
AnAr'+2Cn(4,9) = 0.497 ***
AnAr'Cn = 0.277 ***
KL comparison tests
1−2 : p−val < 1e−04
1−3 : p−val < 1e−04
2−3 : p−val < 1e−04

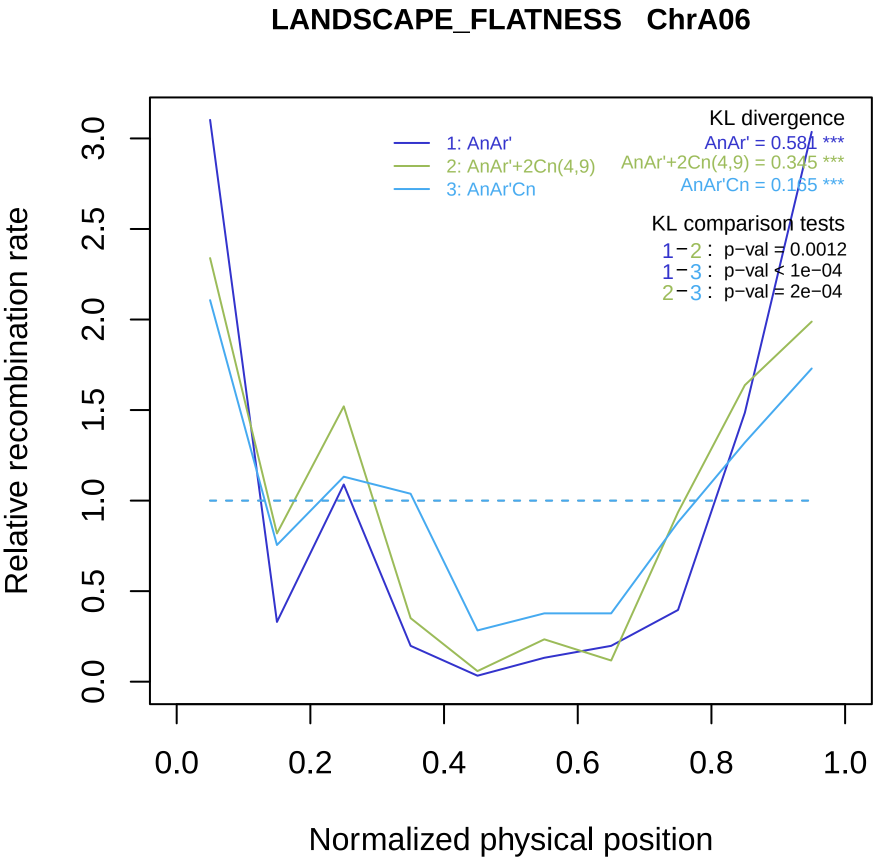
LANDSCAPE_FLATNESS ChrA06
KL divergence
AnAr' = 0.581 ***
AnAr'+2Cn(4,9) = 0.345 ***
AnAr'Cn = 0.165 ***
1: AnAr'
2: AnAr'+2Cn(4,9)
3: AnAr'Cn
KL comparison tests
1−2 : p−val = 0.0012
1−3 : p−val < 1e−04
2−3 : p−val = 2e−04
Relative recombination rate
0.0
0.5
1.0
1.5
2.0
2.5
3.0
0.0
0.2
0.4
0.6
0.8
1.0
Normalized physical position

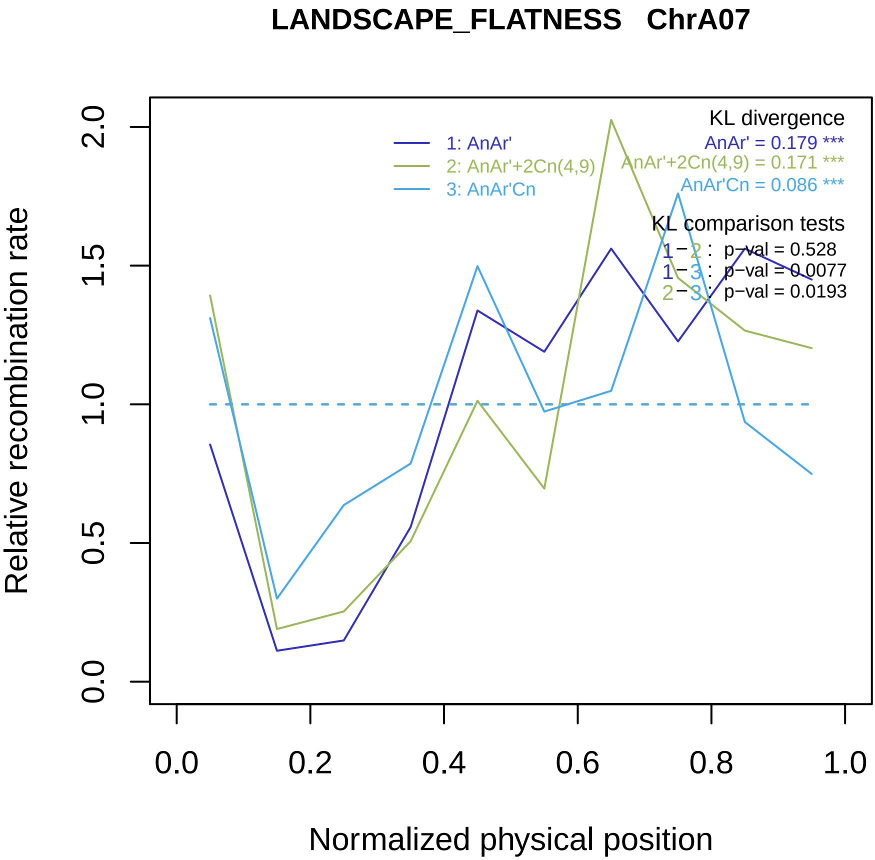

LANDSCAPE_FLATNESS ChrA07
KL divergence
AnAr' = 0.179 ***
AnAr'+2Cn(4,9) = 0.171 ***
AnAr'Cn = 0.086 ***
1: AnAr'
2: AnAr'+2Cn(4,9)
3: AnAr'Cn
KL comparison tests
1 − 2 : p−val = 0.528
1 − 3 : p−val = 0.0077
2 − 3 : p−val = 0.0193
Relative recombination rate
0.0
0.5
1.0
1.5
2.0
0.0
0.2
0.4
0.6
0.8
1.0
Normalized physical position

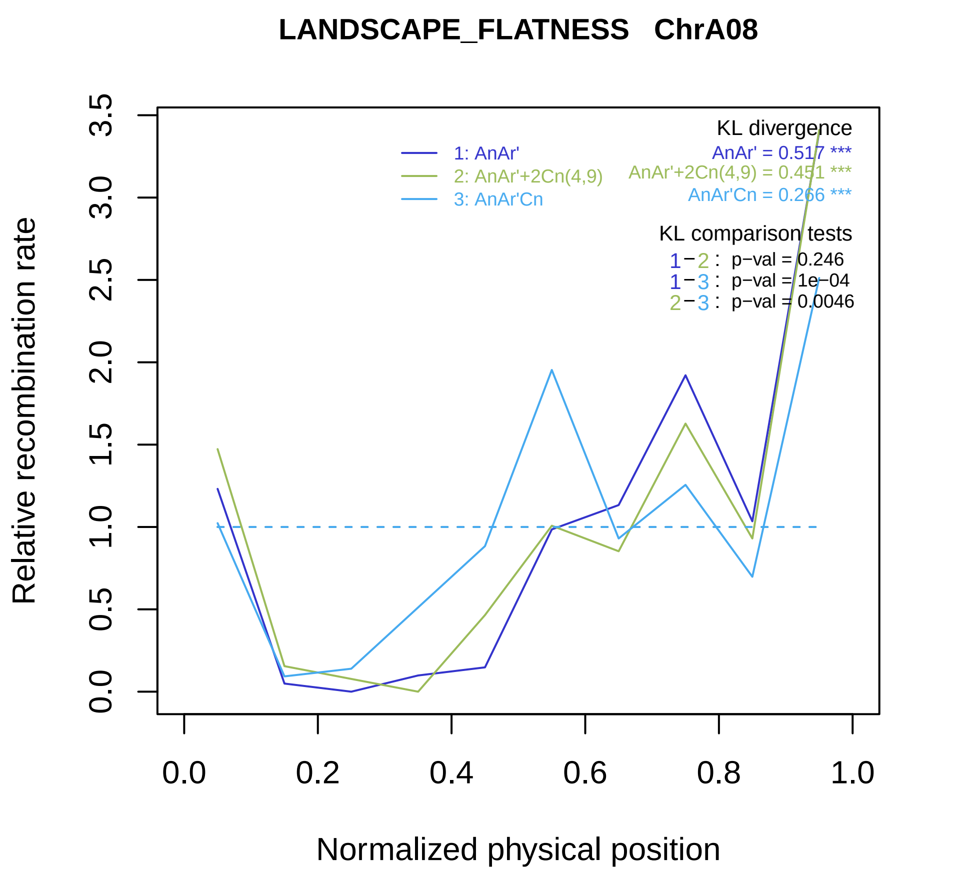
LANDSCAPE_FLATNESS ChrA08
Relative recombination rate
0.0
0.5
1.0
1.5
2.0
2.5
3.0
3.5
0.0
0.2
0.4
0.6
0.8
1.0
Normalized physical position
1: AnAr'
2: AnAr'+2Cn(4,9)
3: AnAr'Cn
KL divergence
AnAr' = 0.517 ***
AnAr'+2Cn(4,9) = 0.451 ***
AnAr'Cn = 0.266 ***
KL comparison tests
1−2 : p−val = 0.246
1−3 : p−val = 1e−04
2−3 : p−val = 0.0046

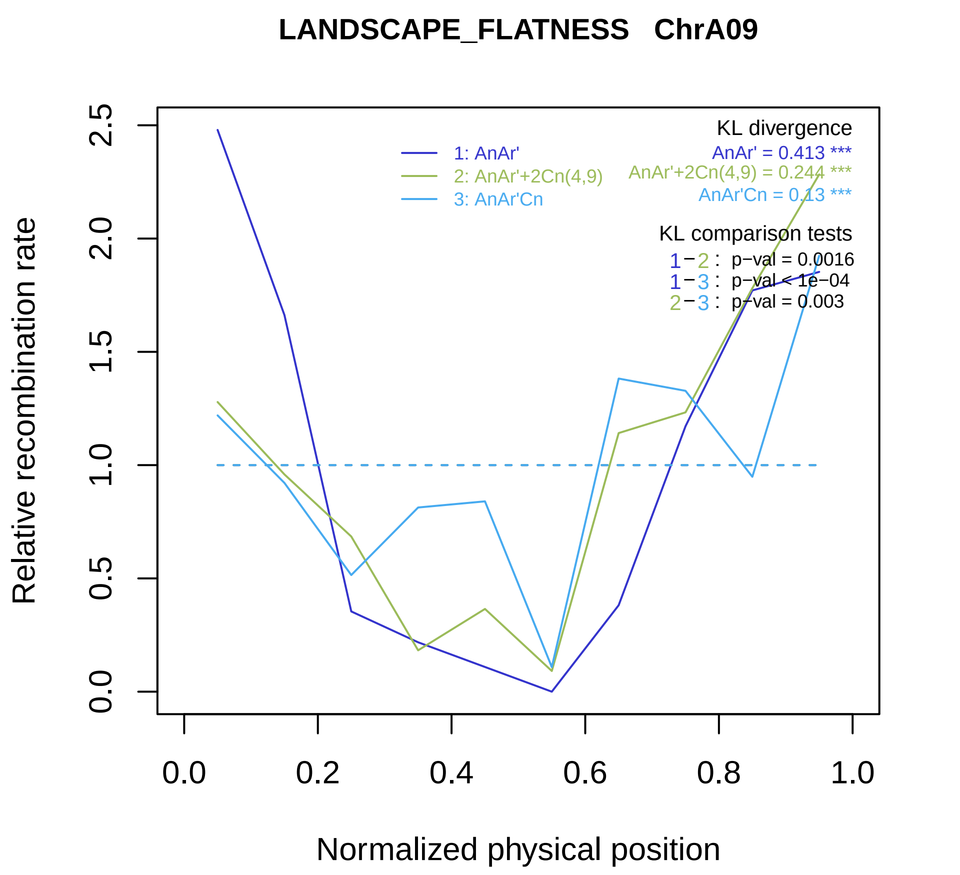

LANDSCAPE_FLATNESS ChrA09
Relative recombination rate
Normalized physical position
0.0
0.5
1.0
1.5
2.0
2.5
0.0
0.2
0.4
0.6
0.8
1.0
1: AnAr'
2: AnAr'+2Cn(4,9)
3: AnAr'Cn
KL divergence
AnAr' = 0.413 ***
AnAr'+2Cn(4,9) = 0.244 ***
AnAr'Cn = 0.13 ***
KL comparison tests
1−2 : p−val = 0.0016
1−3 : p−val < 1e−04
2−3 : p−val = 0.003

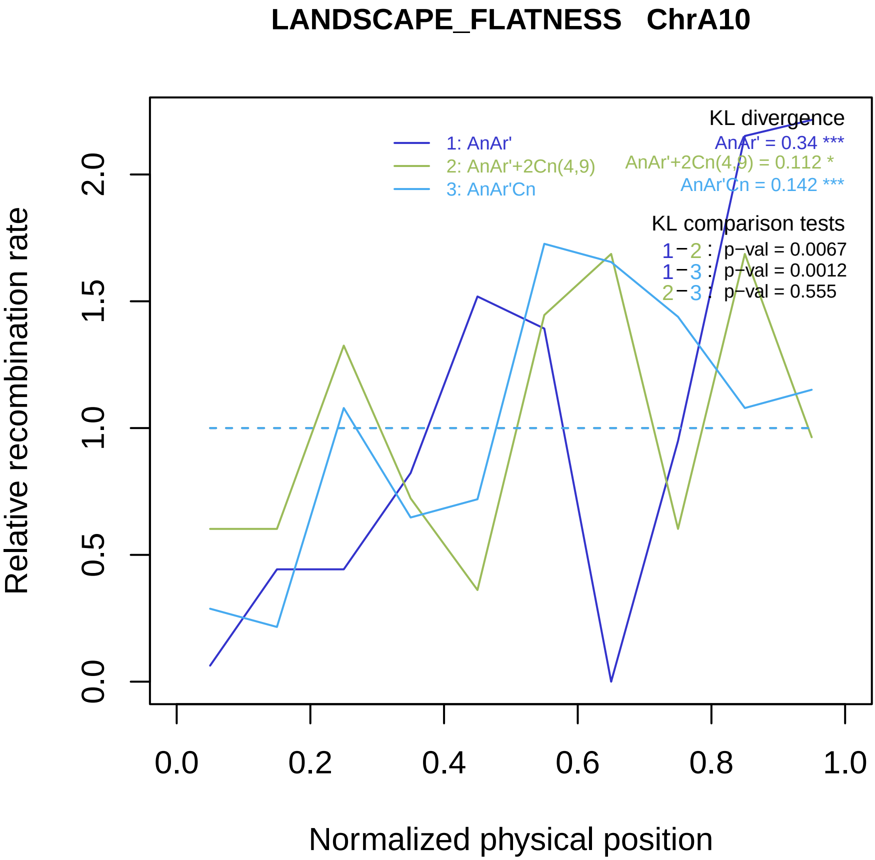

LANDSCAPE_FLATNESS ChrA10
Relative recombination rate
0.0
0.5
1.0
1.5
2.0
Normalized physical position
0.0
0.2
0.4
0.6
0.8
1.0
1: AnAr'
2: AnAr'+2Cn(4,9)
3: AnAr'Cn
KL divergence
AnAr' = 0.34 ***
AnAr'+2Cn(4,9) = 0.112 *
AnAr'Cn = 0.142 ***
KL comparison tests
1−2 : p−val = 0.0067
1−3 : p−val = 0.0012
2−3 : p−val = 0.555

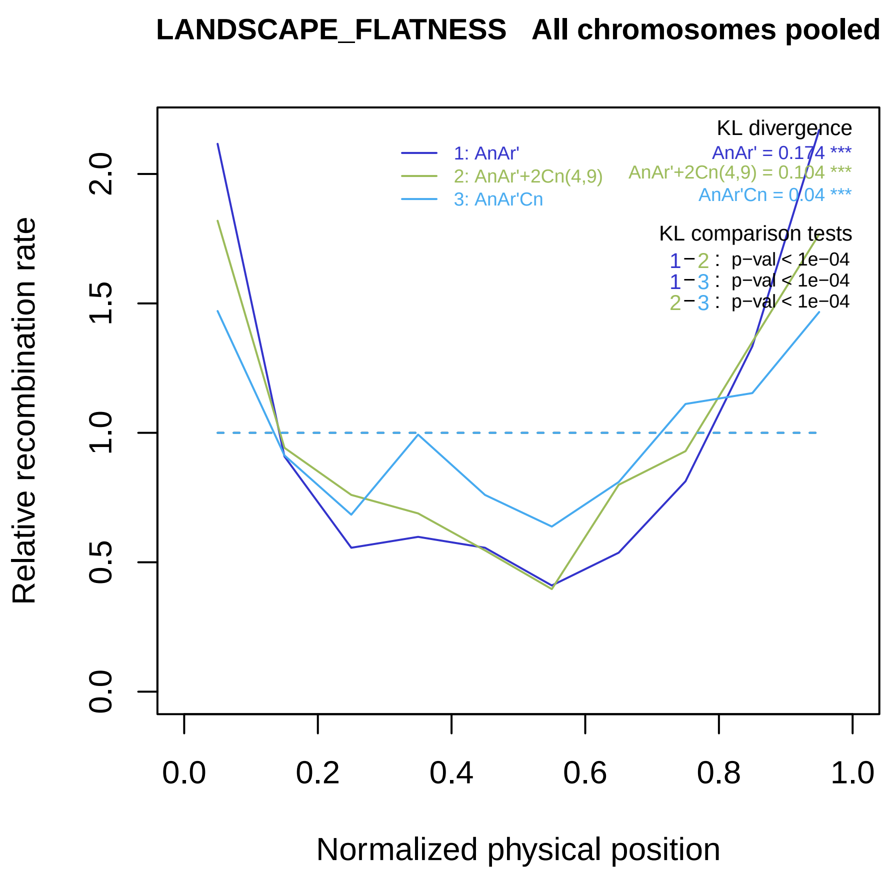

LANDSCAPE_FLATNESS All chromosomes pooled
KL divergence
AnAr' = 0.174 ***
AnAr'+2Cn(4,9) = 0.104 ***
AnAr'Cn = 0.04 ***
1: AnAr'
2: AnAr'+2Cn(4,9)
3: AnAr'Cn
KL comparison tests
1−2 : p−val < 1e−04
1−3 : p−val < 1e−04
2−3 : p−val < 1e−04
Relative recombination rate
Normalized physical position
0.0
0.5
1.0
1.5
2.0
0.2
0.4
0.6
0.8
1.0

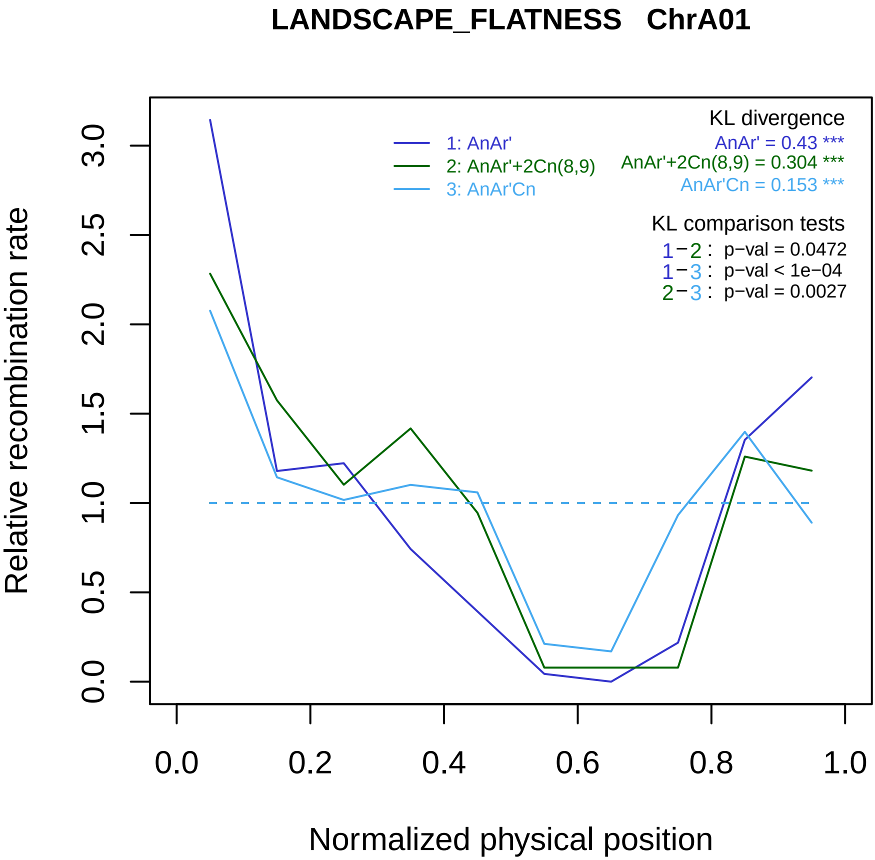

LANDSCAPE_FLATNESS ChrA01
Relative recombination rate
0.0
0.5
1.0
1.5
2.0
2.5
3.0
0.0
0.2
0.4
0.6
0.8
1.0
Normalized physical position
1: AnAr'
2: AnAr'+2Cn(8,9)
3: AnAr'Cn
KL divergence
AnAr' = 0.43 ***
AnAr'+2Cn(8,9) = 0.304 ***
AnAr'Cn = 0.153 ***
KL comparison tests
1−2 : p−val = 0.0472
1−3 : p−val < 1e−04
2−3 : p−val = 0.0027

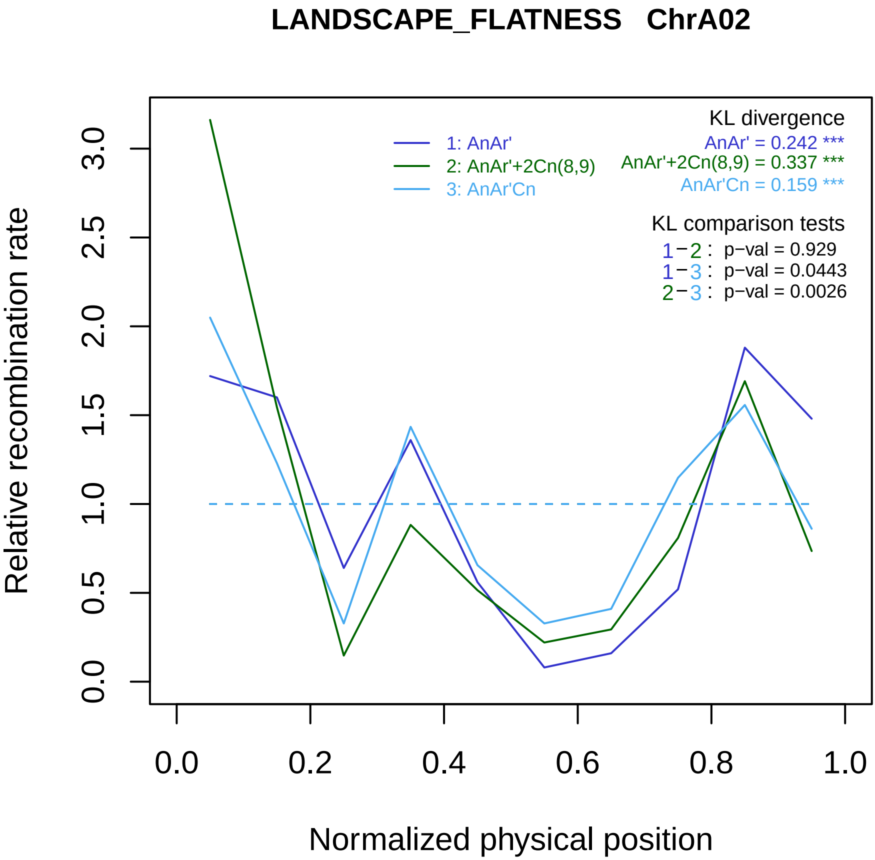
LANDSCAPE_FLATNESS ChrA02
KL divergence
AnAr' = 0.242 ***
AnAr'+2Cn(8,9) = 0.337 ***
AnAr'Cn = 0.159 ***
1: AnAr'
2: AnAr'+2Cn(8,9)
3: AnAr'Cn
KL comparison tests
1−2 : p−val = 0.929
1−3 : p−val = 0.0443
2−3 : p−val = 0.0026
Relative recombination rate
0.0
0.5
1.0
1.5
2.0
2.5
3.0
0.0
0.2
0.4
0.6
0.8
1.0
Normalized physical position

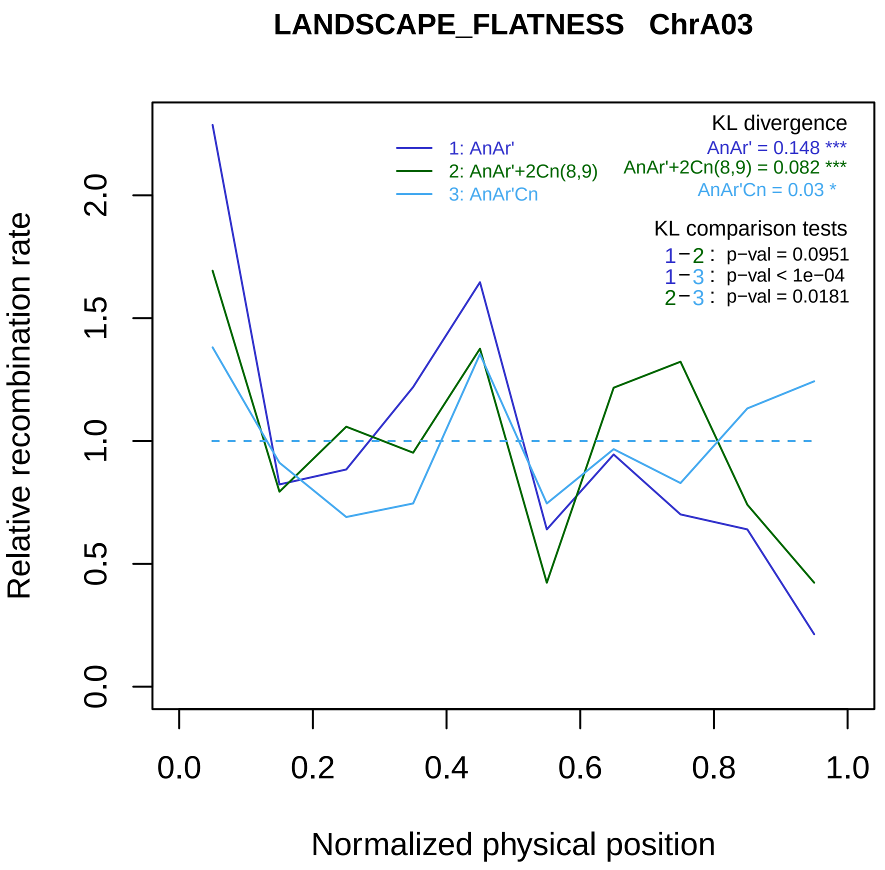

LANDSCAPE_FLATNESS ChrA03
KL divergence
1: AnAr'
2: AnAr'+2Cn(8,9)
3: AnAr'Cn
AnAr' = 0.148 ***
AnAr'+2Cn(8,9) = 0.082 ***
AnAr'Cn = 0.03 *
KL comparison tests
1−2 : p−val = 0.0951
1−3 : p−val < 1e−04
2−3 : p−val = 0.0181
Relative recombination rate
0.0
0.5
1.0
1.5
2.0
0.0
0.2
0.4
0.6
0.8
1.0
Normalized physical position

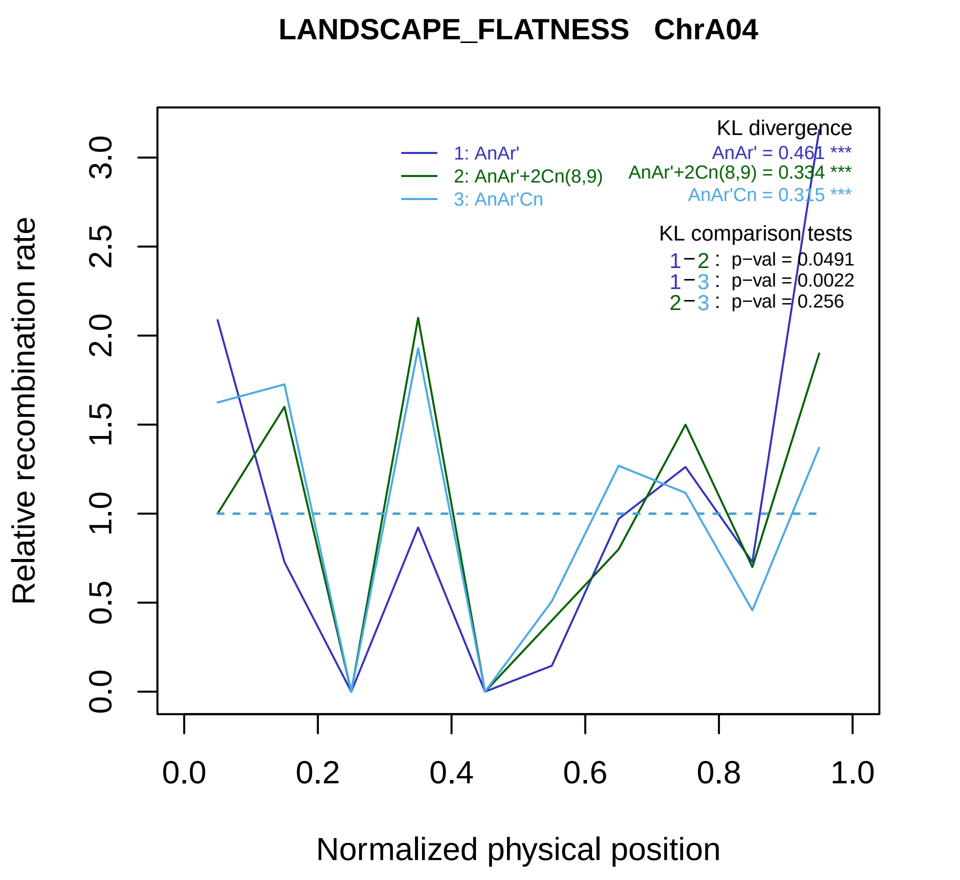

LANDSCAPE_FLATNESS ChrA04
KL divergence
AnAr' = 0.461 ***
AnAr'+2Cn(8,9) = 0.334 ***
AnAr'Cn = 0.315 ***
1: AnAr'
2: AnAr'+2Cn(8,9)
3: AnAr'Cn
KL comparison tests
1−2 : p−val = 0.0491
1−3 : p−val = 0.0022
2−3 : p−val = 0.256
Relative recombination rate
0.0
0.5
1.0
1.5
2.0
2.5
3.0
0.0
0.2
0.4
0.6
0.8
1.0
Normalized physical position

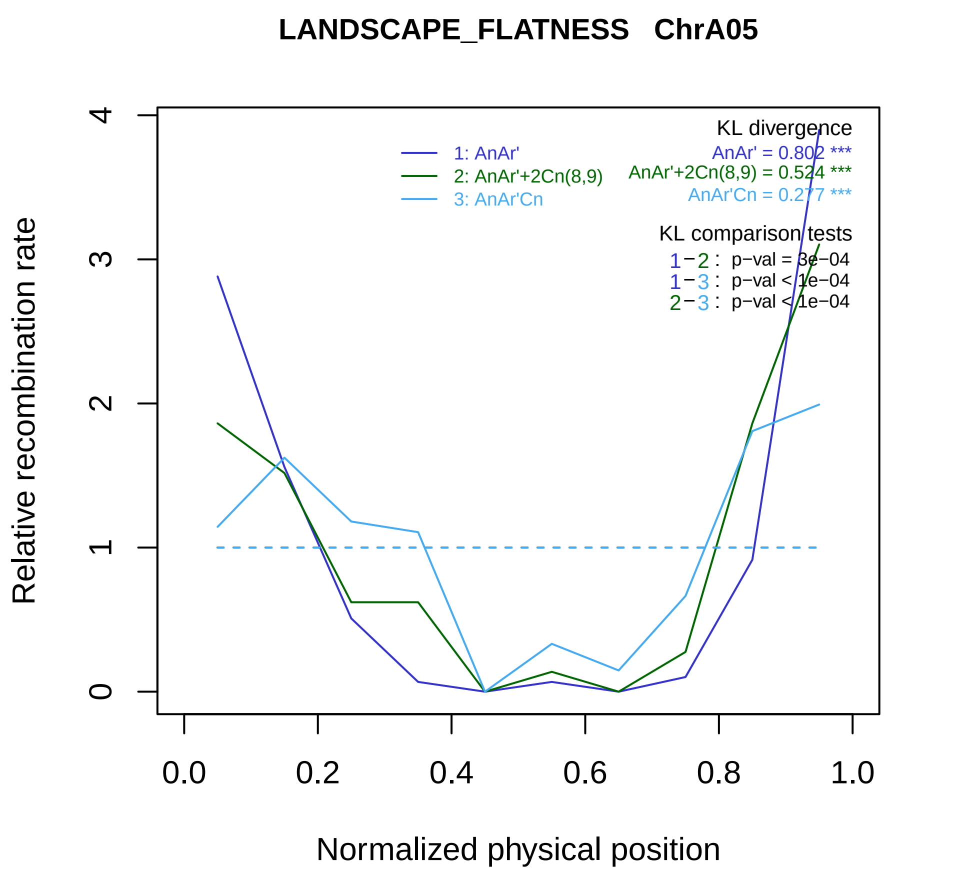
LANDSCAPE_FLATNESS ChrA05
Relative recombination rate
Normalized physical position
0
1
2
3
4
0.0
0.2
0.4
0.6
0.8
1.0
1: AnAr'
2: AnAr'+2Cn(8,9)
3: AnAr'Cn
KL divergence
AnAr' = 0.802 ***
AnAr'+2Cn(8,9) = 0.524 ***
AnAr'Cn = 0.277 ***
KL comparison tests
1−2 : p−val = 3e−04
1−3 : p−val < 1e−04
2−3 : p−val < 1e−04

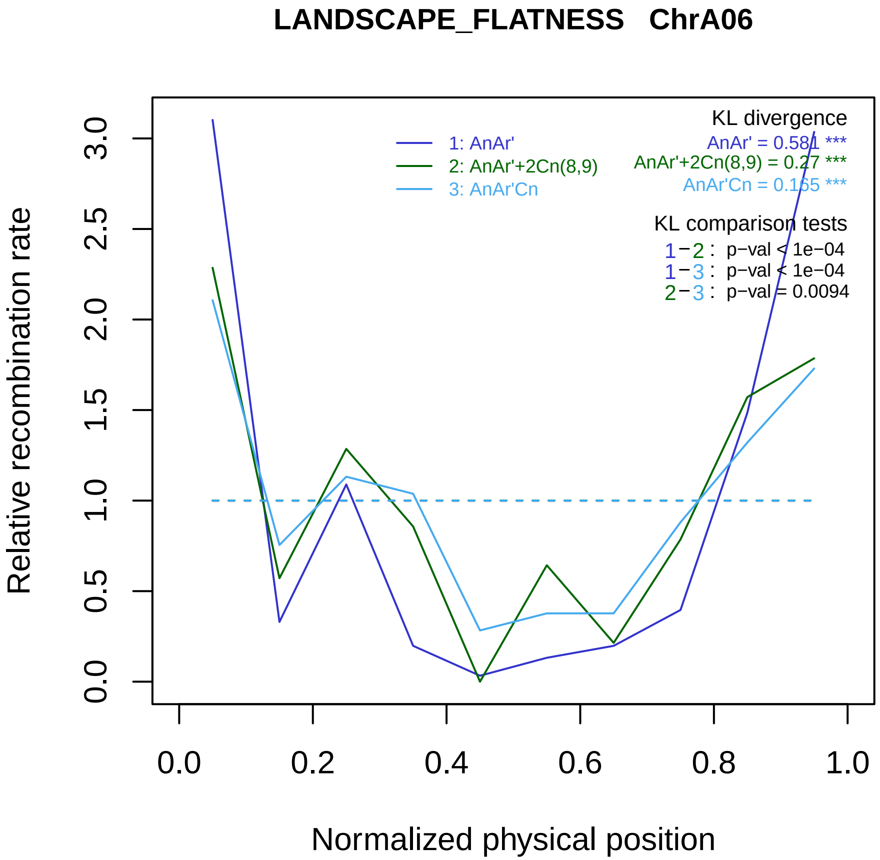
LANDSCAPE_FLATNESS ChrA06
1: AnAr'
2: AnAr'+2Cn(8,9)
3: AnAr'Cn
KL divergence
AnAr' = 0.581 ***
AnAr'+2Cn(8,9) = 0.27 ***
AnAr'Cn = 0.165 ***
KL comparison tests
1−2 : p−val < 1e−04
1−3 : p−val < 1e−04
2−3 : p−val = 0.0094
Relative recombination rate
0.0
0.5
1.0
1.5
2.0
2.5
3.0
Normalized physical position
0.0
0.2
0.4
0.6
0.8
1.0

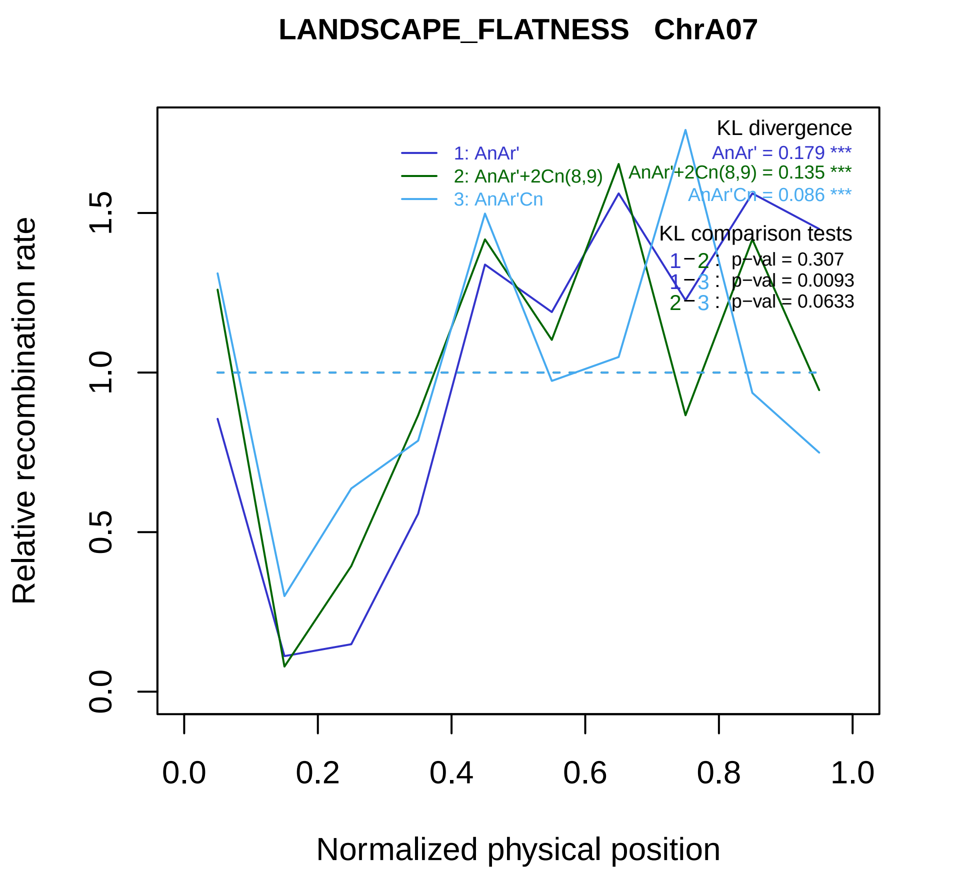

LANDSCAPE_FLATNESS ChrA07
KL divergence
AnAr' = 0.179 ***
AnAr'+2Cn(8,9) = 0.135 ***
AnAr'Cn = 0.086 ***
1: AnAr'
2: AnAr'+2Cn(8,9)
3: AnAr'Cn
KL comparison tests
1−2 : p−val = 0.307
1−3 : p−val = 0.0093
2−3 : p−val = 0.0633
Relative recombination rate
0.0
0.5
1.0
1.5
0.0
0.2
0.4
0.6
0.8
1.0
Normalized physical position

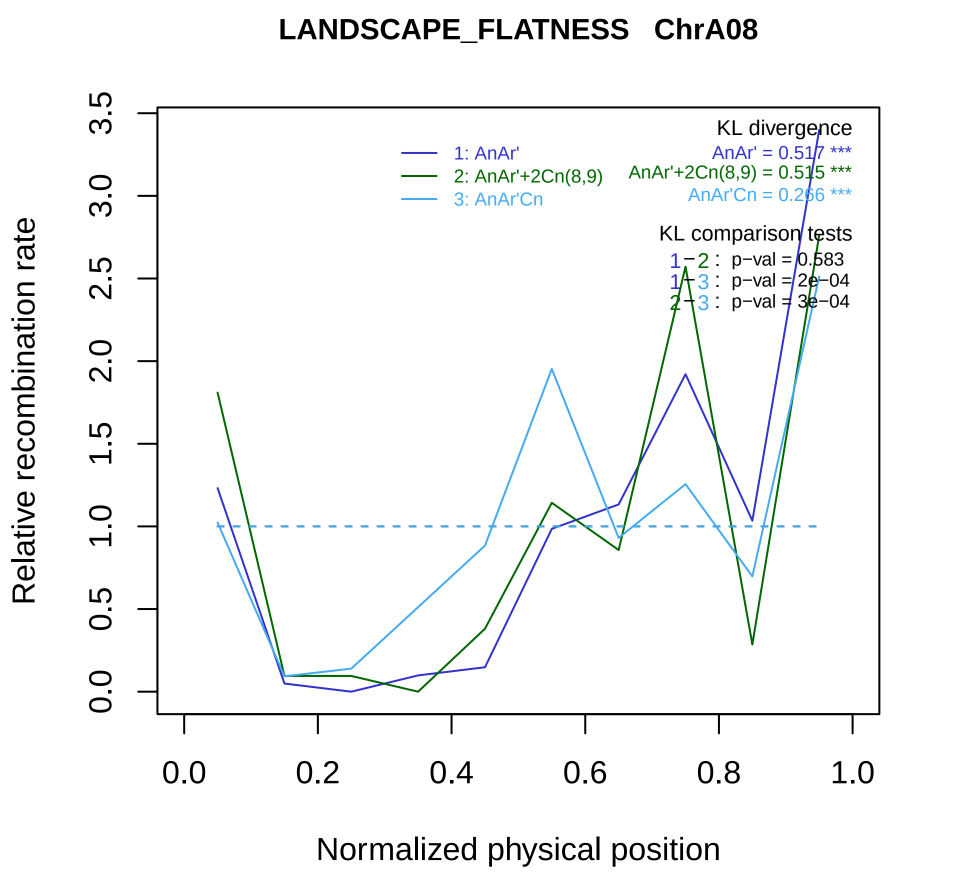
LANDSCAPE_FLATNESS ChrA08
Relative recombination rate
0.0
0.5
1.0
1.5
2.0
2.5
3.0
3.5
Normalized physical position
0.0
0.2
0.4
0.6
0.8
1.0
1: AnAr'
2: AnAr'+2Cn(8,9)
3: AnAr'Cn
KL divergence
AnAr' = 0.517 ***
AnAr'+2Cn(8,9) = 0.515 ***
AnAr'Cn = 0.266 ***
KL comparison tests
1−2 : p−val = 0.583
1−3 : p−val = 2e−04
2−3 : p−val = 3e−04

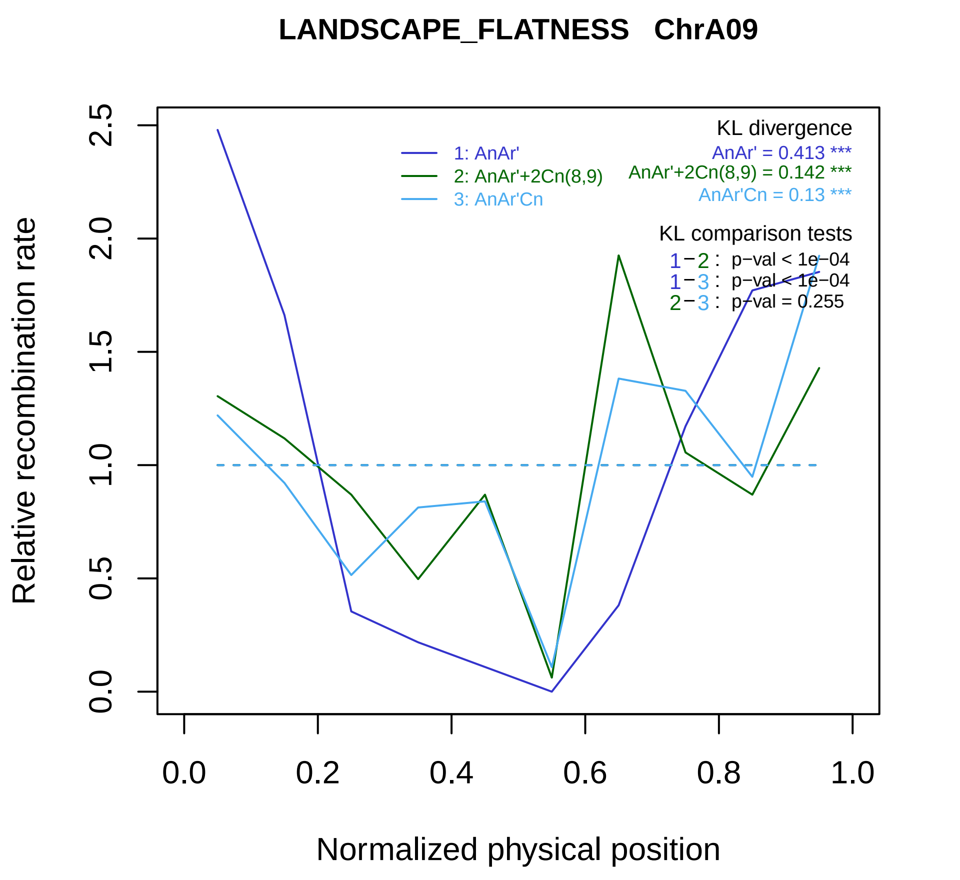
LANDSCAPE_FLATNESS ChrA09
KL divergence
AnAr' = 0.413 ***
AnAr'+2Cn(8,9) = 0.142 ***
AnAr'Cn = 0.13 ***
1: AnAr'
2: AnAr'+2Cn(8,9)
3: AnAr'Cn
KL comparison tests
1−2 : p−val < 1e−04
1−3 : p−val < 1e−04
2−3 : p−val = 0.255
Relative recombination rate
0.0
0.5
1.0
1.5
2.0
2.5
0.0
0.2
0.4
0.6
0.8
1.0
Normalized physical position

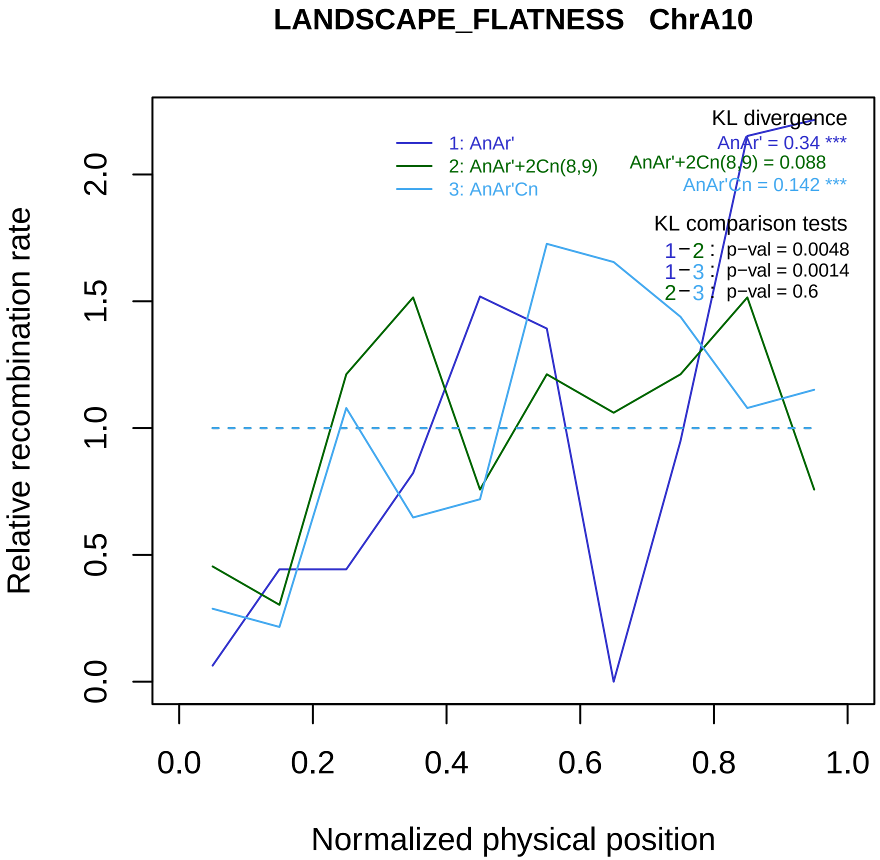

LANDSCAPE_FLATNESS   ChrA10
Relative recombination rate
Normalized physical position
0.0
0.5
1.0
1.5
2.0
0.0
0.2
0.4
0.6
0.8
1.0
1: AnAr'
2: AnAr'+2Cn(8,9)
3: AnAr'Cn
KL divergence
AnAr' = 0.34 ***
AnAr'+2Cn(8,9) = 0.088
AnAr'Cn = 0.142 ***
KL comparison tests
1−2 : p−val = 0.0048
1−3 : p−val = 0.0014
2−3 : p−val = 0.6

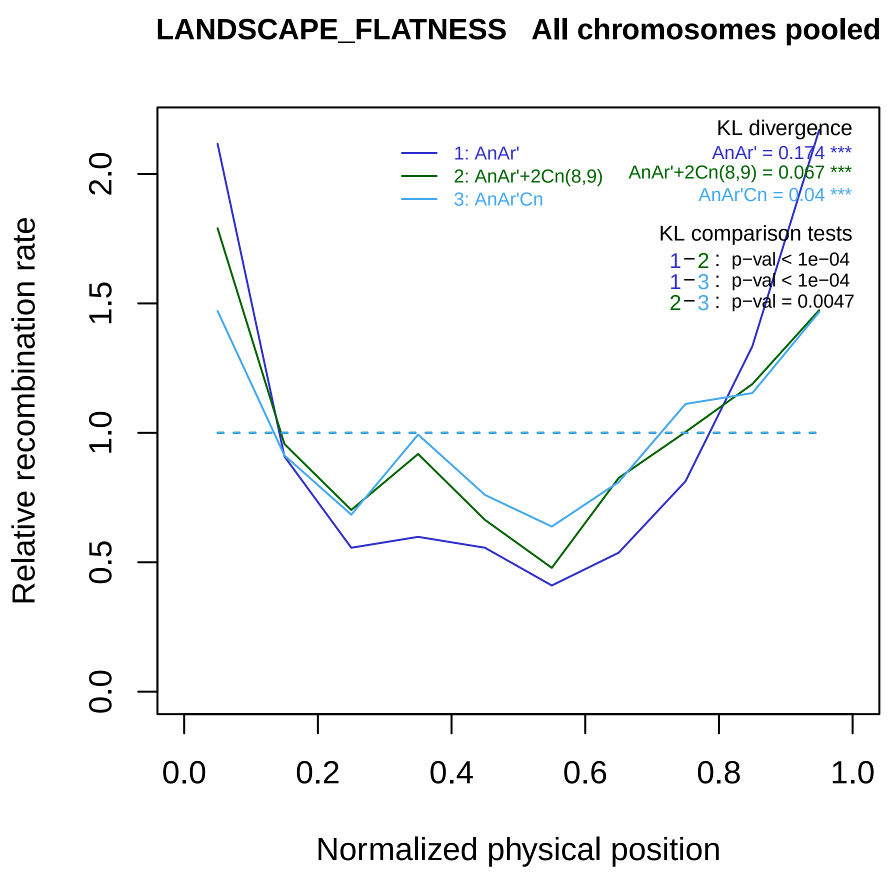
LANDSCAPE_FLATNESS All chromosomes pooled
Relative recombination rate
0.0
0.5
1.0
1.5
2.0
Normalized physical position
0.0
0.2
0.4
0.6
0.8
1.0
1: AnAr'
2: AnAr'+2Cn(8,9)
3: AnAr'Cn
KL divergence
AnAr' = 0.174 ***
AnAr'+2Cn(8,9) = 0.067 ***
AnAr'Cn = 0.04 ***
KL comparison tests
1−2 : p−val < 1e−04
1−3 : p−val < 1e−04
2−3 : p−val = 0.0047

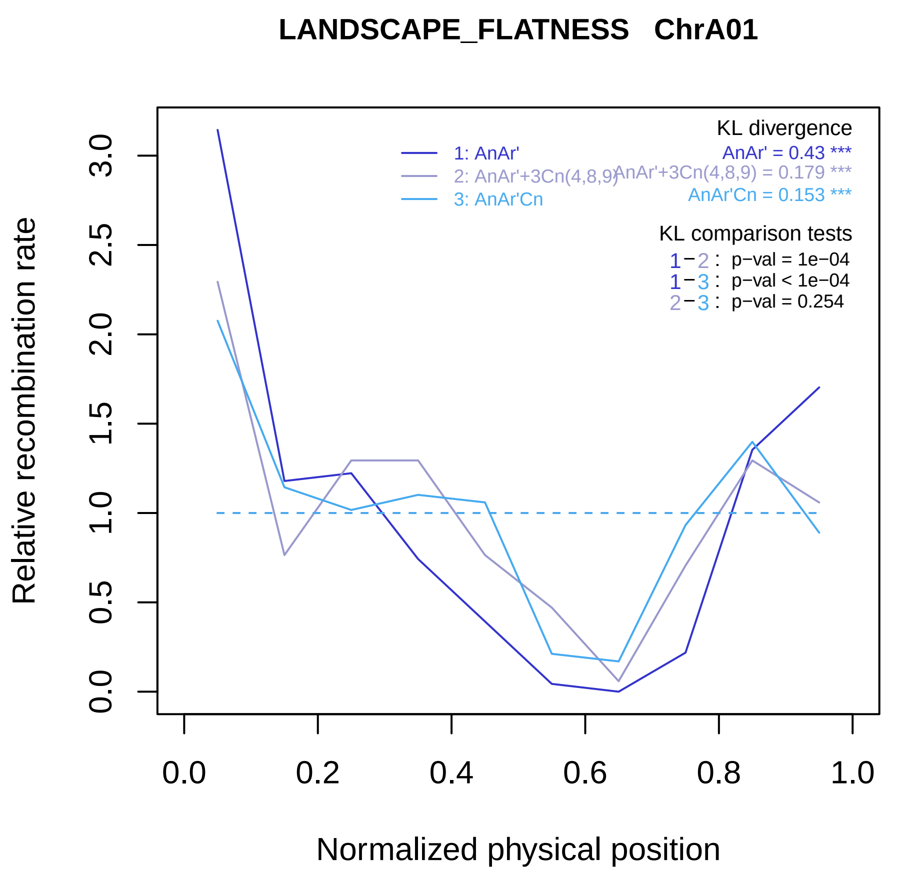

LANDSCAPE_FLATNESS ChrA01
Relative recombination rate
0.0
0.5
1.0
1.5
2.0
2.5
3.0
0.0
0.2
0.4
0.6
0.8
1.0
Normalized physical position
1: AnAr'
2: AnAr'+3Cn(4,8,9)
3: AnAr'Cn
KL divergence
AnAr' = 0.43 ***
AnAr'+3Cn(4,8,9) = 0.179 ***
AnAr'Cn = 0.153 ***
KL comparison tests
1−2 : p−val = 1e−04
1−3 : p−val < 1e−04
2−3 : p−val = 0.254

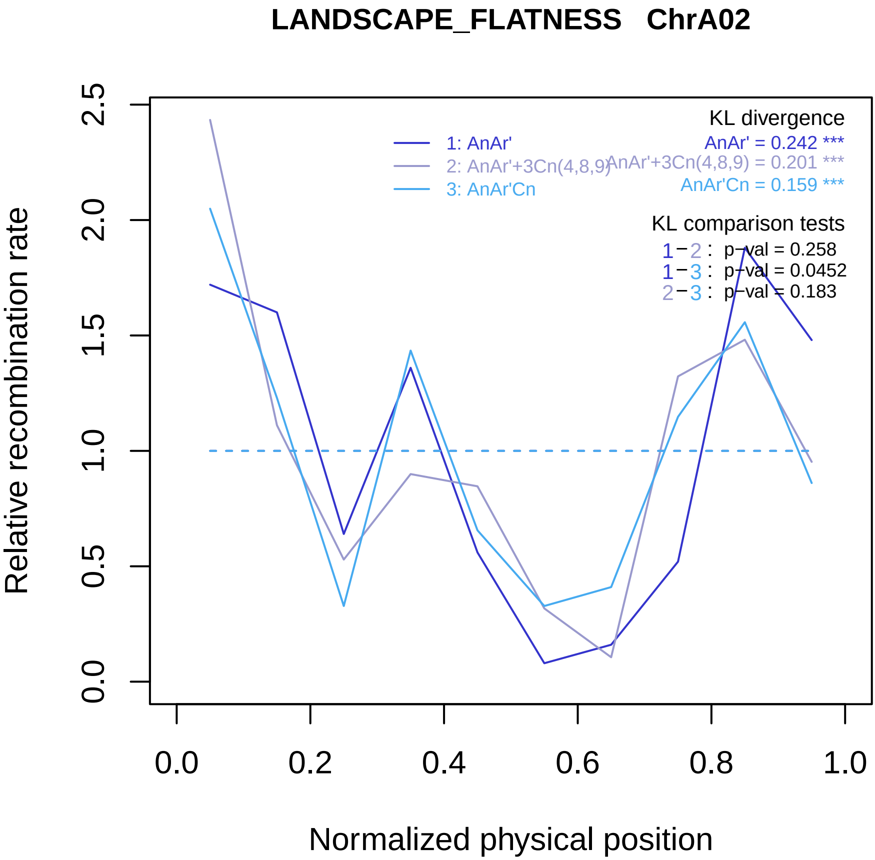
LANDSCAPE_FLATNESS ChrA02
KL divergence
AnAr' = 0.242 ***
AnAr'+3Cn(4,8,9) = 0.201 ***
AnAr'Cn = 0.159 ***
1: AnAr'
2: AnAr'+3Cn(4,8,9)
3: AnAr'Cn
KL comparison tests
1−2 : p−val = 0.258
1−3 : p−val = 0.0452
2−3 : p−val = 0.183
Relative recombination rate
Normalized physical position
0.0
0.5
1.0
1.5
2.0
2.5
0.0
0.2
0.4
0.6
0.8
1.0

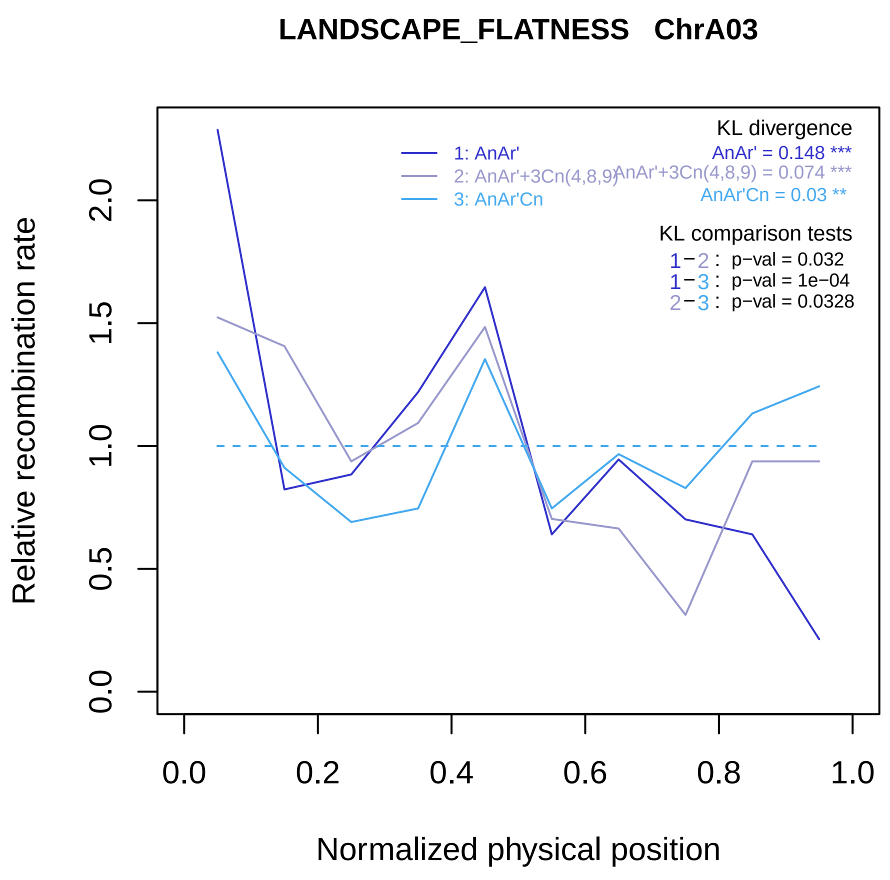

LANDSCAPE_FLATNESS ChrA03
KL divergence
1: AnAr'
2: AnAr'+3Cn(4,8,9)
3: AnAr'Cn
AnAr' = 0.148 ***
AnAr'+3Cn(4,8,9) = 0.074 ***
AnAr'Cn = 0.03 **
KL comparison tests
1−2 : p−val = 0.032
1−3 : p−val = 1e−04
2−3 : p−val = 0.0328
Relative recombination rate
0.0
0.5
1.0
1.5
2.0
0.0
0.2
0.4
0.6
0.8
1.0
Normalized physical position

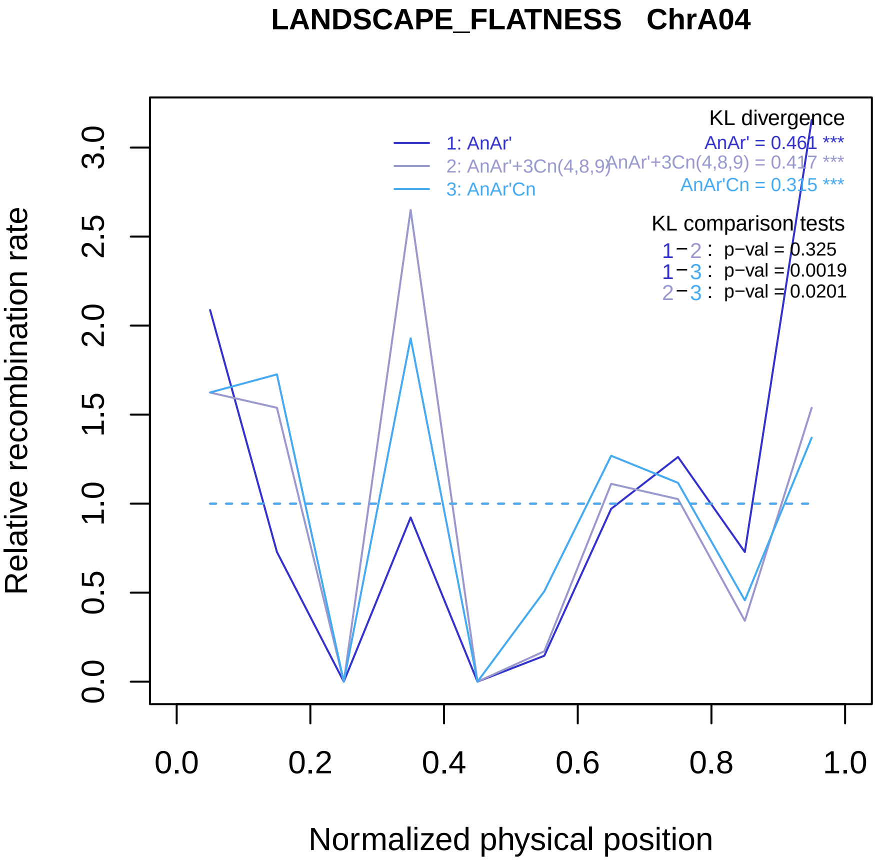

LANDSCAPE_FLATNESS ChrA04
KL divergence
AnAr' = 0.461 ***
AnAr'+3Cn(4,8,9) = 0.417 ***
AnAr'Cn = 0.315 ***
1: AnAr'
2: AnAr'+3Cn(4,8,9)
3: AnAr'Cn
KL comparison tests
1−2 : p−val = 0.325
1−3 : p−val = 0.0019
2−3 : p−val = 0.0201
Relative recombination rate
0.0
0.5
1.0
1.5
2.0
2.5
3.0
0.0
0.2
0.4
0.6
0.8
1.0
Normalized physical position

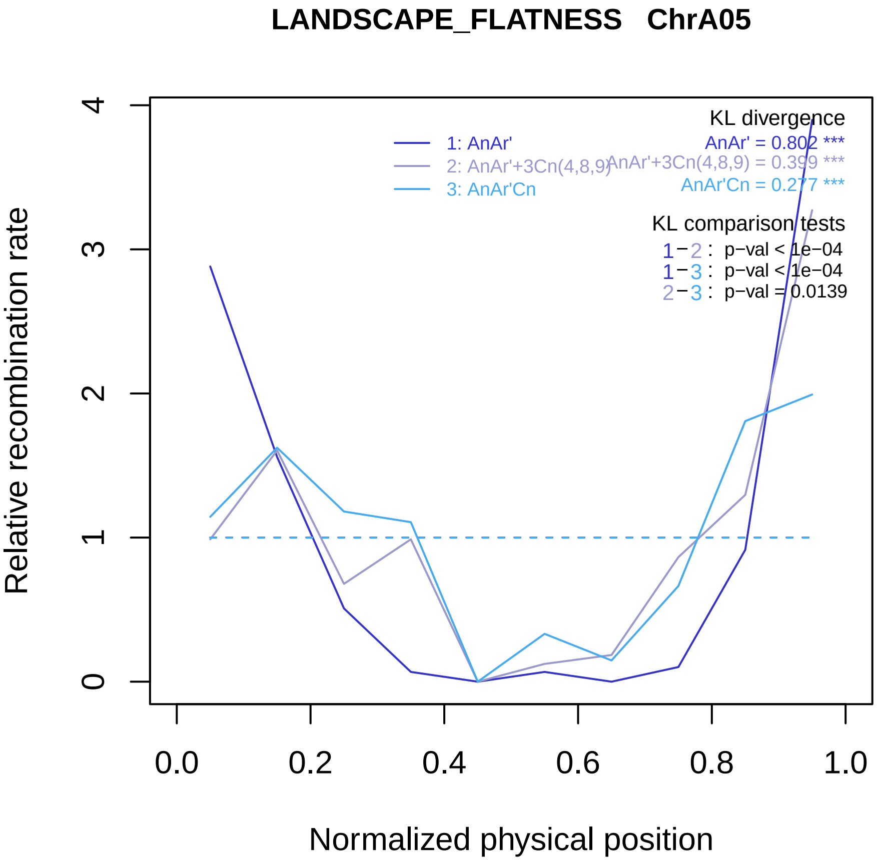

LANDSCAPE_FLATNESS ChrA05
KL divergence
AnAr' = 0.802 ***
AnAr'+3Cn(4,8,9) = 0.399 ***
AnAr'Cn = 0.277 ***
1: AnAr'
2: AnAr'+3Cn(4,8,9)
3: AnAr'Cn
KL comparison tests
1−2 : p−val < 1e−04
1−3 : p−val < 1e−04
2−3 : p−val = 0.0139
Relative recombination rate
0
1
2
3
4
0.0
0.2
0.4
0.6
0.8
1.0
Normalized physical position

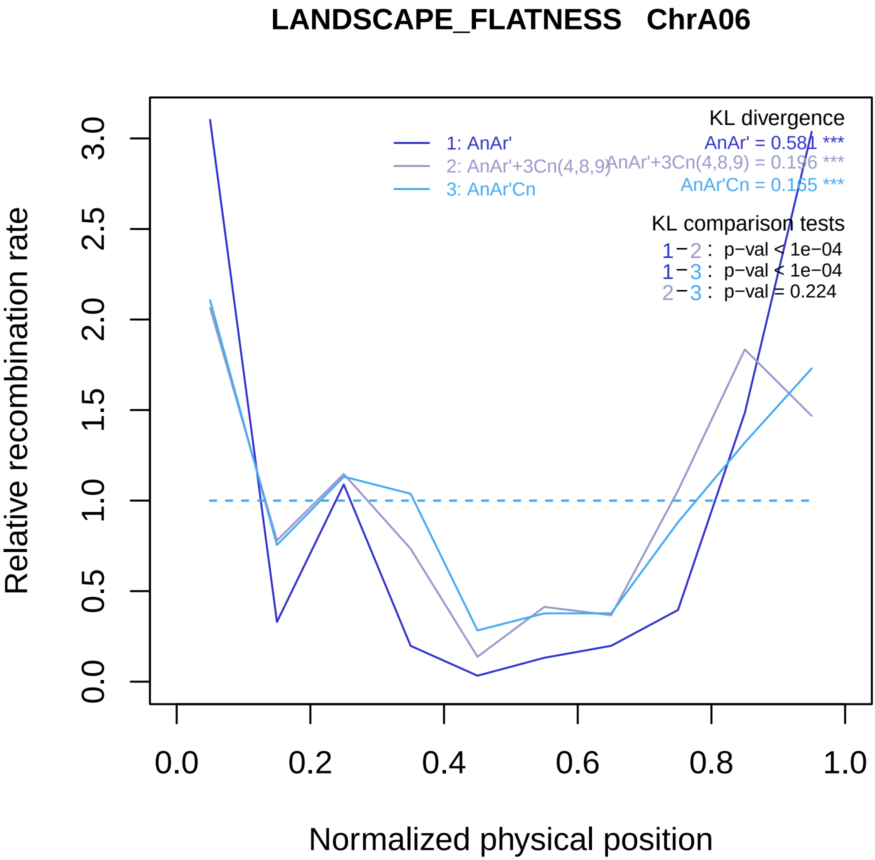
LANDSCAPE_FLATNESS ChrA06
Relative recombination rate
Normalized physical position
0.0
0.5
1.0
1.5
2.0
2.5
3.0
0.0
0.2
0.4
0.6
0.8
1.0
1: AnAr'
2: AnAr'+3Cn(4,8,9)
3: AnAr'Cn
KL divergence
AnAr' = 0.581 ***
AnAr'+3Cn(4,8,9) = 0.196 ***
AnAr'Cn = 0.165 ***
KL comparison tests
1−2 : p−val < 1e−04
1−3 : p−val < 1e−04
2−3 : p−val = 0.224

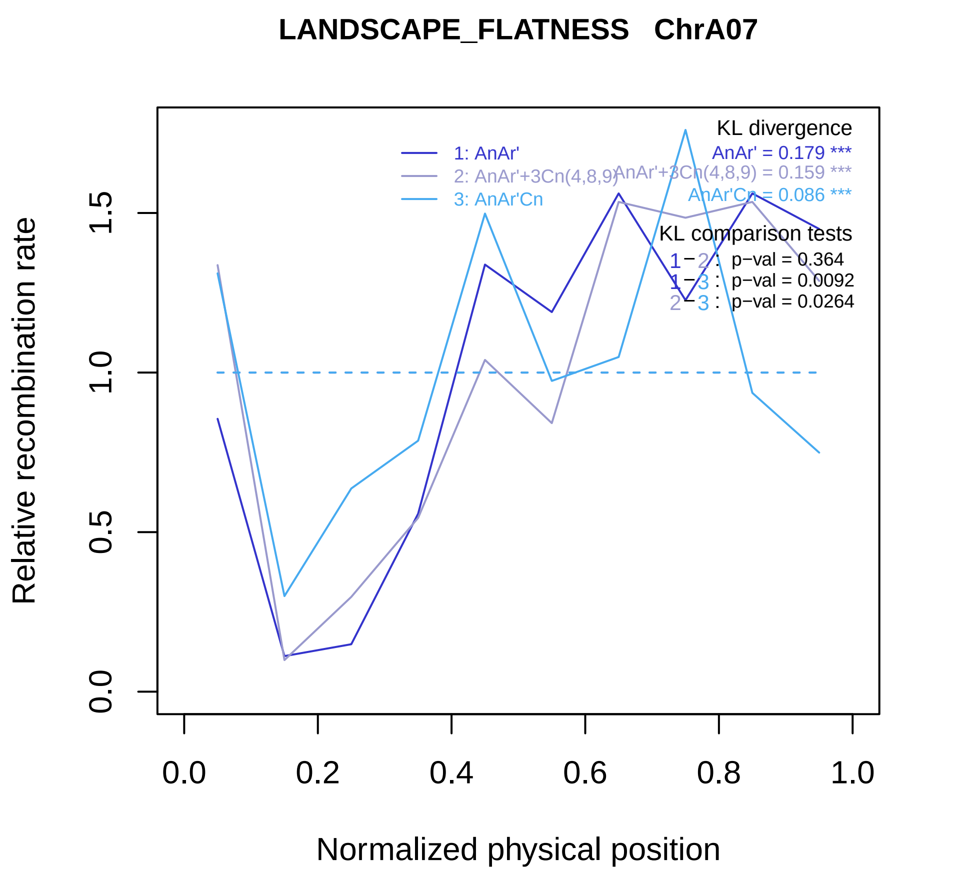

LANDSCAPE_FLATNESS ChrA07
Relative recombination rate
Normalized physical position
0.0
0.5
1.0
1.5
0.0
0.2
0.4
0.6
0.8
1.0
1: AnAr'
2: AnAr'+3Cn(4,8,9)
3: AnAr'Cn
KL divergence
AnAr' = 0.179 ***
AnAr'+3Cn(4,8,9) = 0.159 ***
AnAr'Cn = 0.086 ***
KL comparison tests
1 − 2 : p−val = 0.364
1 − 3 : p−val = 0.0092
2 − 3 : p−val = 0.0264

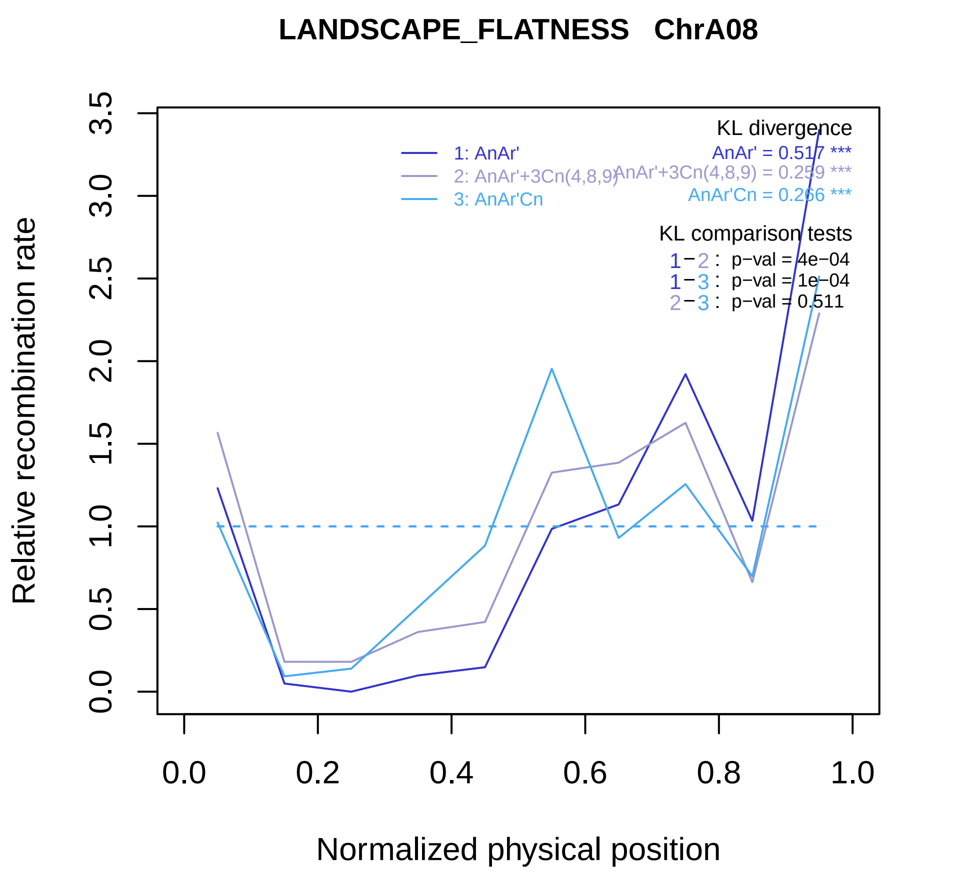
LANDSCAPE_FLATNESS ChrA08
KL divergence
1: AnAr'
2: AnAr'+3Cn(4,8,9)
3: AnAr'Cn
AnAr' = 0.517 ***
AnAr'+3Cn(4,8,9) = 0.259 ***
AnAr'Cn = 0.266 ***
KL comparison tests
1−2 : p−val = 4e−04
1−3 : p−val = 1e−04
2−3 : p−val = 0.511
Relative recombination rate
0.0
0.5
1.0
1.5
2.0
2.5
3.0
3.5
0.0
0.2
0.4
0.6
0.8
1.0
Normalized physical position

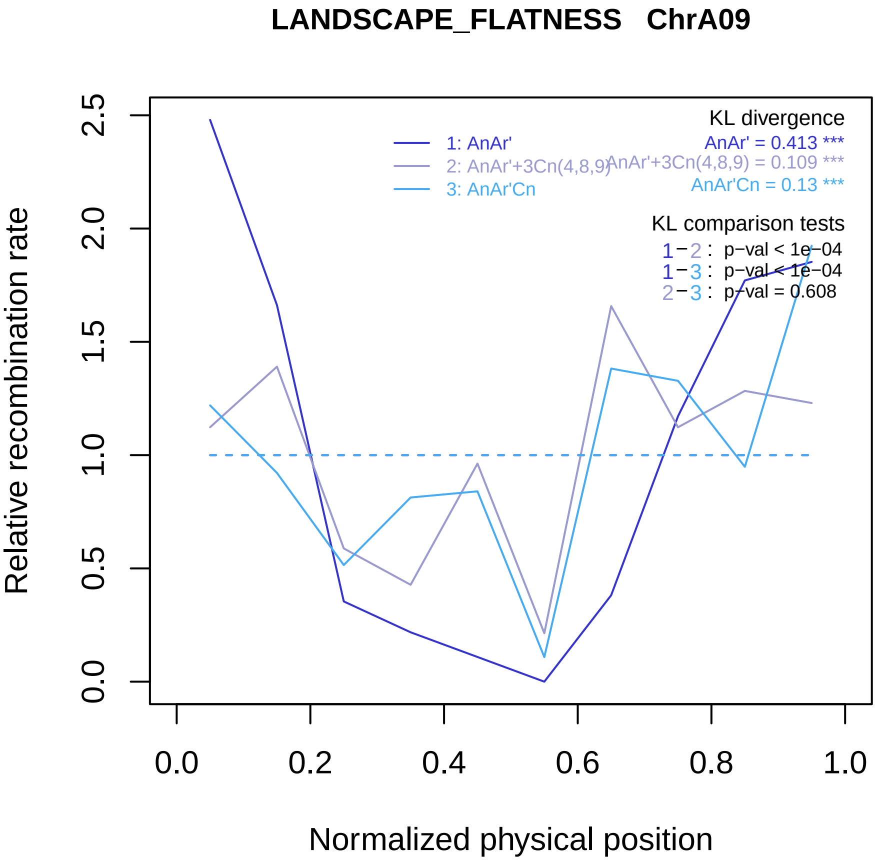

LANDSCAPE_FLATNESS ChrA09
Relative recombination rate
0.0
0.5
1.0
1.5
2.0
2.5
Normalized physical position
0.0
0.2
0.4
0.6
0.8
1.0
1: AnAr'
2: AnAr'+3Cn(4,8,9)
3: AnAr'Cn
KL divergence
AnAr' = 0.413 ***
AnAr'+3Cn(4,8,9) = 0.109 ***
AnAr'Cn = 0.13 ***
KL comparison tests
1−2 : p−val < 1e−04
1−3 : p−val < 1e−04
2−3 : p−val = 0.608

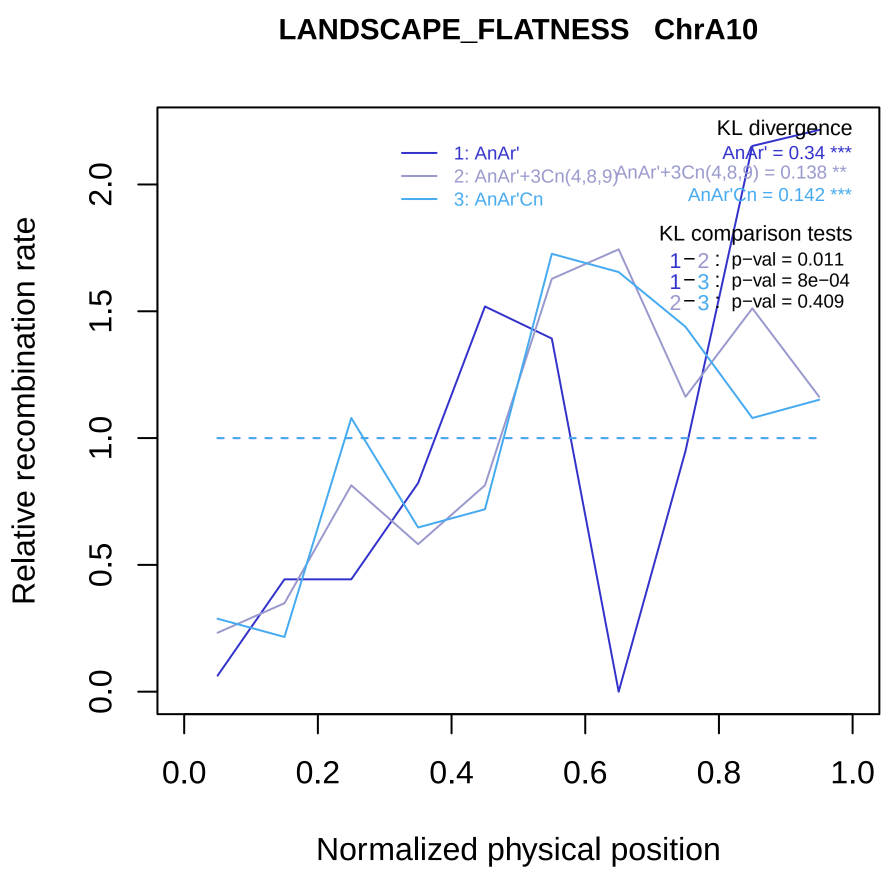

LANDSCAPE_FLATNESS ChrA10
Relative recombination rate
Normalized physical position
0.0
0.5
1.0
1.5
2.0
0.0
0.2
0.4
0.6
0.8
1.0
1: AnAr'
2: AnAr'+3Cn(4,8,9)
3: AnAr'Cn
KL divergence
AnAr' = 0.34 ***
AnAr'+3Cn(4,8,9) = 0.138 **
AnAr'Cn = 0.142 ***
KL comparison tests
1−2 : p−val = 0.011
1−3 : p−val = 8e−04
2−3 : p−val = 0.409

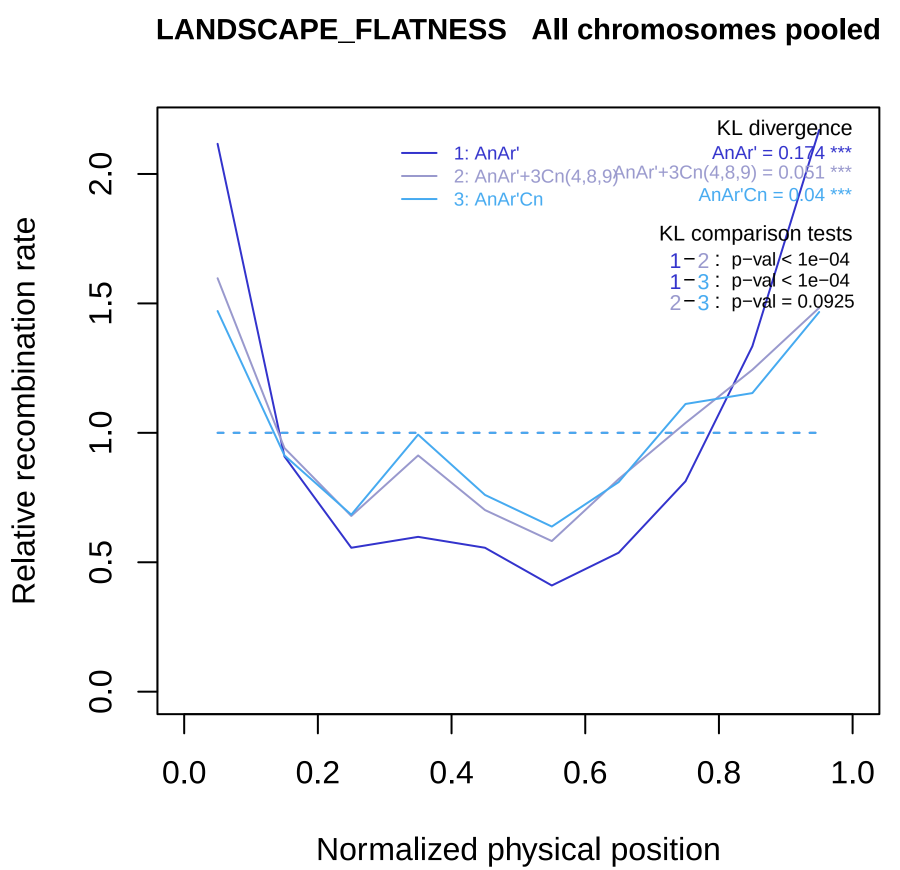
LANDSCAPE_FLATNESS All chromosomes pooled
Relative recombination rate
0.0
0.5
1.0
1.5
2.0
Normalized physical position
0.0
0.2
0.4
0.6
0.8
1.0
1: AnAr'
2: AnAr'+3Cn(4,8,9)
3: AnAr'Cn
KL divergence
AnAr' = 0.174 ***
AnAr'+3Cn(4,8,9) = 0.051 ***
AnAr'Cn = 0.04 ***
KL comparison tests
1−2 : p−val < 1e−04
1−3 : p−val < 1e−04
2−3 : p−val = 0.0925
