## Supplemental Figure 4 for "Genomic divergence shaped the genetic regulation of meiotic homologous recombination in *Brassica* allopolyploids"

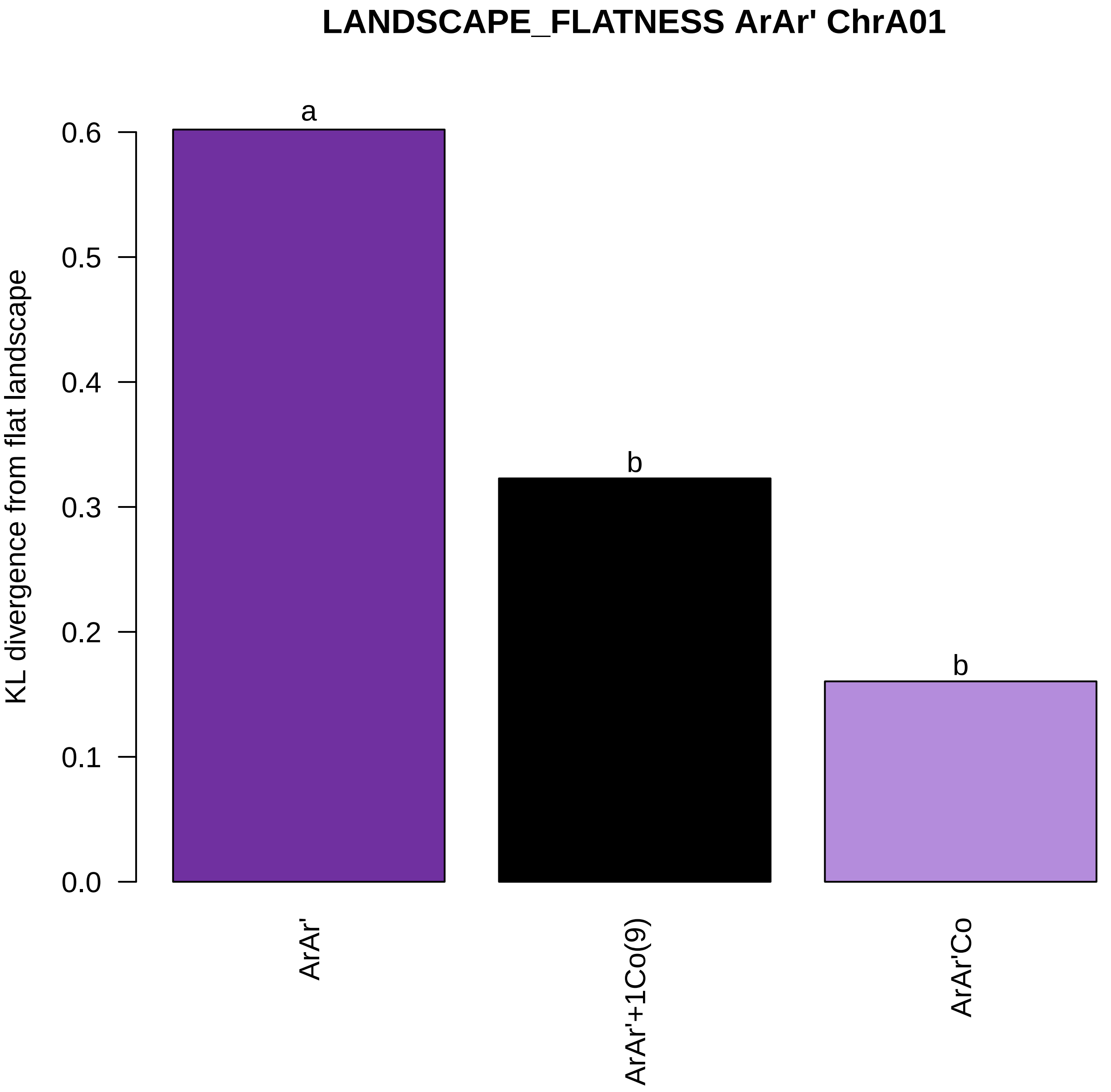

LANDSCAPE_FLATNESS ArAr' ChrA01
KL divergence from flat landscape
0.0
0.1
0.2
0.3
0.4
0.5
0.6
a
b
b
ArAr'
ArAr'+1Co(9)
ArAr'Co

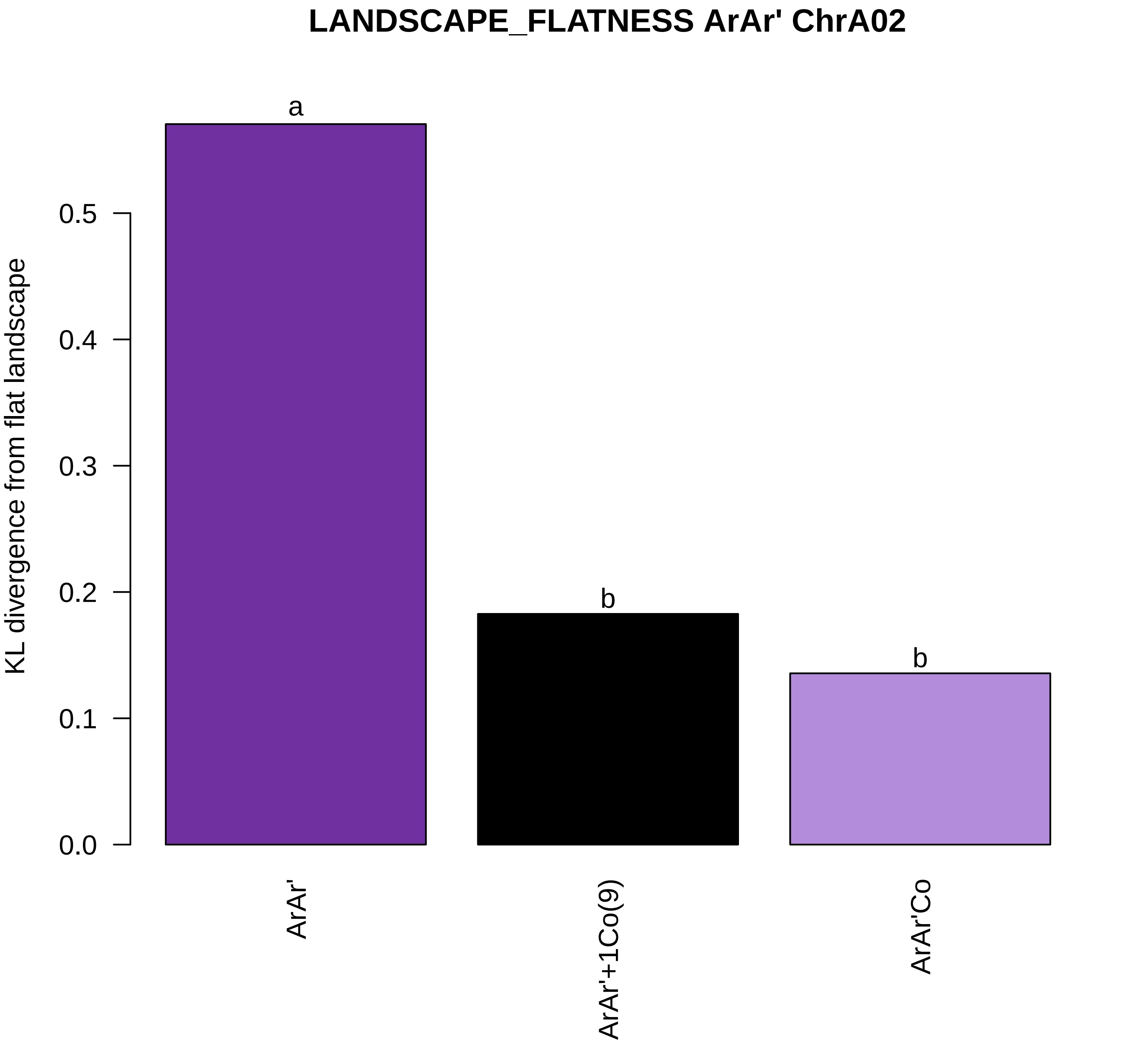

LANDSCAPE_FLATNESS ArAr' ChrA02
KL divergence from flat landscape
0.0
0.1
0.2
0.3
0.4
0.5
a
b
b
ArAr'
ArAr'+1Co(9)
ArAr'Co

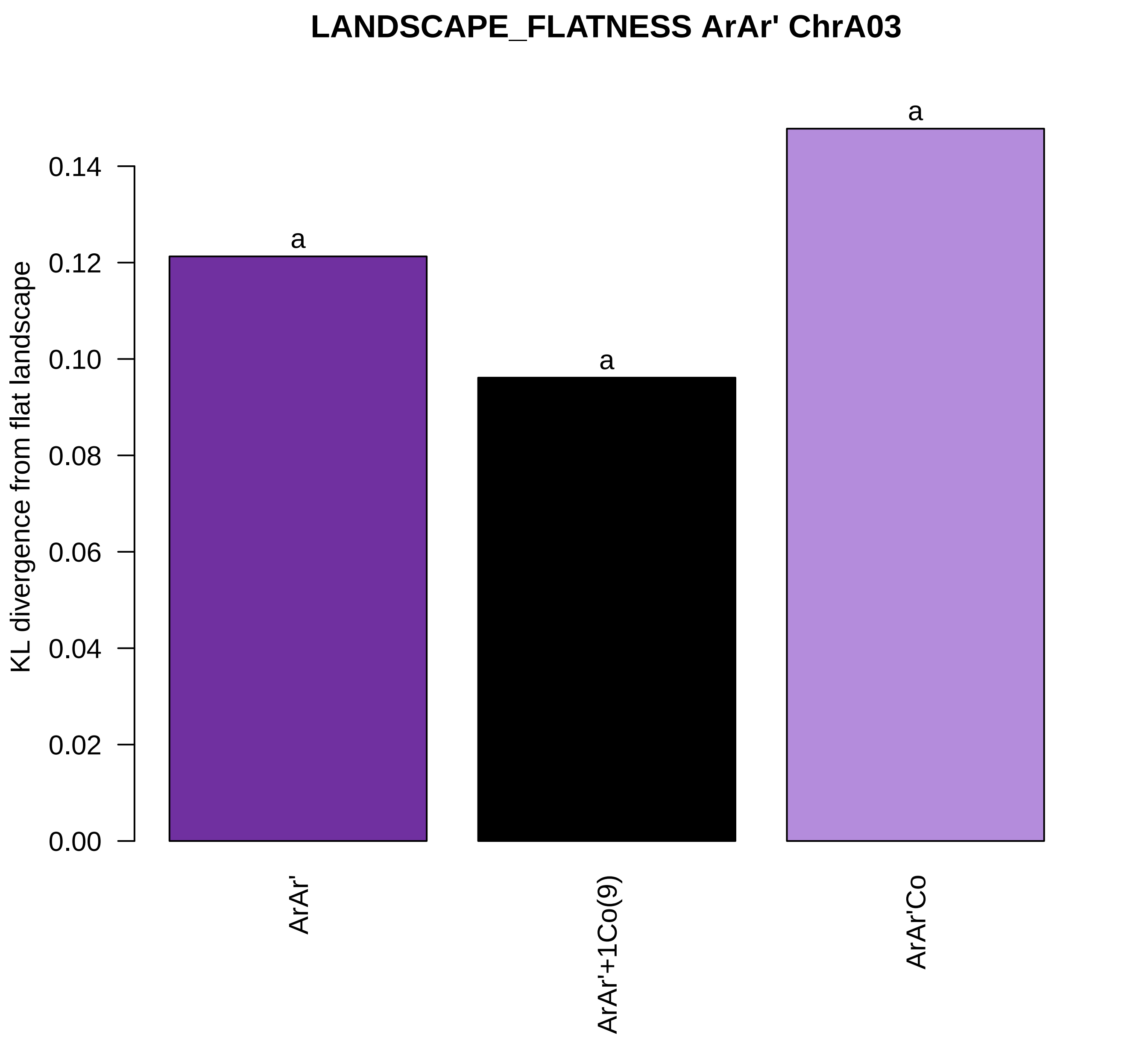
LANDSCAPE_FLATNESS ArAr' ChrA03
KL divergence from flat landscape
0.00
0.02
0.04
0.06
0.08
0.10
0.12
0.14
a
a
a
ArAr'
ArAr'+1Co(9)
ArAr'Co

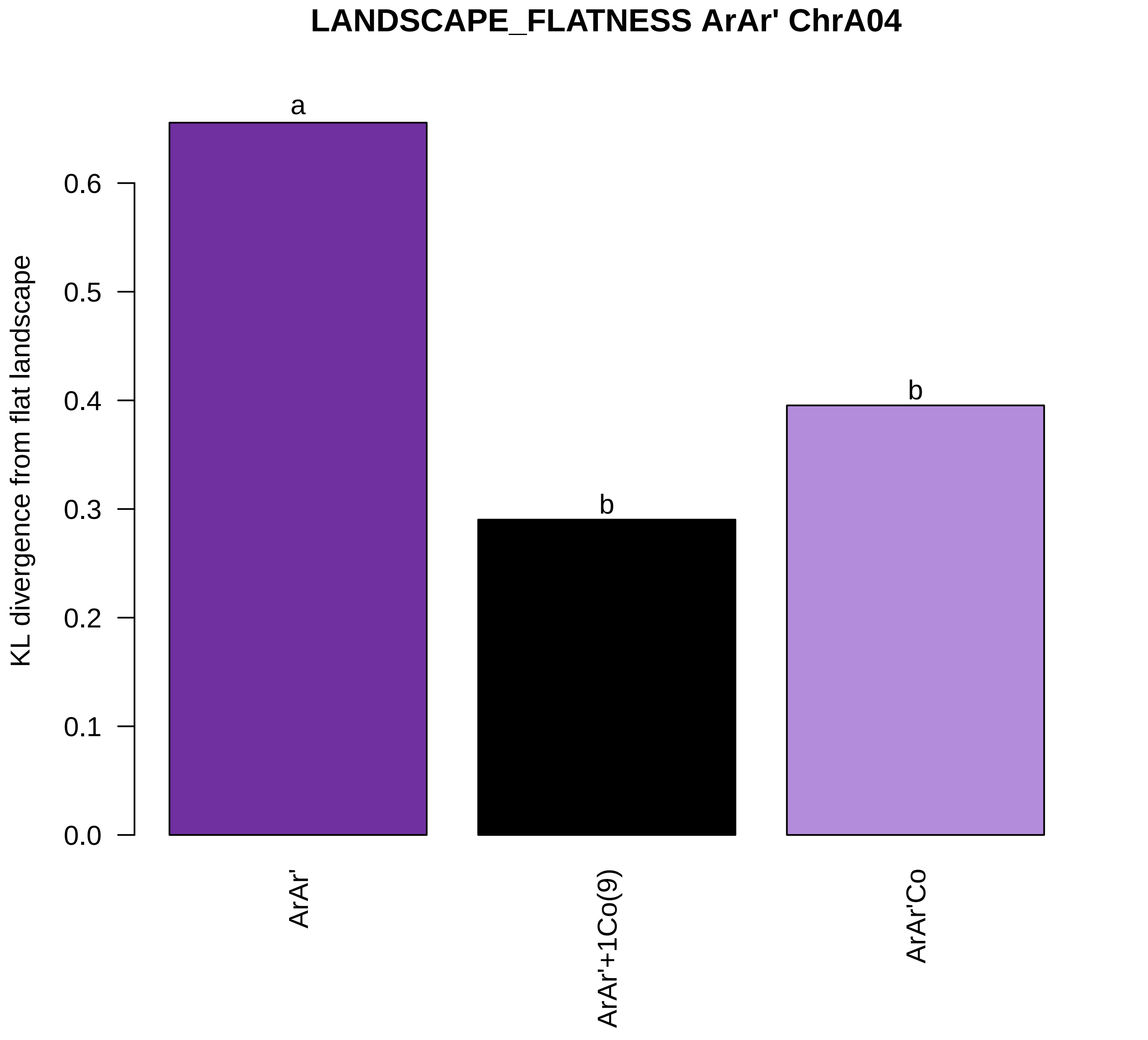

LANDSCAPE_FLATNESS ArAr' ChrA04
KL divergence from flat landscape
0.0
0.1
0.2
0.3
0.4
0.5
0.6
a
b
b
ArAr'
ArAr'+1Co(9)
ArAr'Co

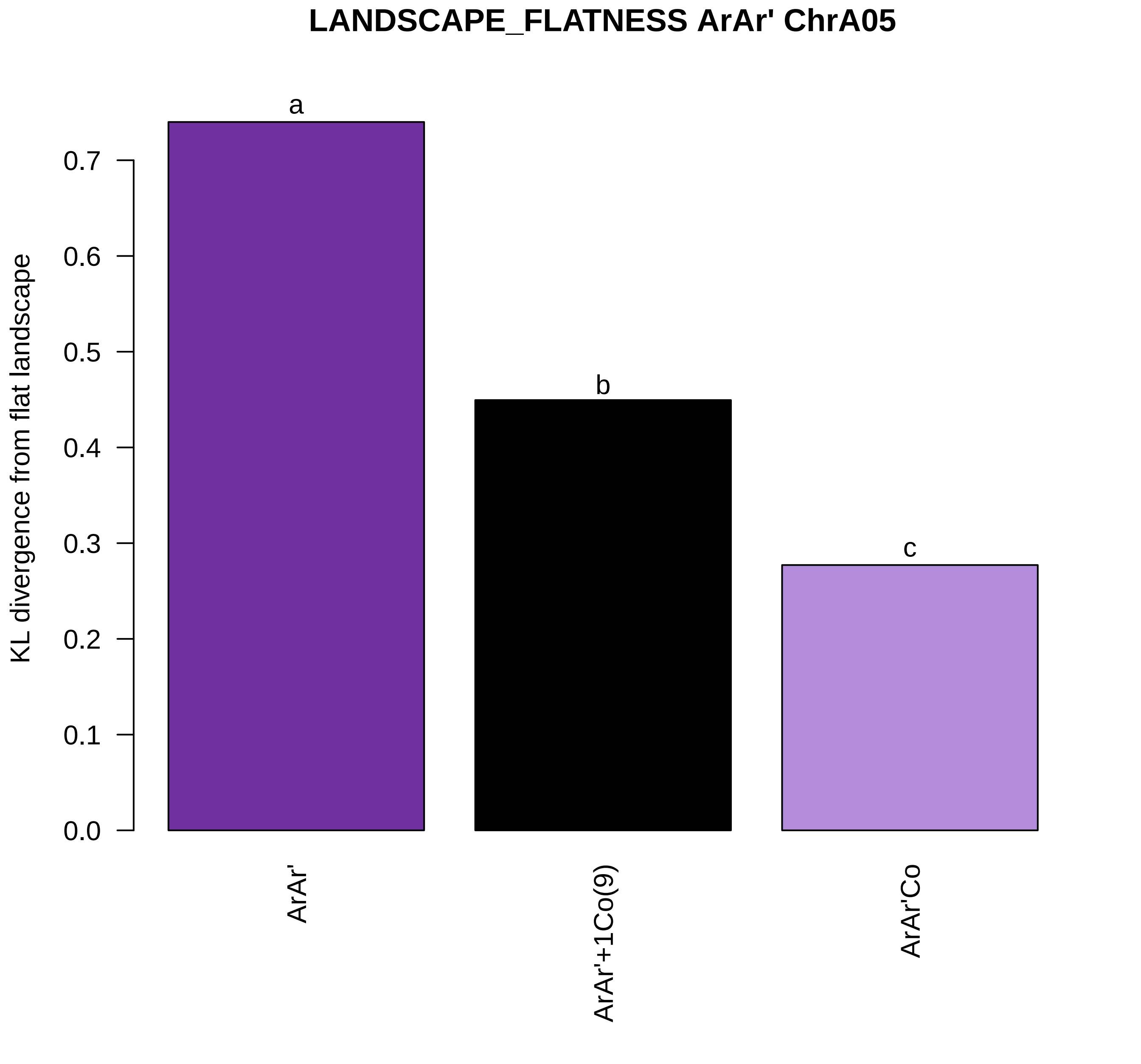

LANDSCAPE_FLATNESS ArAr' ChrA05
KL divergence from flat landscape
0.0
0.1
0.2
0.3
0.4
0.5
0.6
0.7
a
b
c
ArAr'
ArAr'+1Co(9)
ArAr'Co

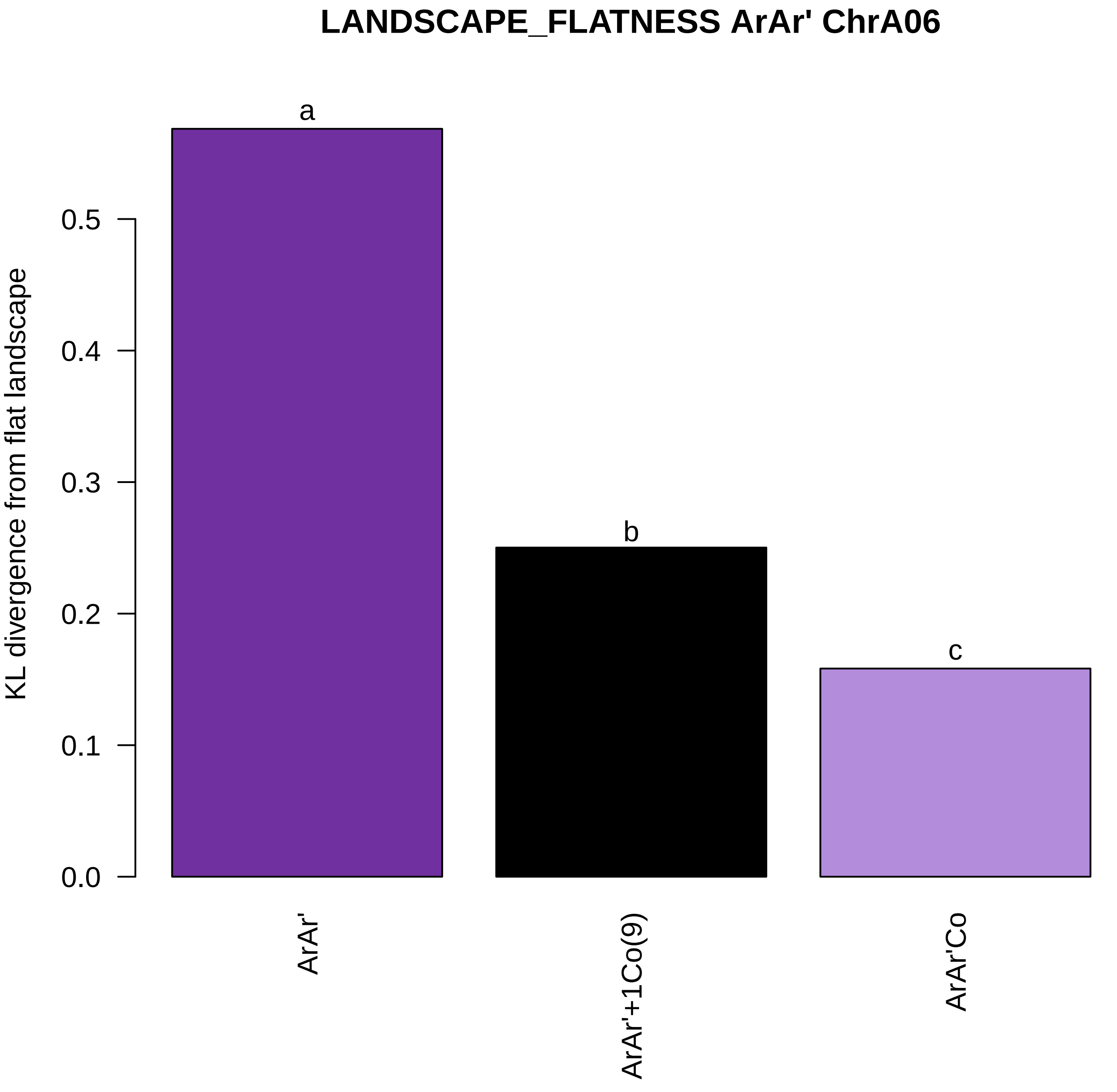

LANDSCAPE_FLATNESS ArAr' ChrA06
KL divergence from flat landscape
0.0
0.1
0.2
0.3
0.4
0.5
a
b
c
ArAr'
ArAr'+1Co(9)
ArAr'Co

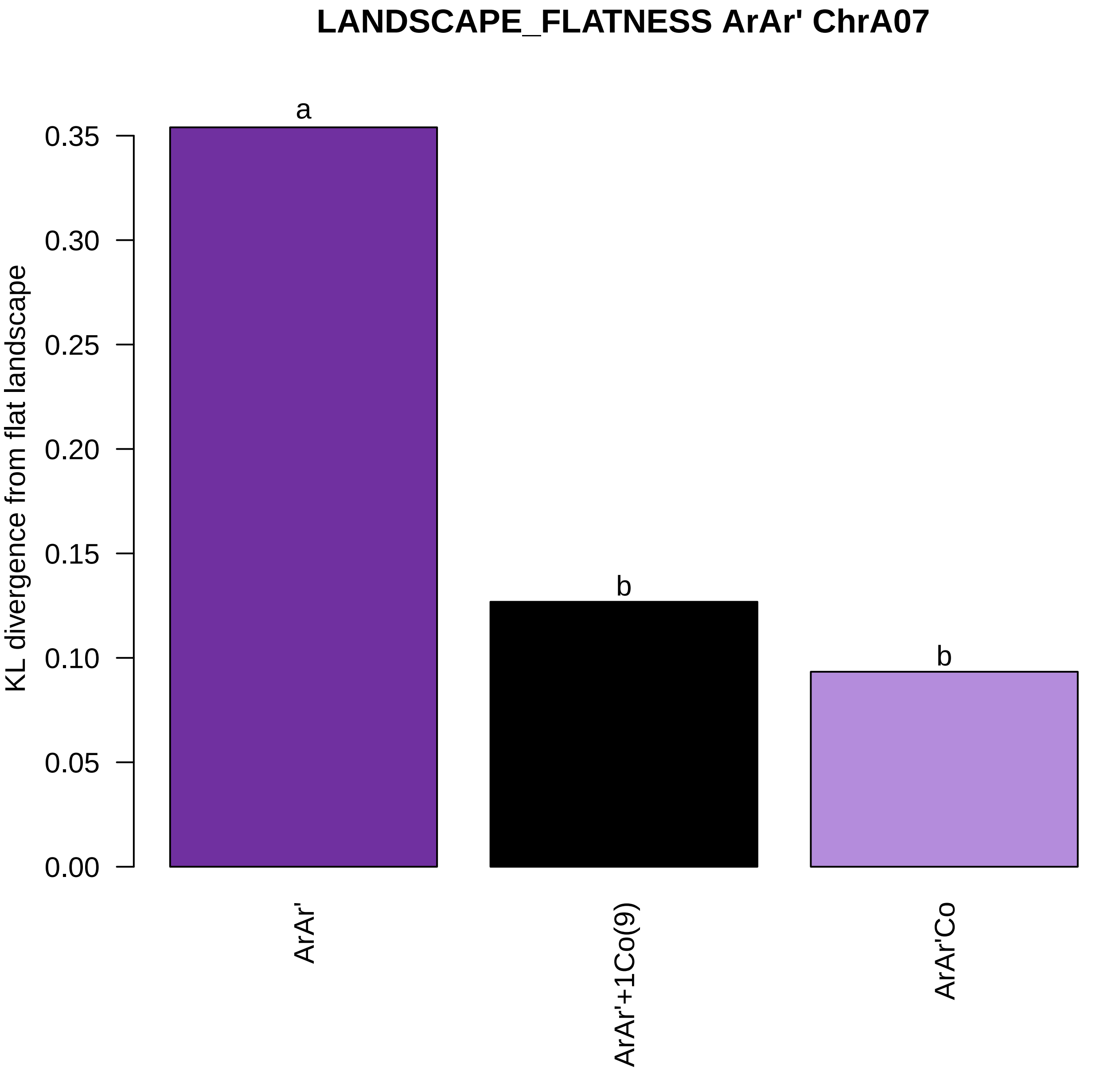
LANDSCAPE_FLATNESS ArAr' ChrA07
KL divergence from flat landscape
0.35
0.30
0.25
0.20
0.15
0.10
0.05
0.00
a
b
b
ArAr'
ArAr'+1Co(9)
ArAr'Co

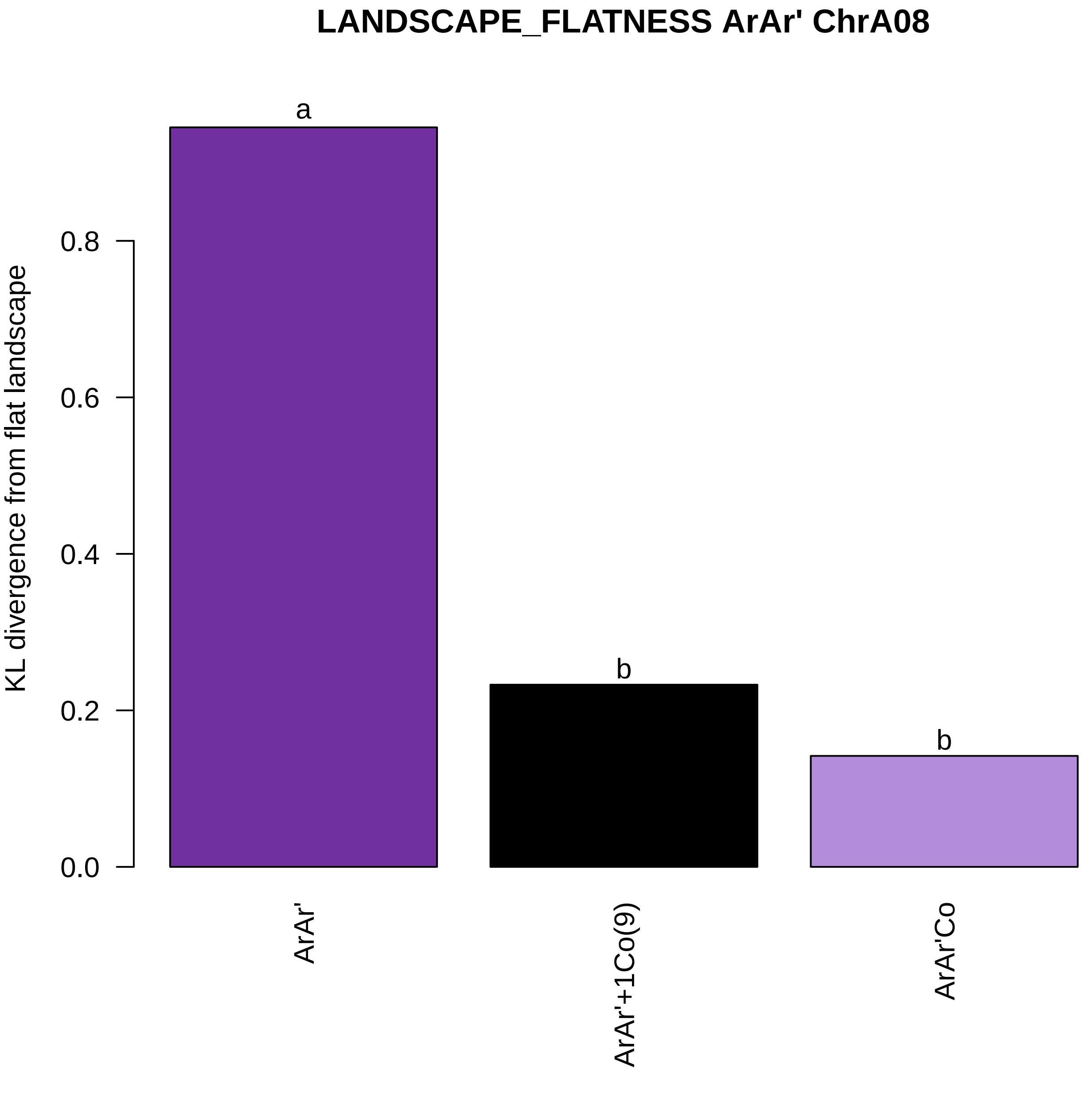
LANDSCAPE_FLATNESS ArAr' ChrA08
KL divergence from flat landscape
0.0
0.2
0.4
0.6
0.8
a
b
b
ArAr'
ArAr'+1Co(9)
ArAr'Co

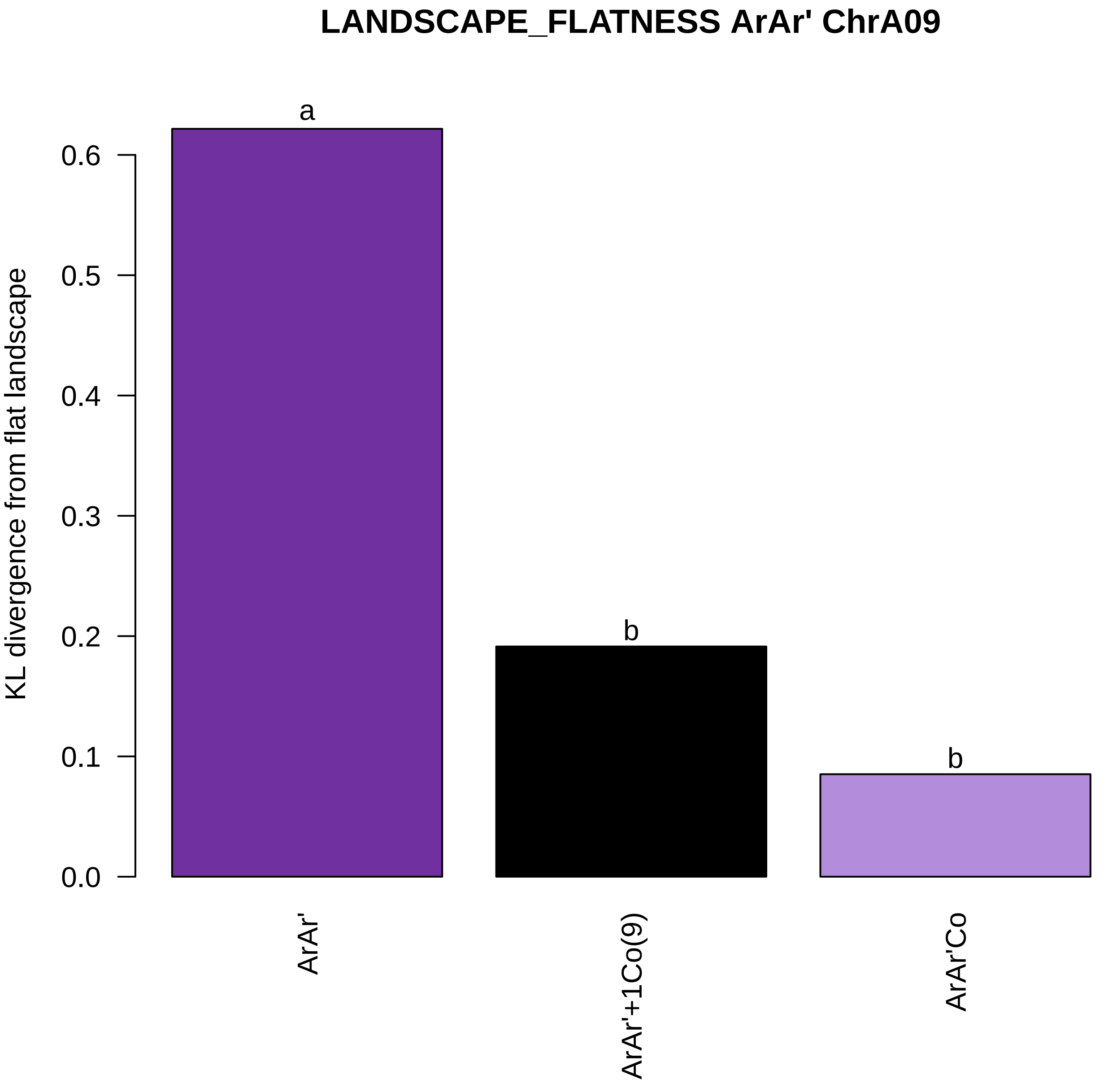
LANDSCAPE_FLATNESS ArAr' ChrA09
KL divergence from flat landscape
0.0
0.1
0.2
0.3
0.4
0.5
0.6
a
b
b
ArAr'
ArAr'+1Co(9)
ArAr'Co

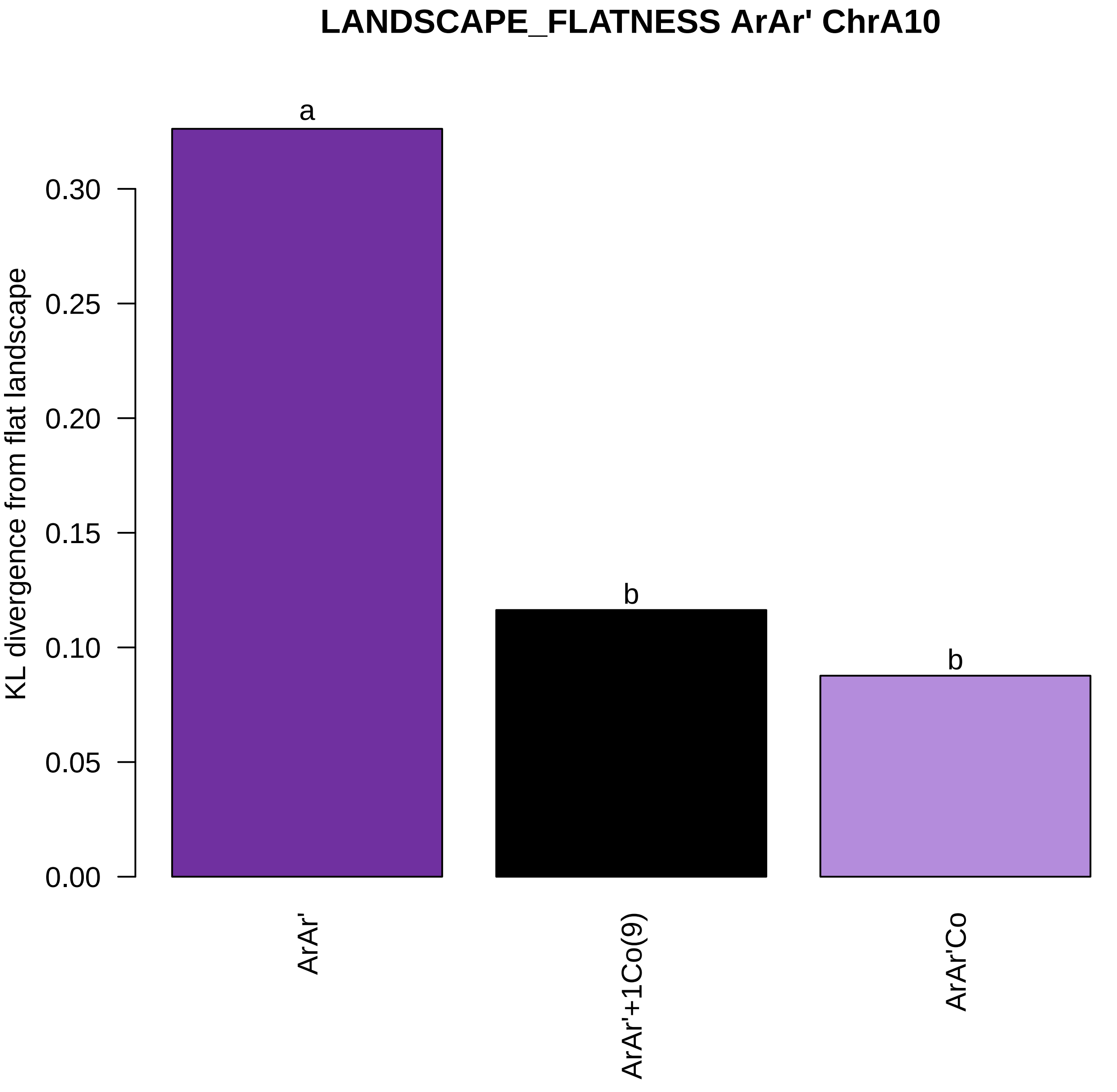

LANDSCAPE_FLATNESS ArAr' ChrA10
KL divergence from flat landscape
0.00
0.05
0.10
0.15
0.20
0.25
0.30
a
b
b
ArAr'
ArAr'+1Co(9)
ArAr'Co

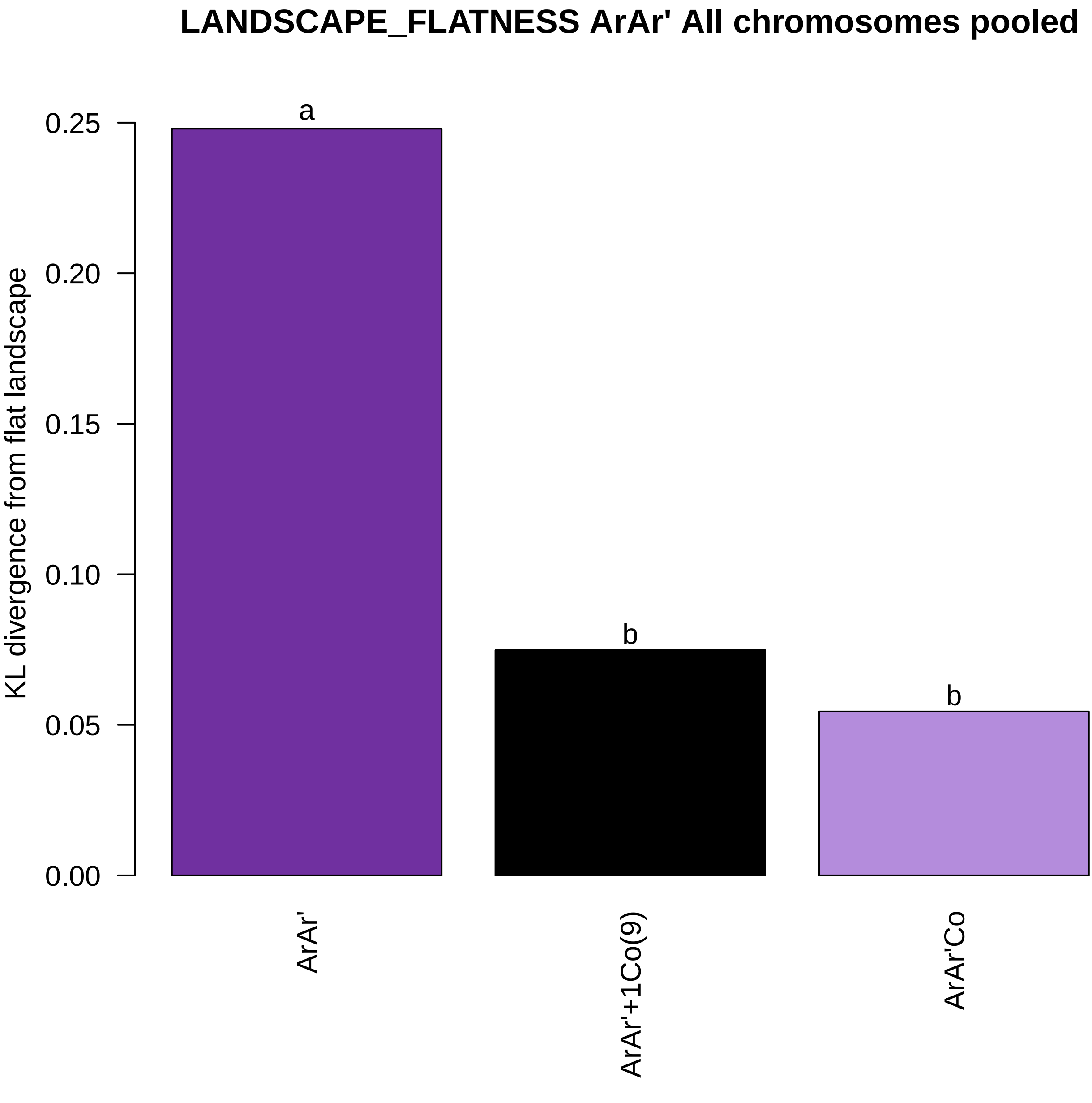
LANDSCAPE_FLATNESS ArAr' All chromosomes pooled
KL divergence from flat landscape
0.00
0.05
0.10
0.15
0.20
0.25
a
b
b
ArAr'
ArAr'+1Co(9)
ArAr'Co

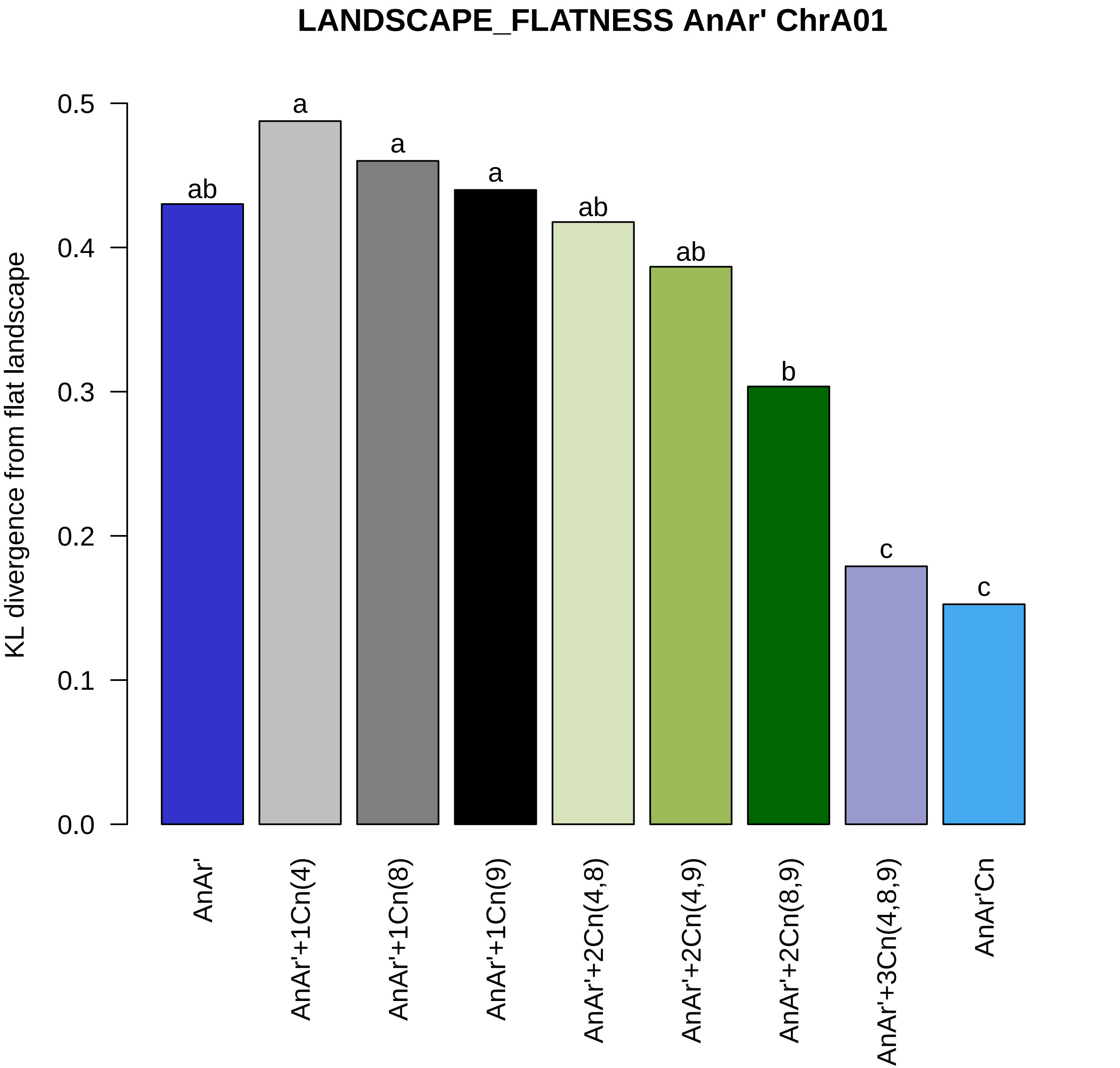

LANDSCAPE_FLATNESS AnAr' ChrA01
KL divergence from flat landscape
0.0
0.1
0.2
0.3
0.4
0.5
ab
a
a
a
ab
ab
b
c
c
AnAr'
AnAr'+1Cn(4)
AnAr'+1Cn(8)
AnAr'+1Cn(9)
AnAr'+2Cn(4,8)
AnAr'+2Cn(4,9)
AnAr'+2Cn(8,9)
AnAr'+3Cn(4,8,9)
AnAr'Cn

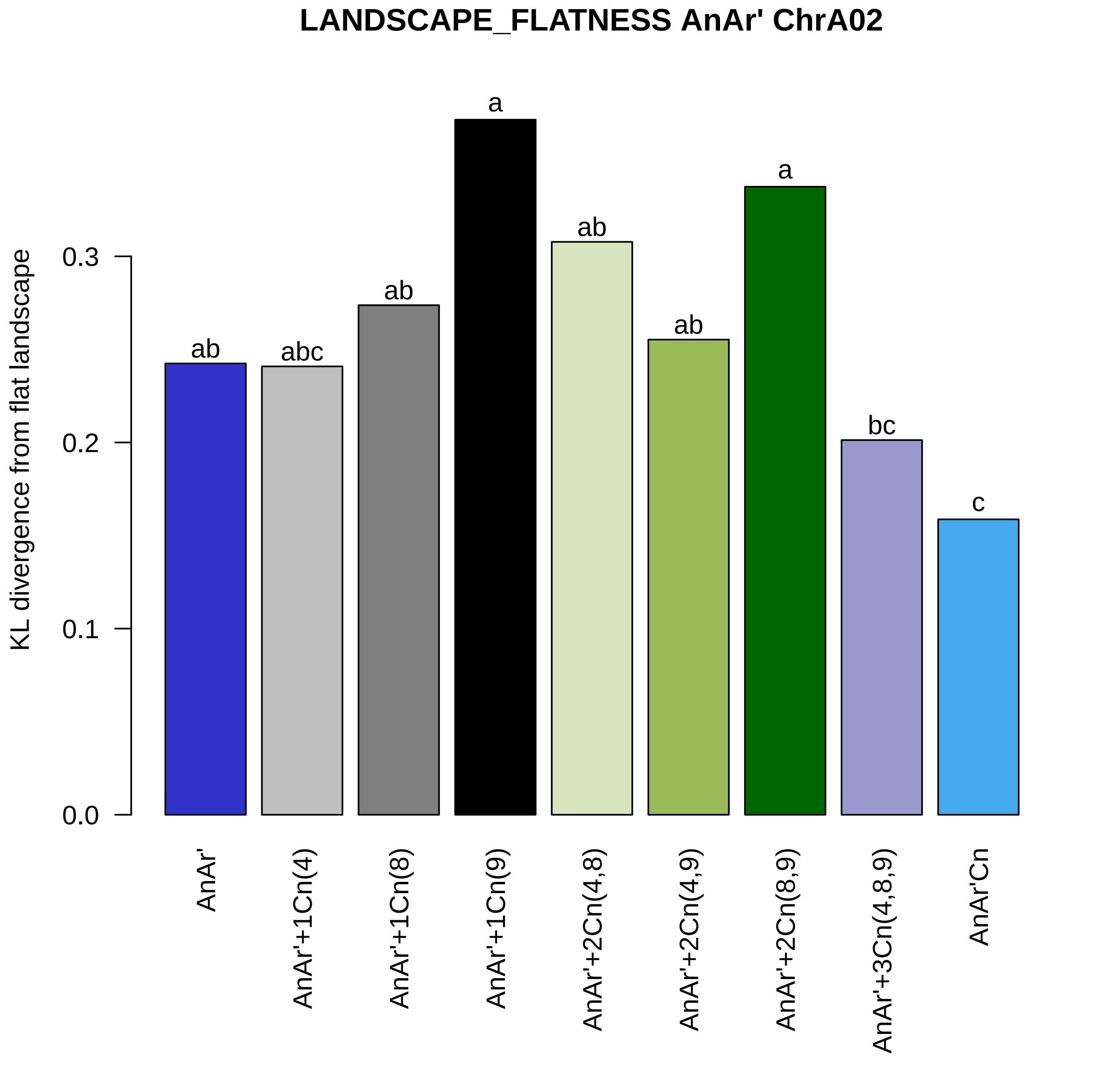
LANDSCAPE_FLATNESS AnAr' ChrA02
KL divergence from flat landscape
0.0
0.1
0.2
0.3
ab
abc
ab
a
ab
ab
a
bc
c
AnAr'
AnAr'+1Cn(4)
AnAr'+1Cn(8)
AnAr'+1Cn(9)
AnAr'+2Cn(4,8)
AnAr'+2Cn(4,9)
AnAr'+2Cn(8,9)
AnAr'+3Cn(4,8,9)
AnAr'Cn

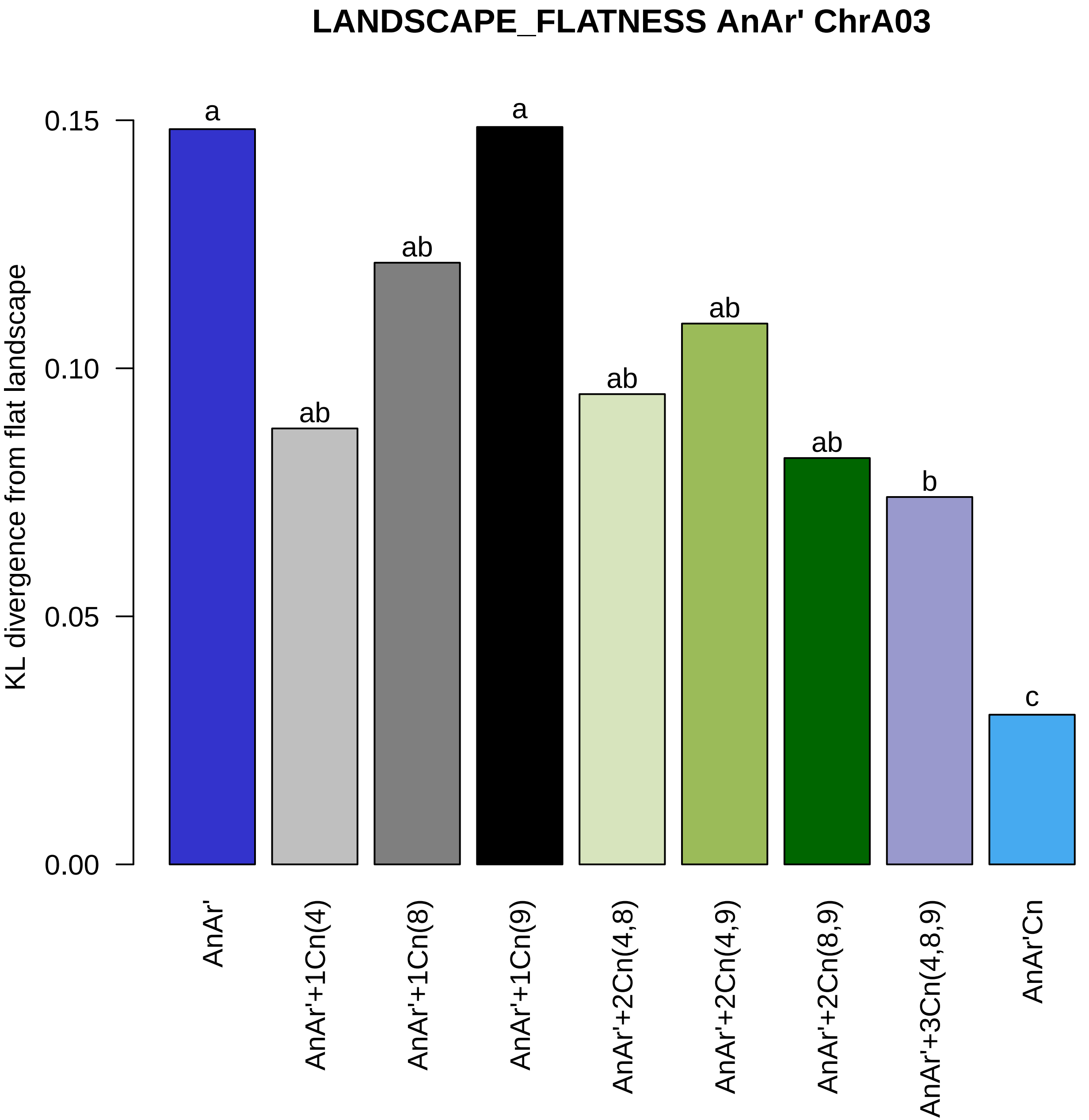
LANDSCAPE_FLATNESS AnAr' ChrA03
KL divergence from flat landscape
0.15
0.10
0.05
0.00
a
ab
ab
a
ab
ab
ab
b
c
AnAr'
AnAr'+1Cn(4)
AnAr'+1Cn(8)
AnAr'+1Cn(9)
AnAr'+2Cn(4,8)
AnAr'+2Cn(4,9)
AnAr'+2Cn(8,9)
AnAr'+3Cn(4,8,9)
AnAr'Cn

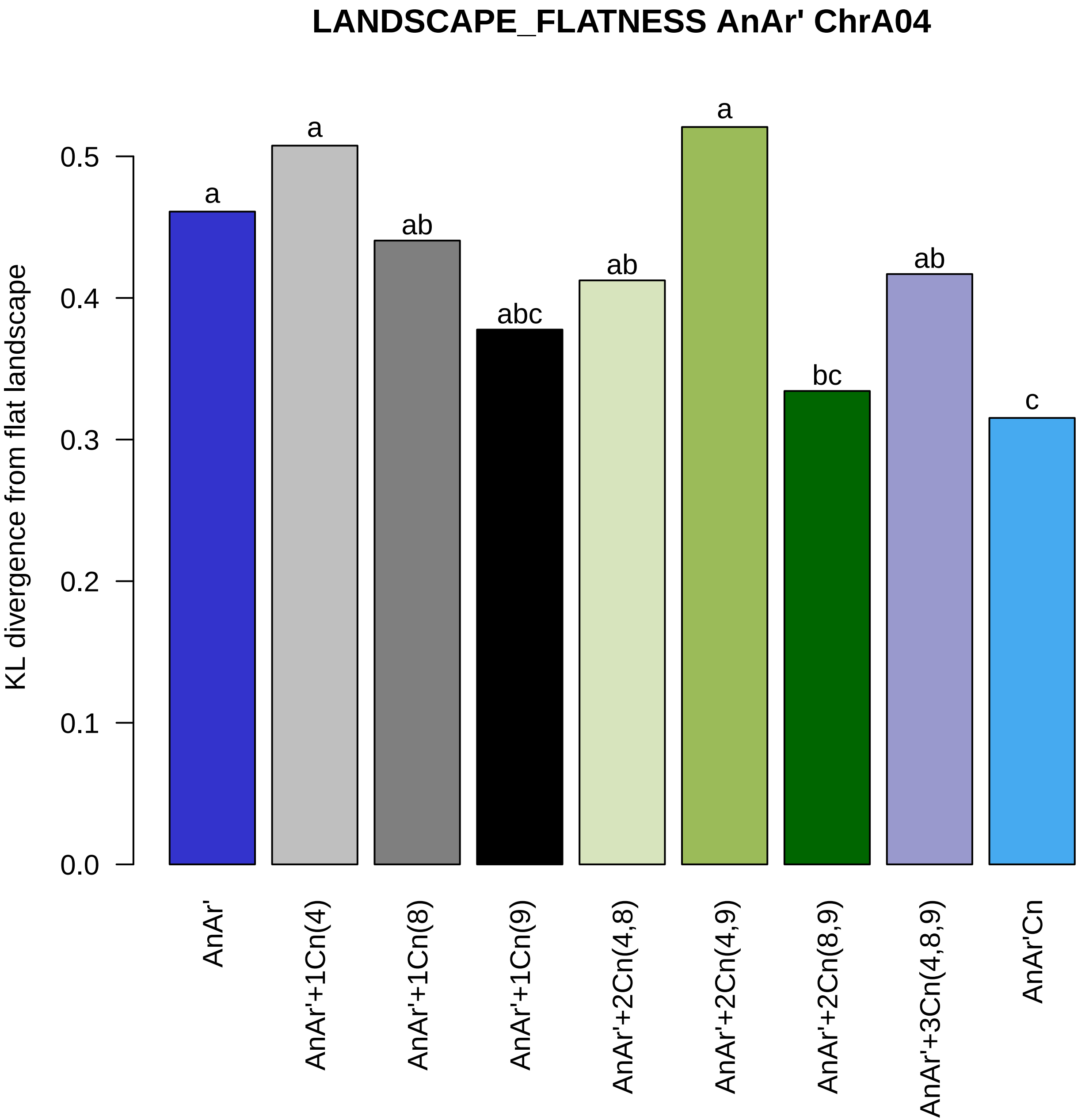

LANDSCAPE_FLATNESS AnAr' ChrA04
KL divergence from flat landscape
0.0
0.1
0.2
0.3
0.4
0.5
a
a
ab
abc
ab
a
bc
ab
c
AnAr'
AnAr'+1Cn(4)
AnAr'+1Cn(8)
AnAr'+1Cn(9)
AnAr'+2Cn(4,8)
AnAr'+2Cn(4,9)
AnAr'+2Cn(8,9)
AnAr'+3Cn(4,8,9)
AnAr'Cn

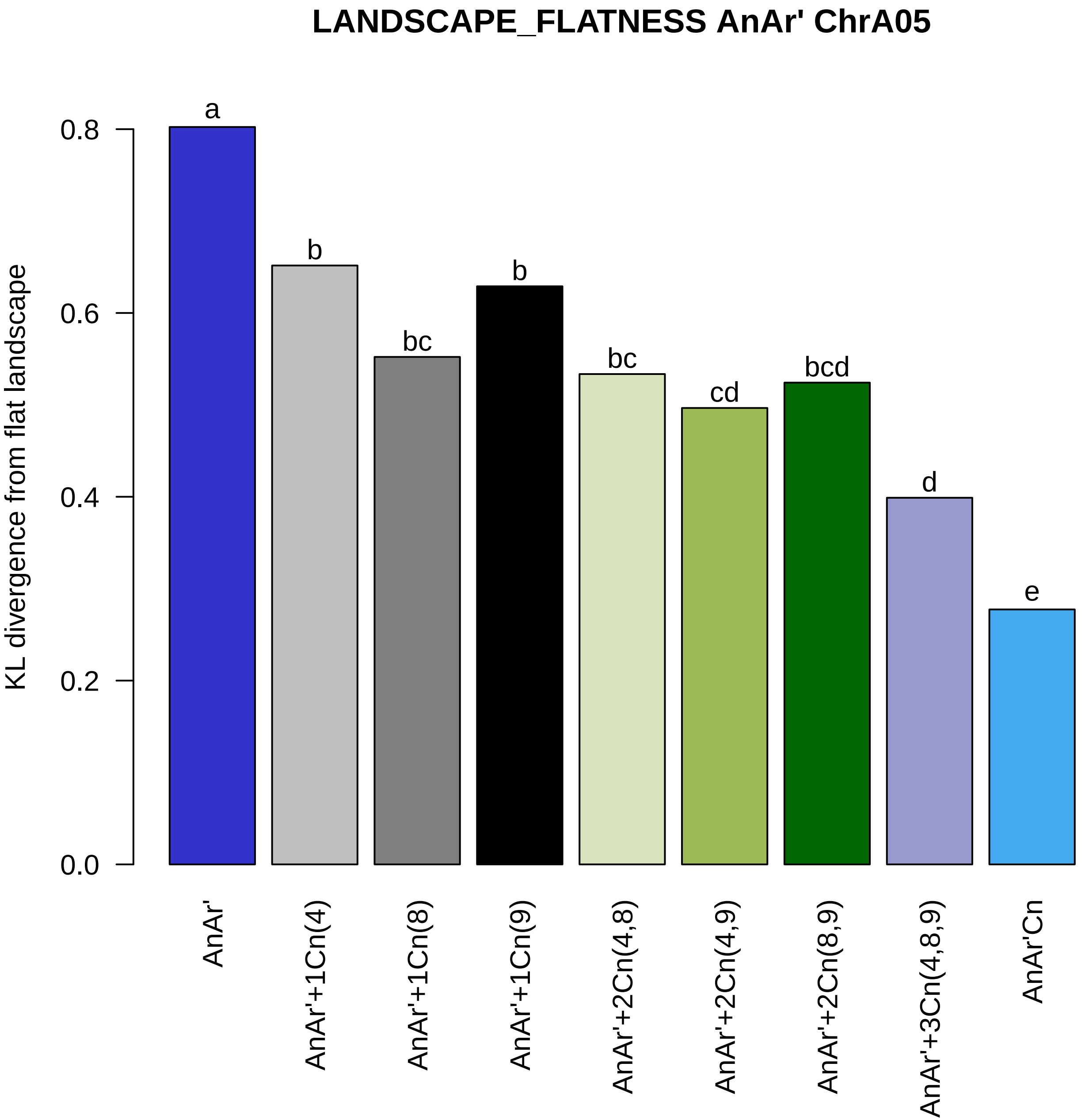

LANDSCAPE_FLATNESS AnAr' ChrA05
KL divergence from flat landscape
0.0
0.2
0.4
0.6
0.8
a
b
bc
b
bc
cd
bcd
d
e
AnAr'
AnAr'+1Cn(4)
AnAr'+1Cn(8)
AnAr'+1Cn(9)
AnAr'+2Cn(4,8)
AnAr'+2Cn(4,9)
AnAr'+2Cn(8,9)
AnAr'+3Cn(4,8,9)
AnAr'Cn

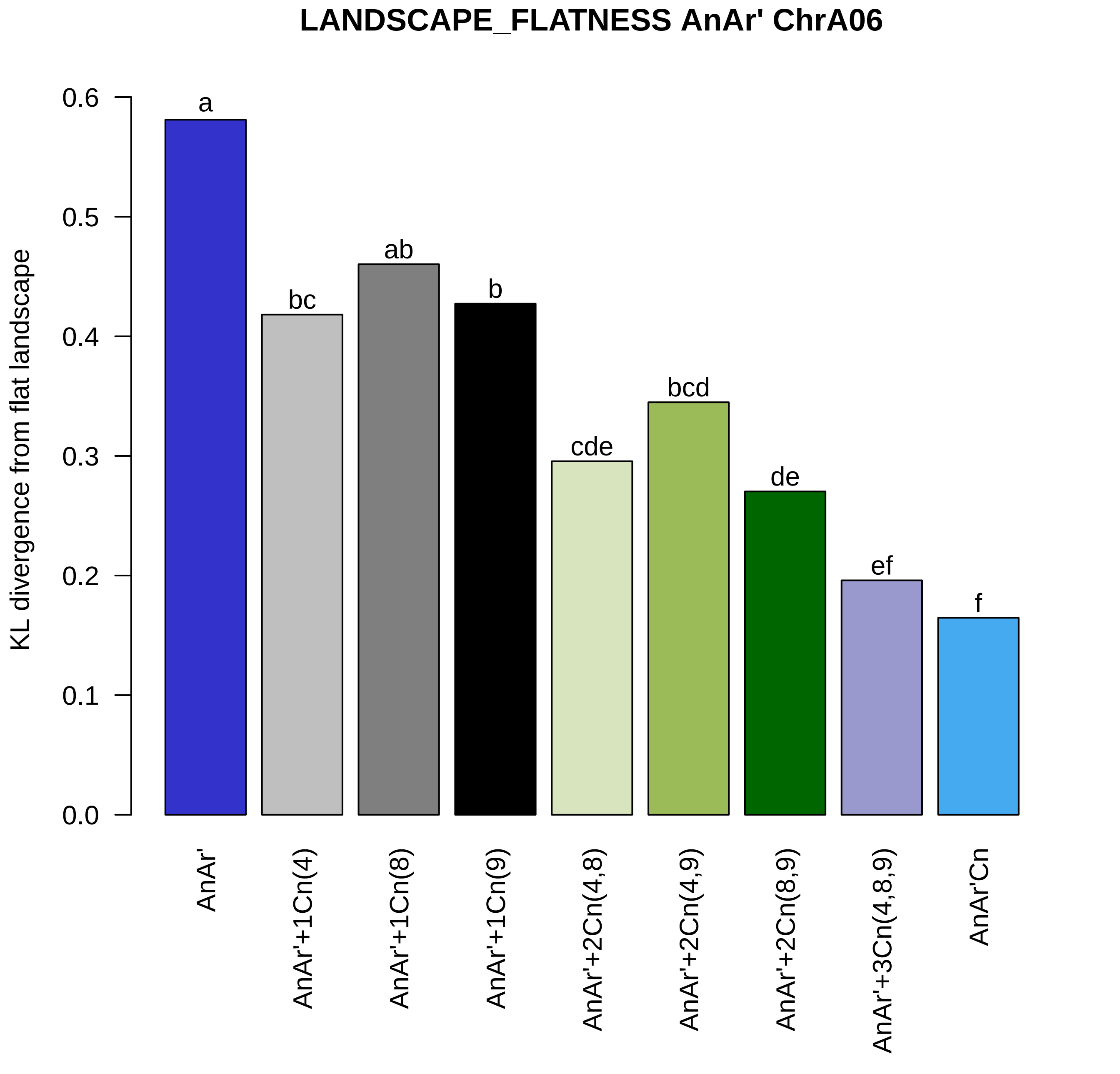

LANDSCAPE_FLATNESS AnAr' ChrA06
KL divergence from flat landscape
0.0
0.1
0.2
0.3
0.4
0.5
0.6
a
bc
ab
b
cde
bcd
de
ef
f
AnAr'
AnAr'+1Cn(4)
AnAr'+1Cn(8)
AnAr'+1Cn(9)
AnAr'+2Cn(4,8)
AnAr'+2Cn(4,9)
AnAr'+2Cn(8,9)
AnAr'+3Cn(4,8,9)
AnAr'Cn

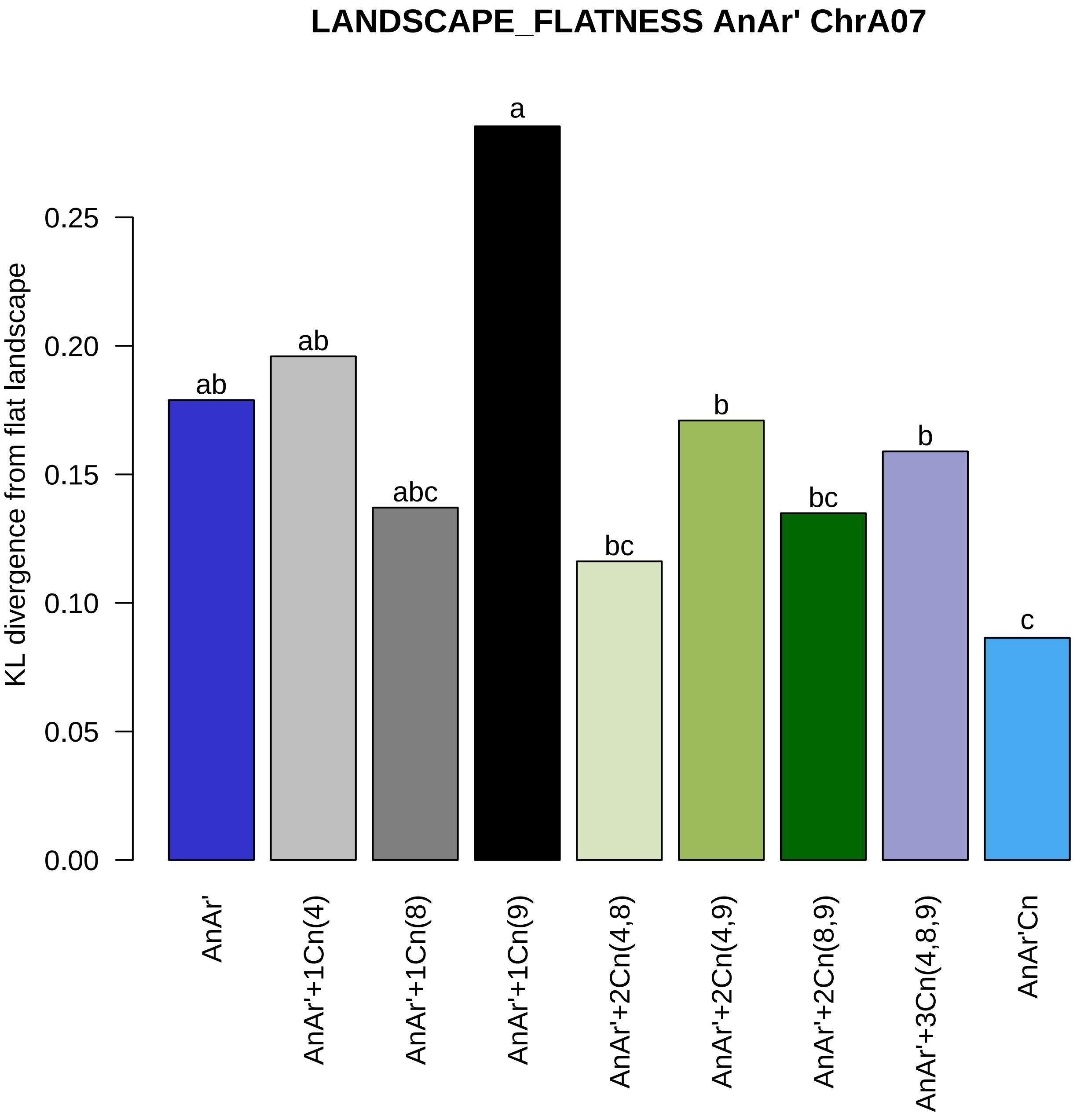

LANDSCAPE_FLATNESS AnAr' ChrA07
KL divergence from flat landscape
0.00
0.05
0.10
0.15
0.20
0.25
ab
ab
abc
a
bc
b
bc
b
c
AnAr'
AnAr'+1Cn(4)
AnAr'+1Cn(8)
AnAr'+1Cn(9)
AnAr'+2Cn(4,8)
AnAr'+2Cn(4,9)
AnAr'+2Cn(8,9)
AnAr'+3Cn(4,8,9)
AnAr'Cn

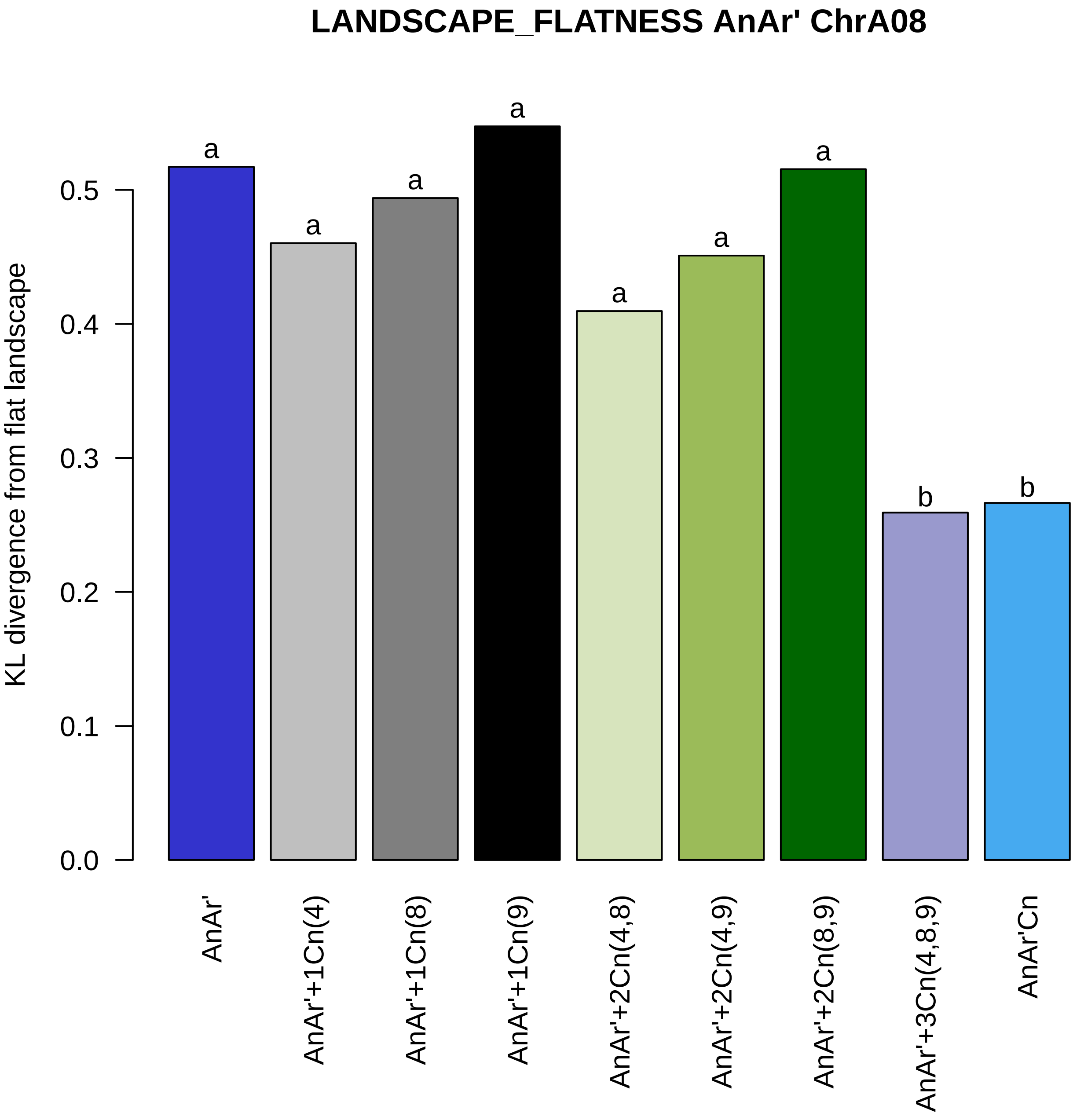

LANDSCAPE_FLATNESS AnAr' ChrA08
KL divergence from flat landscape
0.0
0.1
0.2
0.3
0.4
0.5
a
a
a
a
a
a
a
b
b
AnAr'
AnAr'+1Cn(4)
AnAr'+1Cn(8)
AnAr'+1Cn(9)
AnAr'+2Cn(4,8)
AnAr'+2Cn(4,9)
AnAr'+2Cn(8,9)
AnAr'+3Cn(4,8,9)
AnAr'Cn

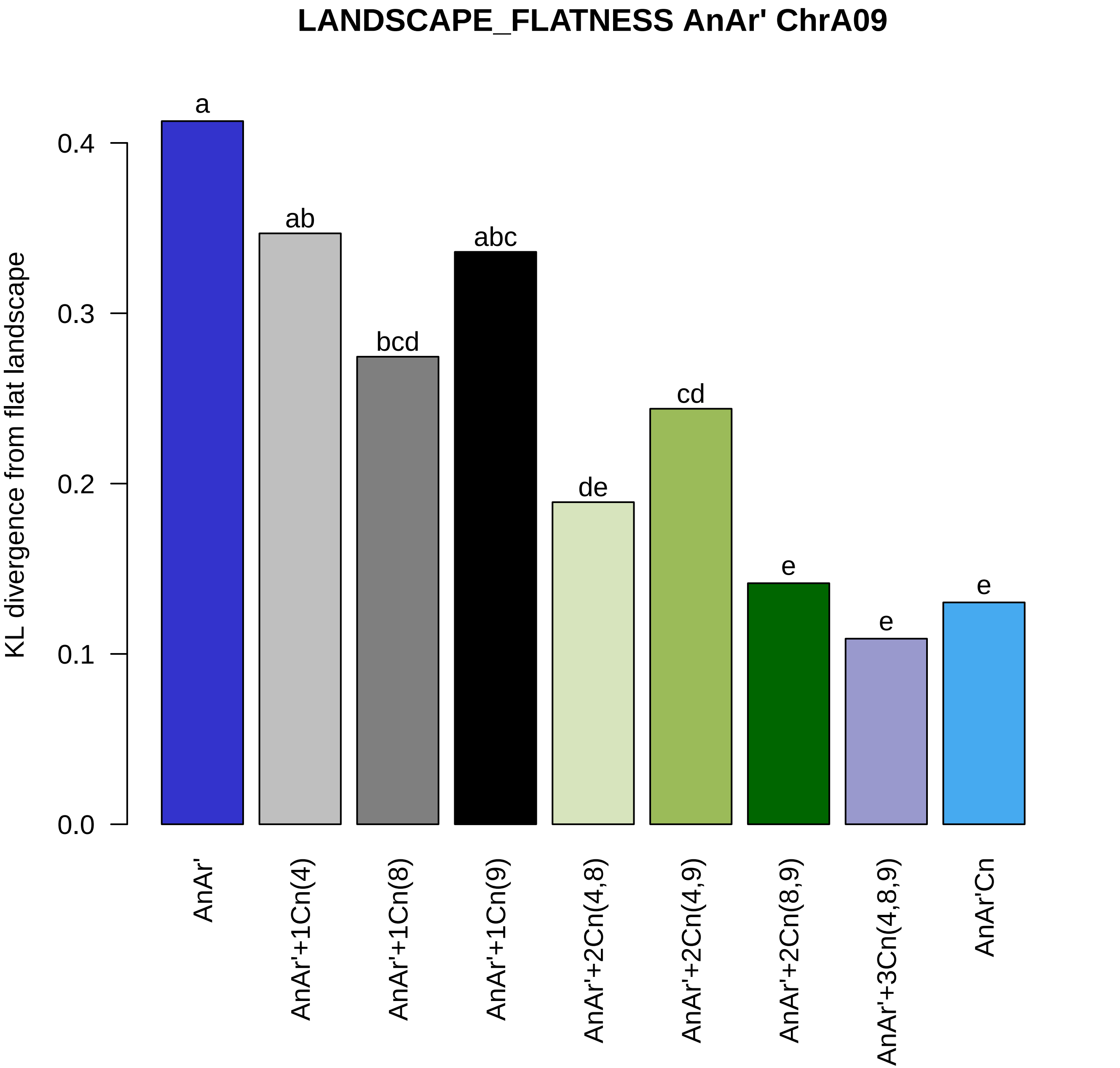
LANDSCAPE_FLATNESS AnAr' ChrA09
KL divergence from flat landscape
0.0
0.1
0.2
0.3
0.4
a
ab
bcd
abc
de
cd
e
e
e
AnAr'
AnAr'+1Cn(4)
AnAr'+1Cn(8)
AnAr'+1Cn(9)
AnAr'+2Cn(4,8)
AnAr'+2Cn(4,9)
AnAr'+2Cn(8,9)
AnAr'+3Cn(4,8,9)
AnAr'Cn

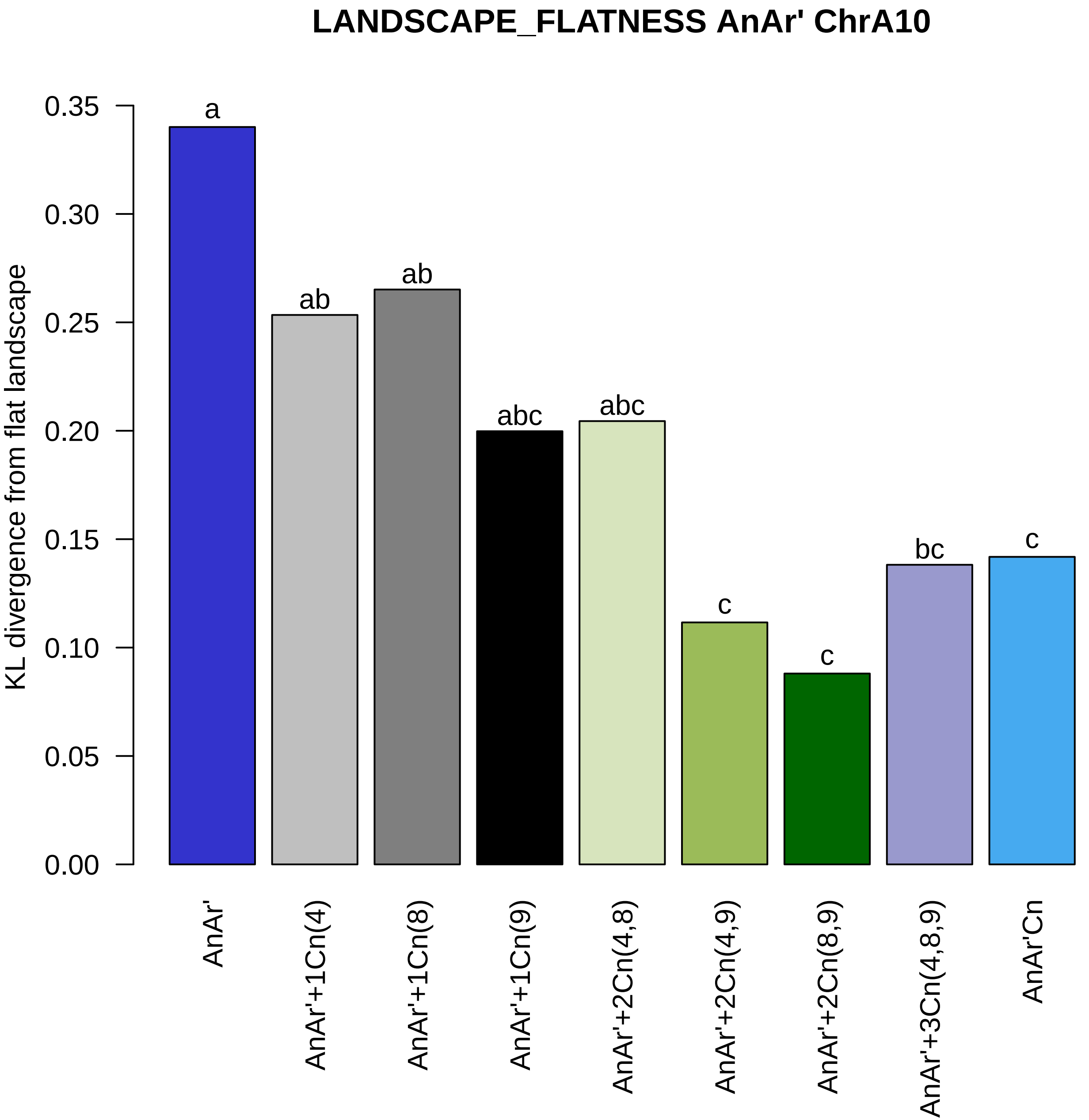
LANDSCAPE_FLATNESS AnAr' ChrA10
KL divergence from flat landscape
0.35
0.30
0.25
0.20
0.15
0.10
0.05
0.00
a
ab
ab
abc
abc
c
c
bc
c
AnAr'
AnAr'+1Cn(4)
AnAr'+1Cn(8)
AnAr'+1Cn(9)
AnAr'+2Cn(4,8)
AnAr'+2Cn(4,9)
AnAr'+2Cn(8,9)
AnAr'+3Cn(4,8,9)
AnAr'Cn

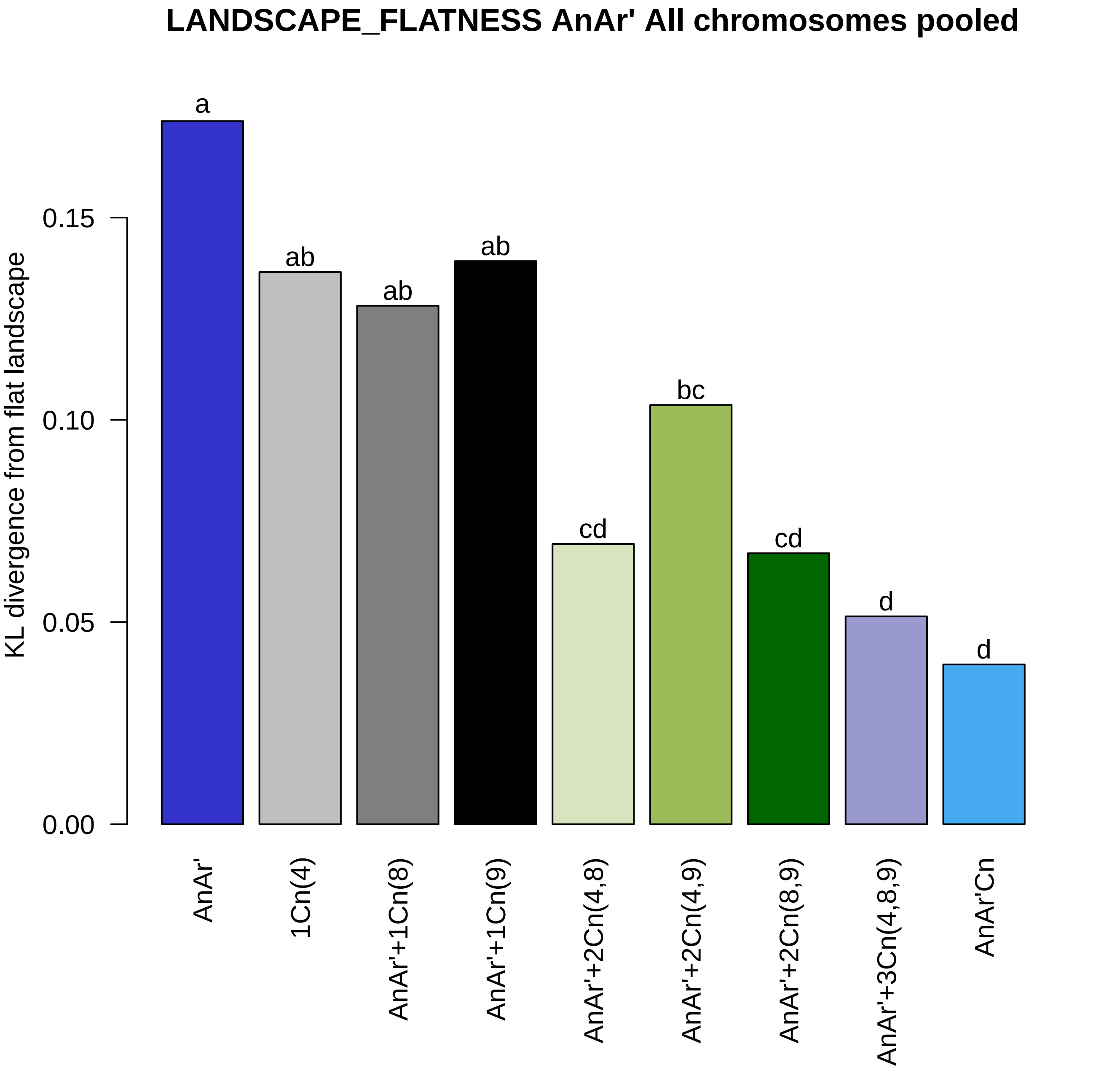

LANDSCAPE_FLATNESS AnAr' All chromosomes pooled
KL divergence from flat landscape
0.00
0.05
0.10
0.15
a
ab
ab
ab
cd
bc
cd
d
d
AnAr'
1Cn(4)
AnAr'+1Cn(8)
AnAr'+1Cn(9)
AnAr'+2Cn(4,8)
AnAr'+2Cn(4,9)
AnAr'+2Cn(8,9)
AnAr'+3Cn(4,8,9)
AnAr'Cn
