## Supplemental Figure 6 for "Genomic divergence shaped the genetic regulation of meiotic homologous recombination in *Brassica* allopolyploids"

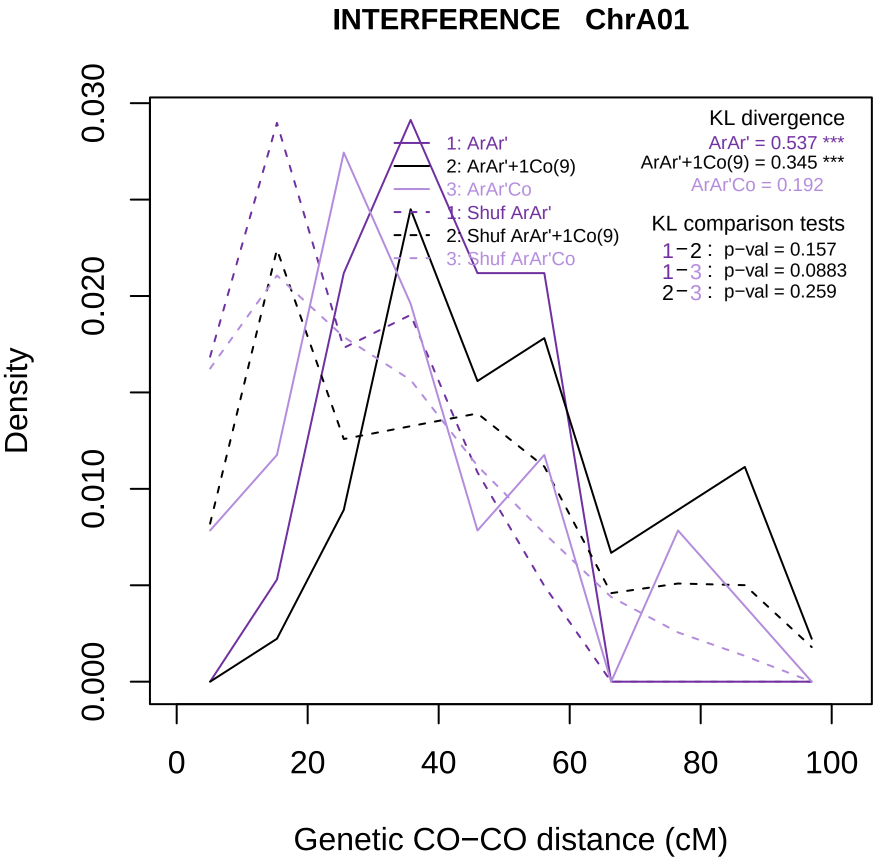
INTERFERENCE ChrA01
Density
0.000
0.010
0.020
0.030
0
20
40
60
80
100
Genetic CO−CO distance (cM)
1: ArAr'
2: ArAr'+1Co(9)
3: ArAr'Co
1: Shuf ArAr'
2: Shuf ArAr'+1Co(9)
3: Shuf ArAr'Co
KL divergence
ArAr' = 0.537 ***
ArAr'+1Co(9) = 0.345 ***
ArAr'Co = 0.192
KL comparison tests
1−2 : p−val = 0.157
1−3 : p−val = 0.0883
2−3 : p−val = 0.259

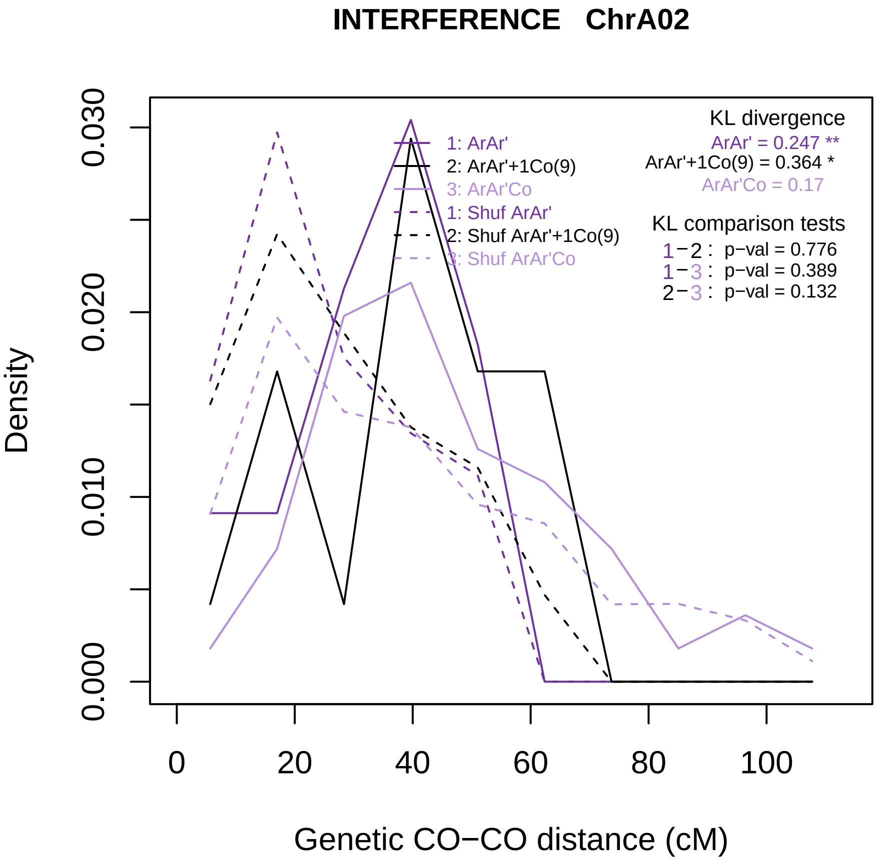
INTERFERENCE ChrA02
Density
0.000
0.010
0.020
0.030
0
20
40
60
80
100
Genetic CO−CO distance (cM)
1: ArAr'
2: ArAr'+1Co(9)
3: ArAr'Co
1: Shuf ArAr'
2: Shuf ArAr'+1Co(9)
3: Shuf ArAr'Co
KL divergence
ArAr' = 0.247 **
ArAr'+1Co(9) = 0.364 *
ArAr'Co = 0.17
KL comparison tests
1−2 : p−val = 0.776
1−3 : p−val = 0.389
2−3 : p−val = 0.132

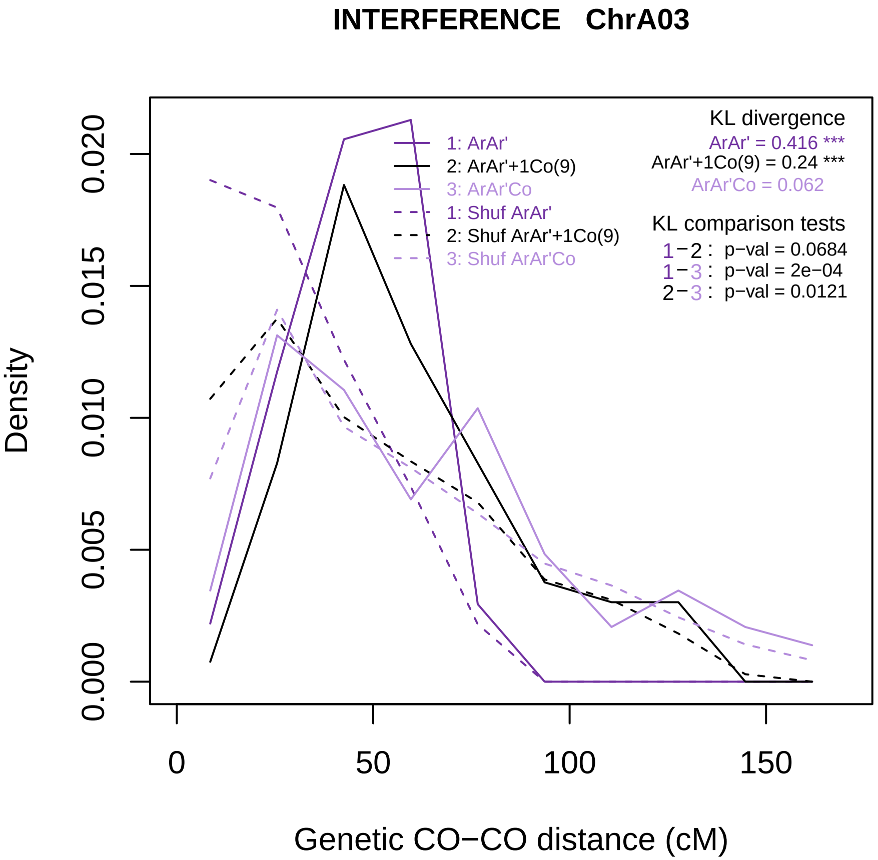

INTERFERENCE ChrA03
Density
Genetic CO−CO distance (cM)
0.000
0.005
0.010
0.015
0.020
0
50
100
150
1: ArAr'
2: ArAr'+1Co(9)
3: ArAr'Co
1: Shuf ArAr'
2: Shuf ArAr'+1Co(9)
3: Shuf ArAr'Co
KL divergence
ArAr' = 0.416 ***
ArAr'+1Co(9) = 0.24 ***
ArAr'Co = 0.062
KL comparison tests
1−2 : p−val = 0.0684
1−3 : p−val = 2e−04
2−3 : p−val = 0.0121

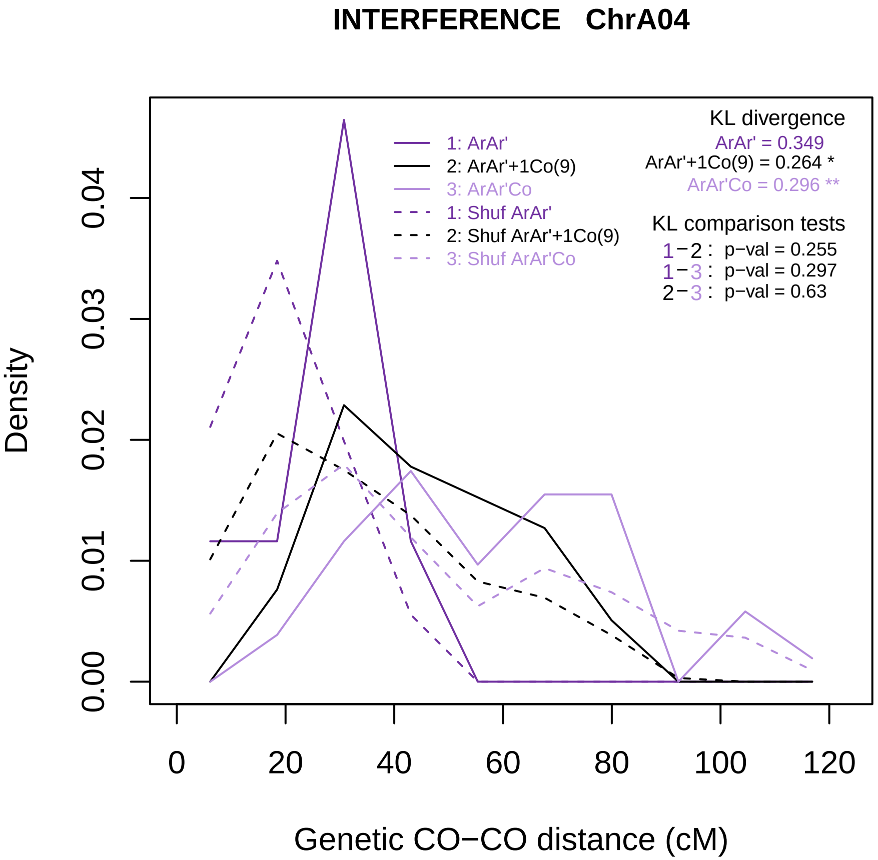
INTERFERENCE ChrA04
Density
0.00
0.01
0.02
0.03
0.04
0
20
40
60
80
100
120
Genetic CO−CO distance (cM)
1: ArAr'
2: ArAr'+1Co(9)
3: ArAr'Co
1: Shuf ArAr'
2: Shuf ArAr'+1Co(9)
3: Shuf ArAr'Co
KL divergence
ArAr' = 0.349
ArAr'+1Co(9) = 0.264 *
ArAr'Co = 0.296 **
KL comparison tests
1−2 : p−val = 0.255
1−3 : p−val = 0.297
2−3 : p−val = 0.63

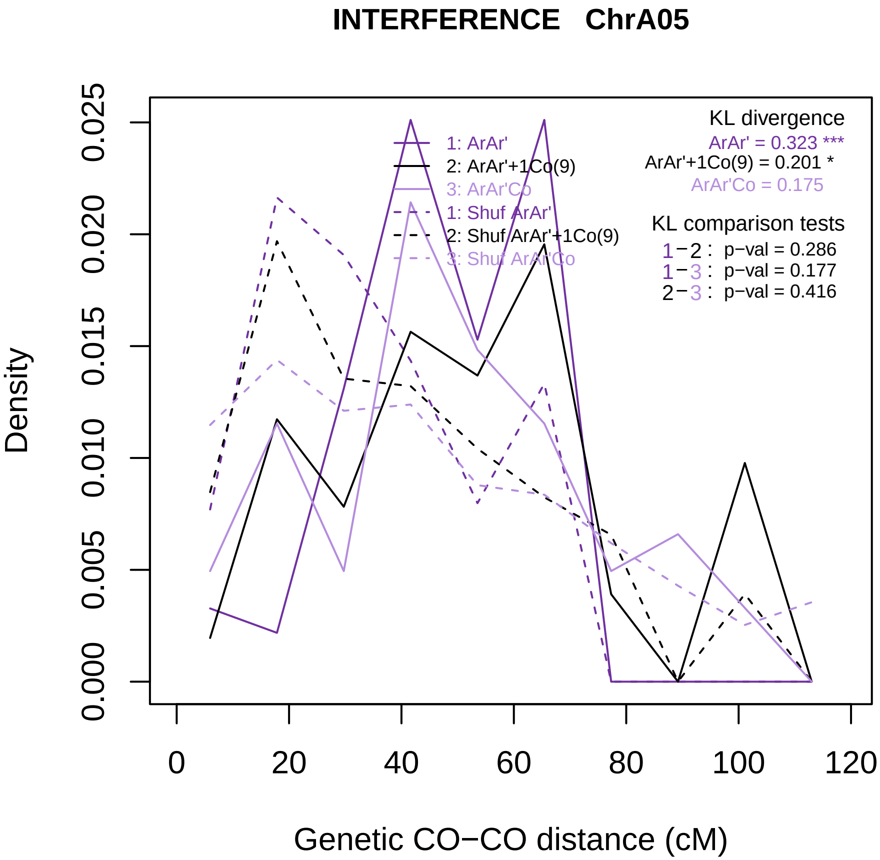
INTERFERENCE ChrA05
Density
0.000
0.005
0.010
0.015
0.020
0.025
0
20
40
60
80
100
120
Genetic CO−CO distance (cM)
1: ArAr'
2: ArAr'+1Co(9)
3: ArAr'Co
1: Shuf ArAr'
2: Shuf ArAr'+1Co(9)
3: Shuf ArAr'Co
KL divergence
ArAr' = 0.323 ***
ArAr'+1Co(9) = 0.201 *
ArAr'Co = 0.175
KL comparison tests
1−2 : p−val = 0.286
1−3 : p−val = 0.177
2−3 : p−val = 0.416

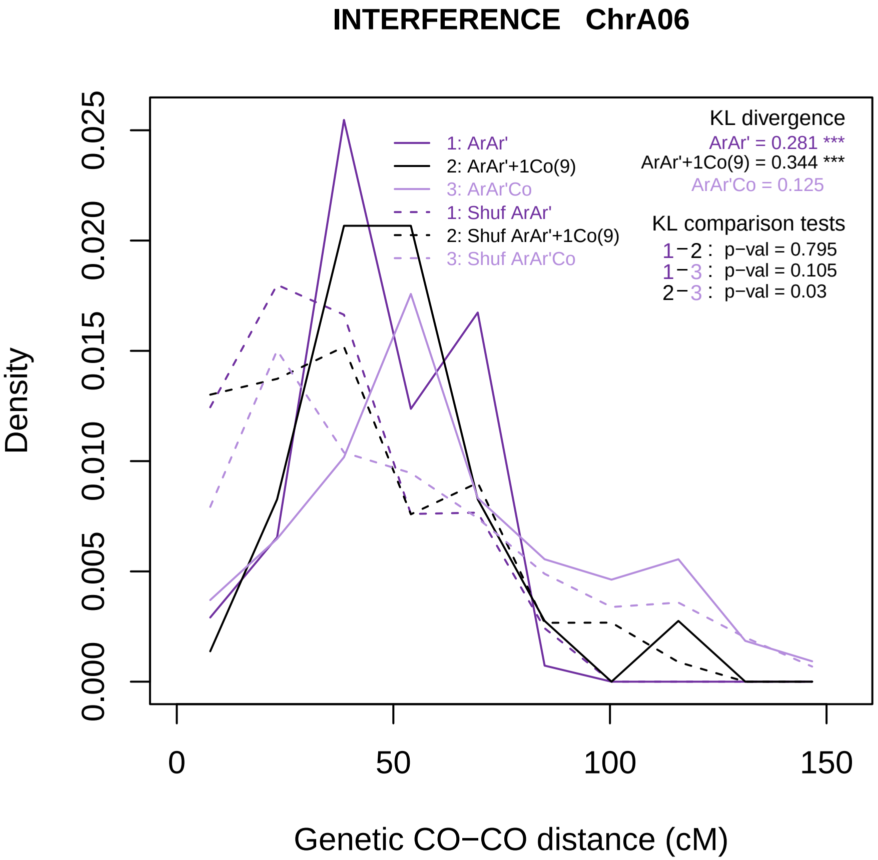

INTERFERENCE ChrA06
Density
Genetic CO−CO distance (cM)
0.000
0.005
0.010
0.015
0.020
0.025
0
50
100
150
1: ArAr'
2: ArAr'+1Co(9)
3: ArAr'Co
1: Shuf ArAr'
2: Shuf ArAr'+1Co(9)
3: Shuf ArAr'Co
KL divergence
ArAr' = 0.281 ***
ArAr'+1Co(9) = 0.344 ***
ArAr'Co = 0.125
KL comparison tests
1−2 : p−val = 0.795
1−3 : p−val = 0.105
2−3 : p−val = 0.03

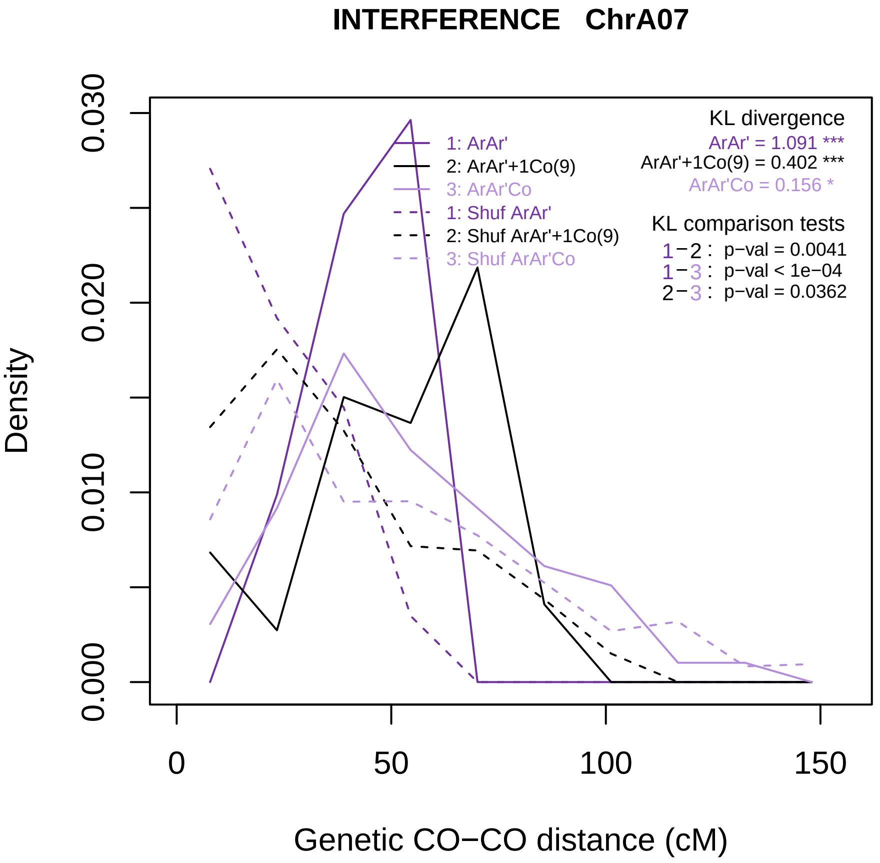
INTERFERENCE ChrA07
Density
0.000
0.010
0.020
0.030
0
50
100
150
Genetic CO−CO distance (cM)
1: ArAr'
2: ArAr'+1Co(9)
3: ArAr'Co
1: Shuf ArAr'
2: Shuf ArAr'+1Co(9)
3: Shuf ArAr'Co
KL divergence
ArAr' = 1.091 ***
ArAr'+1Co(9) = 0.402 ***
ArAr'Co = 0.156 *
KL comparison tests
1−2 : p−val = 0.0041
1−3 : p−val < 1e−04
2−3 : p−val = 0.0362

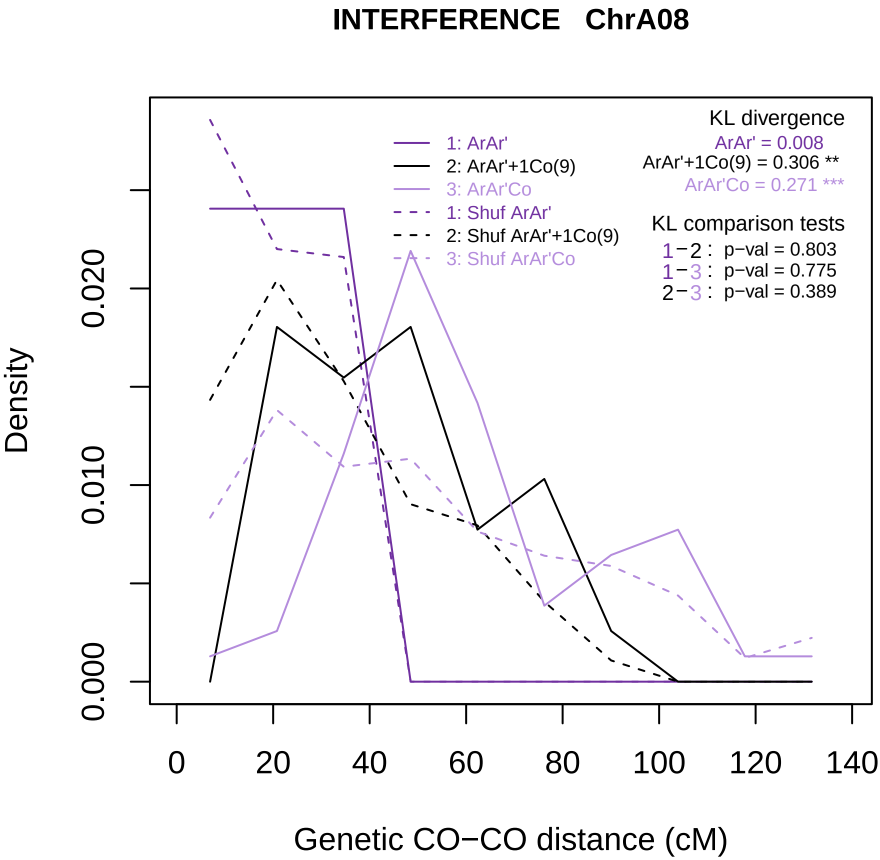

INTERFERENCE ChrA08
1: ArAr'
2: ArAr'+1Co(9)
3: ArAr'Co
1: Shuf ArAr'
2: Shuf ArAr'+1Co(9)
3: Shuf ArAr'Co
KL divergence
ArAr' = 0.008
ArAr'+1Co(9) = 0.306 **
ArAr'Co = 0.271 ***
KL comparison tests
1−2 : p−val = 0.803
1−3 : p−val = 0.775
2−3 : p−val = 0.389
Density
0.000
0.010
0.020
0
20
40
60
80
100
120
140
Genetic CO−CO distance (cM)

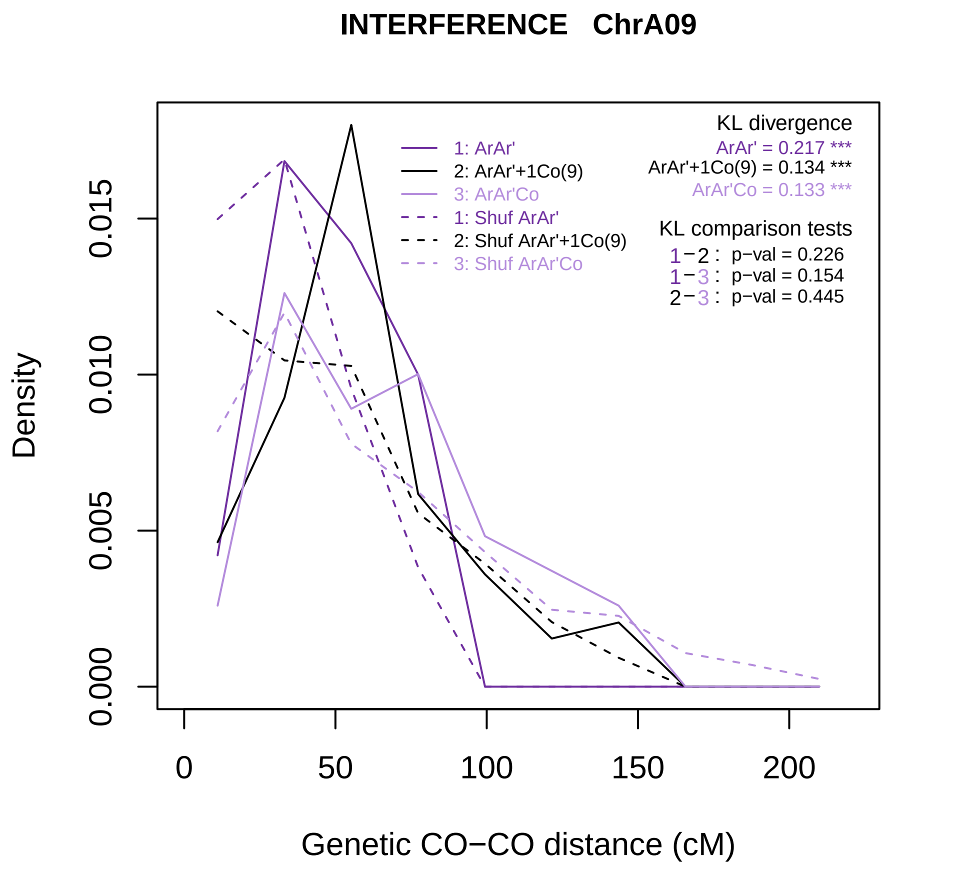

INTERFERENCE ChrA09
KL divergence
ArAr' = 0.217 ***
ArAr'+1Co(9) = 0.134 ***
ArAr'Co = 0.133 ***
1: ArAr'
2: ArAr'+1Co(9)
3: ArAr'Co
1: Shuf ArAr'
2: Shuf ArAr'+1Co(9)
3: Shuf ArAr'Co
KL comparison tests
1−2 : p−val = 0.226
1−3 : p−val = 0.154
2−3 : p−val = 0.445
Density
0.000
0.005
0.010
0.015
0
50
100
150
200
Genetic CO−CO distance (cM)

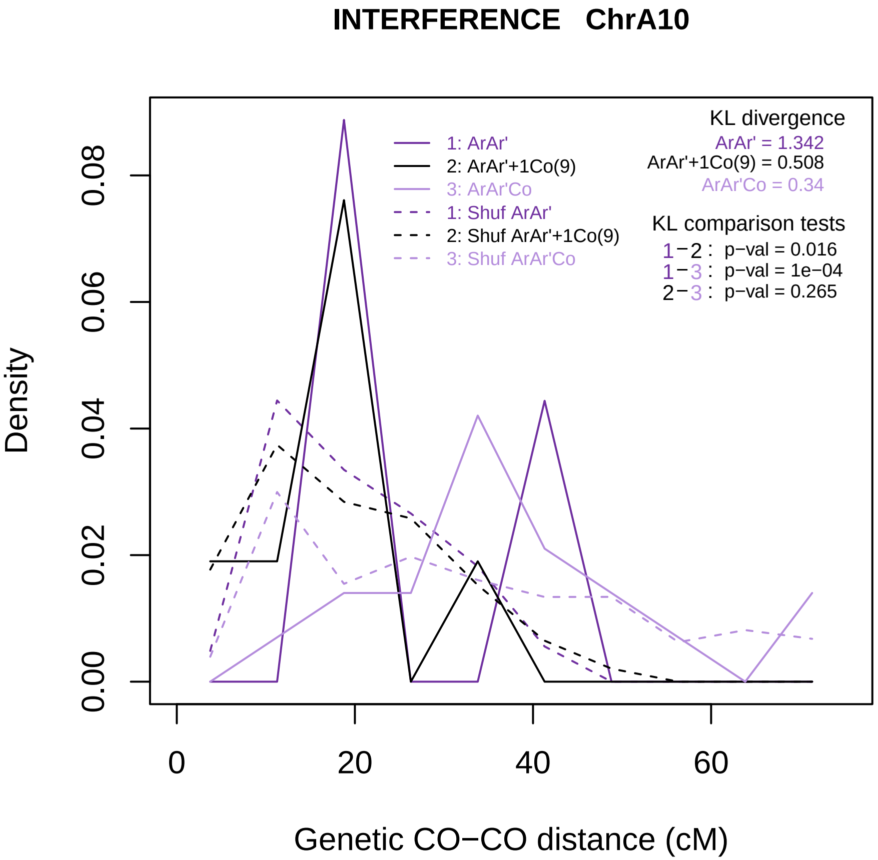

INTERFERENCE ChrA10
Density
0.00
0.02
0.04
0.06
0.08
0
20
40
60
Genetic CO−CO distance (cM)
1: ArAr'
2: ArAr'+1Co(9)
3: ArAr'Co
1: Shuf ArAr'
2: Shuf ArAr'+1Co(9)
3: Shuf ArAr'Co
KL divergence
ArAr' = 1.342
ArAr'+1Co(9) = 0.508
ArAr'Co = 0.34
KL comparison tests
1−2 : p−val = 0.016
1−3 : p−val = 1e−04
2−3 : p−val = 0.265

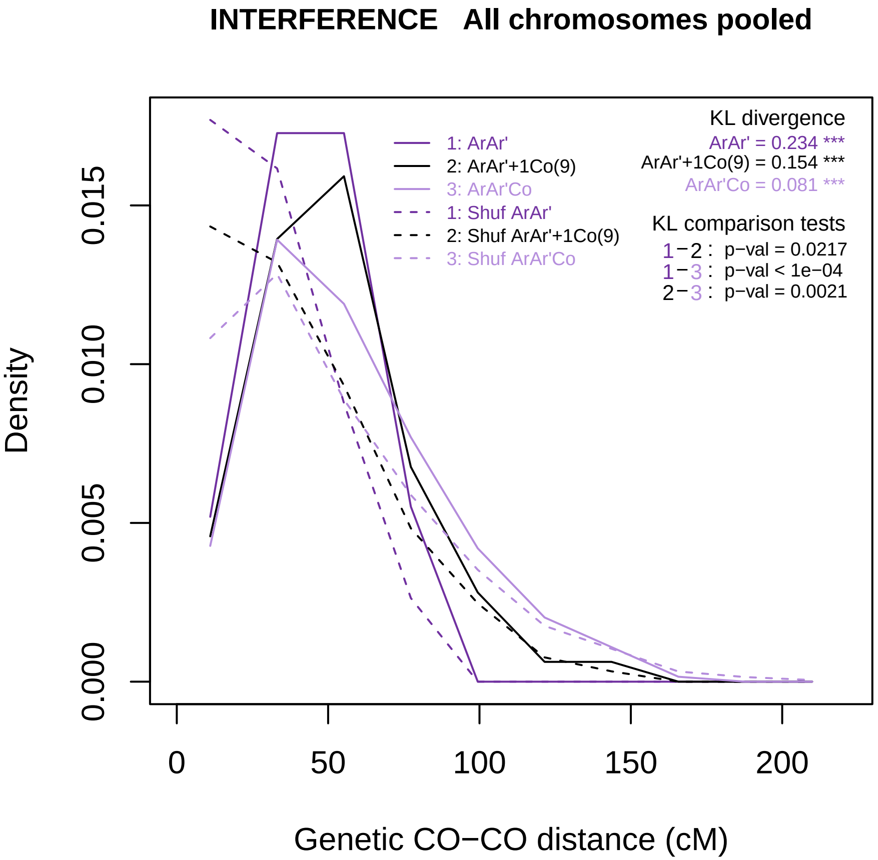
INTERFERENCE All chromosomes pooled
Density
0.000
0.005
0.010
0.015
0
50
100
150
200
Genetic CO−CO distance (cM)
1: ArAr'
2: ArAr'+1Co(9)
3: ArAr'Co
1: Shuf ArAr'
2: Shuf ArAr'+1Co(9)
3: Shuf ArAr'Co
KL divergence
ArAr' = 0.234 ***
ArAr'+1Co(9) = 0.154 ***
ArAr'Co = 0.081 ***
KL comparison tests
1−2 : p−val = 0.0217
1−3 : p−val < 1e−04
2−3 : p−val = 0.0021

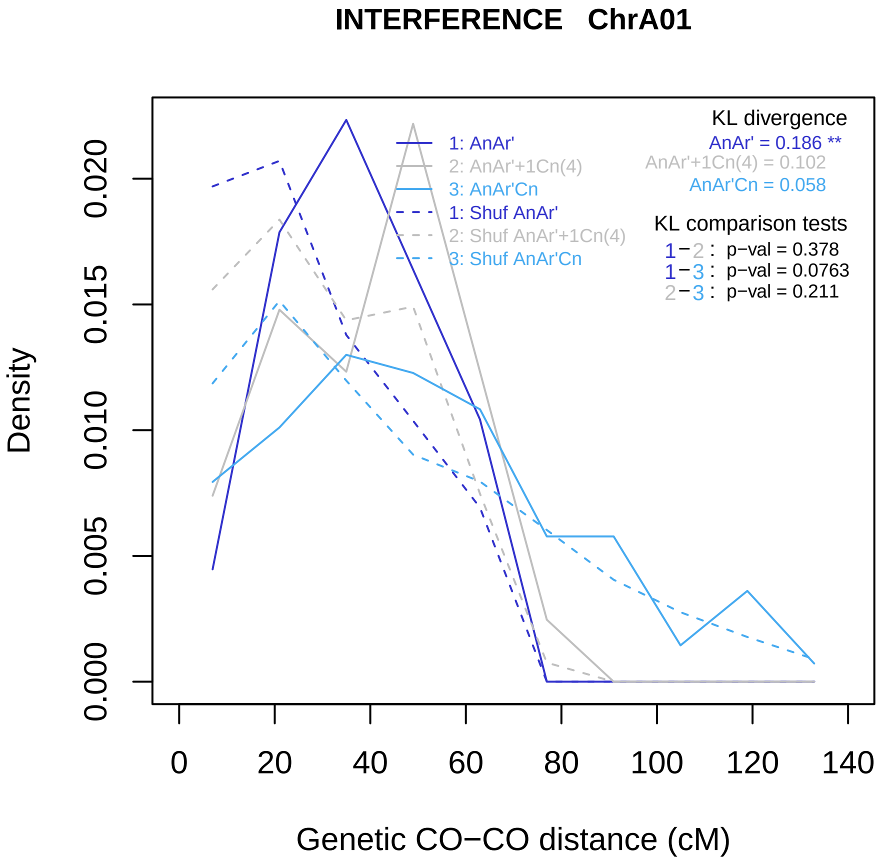

INTERFERENCE ChrA01
Density
0.000
0.005
0.010
0.015
0.020
0
20
40
60
80
100
120
140
Genetic CO−CO distance (cM)
1: AnAr'
2: AnAr'+1Cn(4)
3: AnAr'Cn
1: Shuf AnAr'
2: Shuf AnAr'+1Cn(4)
3: Shuf AnAr'Cn
KL divergence
AnAr' = 0.186 **
AnAr'+1Cn(4) = 0.102
AnAr'Cn = 0.058
KL comparison tests
1−2 : p−val = 0.378
1−3 : p−val = 0.0763
2−3 : p−val = 0.211

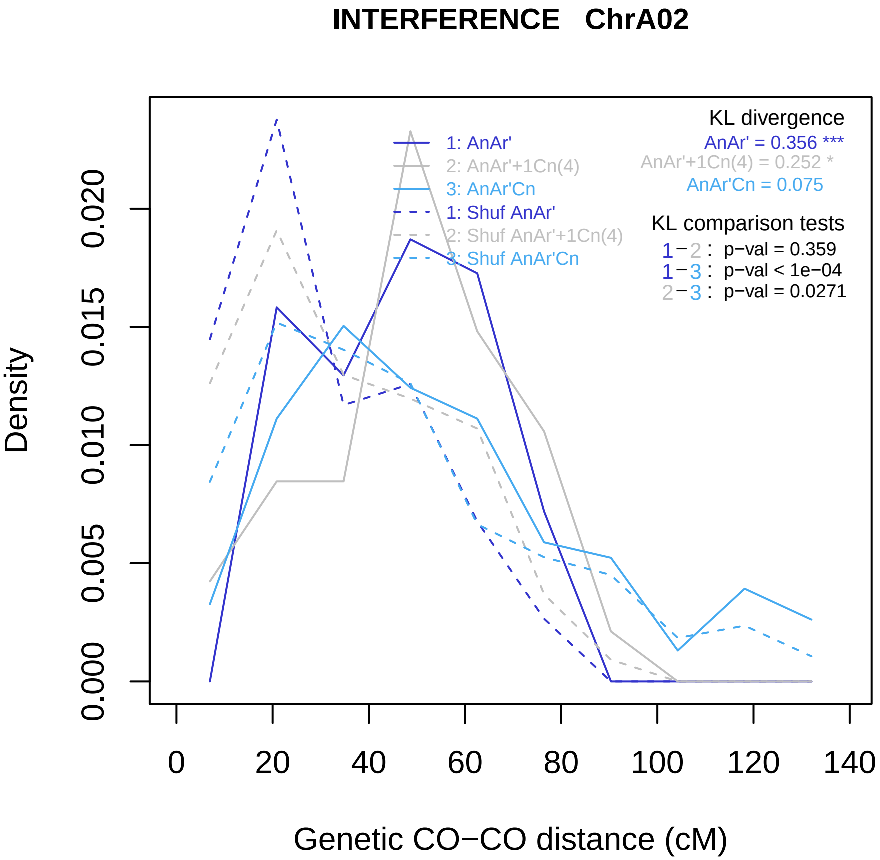

INTERFERENCE ChrA02
KL divergence
AnAr' = 0.356 ***
AnAr'+1Cn(4) = 0.252 *
AnAr'Cn = 0.075
1: AnAr'
2: AnAr'+1Cn(4)
3: AnAr'Cn
1: Shuf AnAr'
2: Shuf AnAr'+1Cn(4)
3: Shuf AnAr'Cn
KL comparison tests
1−2 : p−val = 0.359
1−3 : p−val < 1e−04
2−3 : p−val = 0.0271
Density
0.000
0.005
0.010
0.015
0.020
0
20
40
60
80
100
120
140
Genetic CO−CO distance (cM)

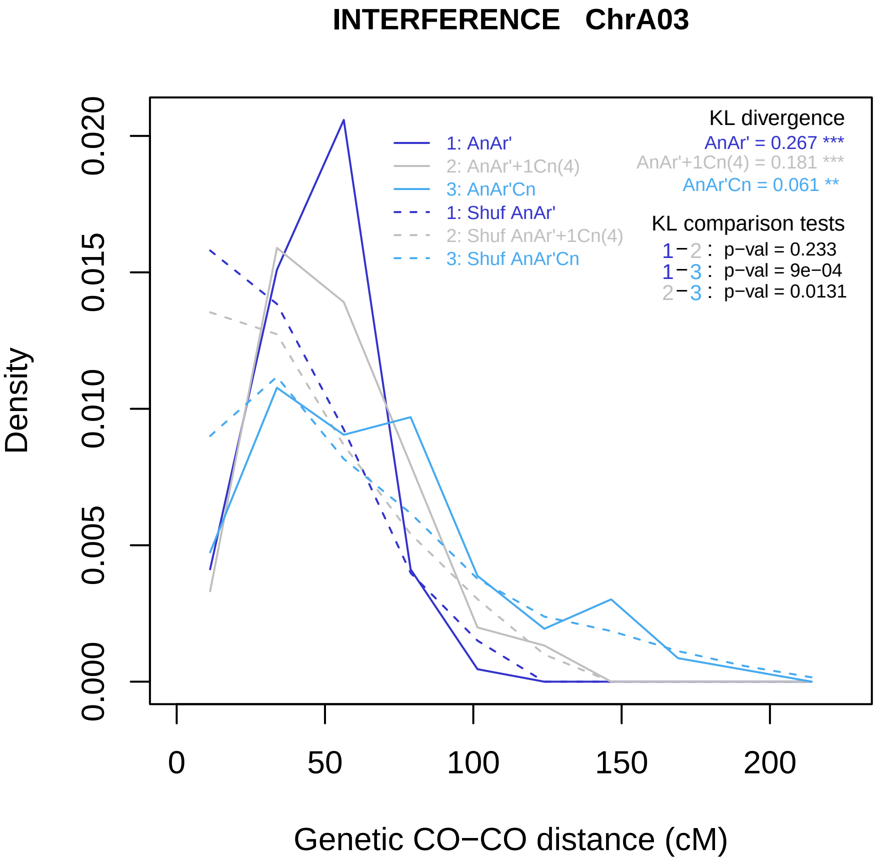

INTERFERENCE ChrA03
KL divergence
AnAr' = 0.267 ***
AnAr'+1Cn(4) = 0.181 ***
AnAr'Cn = 0.061 **
1: AnAr'
2: AnAr'+1Cn(4)
3: AnAr'Cn
1: Shuf AnAr'
2: Shuf AnAr'+1Cn(4)
3: Shuf AnAr'Cn
KL comparison tests
1−2 : p−val = 0.233
1−3 : p−val = 9e−04
2−3 : p−val = 0.0131
Density
0.000
0.005
0.010
0.015
0.020
0
50
100
150
200
Genetic CO−CO distance (cM)

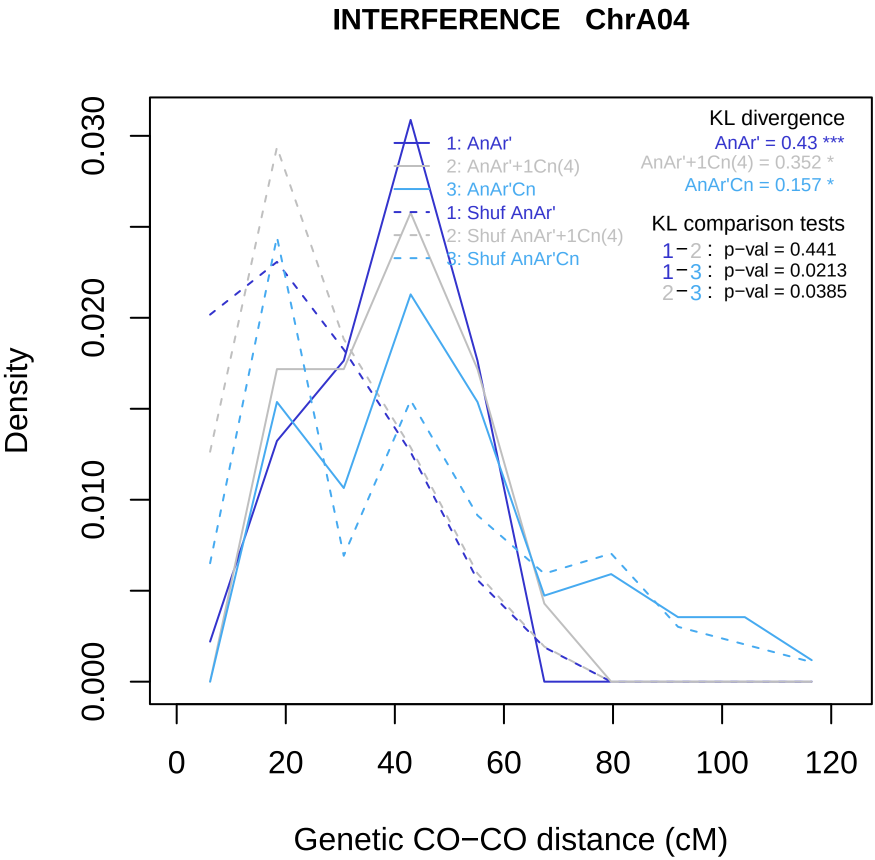
INTERFERENCE ChrA04
Density
0.000
0.010
0.020
0.030
0
20
40
60
80
100
120
Genetic CO−CO distance (cM)
1: AnAr'
2: AnAr'+1Cn(4)
3: AnAr'Cn
1: Shuf AnAr'
2: Shuf AnAr'+1Cn(4)
3: Shuf AnAr'Cn
KL divergence
AnAr' = 0.43 ***
AnAr'+1Cn(4) = 0.352 *
AnAr'Cn = 0.157 *
KL comparison tests
1−2 : p−val = 0.441
1−3 : p−val = 0.0213
2−3 : p−val = 0.0385

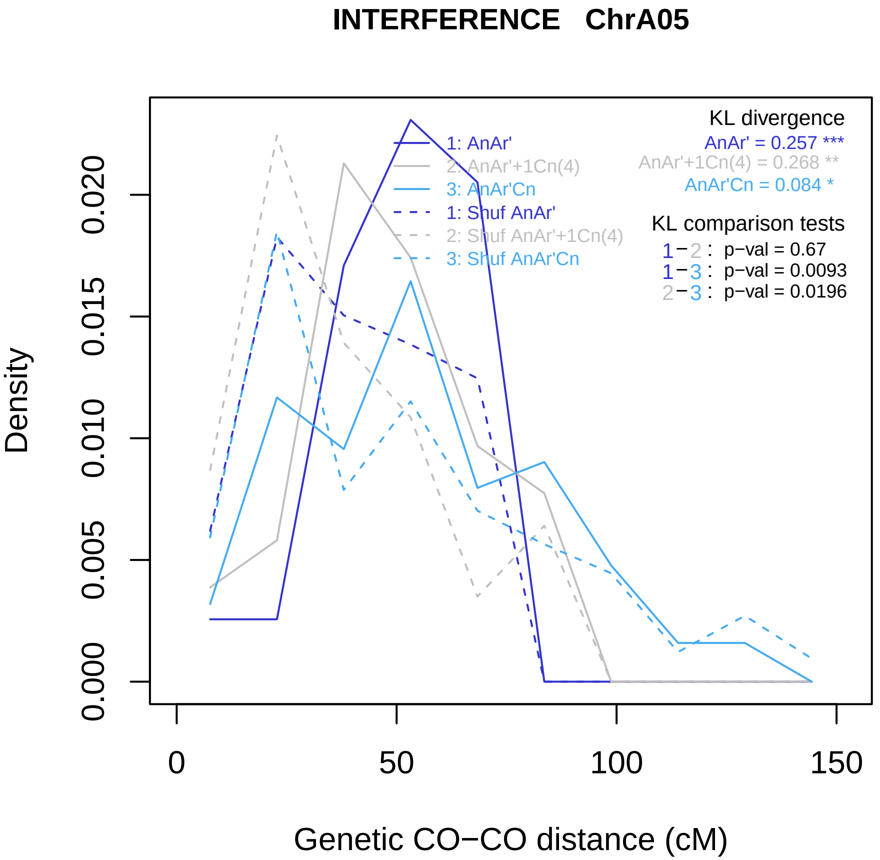
INTERFERENCE   ChrA05
Density
Genetic CO−CO distance (cM)
0.000
0.005
0.010
0.015
0.020
0
50
100
150
1: AnAr'
2: AnAr'+1Cn(4)
3: AnAr'Cn
1: Shuf AnAr'
2: Shuf AnAr'+1Cn(4)
3: Shuf AnAr'Cn
KL divergence
AnAr' = 0.257 ***
AnAr'+1Cn(4) = 0.268 **
AnAr'Cn = 0.084 *
KL comparison tests
1−2 : p−val = 0.67
1−3 : p−val = 0.0093
2−3 : p−val = 0.0196

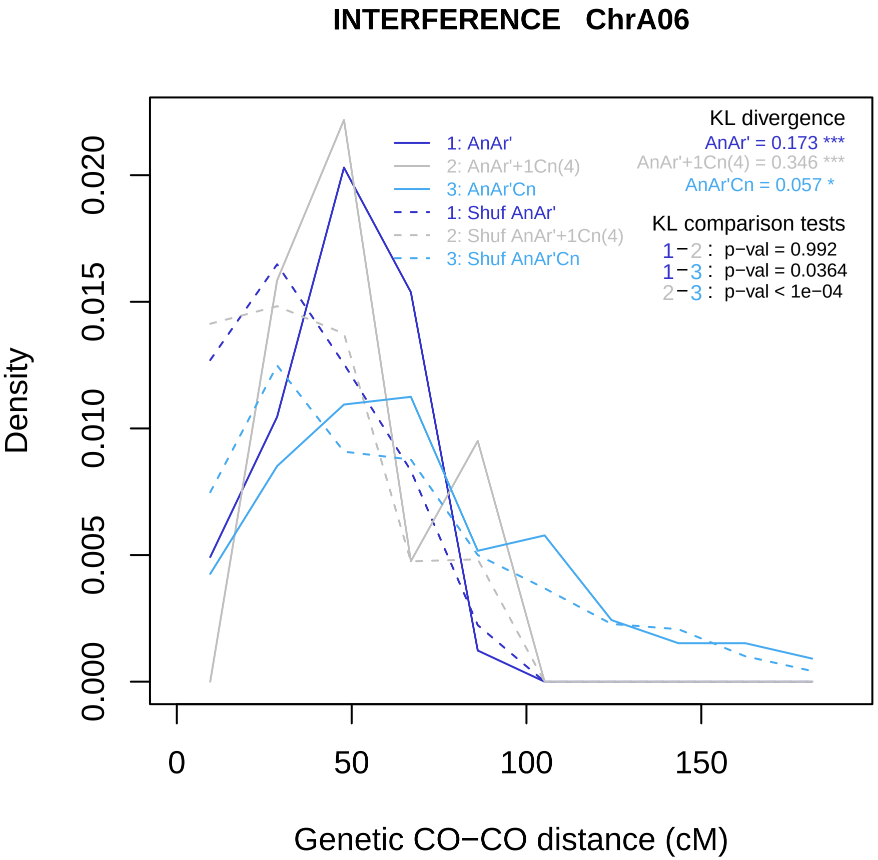
INTERFERENCE ChrA06
Density
0.000
0.005
0.010
0.015
0.020
0
50
100
150
Genetic CO−CO distance (cM)
1: AnAr'
2: AnAr'+1Cn(4)
3: AnAr'Cn
1: Shuf AnAr'
2: Shuf AnAr'+1Cn(4)
3: Shuf AnAr'Cn
KL divergence
AnAr' = 0.173 ***
AnAr'+1Cn(4) = 0.346 ***
AnAr'Cn = 0.057 *
KL comparison tests
1−2 : p−val = 0.992
1−3 : p−val = 0.0364
2−3 : p−val < 1e−04

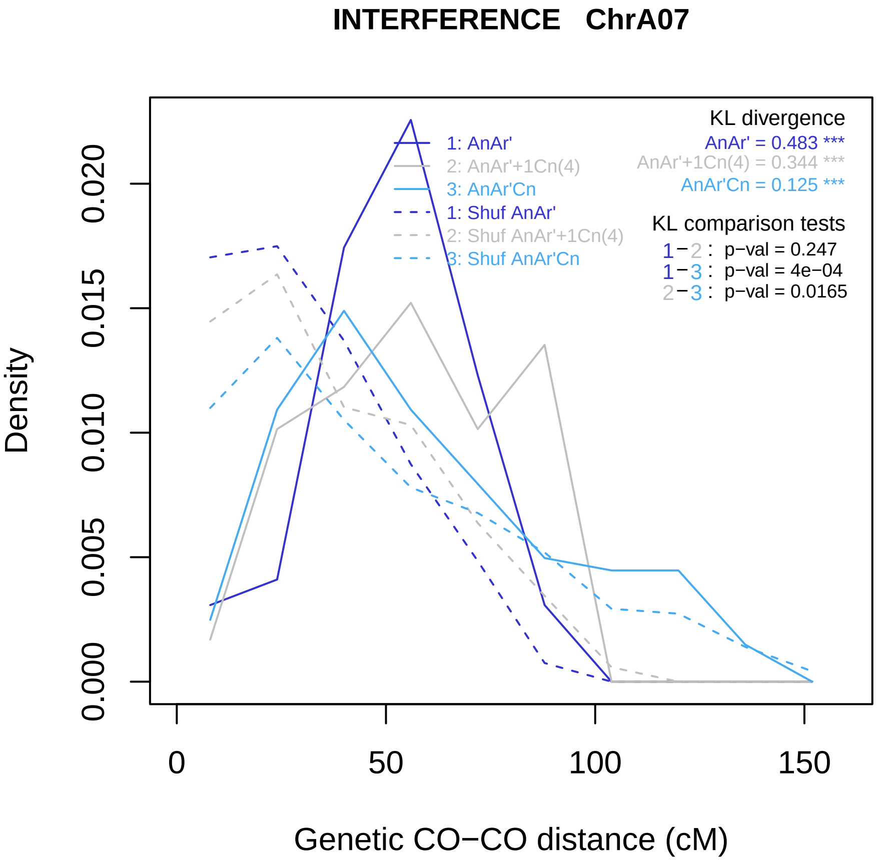

INTERFERENCE ChrA07
KL divergence
AnAr' = 0.483 ***
AnAr'+1Cn(4) = 0.344 ***
AnAr'Cn = 0.125 ***
1: AnAr'
2: AnAr'+1Cn(4)
3: AnAr'Cn
1: Shuf AnAr'
2: Shuf AnAr'+1Cn(4)
3: Shuf AnAr'Cn
KL comparison tests
1−2 : p−val = 0.247
1−3 : p−val = 4e−04
2−3 : p−val = 0.0165
Density
0.000
0.005
0.010
0.015
0.020
0
50
100
150
Genetic CO−CO distance (cM)

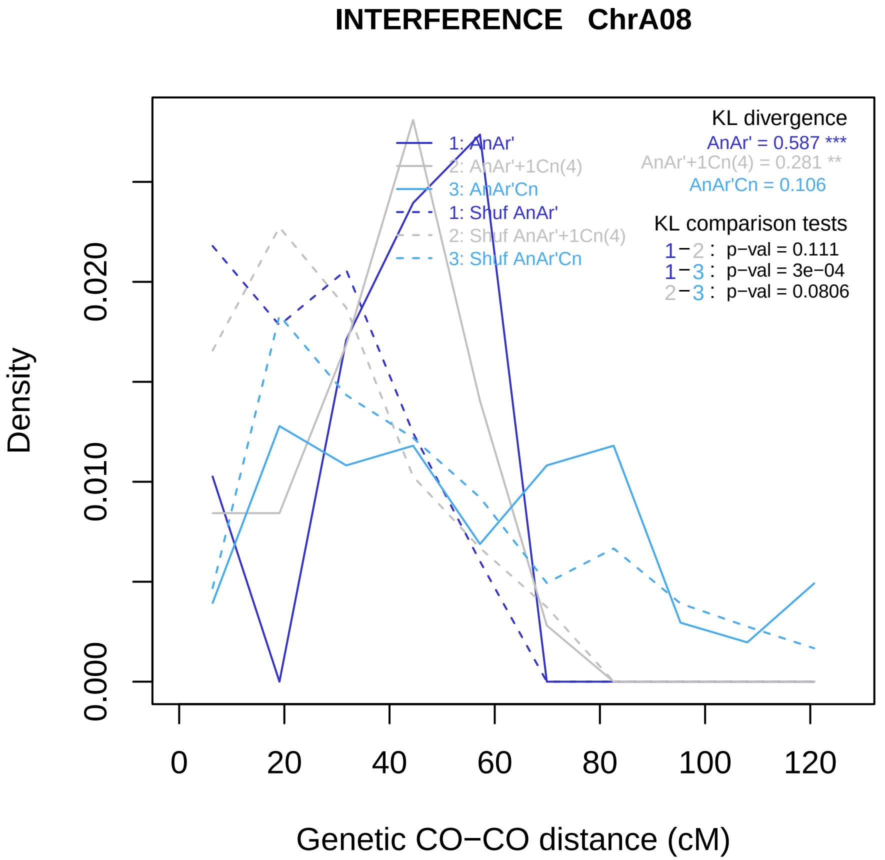
INTERFERENCE ChrA08
KL divergence
AnAr' = 0.587 ***
AnAr'+1Cn(4) = 0.281 **
AnAr'Cn = 0.106
KL comparison tests
1−2 : p−val = 0.111
1−3 : p−val = 3e−04
2−3 : p−val = 0.0806
1: AnAr'
2: AnAr'+1Cn(4)
3: AnAr'Cn
1: Shuf AnAr'
2: Shuf AnAr'+1Cn(4)
3: Shuf AnAr'Cn
Density
0.000
0.010
0.020
0
20
40
60
80
100
120
Genetic CO−CO distance (cM)

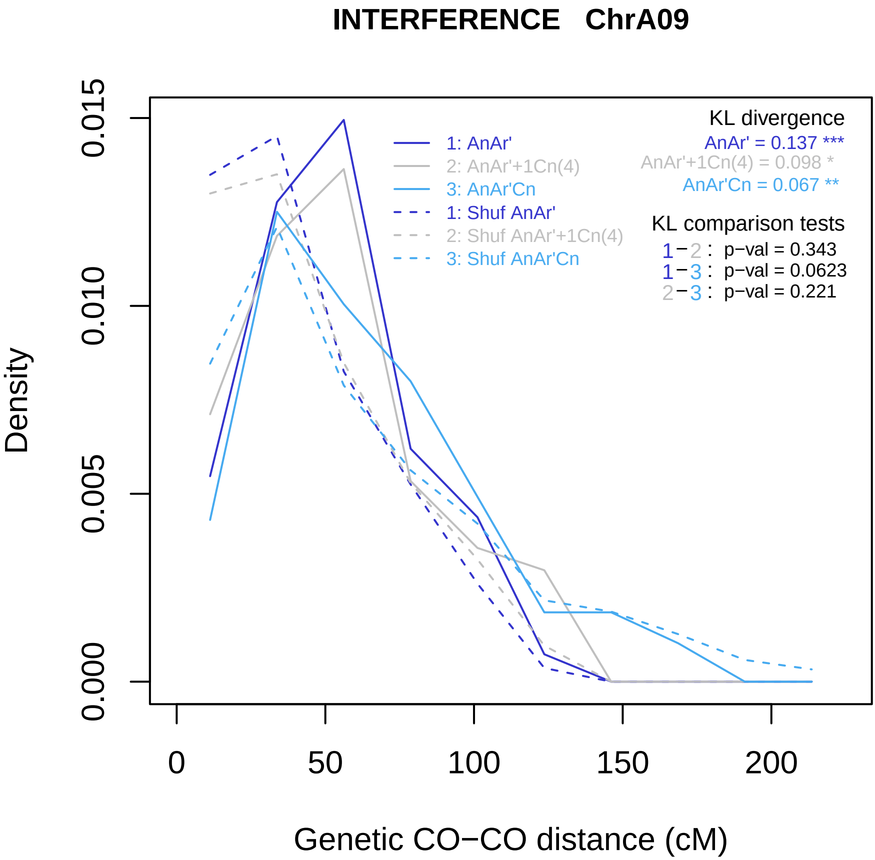
INTERFERENCE ChrA09
Density
0.000
0.005
0.010
0.015
0
50
100
150
200
Genetic CO−CO distance (cM)
1: AnAr'
2: AnAr'+1Cn(4)
3: AnAr'Cn
1: Shuf AnAr'
2: Shuf AnAr'+1Cn(4)
3: Shuf AnAr'Cn
KL divergence
AnAr' = 0.137 ***
AnAr'+1Cn(4) = 0.098 *
AnAr'Cn = 0.067 **
KL comparison tests
1−2 : p−val = 0.343
1−3 : p−val = 0.0623
2−3 : p−val = 0.221

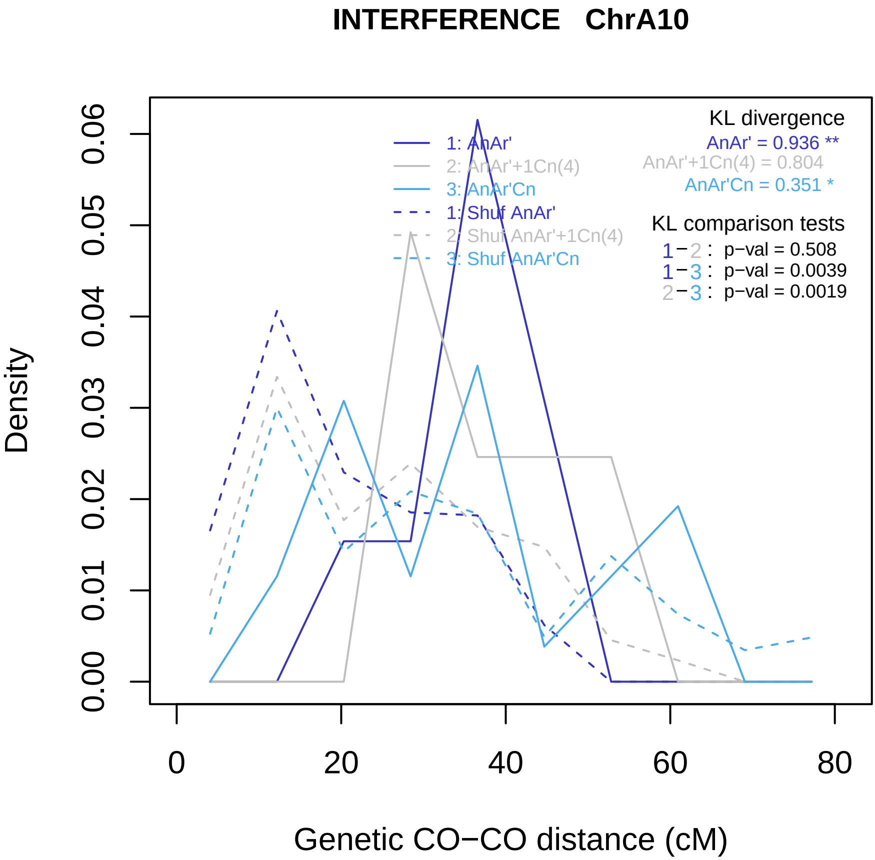

INTERFERENCE ChrA10
Density
Genetic CO−CO distance (cM)
0.00
0.01
0.02
0.03
0.04
0.05
0.06
0
20
40
60
80
1: AnAr'
2: AnAr'+1Cn(4)
3: AnAr'Cn
1: Shuf AnAr'
2: Shuf AnAr'+1Cn(4)
3: Shuf AnAr'Cn
KL divergence
AnAr' = 0.936 **
AnAr'+1Cn(4) = 0.804
AnAr'Cn = 0.351 *
KL comparison tests
1−2 : p−val = 0.508
1−3 : p−val = 0.0039
2−3 : p−val = 0.0019

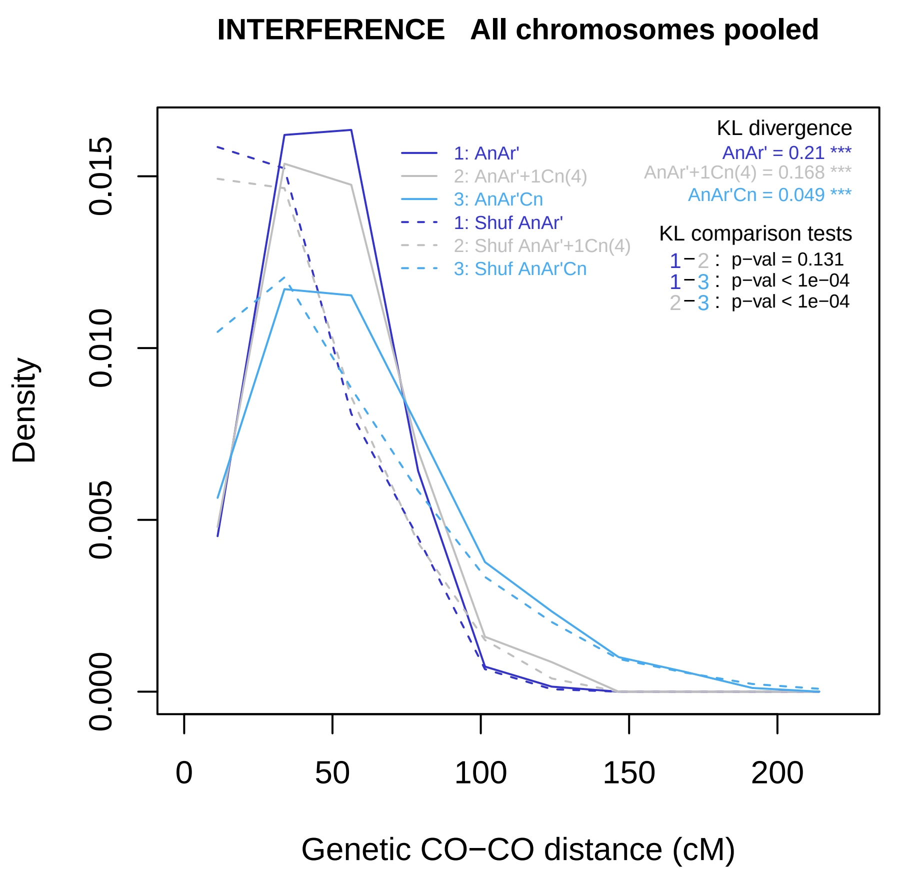

INTERFERENCE All chromosomes pooled
Density
0.000
0.005
0.010
0.015
0
50
100
150
200
Genetic CO−CO distance (cM)
1: AnAr'
2: AnAr'+1Cn(4)
3: AnAr'Cn
1: Shuf AnAr'
2: Shuf AnAr'+1Cn(4)
3: Shuf AnAr'Cn
KL divergence
AnAr' = 0.21 ***
AnAr'+1Cn(4) = 0.168 ***
AnAr'Cn = 0.049 ***
KL comparison tests
1−2 : p−val = 0.131
1−3 : p−val < 1e−04
2−3 : p−val < 1e−04

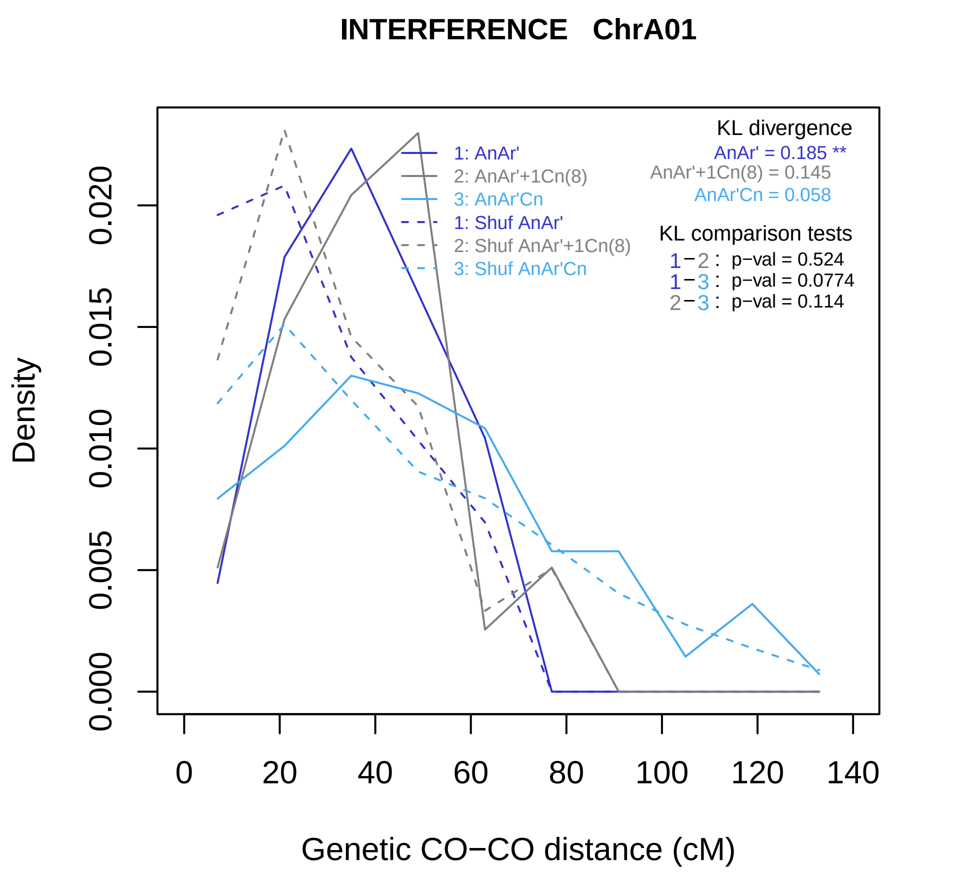
INTERFERENCE ChrA01
KL divergence
AnAr' = 0.185 **
AnAr'+1Cn(8) = 0.145
AnAr'Cn = 0.058
1: AnAr'
2: AnAr'+1Cn(8)
3: AnAr'Cn
1: Shuf AnAr'
2: Shuf AnAr'+1Cn(8)
3: Shuf AnAr'Cn
KL comparison tests
1−2 : p−val = 0.524
1−3 : p−val = 0.0774
2−3 : p−val = 0.114
Density
0.000
0.005
0.010
0.015
0.020
0
20
40
60
80
100
120
140
Genetic CO−CO distance (cM)

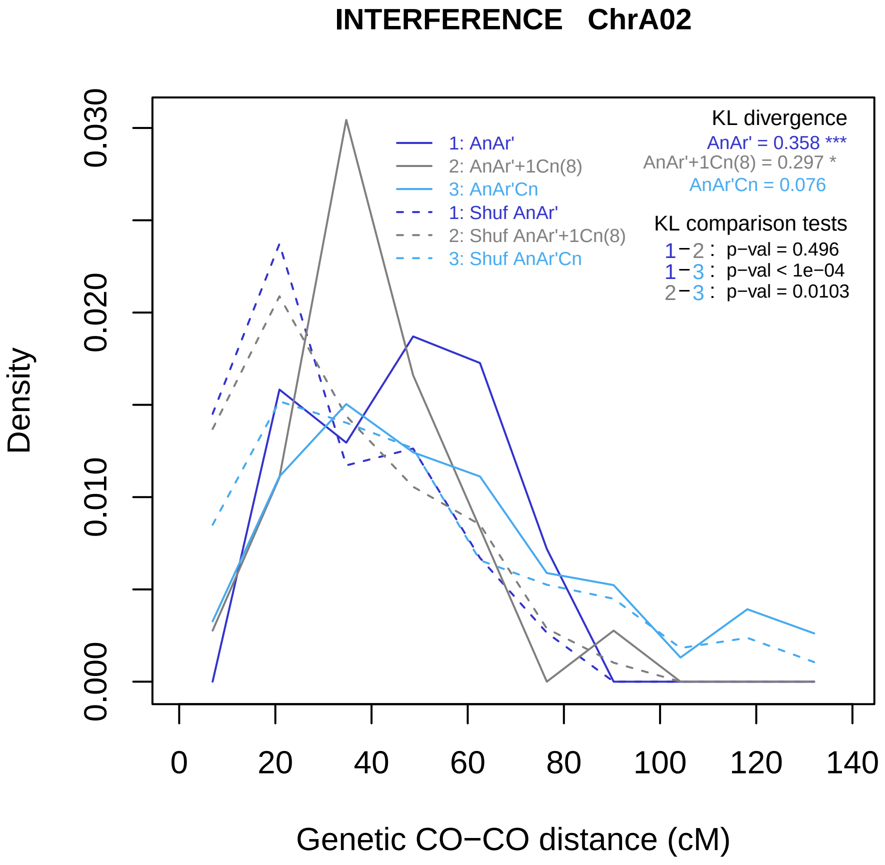
INTERFERENCE ChrA02
Density
0.000
0.010
0.020
0.030
0
20
40
60
80
100
120
140
Genetic CO−CO distance (cM)
1: AnAr'
2: AnAr'+1Cn(8)
3: AnAr'Cn
1: Shuf AnAr'
2: Shuf AnAr'+1Cn(8)
3: Shuf AnAr'Cn
KL divergence
AnAr' = 0.358 ***
AnAr'+1Cn(8) = 0.297 *
AnAr'Cn = 0.076
KL comparison tests
1−2 : p−val = 0.496
1−3 : p−val < 1e−04
2−3 : p−val = 0.0103

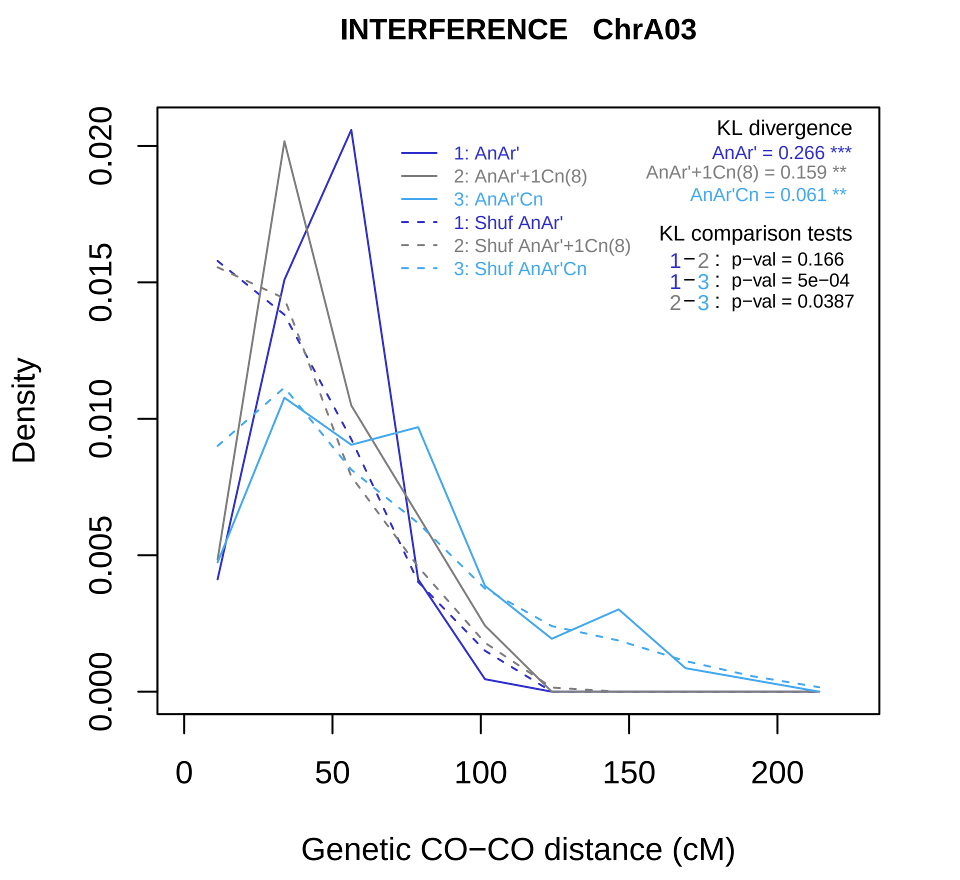

INTERFERENCE ChrA03
KL divergence
AnAr' = 0.266 ***
AnAr'+1Cn(8) = 0.159 **
AnAr'Cn = 0.061 **
1: AnAr'
2: AnAr'+1Cn(8)
3: AnAr'Cn
1: Shuf AnAr'
2: Shuf AnAr'+1Cn(8)
3: Shuf AnAr'Cn
KL comparison tests
1−2 : p−val = 0.166
1−3 : p−val = 5e−04
2−3 : p−val = 0.0387
Density
0.000
0.005
0.010
0.015
0.020
0
50
100
150
200
Genetic CO−CO distance (cM)

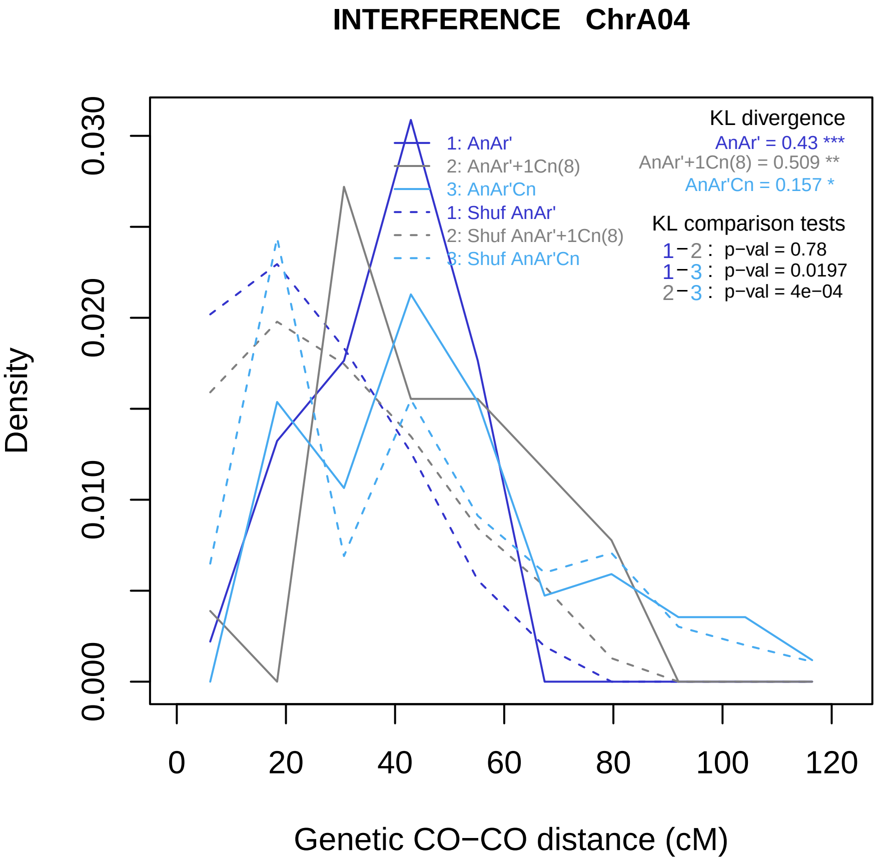

INTERFERENCE ChrA04
Density
0.000
0.010
0.020
0.030
0
20
40
60
80
100
120
Genetic CO−CO distance (cM)
1: AnAr'
2: AnAr'+1Cn(8)
3: AnAr'Cn
1: Shuf AnAr'
2: Shuf AnAr'+1Cn(8)
3: Shuf AnAr'Cn
KL divergence
AnAr' = 0.43 ***
AnAr'+1Cn(8) = 0.509 **
AnAr'Cn = 0.157 *
KL comparison tests
1−2 : p−val = 0.78
1−3 : p−val = 0.0197
2−3 : p−val = 4e−04

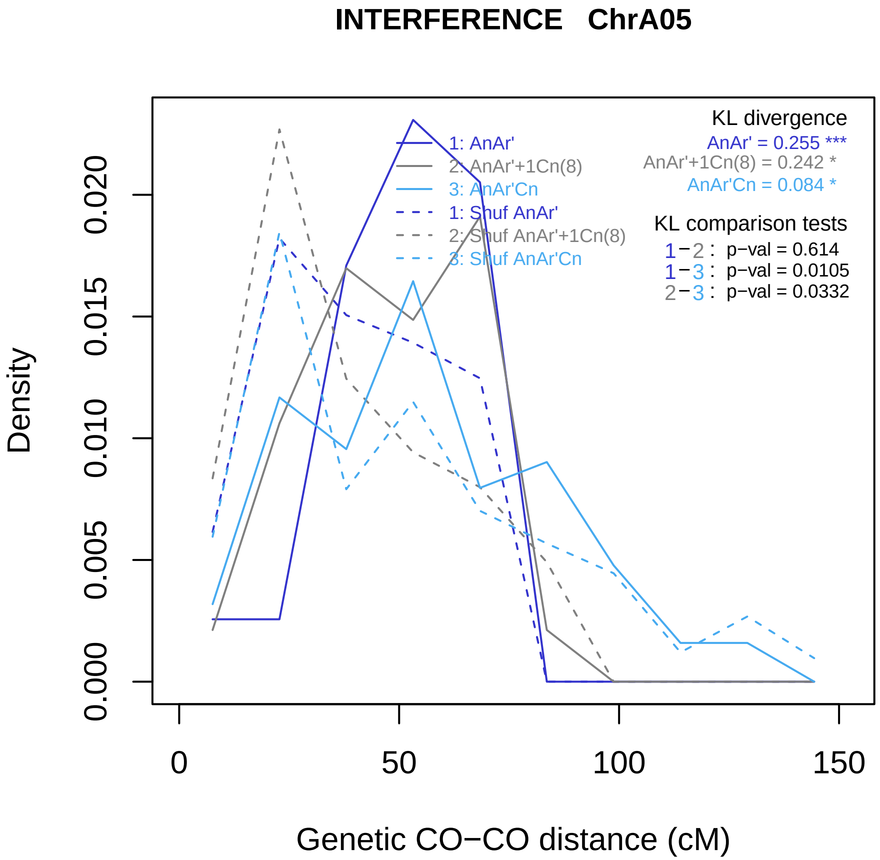

INTERFERENCE ChrA05
Density
0.000
0.005
0.010
0.015
0.020
Genetic CO−CO distance (cM)
0
50
100
150
1: AnAr'
2: AnAr'+1Cn(8)
3: AnAr'Cn
1: Shuf AnAr'
2: Shuf AnAr'+1Cn(8)
3: Shuf AnAr'Cn
KL divergence
AnAr' = 0.255 ***
AnAr'+1Cn(8) = 0.242 *
AnAr'Cn = 0.084 *
KL comparison tests
1−2 : p−val = 0.614
1−3 : p−val = 0.0105
2−3 : p−val = 0.0332

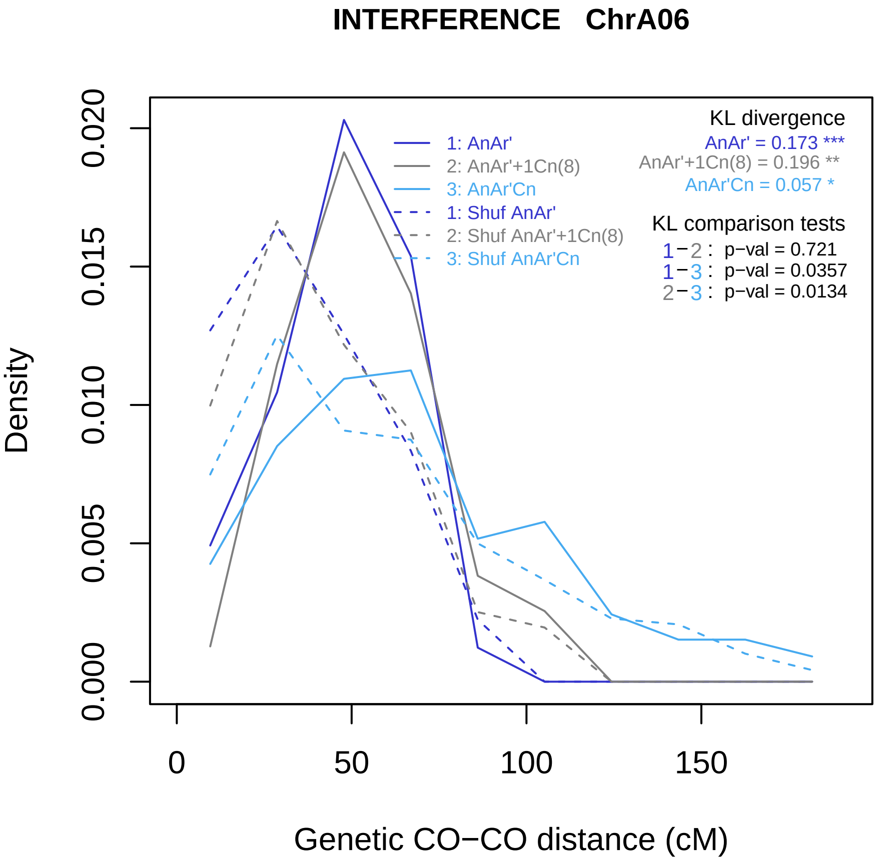
INTERFERENCE ChrA06
Density
0.000
0.005
0.010
0.015
0.020
0
50
100
150
Genetic CO−CO distance (cM)
1: AnAr'
2: AnAr'+1Cn(8)
3: AnAr'Cn
1: Shuf AnAr'
2: Shuf AnAr'+1Cn(8)
3: Shuf AnAr'Cn
KL divergence
AnAr' = 0.173 ***
AnAr'+1Cn(8) = 0.196 **
AnAr'Cn = 0.057 *
KL comparison tests
1−2 : p−val = 0.721
1−3 : p−val = 0.0357
2−3 : p−val = 0.0134

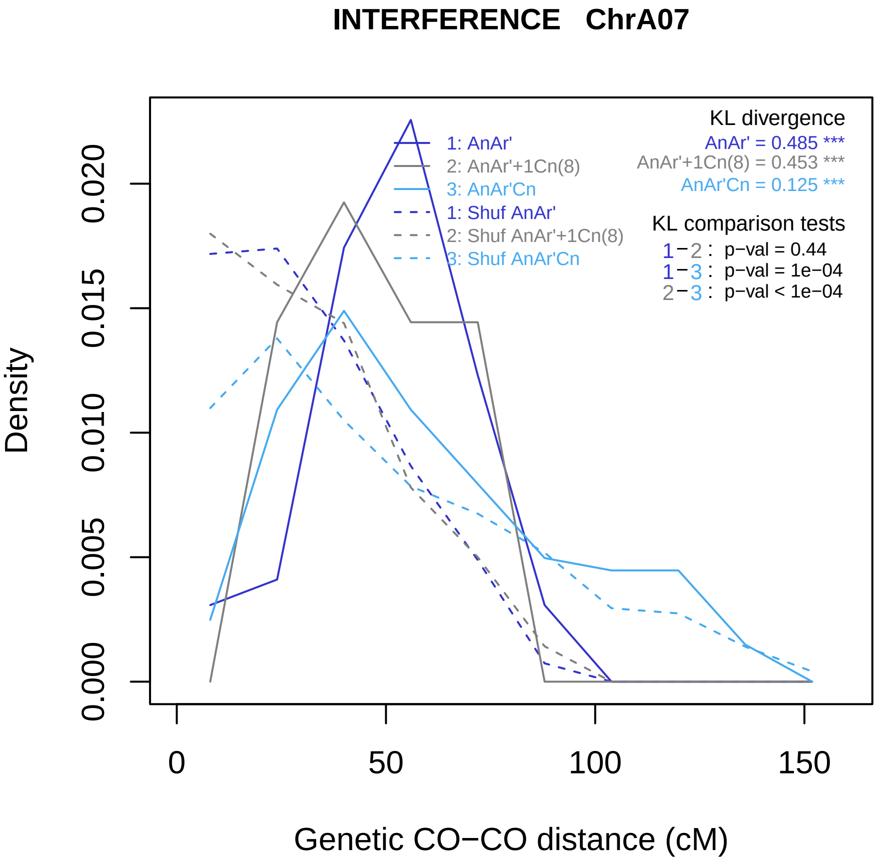
INTERFERENCE ChrA07
KL divergence
AnAr' = 0.485 ***
AnAr'+1Cn(8) = 0.453 ***
AnAr'Cn = 0.125 ***
KL comparison tests
1−2 : p−val = 0.44
1−3 : p−val = 1e−04
2−3 : p−val < 1e−04
1: AnAr'
2: AnAr'+1Cn(8)
3: AnAr'Cn
1: Shuf AnAr'
2: Shuf AnAr'+1Cn(8)
3: Shuf AnAr'Cn
Density
0.000
0.005
0.010
0.015
0.020
0
50
100
150
Genetic CO−CO distance (cM)

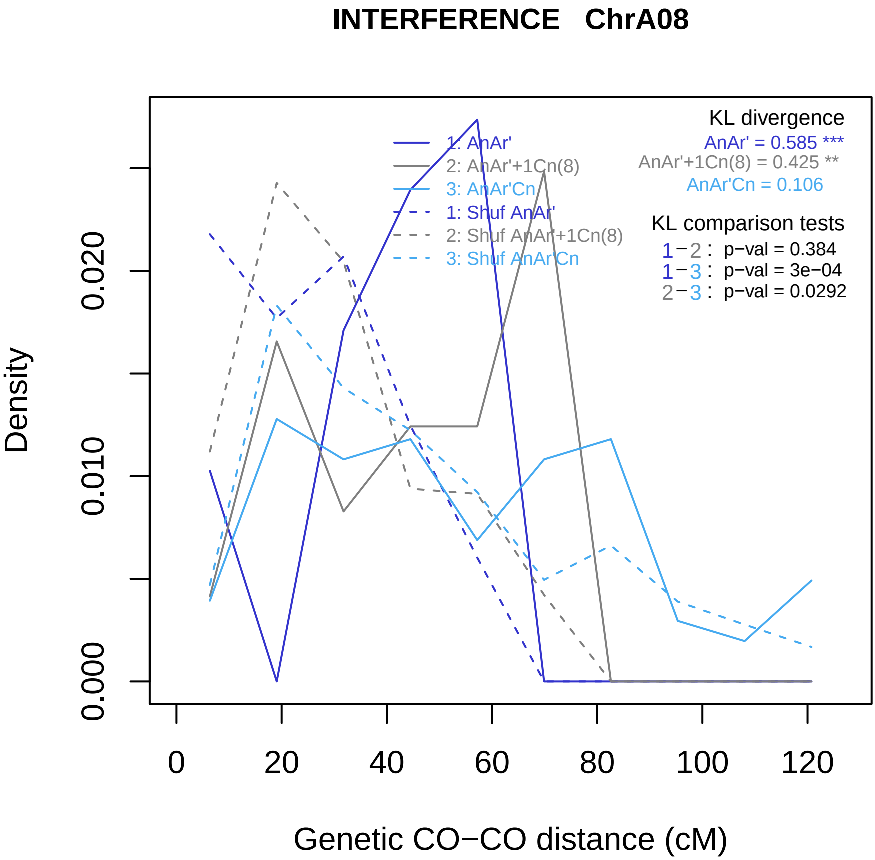
INTERFERENCE ChrA08
Density
0.000
0.010
0.020
Genetic CO−CO distance (cM)
0
20
40
60
80
100
120
1: AnAr'
2: AnAr'+1Cn(8)
3: AnAr'Cn
1: Shuf AnAr'
2: Shuf AnAr'+1Cn(8)
3: Shuf AnAr'Cn
KL divergence
AnAr' = 0.585 ***
AnAr'+1Cn(8) = 0.425 **
AnAr'Cn = 0.106
KL comparison tests
1−2 : p−val = 0.384
1−3 : p−val = 3e−04
2−3 : p−val = 0.0292

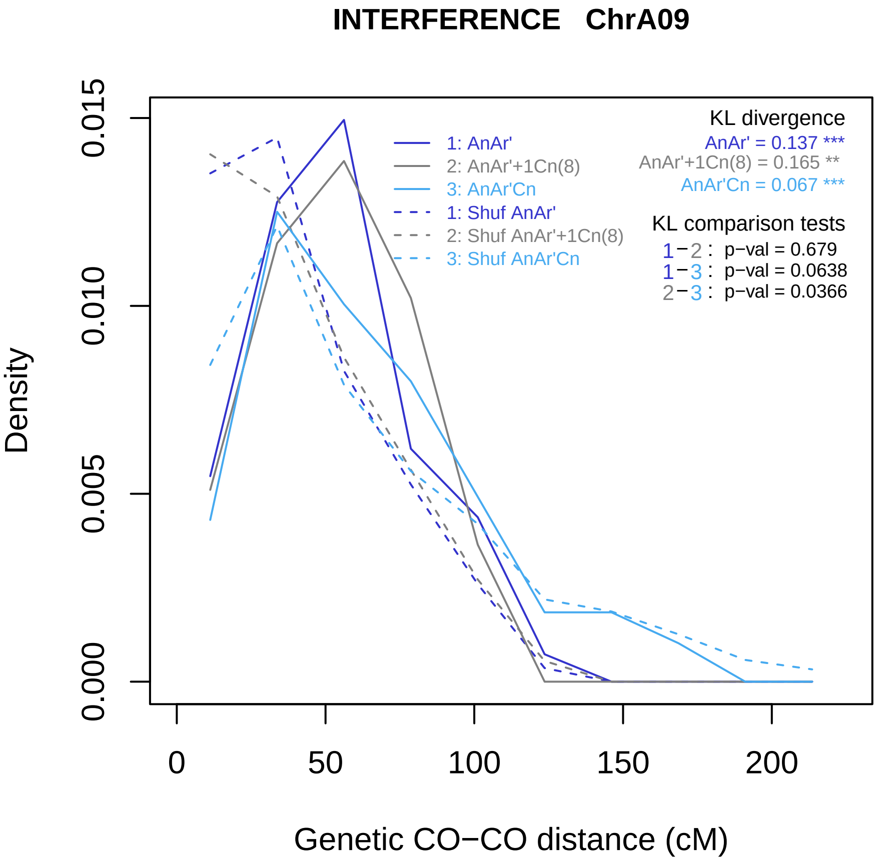
INTERFERENCE ChrA09
Density
0.000
0.005
0.010
0.015
0
50
100
150
200
Genetic CO−CO distance (cM)
1: AnAr'
2: AnAr'+1Cn(8)
3: AnAr'Cn
1: Shuf AnAr'
2: Shuf AnAr'+1Cn(8)
3: Shuf AnAr'Cn
KL divergence
AnAr' = 0.137 ***
AnAr'+1Cn(8) = 0.165 **
AnAr'Cn = 0.067 ***
KL comparison tests
1−2 : p−val = 0.679
1−3 : p−val = 0.0638
2−3 : p−val = 0.0366

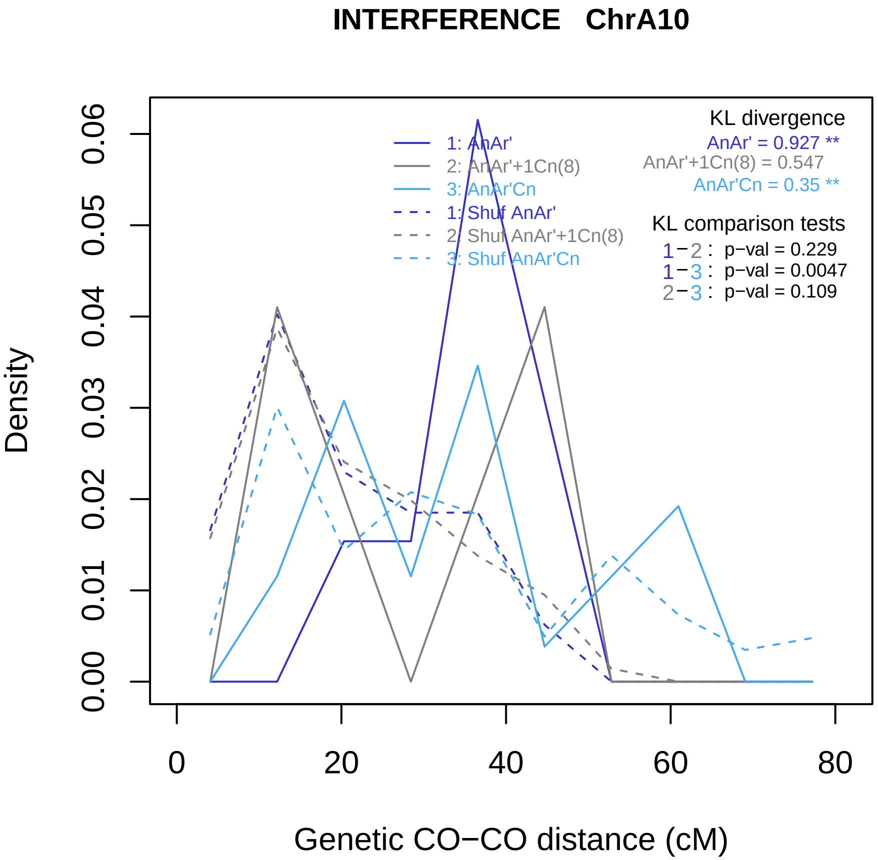

INTERFERENCE ChrA10
KL divergence
AnAr' = 0.927 **
AnAr'+1Cn(8) = 0.547
AnAr'Cn = 0.35 **
KL comparison tests
1−2 : p−val = 0.229
1−3 : p−val = 0.0047
2−3 : p−val = 0.109
1: AnAr'
2: AnAr'+1Cn(8)
3: AnAr'Cn
1: Shuf AnAr'
2: Shuf AnAr'+1Cn(8)
3: Shuf AnAr'Cn
Density
0.00
0.01
0.02
0.03
0.04
0.05
0.06
0
20
40
60
80
Genetic CO−CO distance (cM)

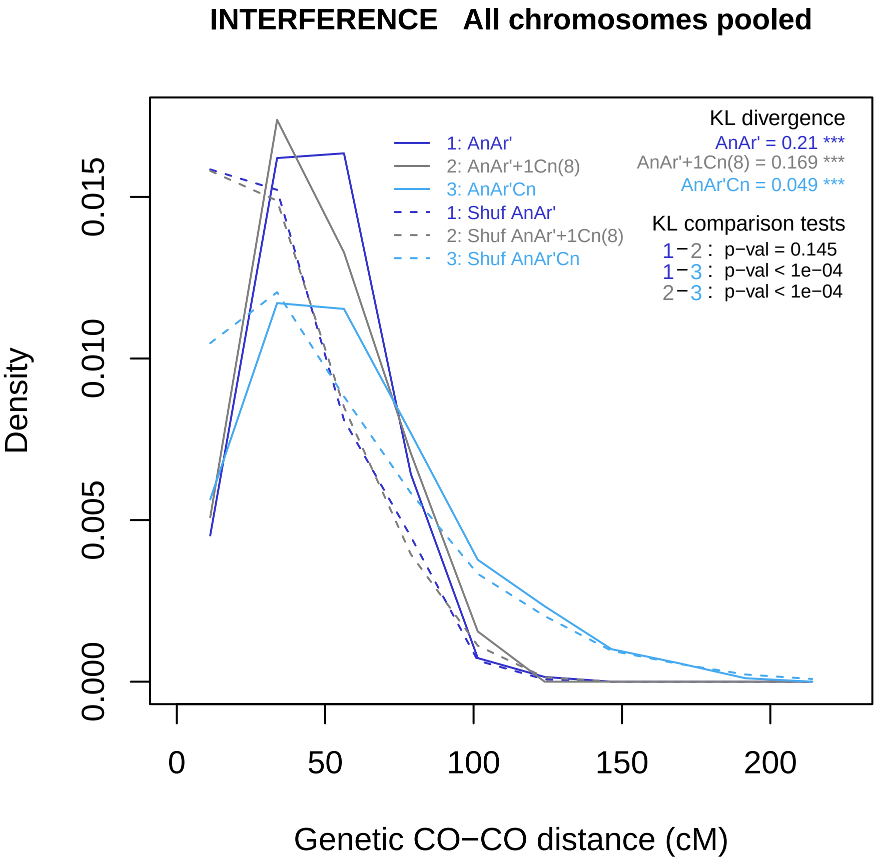
INTERFERENCE   All chromosomes pooled
Density
0.000
0.005
0.010
0.015
0
50
100
150
200
Genetic CO−CO distance (cM)
1: AnAr'
2: AnAr'+1Cn(8)
3: AnAr'Cn
1: Shuf AnAr'
2: Shuf AnAr'+1Cn(8)
3: Shuf AnAr'Cn
KL divergence
AnAr' = 0.21 ***
AnAr'+1Cn(8) = 0.169 ***
AnAr'Cn = 0.049 ***
KL comparison tests
1−2 : p−val = 0.145
1−3 : p−val < 1e−04
2−3 : p−val < 1e−04

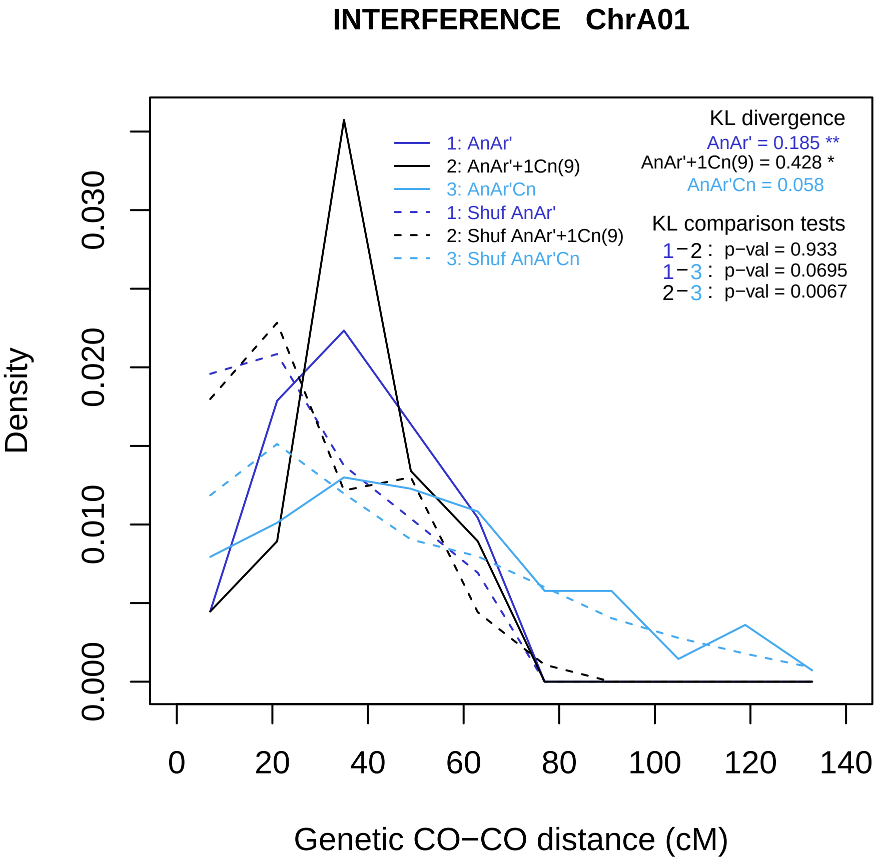

INTERFERENCE ChrA01
Density
0.000
0.010
0.020
0.030
0
20
40
60
80
100
120
140
Genetic CO−CO distance (cM)
1: AnAr'
2: AnAr'+1Cn(9)
3: AnAr'Cn
1: Shuf AnAr'
2: Shuf AnAr'+1Cn(9)
3: Shuf AnAr'Cn
KL divergence
AnAr' = 0.185 **
AnAr'+1Cn(9) = 0.428 *
AnAr'Cn = 0.058
KL comparison tests
1−2 : p−val = 0.933
1−3 : p−val = 0.0695
2−3 : p−val = 0.0067

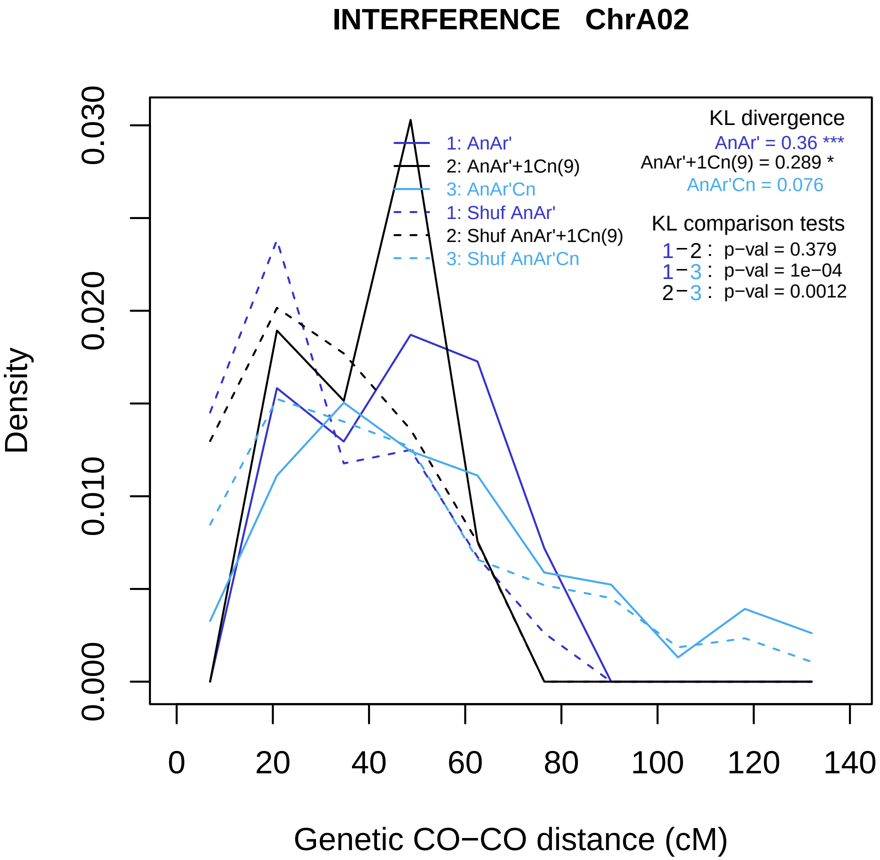
INTERFERENCE ChrA02
KL divergence
AnAr' = 0.36 ***
AnAr'+1Cn(9) = 0.289 *
AnAr'Cn = 0.076
1: AnAr'
2: AnAr'+1Cn(9)
3: AnAr'Cn
1: Shuf AnAr'
2: Shuf AnAr'+1Cn(9)
3: Shuf AnAr'Cn
KL comparison tests
1−2 : p−val = 0.379
1−3 : p−val = 1e−04
2−3 : p−val = 0.0012
Density
0.000
0.010
0.020
0.030
0
20
40
60
80
100
120
140
Genetic CO−CO distance (cM)

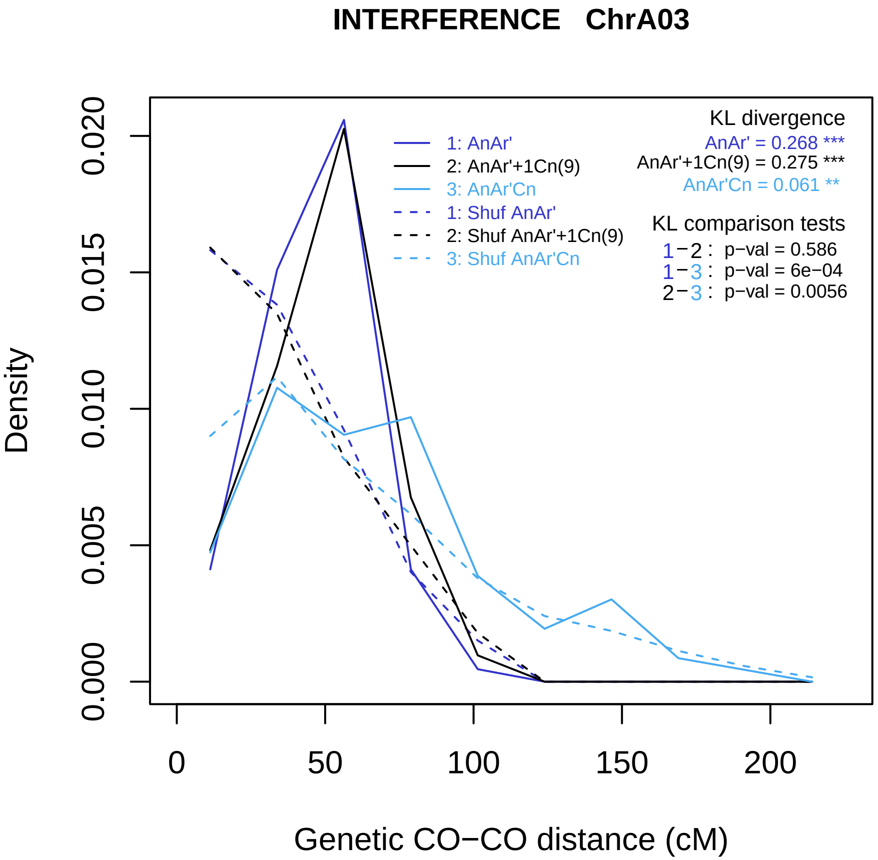
INTERFERENCE ChrA03
Density
0.000
0.005
0.010
0.015
0.020
0
50
100
150
200
Genetic CO−CO distance (cM)
1: AnAr'
2: AnAr'+1Cn(9)
3: AnAr'Cn
1: Shuf AnAr'
2: Shuf AnAr'+1Cn(9)
3: Shuf AnAr'Cn
KL divergence
AnAr' = 0.268 ***
AnAr'+1Cn(9) = 0.275 ***
AnAr'Cn = 0.061 **
KL comparison tests
1−2 : p−val = 0.586
1−3 : p−val = 6e−04
2−3 : p−val = 0.0056

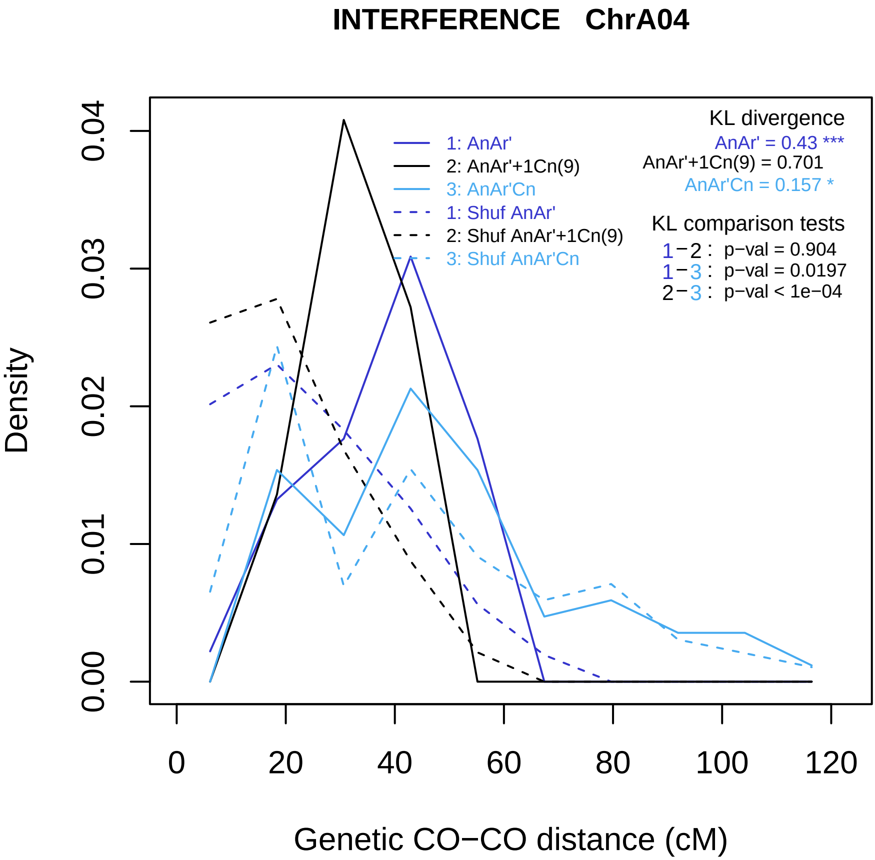
INTERFERENCE ChrA04
Density
0.00
0.01
0.02
0.03
0.04
0
20
40
60
80
100
120
Genetic CO−CO distance (cM)
1: AnAr'
2: AnAr'+1Cn(9)
3: AnAr'Cn
1: Shuf AnAr'
2: Shuf AnAr'+1Cn(9)
3: Shuf AnAr'Cn
KL divergence
AnAr' = 0.43 ***
AnAr'+1Cn(9) = 0.701
AnAr'Cn = 0.157 *
KL comparison tests
1−2 : p−val = 0.904
1−3 : p−val = 0.0197
2−3 : p−val < 1e−04

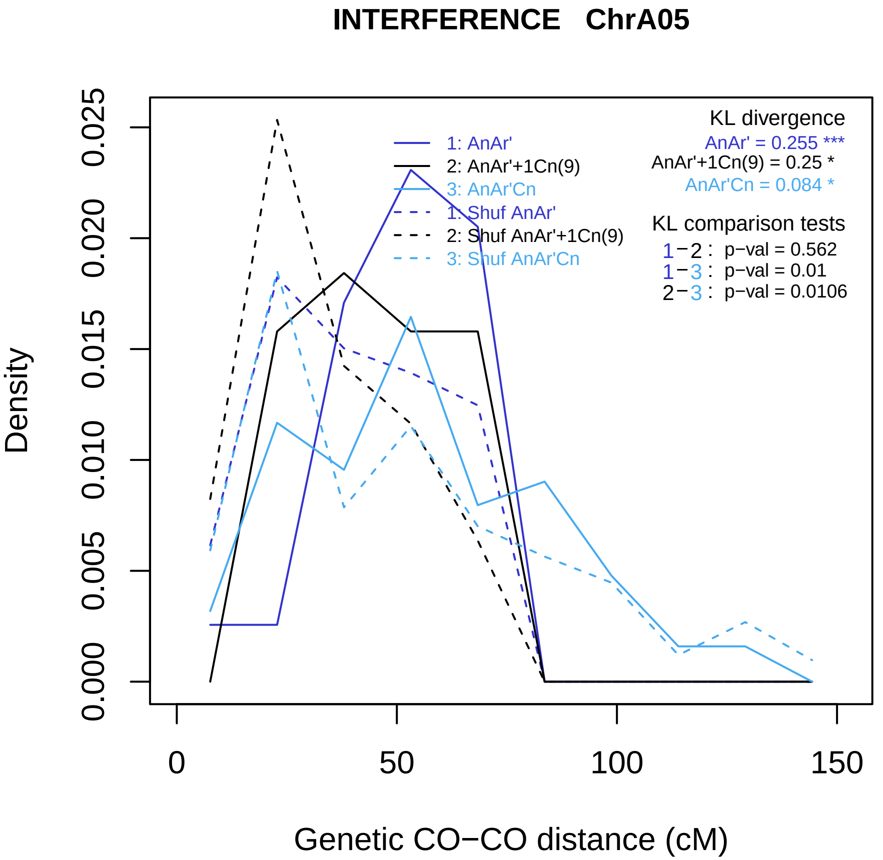

INTERFERENCE ChrA05
1: AnAr'
2: AnAr'+1Cn(9)
3: AnAr'Cn
1: Shuf AnAr'
2: Shuf AnAr'+1Cn(9)
3: Shuf AnAr'Cn
KL divergence
AnAr' = 0.255 ***
AnAr'+1Cn(9) = 0.25 *
AnAr'Cn = 0.084 *
KL comparison tests
1−2 : p−val = 0.562
1−3 : p−val = 0.01
2−3 : p−val = 0.0106
Density
0.000
0.005
0.010
0.015
0.020
0.025
0
50
100
150
Genetic CO−CO distance (cM)

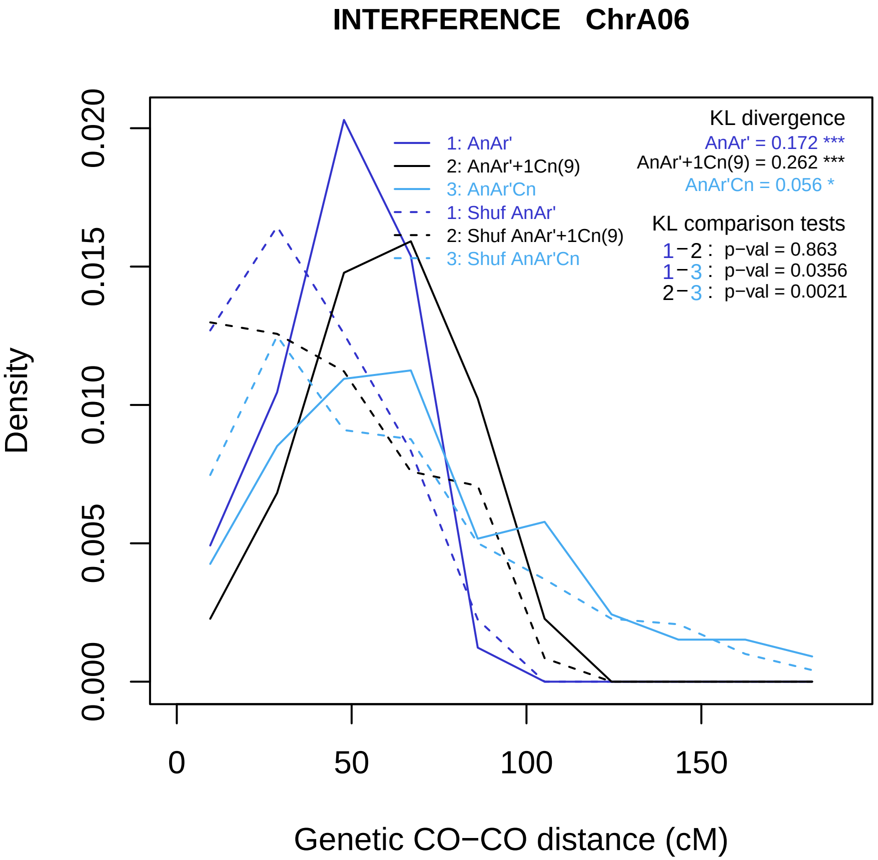
INTERFERENCE ChrA06
Density
Genetic CO−CO distance (cM)
0.000
0.005
0.010
0.015
0.020
0
50
100
150
1: AnAr'
2: AnAr'+1Cn(9)
3: AnAr'Cn
1: Shuf AnAr'
2: Shuf AnAr'+1Cn(9)
3: Shuf AnAr'Cn
KL divergence
AnAr' = 0.172 ***
AnAr'+1Cn(9) = 0.262 ***
AnAr'Cn = 0.056 *
KL comparison tests
1−2 : p−val = 0.863
1−3 : p−val = 0.0356
2−3 : p−val = 0.0021

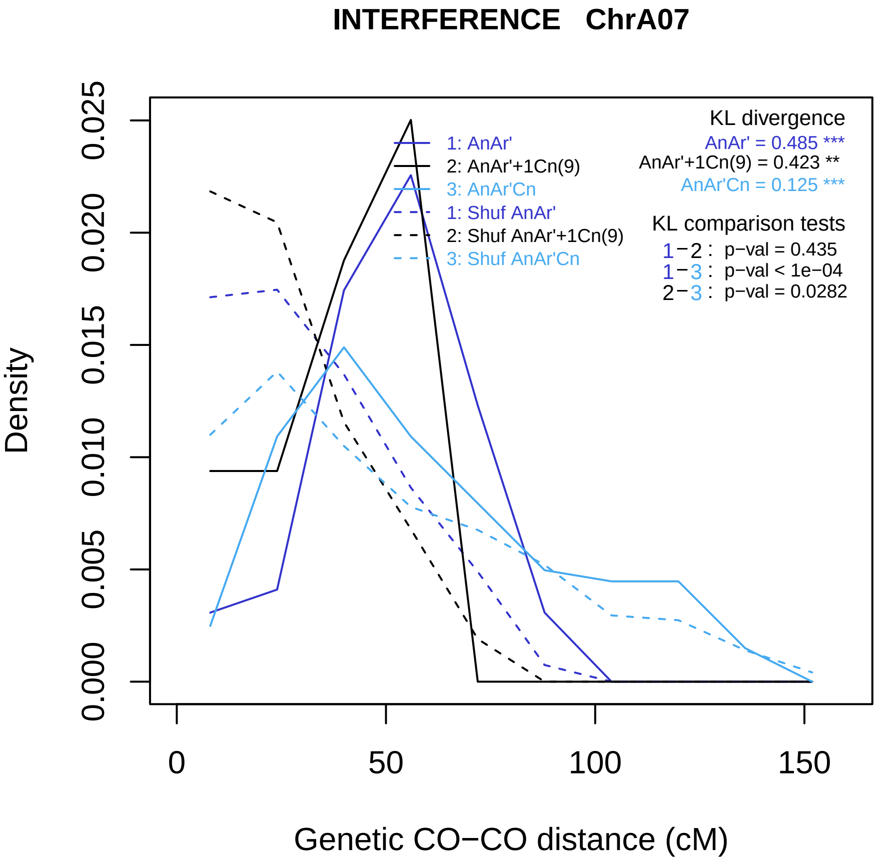
INTERFERENCE ChrA07
KL divergence
AnAr' = 0.485 ***
AnAr'+1Cn(9) = 0.423 **
AnAr'Cn = 0.125 ***
1: AnAr'
2: AnAr'+1Cn(9)
3: AnAr'Cn
1: Shuf AnAr'
2: Shuf AnAr'+1Cn(9)
3: Shuf AnAr'Cn
KL comparison tests
1−2 : p−val = 0.435
1−3 : p−val < 1e−04
2−3 : p−val = 0.0282
Density
0.000
0.005
0.010
0.015
0.020
0.025
0
50
100
150
Genetic CO−CO distance (cM)

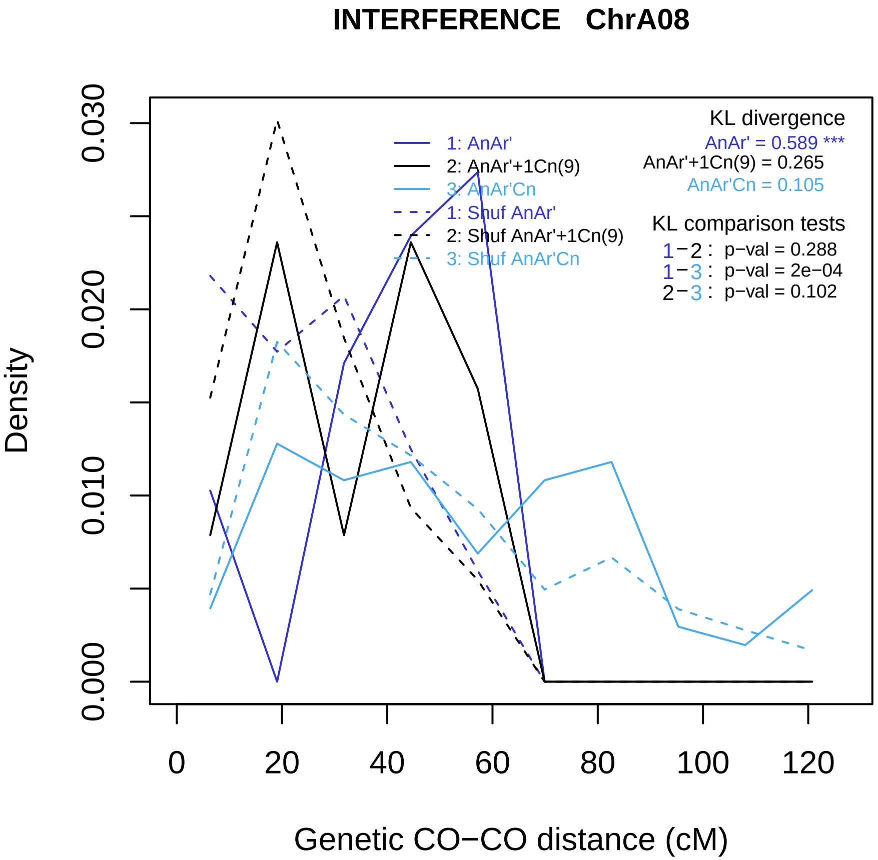
INTERFERENCE ChrA08
KL divergence
AnAr' = 0.589 ***
AnAr'+1Cn(9) = 0.265
AnAr'Cn = 0.105
1: AnAr'
2: AnAr'+1Cn(9)
3: AnAr'Cn
1: Shuf AnAr'
2: Shuf AnAr'+1Cn(9)
3: Shuf AnAr'Cn
KL comparison tests
1−2 : p−val = 0.288
1−3 : p−val = 2e−04
2−3 : p−val = 0.102
Density
0.000
0.010
0.020
0.030
0
20
40
60
80
100
120
Genetic CO−CO distance (cM)

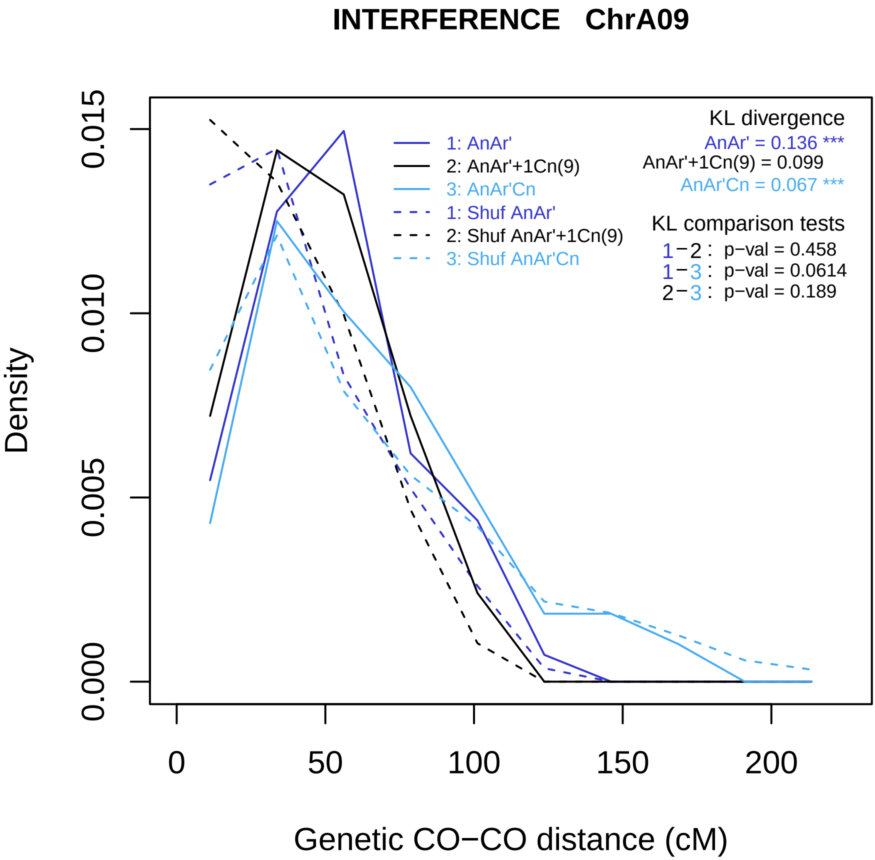

INTERFERENCE ChrA09
Density
0.000
0.005
0.010
0.015
0
50
100
150
200
Genetic CO−CO distance (cM)
1: AnAr'
2: AnAr'+1Cn(9)
3: AnAr'Cn
1: Shuf AnAr'
2: Shuf AnAr'+1Cn(9)
3: Shuf AnAr'Cn
KL divergence
AnAr' = 0.136 ***
AnAr'+1Cn(9) = 0.099
AnAr'Cn = 0.067 ***
KL comparison tests
1−2 : p−val = 0.458
1−3 : p−val = 0.0614
2−3 : p−val = 0.189

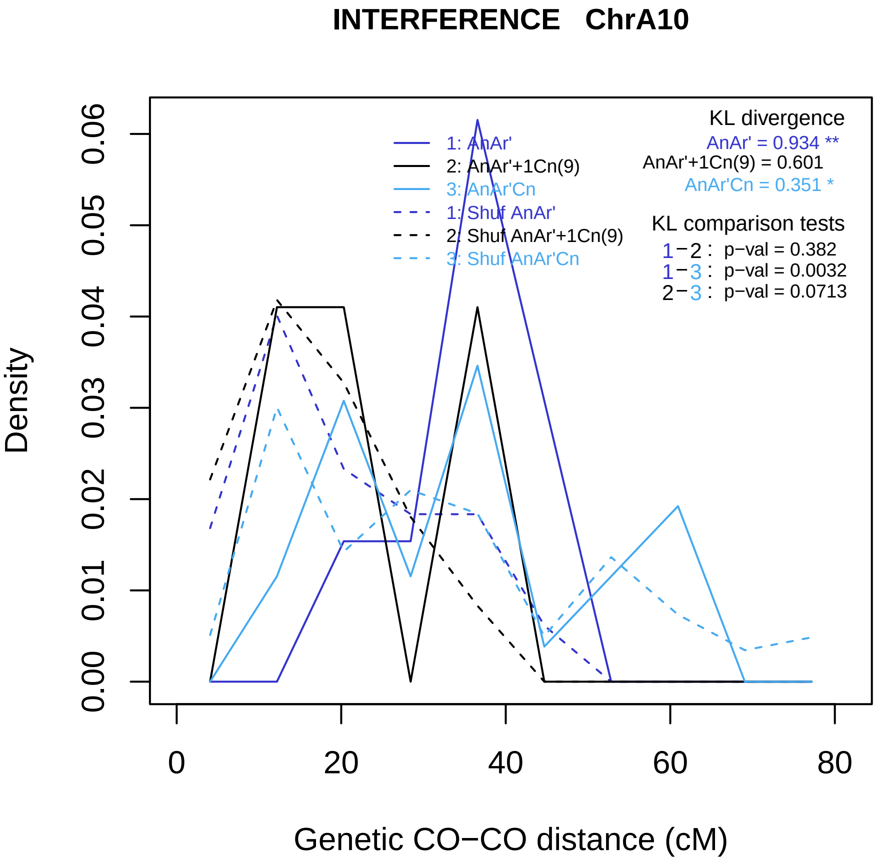
INTERFERENCE ChrA10
Density
0.00
0.01
0.02
0.03
0.04
0.05
0.06
0
20
40
60
80
Genetic CO−CO distance (cM)
1: AnAr'
2: AnAr'+1Cn(9)
3: AnAr'Cn
1: Shuf AnAr'
2: Shuf AnAr'+1Cn(9)
3: Shuf AnAr'Cn
KL divergence
AnAr' = 0.934 **
AnAr'+1Cn(9) = 0.601
AnAr'Cn = 0.351 *
KL comparison tests
1−2 : p−val = 0.382
1−3 : p−val = 0.0032
2−3 : p−val = 0.0713

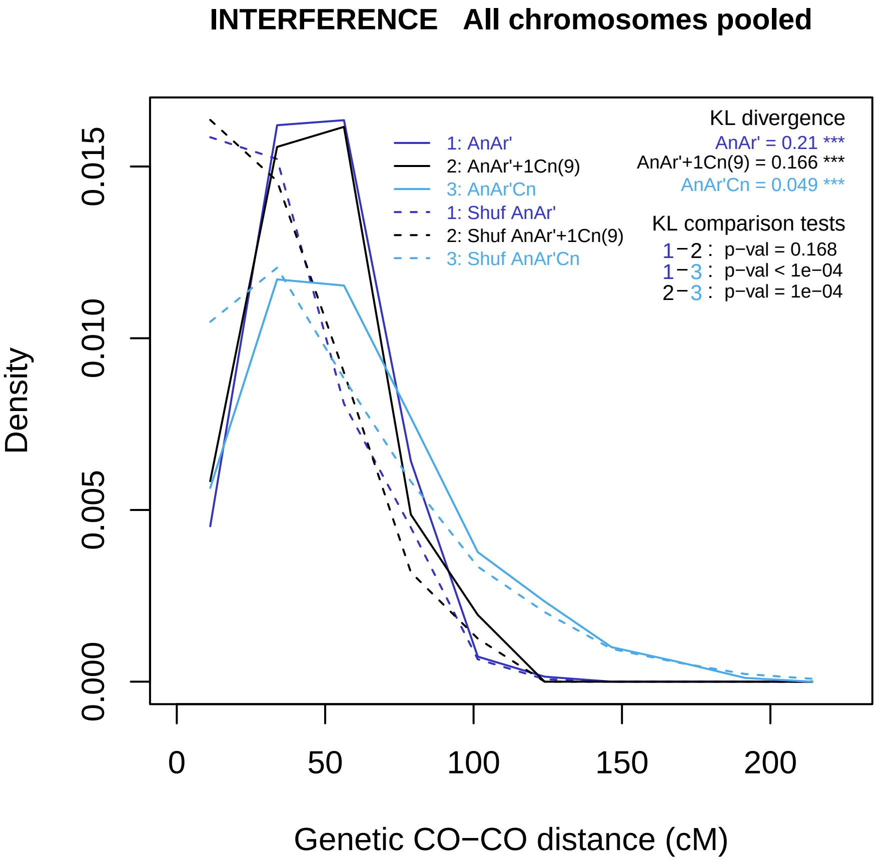
INTERFERENCE All chromosomes pooled
Density
0.000
0.005
0.010
0.015
0
50
100
150
200
Genetic CO−CO distance (cM)
1: AnAr'
2: AnAr'+1Cn(9)
3: AnAr'Cn
1: Shuf AnAr'
2: Shuf AnAr'+1Cn(9)
3: Shuf AnAr'Cn
KL divergence
AnAr' = 0.21 ***
AnAr'+1Cn(9) = 0.166 ***
AnAr'Cn = 0.049 ***
KL comparison tests
1−2 : p−val = 0.168
1−3 : p−val < 1e−04
2−3 : p−val = 1e−04

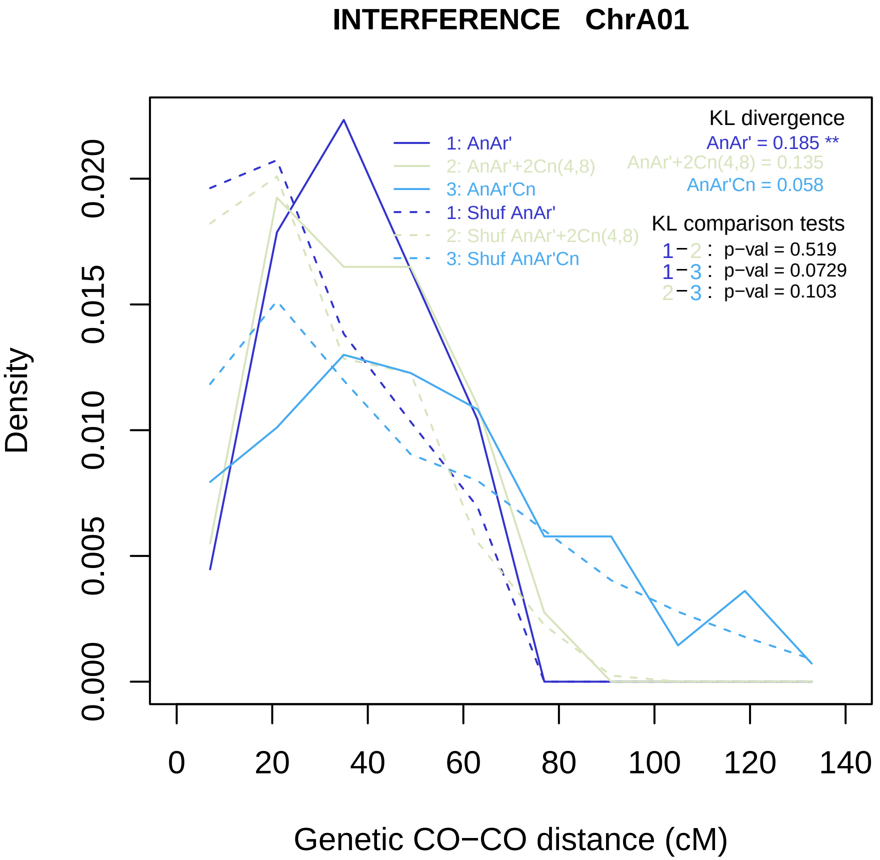

INTERFERENCE ChrA01
Density
0.000
0.005
0.010
0.015
0.020
0
20
40
60
80
100
120
140
Genetic CO−CO distance (cM)
1: AnAr'
2: AnAr'+2Cn(4,8)
3: AnAr'Cn
1: Shuf AnAr'
2: Shuf AnAr'+2Cn(4,8)
3: Shuf AnAr'Cn
KL divergence
AnAr' = 0.185 **
AnAr'+2Cn(4,8) = 0.135
AnAr'Cn = 0.058
KL comparison tests
1−2 : p−val = 0.519
1−3 : p−val = 0.0729
2−3 : p−val = 0.103

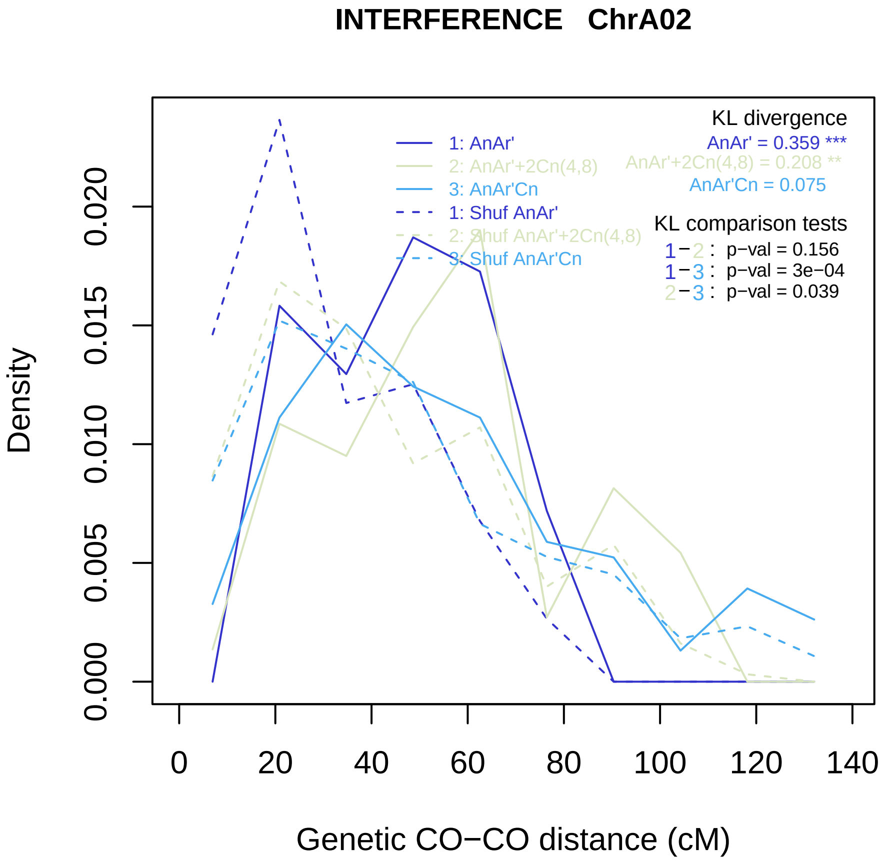
INTERFERENCE ChrA02
KL divergence
AnAr' = 0.359 ***
AnAr'+2Cn(4,8) = 0.208 **
AnAr'Cn = 0.075
1: AnAr'
2: AnAr'+2Cn(4,8)
3: AnAr'Cn
1: Shuf AnAr'
2: Shuf AnAr'+2Cn(4,8)
3: Shuf AnAr'Cn
KL comparison tests
1−2 : p−val = 0.156
1−3 : p−val = 3e−04
2−3 : p−val = 0.039
Density
0.000
0.005
0.010
0.015
0.020
0
20
40
60
80
100
120
140
Genetic CO−CO distance (cM)

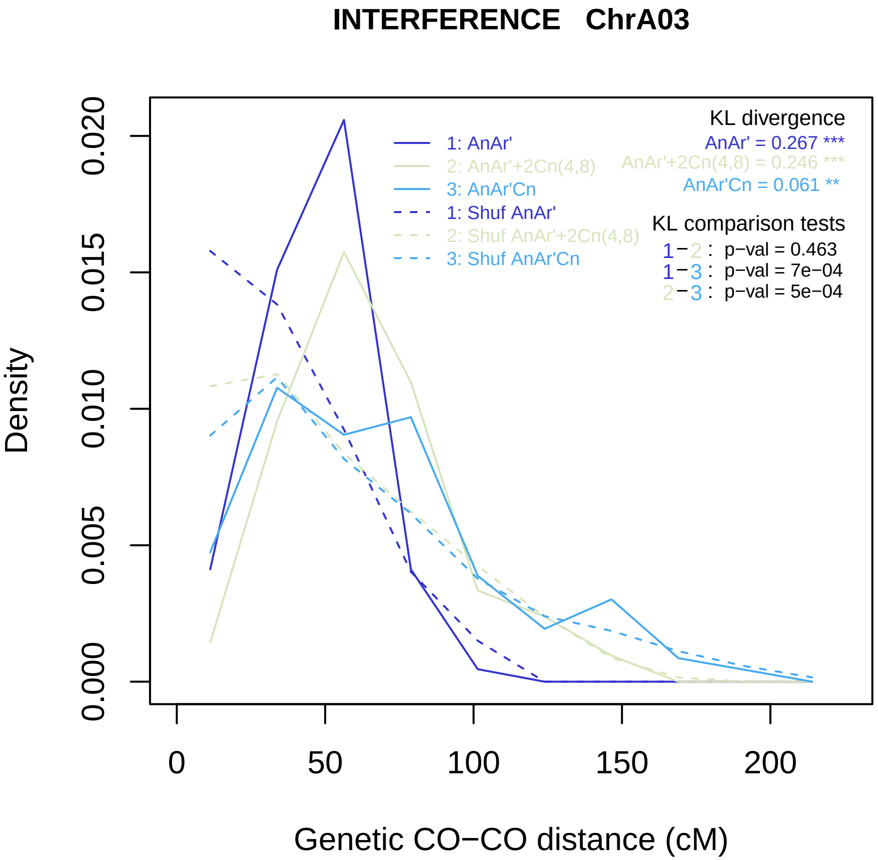
INTERFERENCE ChrA03
Density
Genetic CO−CO distance (cM)
0.000
0.005
0.010
0.015
0.020
0
50
100
150
200
1: AnAr'
2: AnAr'+2Cn(4,8)
3: AnAr'Cn
1: Shuf AnAr'
2: Shuf AnAr'+2Cn(4,8)
3: Shuf AnAr'Cn
KL divergence
AnAr' = 0.267 ***
AnAr'+2Cn(4,8) = 0.246 ***
AnAr'Cn = 0.061 **
KL comparison tests
1−2 : p−val = 0.463
1−3 : p−val = 7e−04
2−3 : p−val = 5e−04

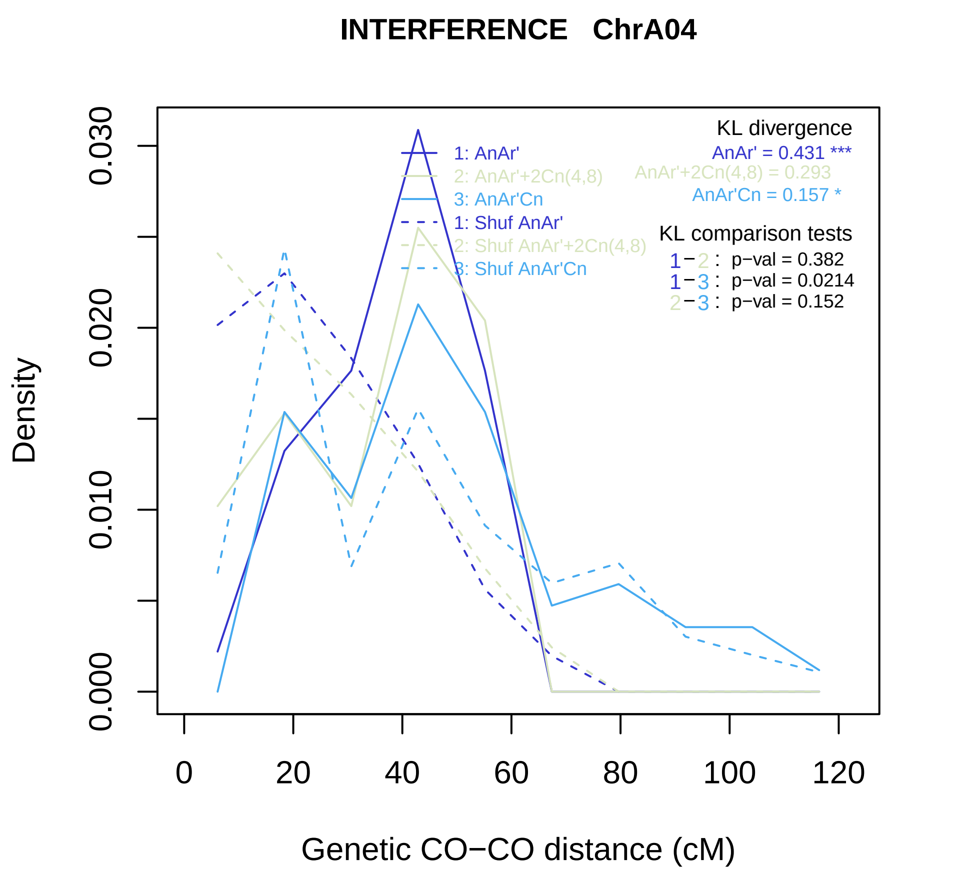
INTERFERENCE ChrA04
Density
0.000
0.010
0.020
0.030
0
20
40
60
80
100
120
Genetic CO−CO distance (cM)
1: AnAr'
2: AnAr'+2Cn(4,8)
3: AnAr'Cn
1: Shuf AnAr'
2: Shuf AnAr'+2Cn(4,8)
3: Shuf AnAr'Cn
KL divergence
AnAr' = 0.431 ***
AnAr'+2Cn(4,8) = 0.293
AnAr'Cn = 0.157 *
KL comparison tests
1−2 : p−val = 0.382
1−3 : p−val = 0.0214
2−3 : p−val = 0.152

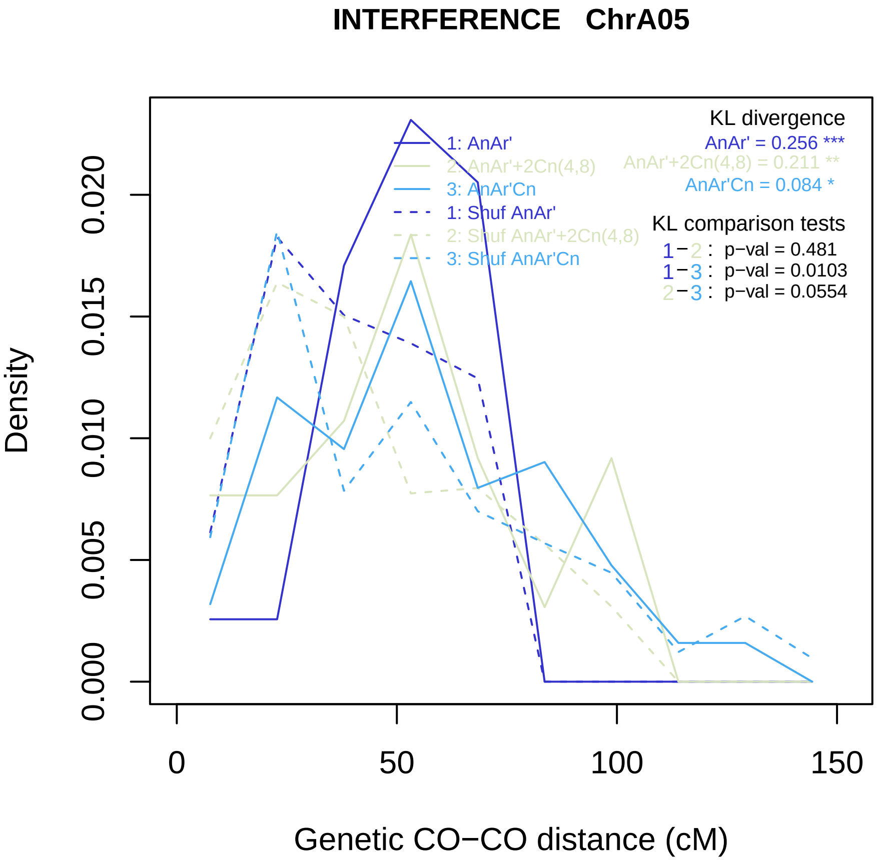

INTERFERENCE ChrA05
KL divergence
AnAr' = 0.256 ***
AnAr'+2Cn(4,8) = 0.211 **
AnAr'Cn = 0.084 *
1: AnAr'
2: AnAr'+2Cn(4,8)
3: AnAr'Cn
1: Shuf AnAr'
2: Shuf AnAr'+2Cn(4,8)
3: Shuf AnAr'Cn
KL comparison tests
1−2 : p−val = 0.481
1−3 : p−val = 0.0103
2−3 : p−val = 0.0554
Density
0.000
0.005
0.010
0.015
0.020
0
50
100
150
Genetic CO−CO distance (cM)

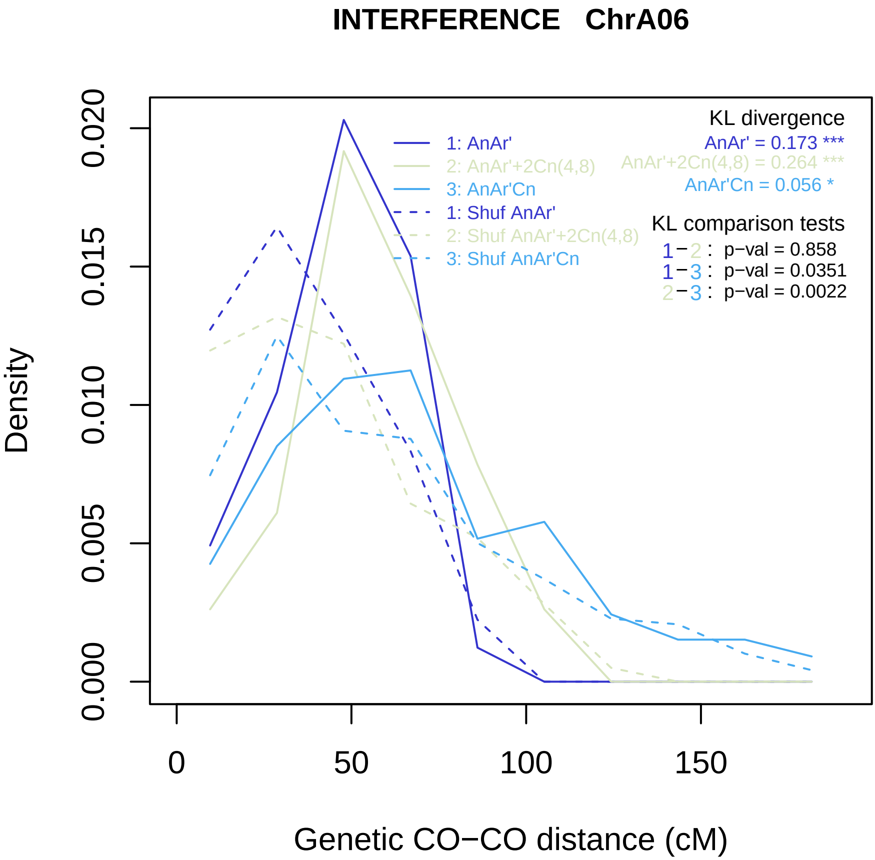
INTERFERENCE ChrA06
Density
0.000
0.005
0.010
0.015
0.020
0
50
100
150
Genetic CO−CO distance (cM)
1: AnAr'
2: AnAr'+2Cn(4,8)
3: AnAr'Cn
1: Shuf AnAr'
2: Shuf AnAr'+2Cn(4,8)
3: Shuf AnAr'Cn
KL divergence
AnAr' = 0.173 ***
AnAr'+2Cn(4,8) = 0.264 ***
AnAr'Cn = 0.056 *
KL comparison tests
1−2 : p−val = 0.858
1−3 : p−val = 0.0351
2−3 : p−val = 0.0022

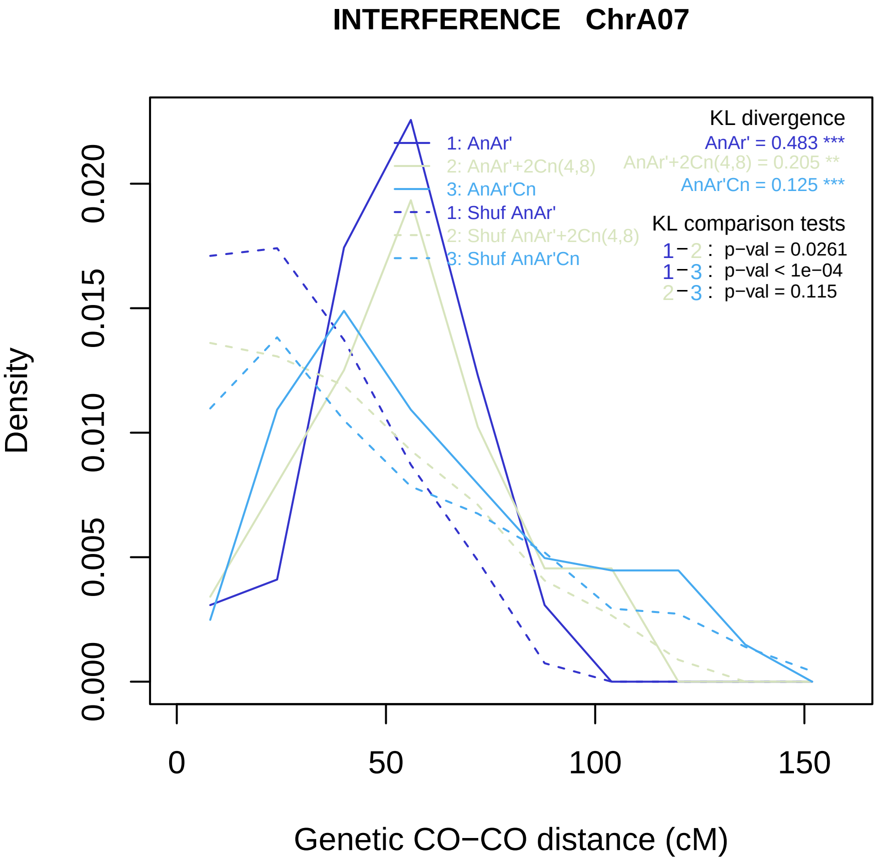
INTERFERENCE ChrA07
KL divergence
AnAr' = 0.483 ***
AnAr'+2Cn(4,8) = 0.205 **
AnAr'Cn = 0.125 ***
1: AnAr'
2: AnAr'+2Cn(4,8)
3: AnAr'Cn
1: Shuf AnAr'
2: Shuf AnAr'+2Cn(4,8)
3: Shuf AnAr'Cn
KL comparison tests
1−2 : p−val = 0.0261
1−3 : p−val < 1e−04
2−3 : p−val = 0.115
Density
0.000
0.005
0.010
0.015
0.020
0
50
100
150
Genetic CO−CO distance (cM)

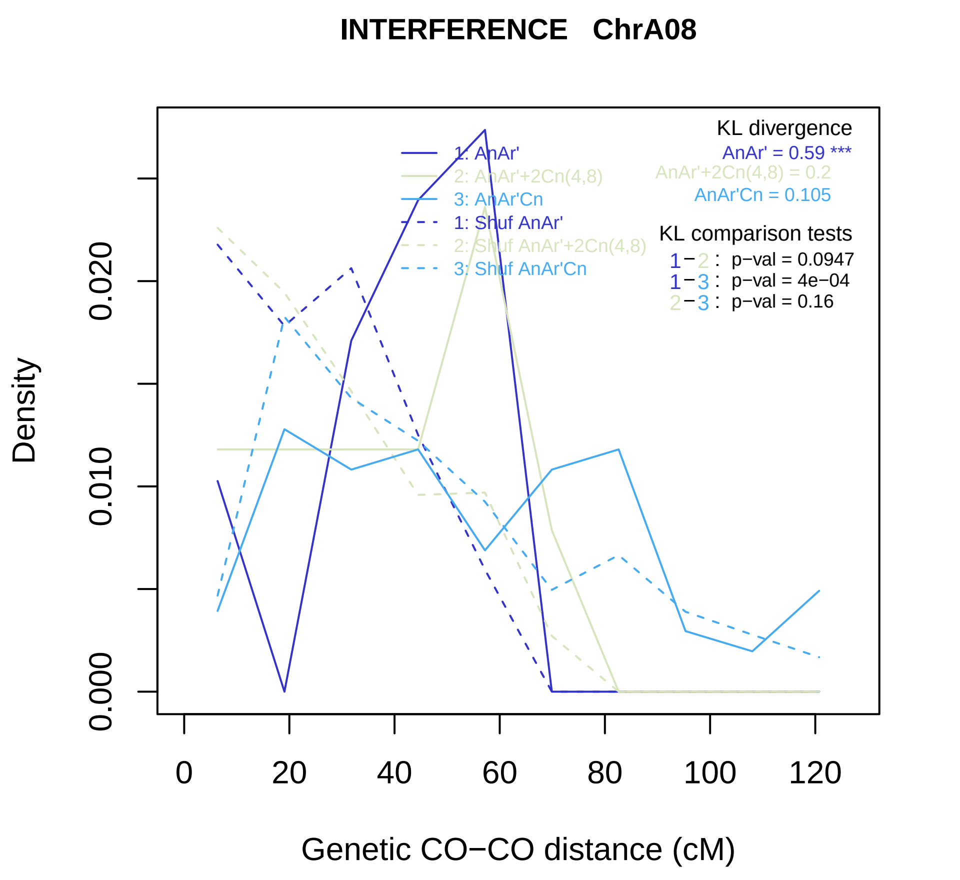

INTERFERENCE ChrA08
KL divergence
AnAr' = 0.59 ***
AnAr'+2Cn(4,8) = 0.2
AnAr'Cn = 0.105
1: AnAr'
2: AnAr'+2Cn(4,8)
3: AnAr'Cn
1: Shuf AnAr'
2: Shuf AnAr'+2Cn(4,8)
3: Shuf AnAr'Cn
KL comparison tests
1−2 : p−val = 0.0947
1−3 : p−val = 4e−04
2−3 : p−val = 0.16
Density
0.000
0.010
0.020
0
20
40
60
80
100
120
Genetic CO−CO distance (cM)

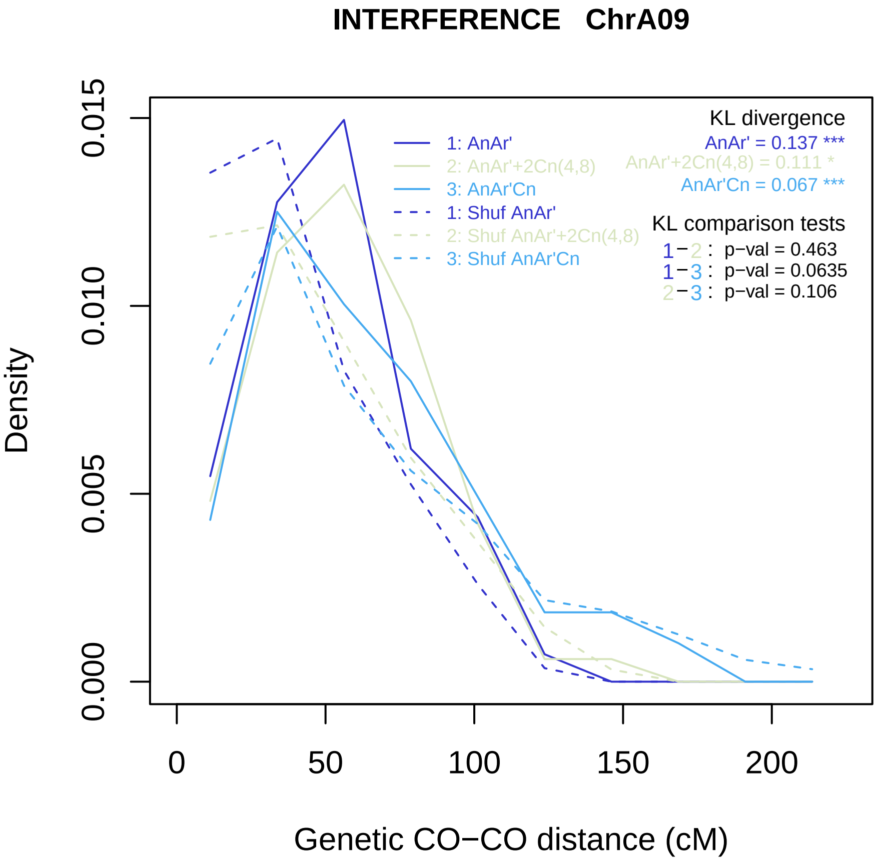

INTERFERENCE ChrA09
Density
Genetic CO−CO distance (cM)
0.000
0.005
0.010
0.015
0
50
100
150
200
1: AnAr'
2: AnAr'+2Cn(4,8)
3: AnAr'Cn
1: Shuf AnAr'
2: Shuf AnAr'+2Cn(4,8)
3: Shuf AnAr'Cn
KL divergence
AnAr' = 0.137 ***
AnAr'+2Cn(4,8) = 0.111 *
AnAr'Cn = 0.067 ***
KL comparison tests
1−2 : p−val = 0.463
1−3 : p−val = 0.0635
2−3 : p−val = 0.106

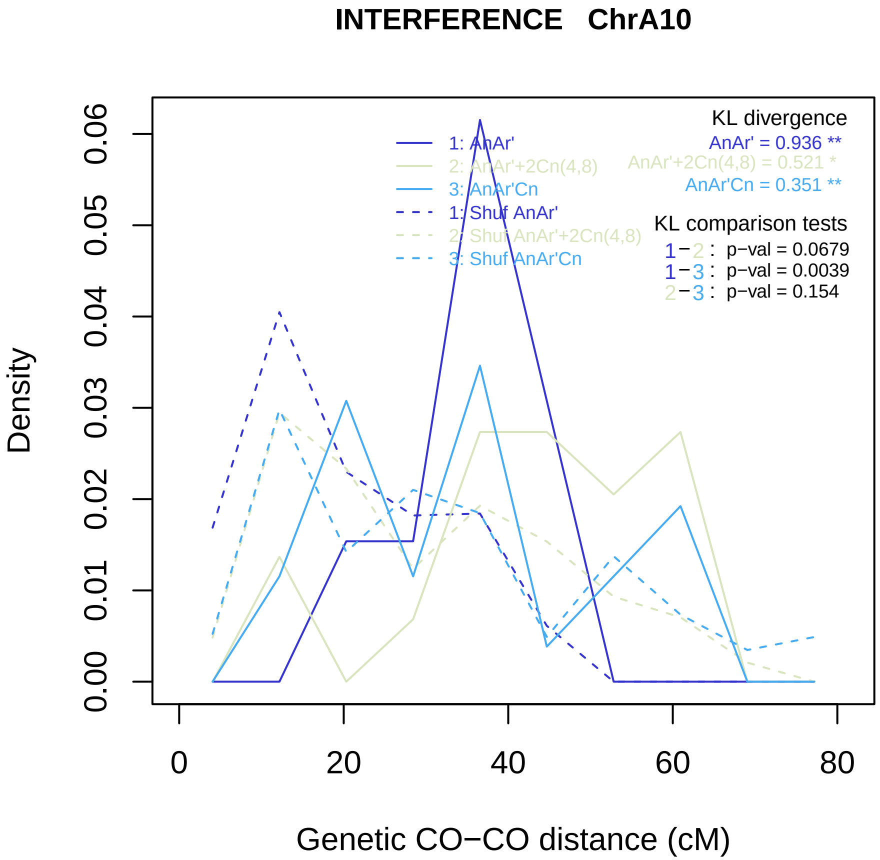
INTERFERENCE ChrA10
Density
0.00
0.01
0.02
0.03
0.04
0.05
0.06
0
20
40
60
80
Genetic CO−CO distance (cM)
1: AnAr'
2: AnAr'+2Cn(4,8)
3: AnAr'Cn
1: Shuf AnAr'
2: Shuf AnAr'+2Cn(4,8)
3: Shuf AnAr'Cn
KL divergence
AnAr' = 0.936 **
AnAr'+2Cn(4,8) = 0.521 *
AnAr'Cn = 0.351 **
KL comparison tests
1−2 : p−val = 0.0679
1−3 : p−val = 0.0039
2−3 : p−val = 0.154

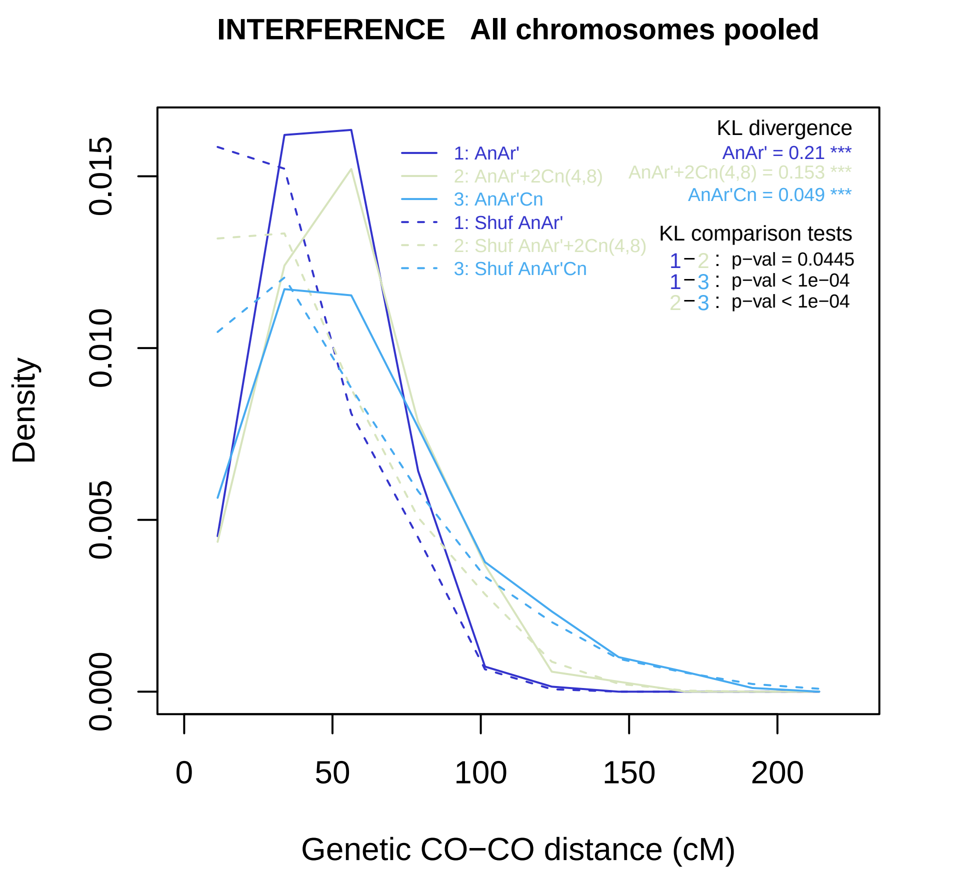
INTERFERENCE   All chromosomes pooled
Density
0.000
0.005
0.010
0.015
0
50
100
150
200
Genetic CO−CO distance (cM)
1: AnAr'
2: AnAr'+2Cn(4,8)
3: AnAr'Cn
1: Shuf AnAr'
2: Shuf AnAr'+2Cn(4,8)
3: Shuf AnAr'Cn
KL divergence
AnAr' = 0.21 ***
AnAr'+2Cn(4,8) = 0.153 ***
AnAr'Cn = 0.049 ***
KL comparison tests
1−2 : p−val = 0.0445
1−3 : p−val < 1e−04
2−3 : p−val < 1e−04

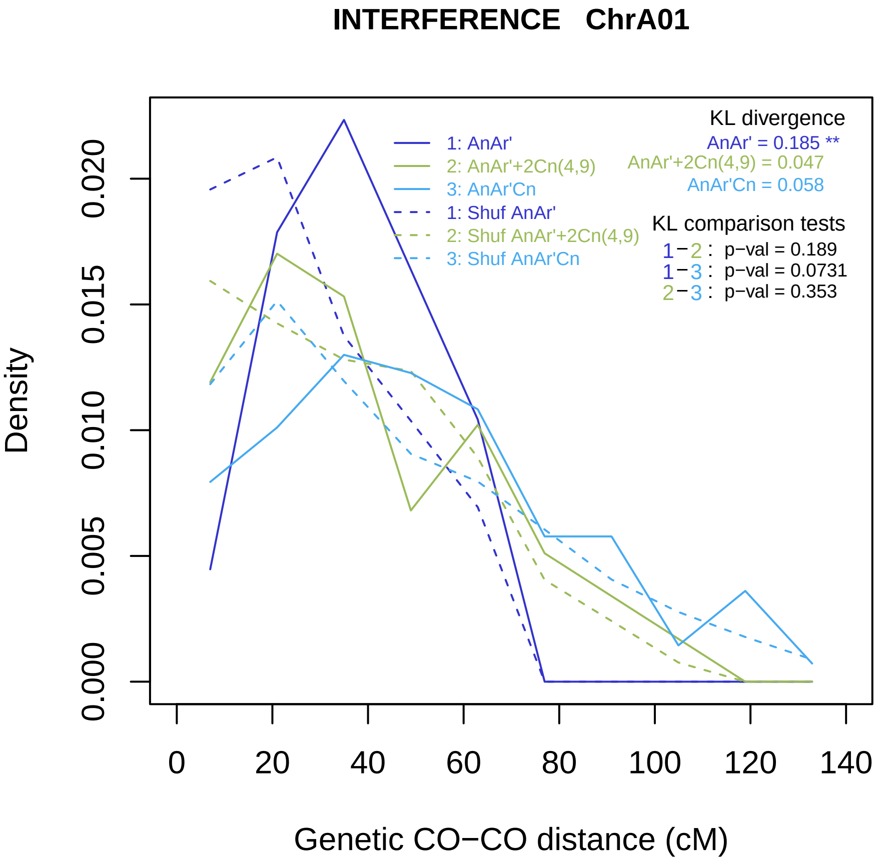
INTERFERENCE ChrA01
KL divergence
AnAr' = 0.185 **
AnAr'+2Cn(4,9) = 0.047
AnAr'Cn = 0.058
1: AnAr'
2: AnAr'+2Cn(4,9)
3: AnAr'Cn
1: Shuf AnAr'
2: Shuf AnAr'+2Cn(4,9)
3: Shuf AnAr'Cn
KL comparison tests
1−2 : p−val = 0.189
1−3 : p−val = 0.0731
2−3 : p−val = 0.353
Density
0.000
0.005
0.010
0.015
0.020
0
20
40
60
80
100
120
140
Genetic CO−CO distance (cM)

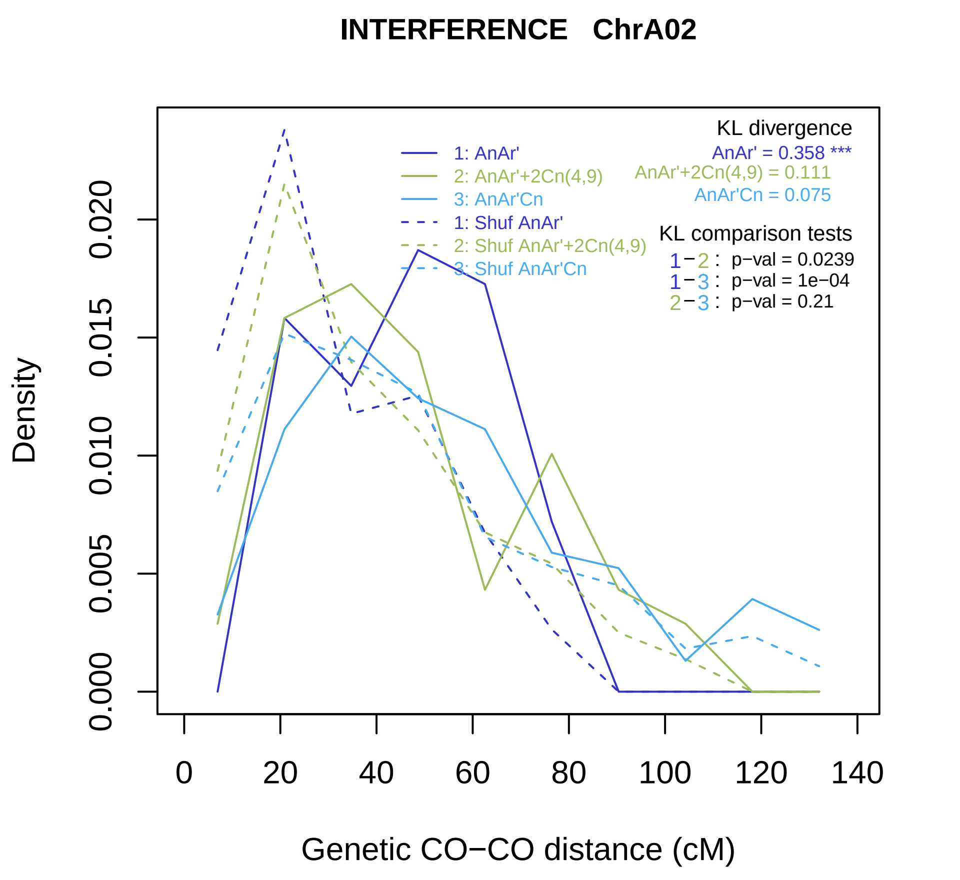
INTERFERENCE ChrA02
Density
0.000
0.005
0.010
0.015
0.020
0
20
40
60
80
100
120
140
Genetic CO−CO distance (cM)
1: AnAr'
2: AnAr'+2Cn(4,9)
3: AnAr'Cn
1: Shuf AnAr'
2: Shuf AnAr'+2Cn(4,9)
3: Shuf AnAr'Cn
KL divergence
AnAr' = 0.358 ***
AnAr'+2Cn(4,9) = 0.111
AnAr'Cn = 0.075
KL comparison tests
1−2 : p−val = 0.0239
1−3 : p−val = 1e−04
2−3 : p−val = 0.21

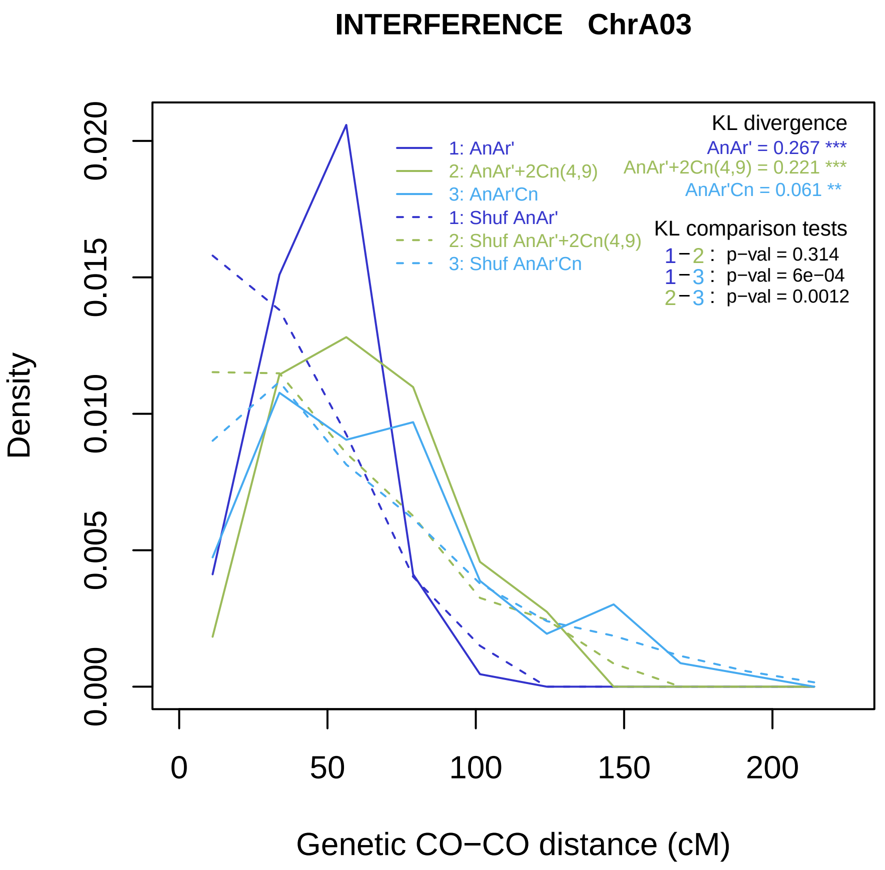
INTERFERENCE ChrA03
Density
0.000
0.005
0.010
0.015
0.020
0
50
100
150
200
Genetic CO−CO distance (cM)
1: AnAr'
2: AnAr'+2Cn(4,9)
3: AnAr'Cn
1: Shuf AnAr'
2: Shuf AnAr'+2Cn(4,9)
3: Shuf AnAr'Cn
KL divergence
AnAr' = 0.267 ***
AnAr'+2Cn(4,9) = 0.221 ***
AnAr'Cn = 0.061 **
KL comparison tests
1−2 : p−val = 0.314
1−3 : p−val = 6e−04
2−3 : p−val = 0.0012

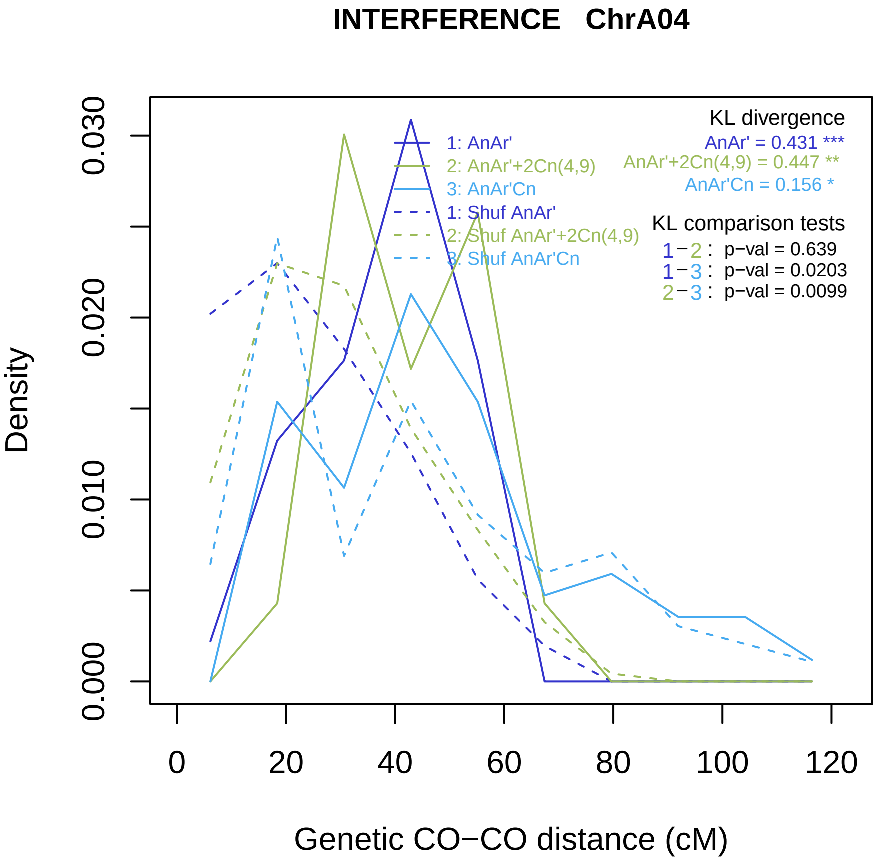

INTERFERENCE ChrA04
Density
0.000
0.010
0.020
0.030
0
20
40
60
80
100
120
Genetic CO−CO distance (cM)
1: AnAr'
2: AnAr'+2Cn(4,9)
3: AnAr'Cn
1: Shuf AnAr'
2: Shuf AnAr'+2Cn(4,9)
3: Shuf AnAr'Cn
KL divergence
AnAr' = 0.431 ***
AnAr'+2Cn(4,9) = 0.447 **
AnAr'Cn = 0.156 *
KL comparison tests
1−2 : p−val = 0.639
1−3 : p−val = 0.0203
2−3 : p−val = 0.0099

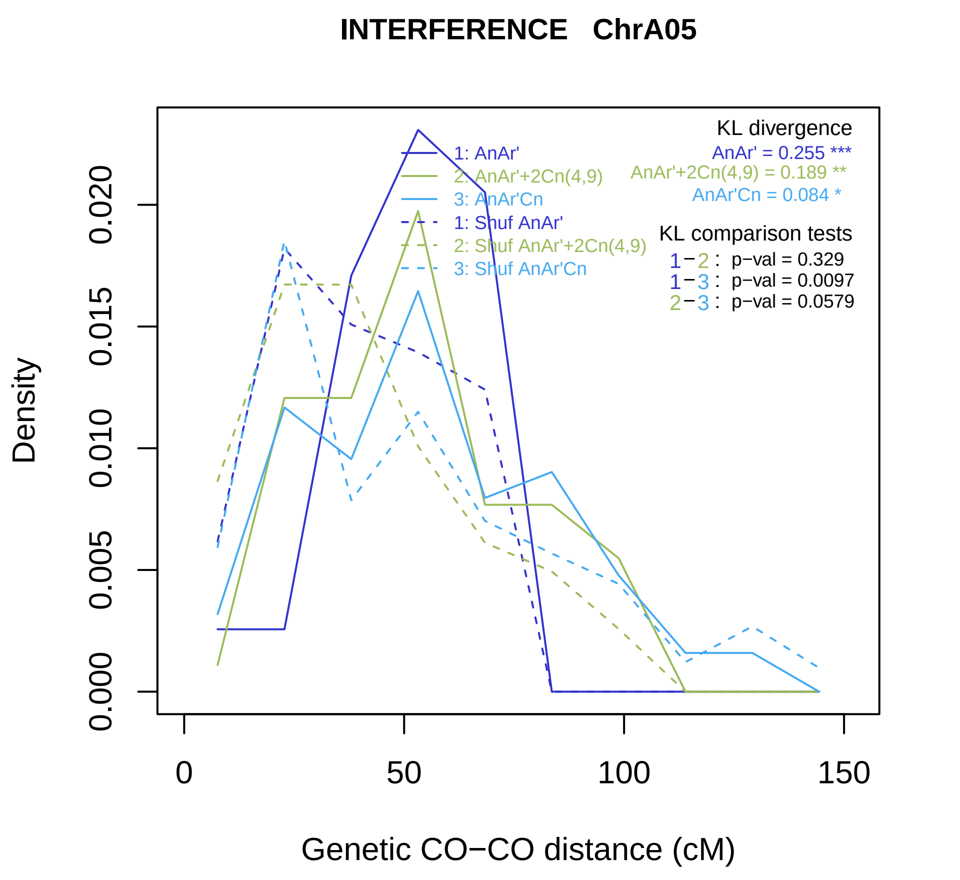
INTERFERENCE ChrA05
Density
0.000
0.005
0.010
0.015
0.020
0
50
100
150
Genetic CO−CO distance (cM)
1: AnAr'
2: AnAr'+2Cn(4,9)
3: AnAr'Cn
1: Shuf AnAr'
2: Shuf AnAr'+2Cn(4,9)
3: Shuf AnAr'Cn
KL divergence
AnAr' = 0.255 ***
AnAr'+2Cn(4,9) = 0.189 **
AnAr'Cn = 0.084 *
KL comparison tests
1−2 : p−val = 0.329
1−3 : p−val = 0.0097
2−3 : p−val = 0.0579

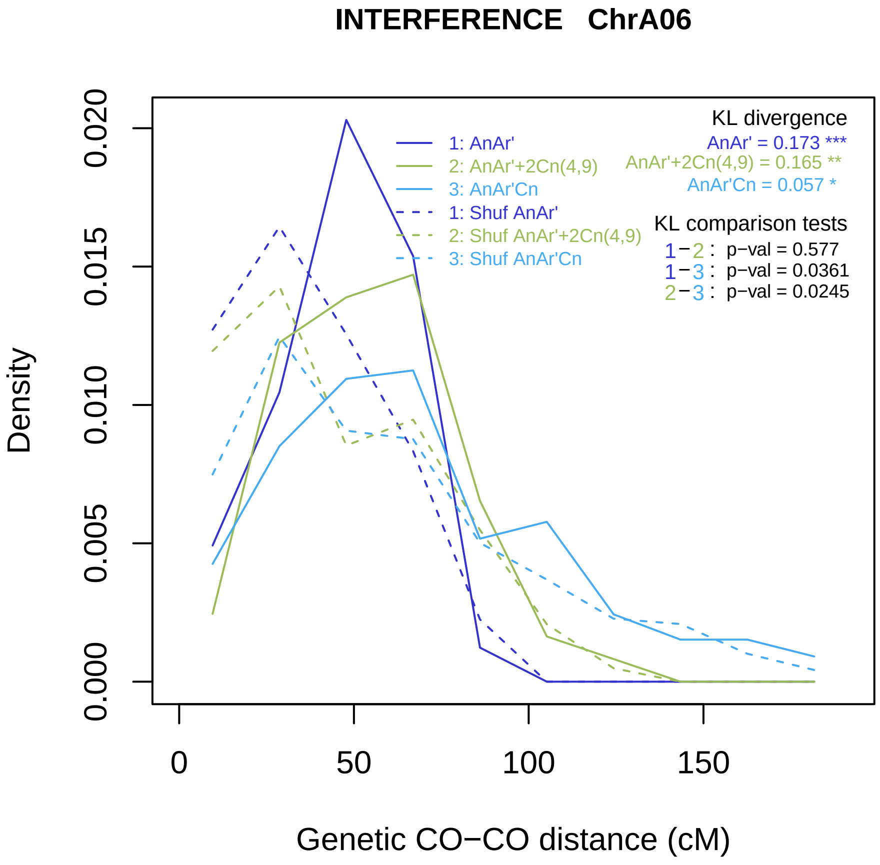
INTERFERENCE ChrA06
Density
0.000
0.005
0.010
0.015
0.020
0
50
100
150
Genetic CO−CO distance (cM)
1: AnAr'
2: AnAr'+2Cn(4,9)
3: AnAr'Cn
1: Shuf AnAr'
2: Shuf AnAr'+2Cn(4,9)
3: Shuf AnAr'Cn
KL divergence
AnAr' = 0.173 ***
AnAr'+2Cn(4,9) = 0.165 **
AnAr'Cn = 0.057 *
KL comparison tests
1−2 : p−val = 0.577
1−3 : p−val = 0.0361
2−3 : p−val = 0.0245

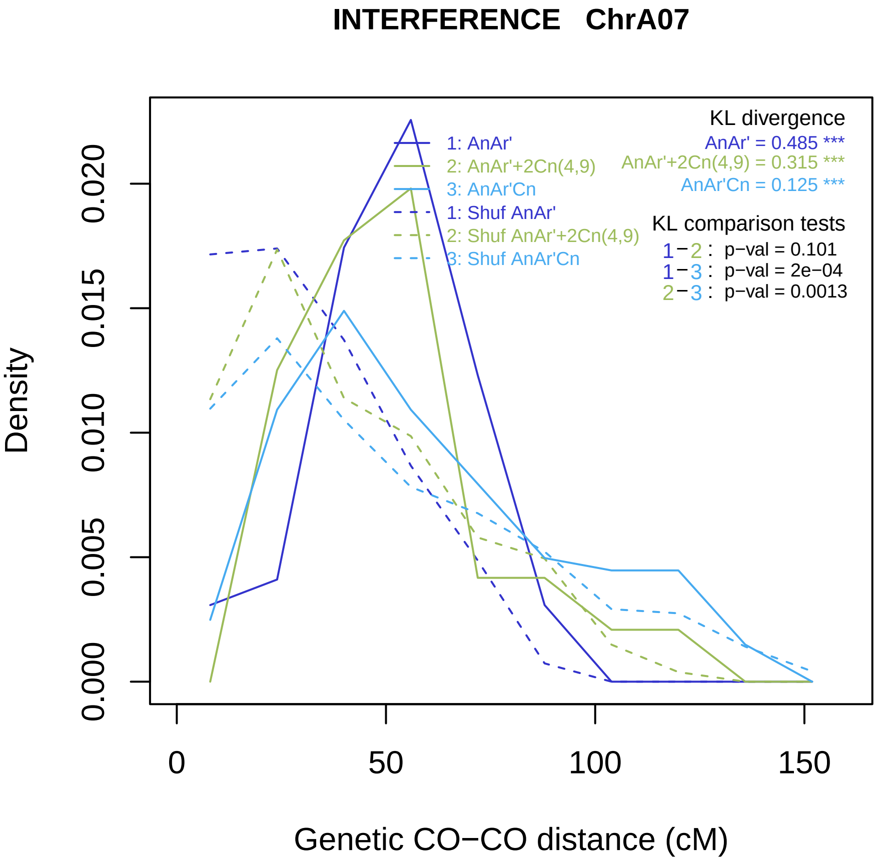
INTERFERENCE ChrA07
Density
0.000
0.005
0.010
0.015
0.020
0
50
100
150
Genetic CO−CO distance (cM)
1: AnAr'
2: AnAr'+2Cn(4,9)
3: AnAr'Cn
1: Shuf AnAr'
2: Shuf AnAr'+2Cn(4,9)
3: Shuf AnAr'Cn
KL divergence
AnAr' = 0.485 ***
AnAr'+2Cn(4,9) = 0.315 ***
AnAr'Cn = 0.125 ***
KL comparison tests
1−2 : p−val = 0.101
1−3 : p−val = 2e−04
2−3 : p−val = 0.0013

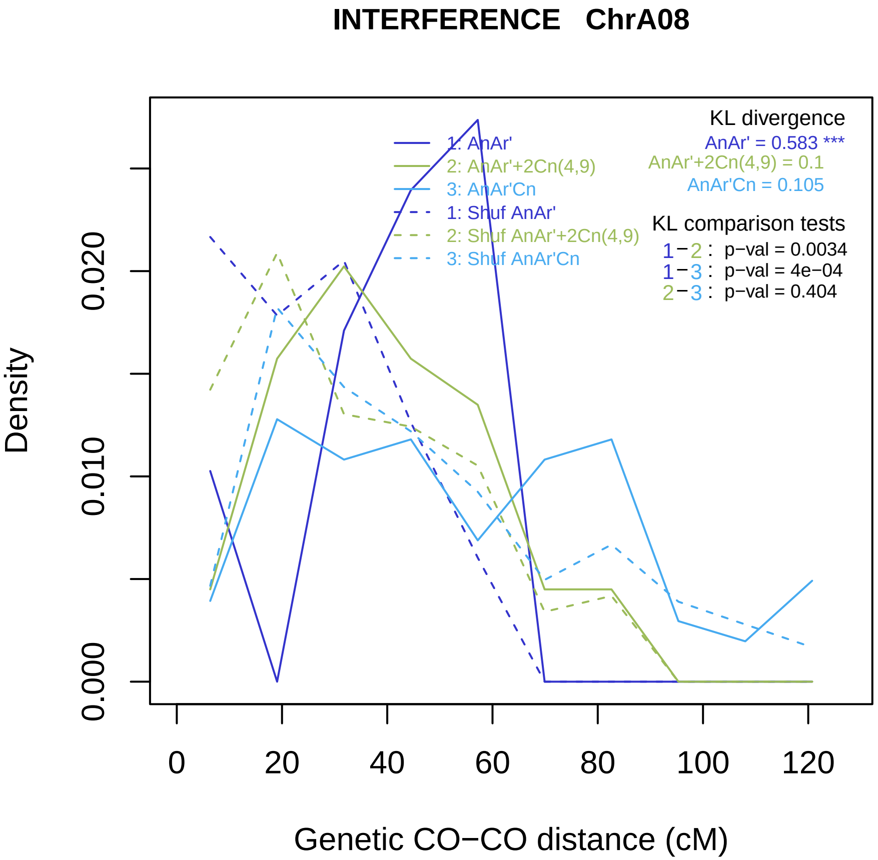
INTERFERENCE ChrA08
KL divergence
AnAr' = 0.583 ***
AnAr'+2Cn(4,9) = 0.1
AnAr'Cn = 0.105
KL comparison tests
1−2 : p−val = 0.0034
1−3 : p−val = 4e−04
2−3 : p−val = 0.404
1: AnAr'
2: AnAr'+2Cn(4,9)
3: AnAr'Cn
1: Shuf AnAr'
2: Shuf AnAr'+2Cn(4,9)
3: Shuf AnAr'Cn
Density
0.000
0.010
0.020
0
20
40
60
80
100
120
Genetic CO−CO distance (cM)

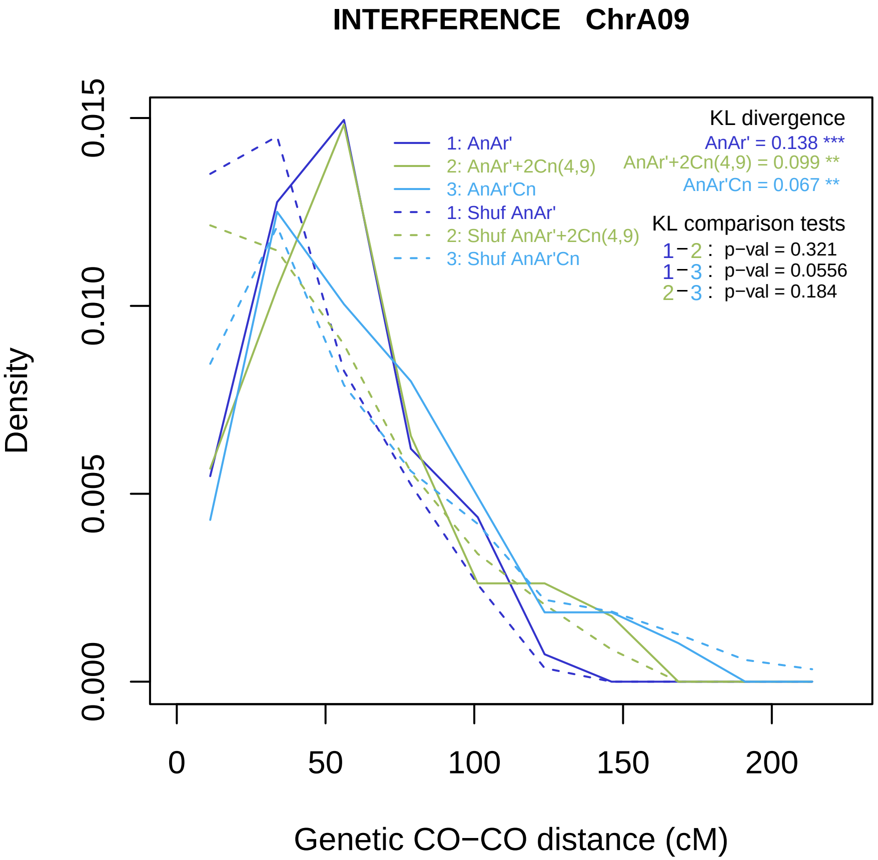

INTERFERENCE ChrA09
Density
0.000
0.005
0.010
0.015
0
50
100
150
200
Genetic CO−CO distance (cM)
1: AnAr'
2: AnAr'+2Cn(4,9)
3: AnAr'Cn
1: Shuf AnAr'
2: Shuf AnAr'+2Cn(4,9)
3: Shuf AnAr'Cn
KL divergence
AnAr' = 0.138 ***
AnAr'+2Cn(4,9) = 0.099 **
AnAr'Cn = 0.067 **
KL comparison tests
1−2 : p−val = 0.321
1−3 : p−val = 0.0556
2−3 : p−val = 0.184

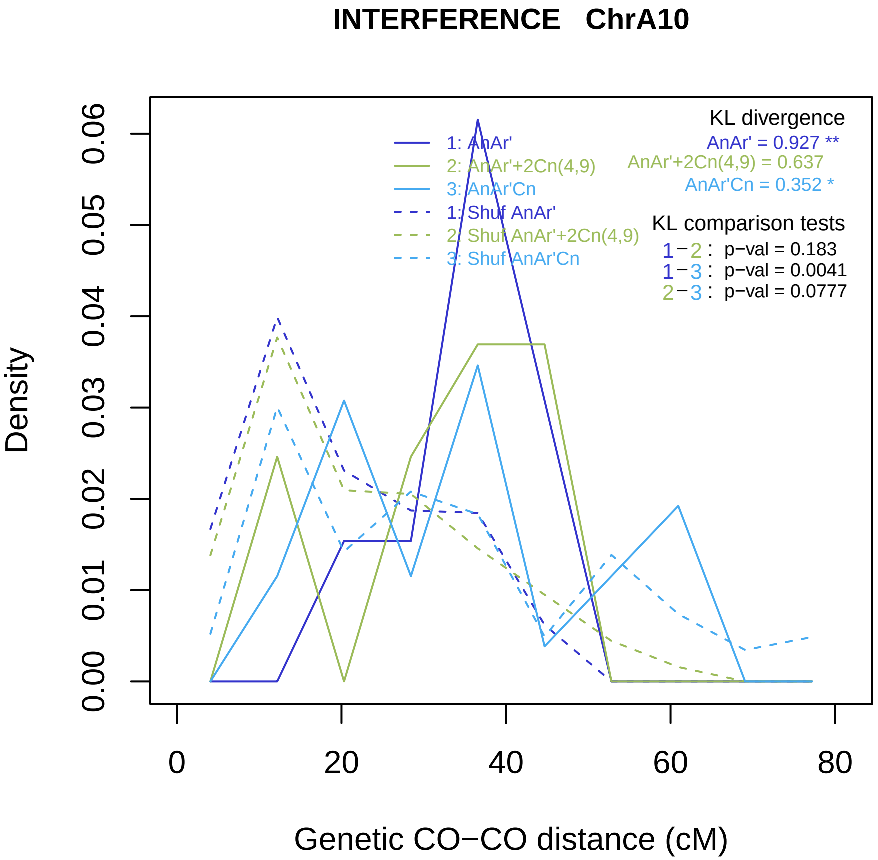

INTERFERENCE ChrA10
KL divergence
AnAr' = 0.927 **
AnAr'+2Cn(4,9) = 0.637
AnAr'Cn = 0.352 *
1: AnAr'
2: AnAr'+2Cn(4,9)
3: AnAr'Cn
1: Shuf AnAr'
2: Shuf AnAr'+2Cn(4,9)
3: Shuf AnAr'Cn
KL comparison tests
1−2 : p−val = 0.183
1−3 : p−val = 0.0041
2−3 : p−val = 0.0777
Density
0.00
0.01
0.02
0.03
0.04
0.05
0.06
0
20
40
60
80
Genetic CO−CO distance (cM)

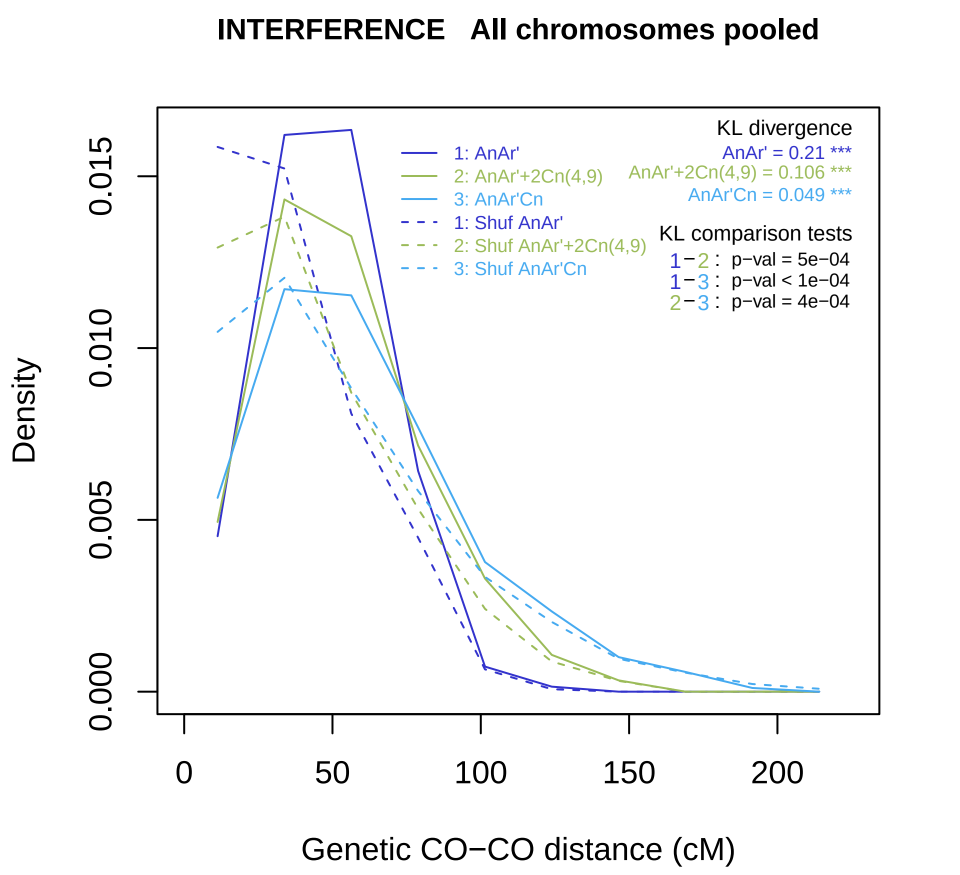
INTERFERENCE All chromosomes pooled
Density
0.000
0.005
0.010
0.015
0
50
100
150
200
Genetic CO−CO distance (cM)
1: AnAr'
2: AnAr'+2Cn(4,9)
3: AnAr'Cn
1: Shuf AnAr'
2: Shuf AnAr'+2Cn(4,9)
3: Shuf AnAr'Cn
KL divergence
AnAr' = 0.21 ***
AnAr'+2Cn(4,9) = 0.106 ***
AnAr'Cn = 0.049 ***
KL comparison tests
1−2 : p−val = 5e−04
1−3 : p−val < 1e−04
2−3 : p−val = 4e−04

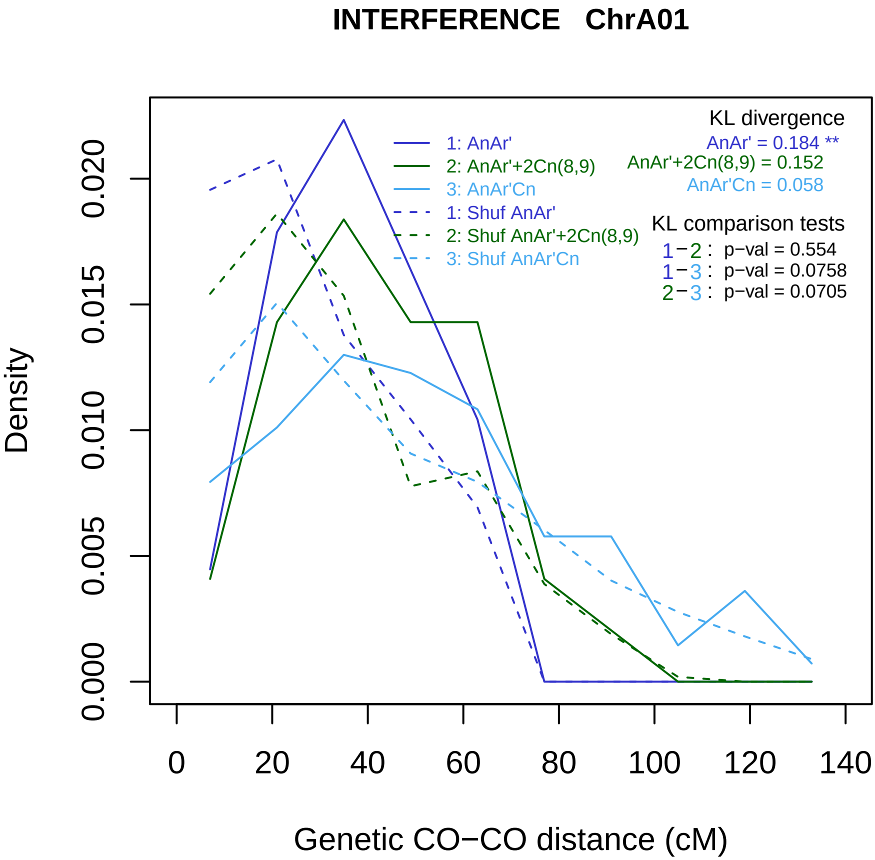
INTERFERENCE ChrA01
KL divergence
AnAr' = 0.184 **
AnAr'+2Cn(8,9) = 0.152
AnAr'Cn = 0.058
1: AnAr'
2: AnAr'+2Cn(8,9)
3: AnAr'Cn
1: Shuf AnAr'
2: Shuf AnAr'+2Cn(8,9)
3: Shuf AnAr'Cn
KL comparison tests
1−2 : p−val = 0.554
1−3 : p−val = 0.0758
2−3 : p−val = 0.0705
Density
0.000
0.005
0.010
0.015
0.020
0
20
40
60
80
100
120
140
Genetic CO−CO distance (cM)

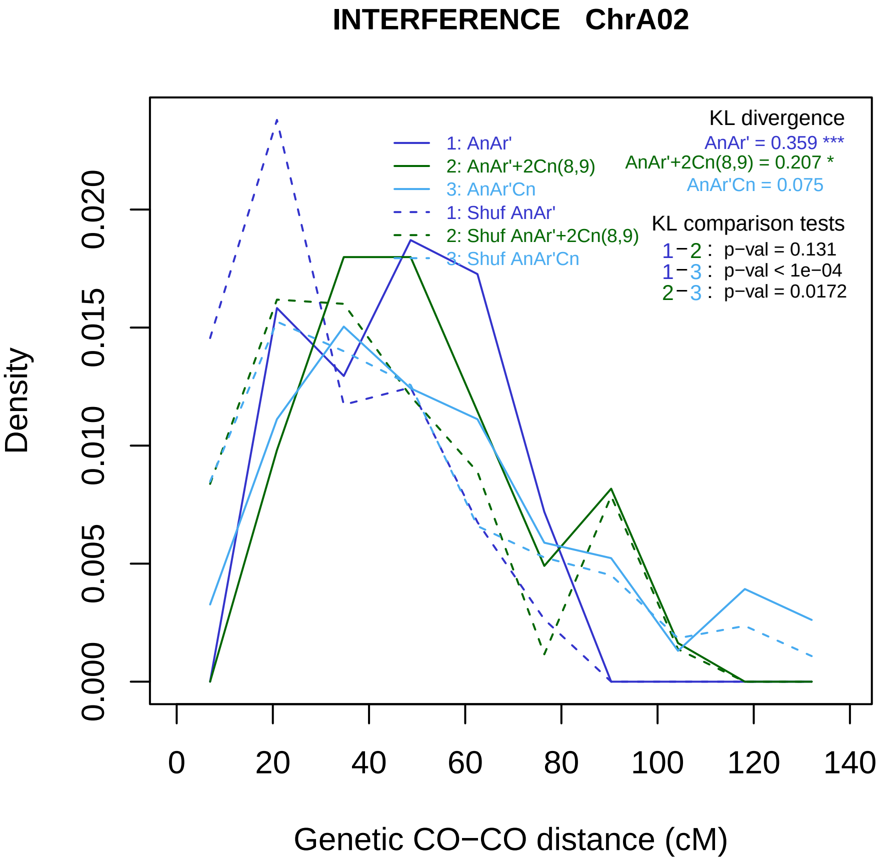
INTERFERENCE ChrA02
KL divergence
AnAr' = 0.359 ***
AnAr'+2Cn(8,9) = 0.207 *
AnAr'Cn = 0.075
1: AnAr'
2: AnAr'+2Cn(8,9)
3: AnAr'Cn
1: Shuf AnAr'
2: Shuf AnAr'+2Cn(8,9)
3: Shuf AnAr'Cn
KL comparison tests
1−2 : p−val = 0.131
1−3 : p−val < 1e−04
2−3 : p−val = 0.0172
Density
0.000
0.005
0.010
0.015
0.020
0
20
40
60
80
100
120
140
Genetic CO−CO distance (cM)

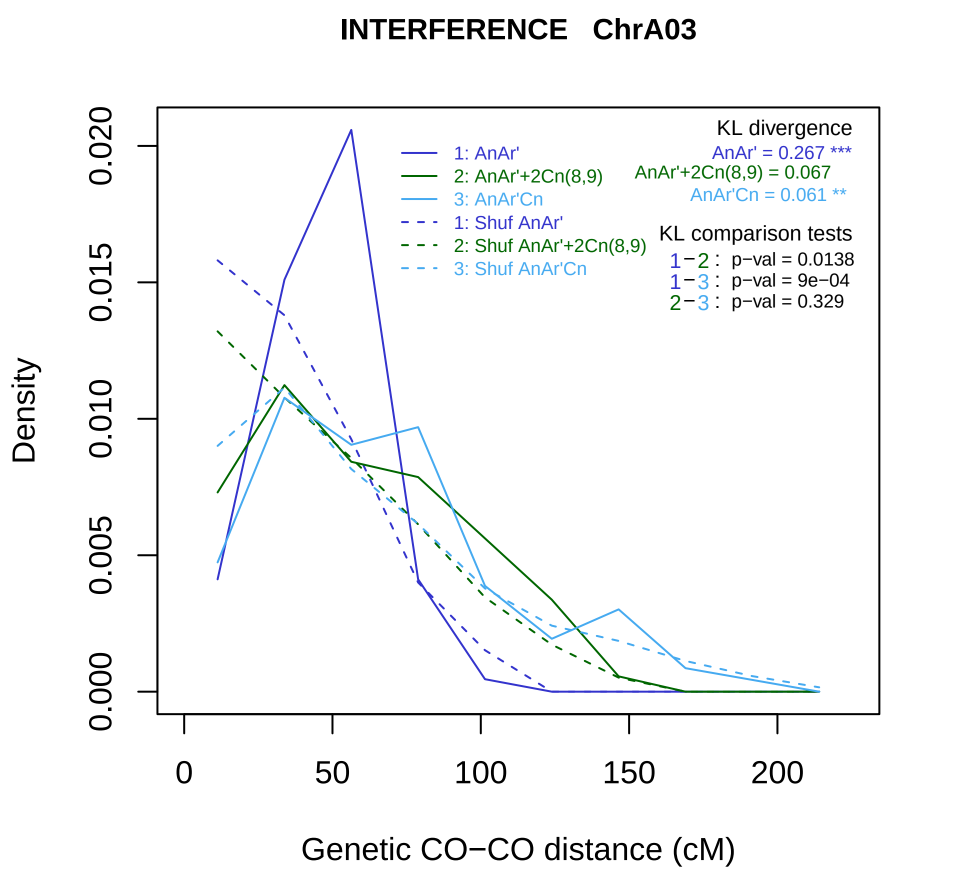
INTERFERENCE ChrA03
Density
Genetic CO−CO distance (cM)
0.000
0.005
0.010
0.015
0.020
0
50
100
150
200
1: AnAr'
2: AnAr'+2Cn(8,9)
3: AnAr'Cn
1: Shuf AnAr'
2: Shuf AnAr'+2Cn(8,9)
3: Shuf AnAr'Cn
KL divergence
AnAr' = 0.267 ***
AnAr'+2Cn(8,9) = 0.067
AnAr'Cn = 0.061 **
KL comparison tests
1−2 : p−val = 0.0138
1−3 : p−val = 9e−04
2−3 : p−val = 0.329

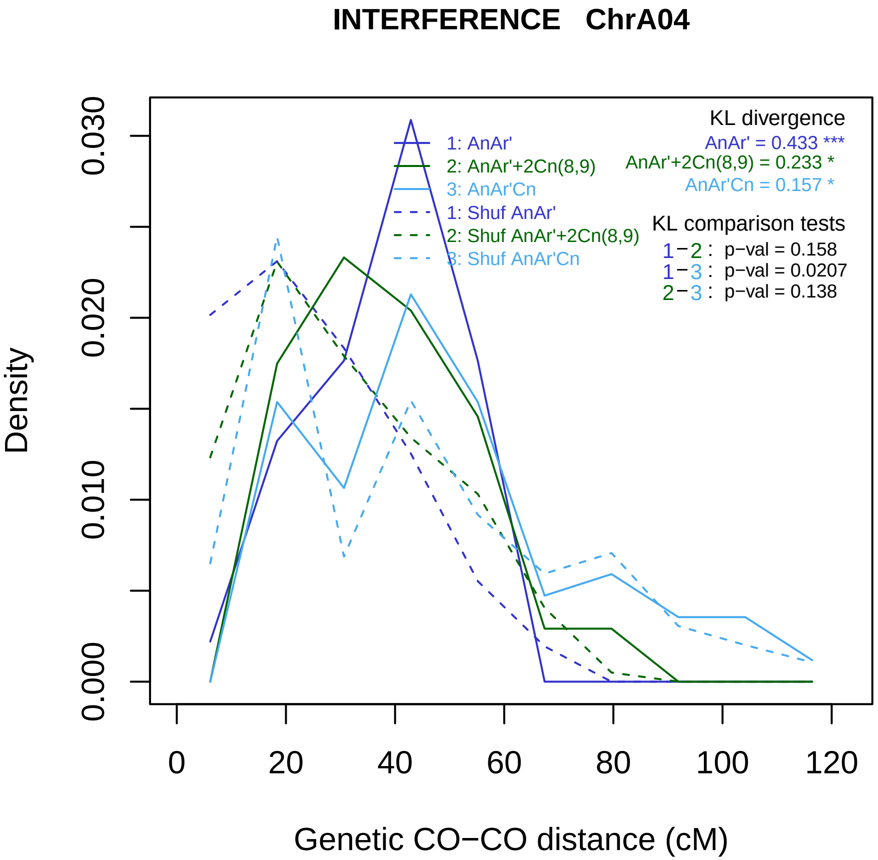

INTERFERENCE ChrA04
KL divergence
AnAr' = 0.433 ***
AnAr'+2Cn(8,9) = 0.233 *
AnAr'Cn = 0.157 *
1: AnAr'
2: AnAr'+2Cn(8,9)
3: AnAr'Cn
1: Shuf AnAr'
2: Shuf AnAr'+2Cn(8,9)
3: Shuf AnAr'Cn
KL comparison tests
1−2 : p−val = 0.158
1−3 : p−val = 0.0207
2−3 : p−val = 0.138
Density
0.000
0.010
0.020
0.030
0
20
40
60
80
100
120
Genetic CO−CO distance (cM)

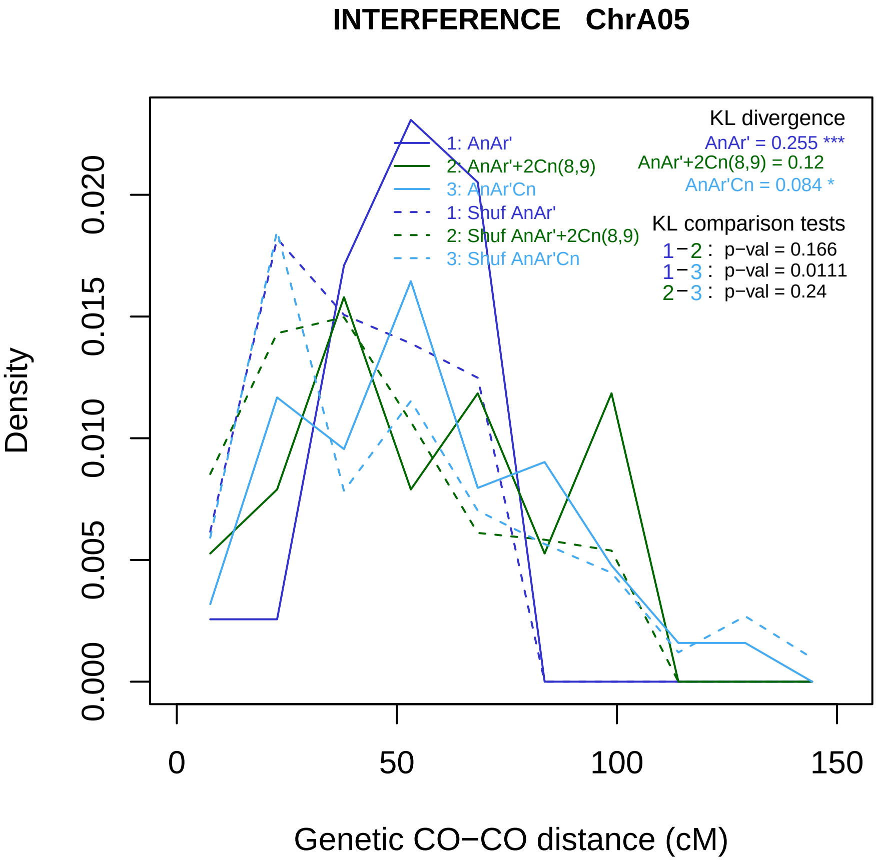
INTERFERENCE ChrA05
Density
0.000
0.005
0.010
0.015
0.020
0
50
100
150
Genetic CO−CO distance (cM)
1: AnAr'
2: AnAr'+2Cn(8,9)
3: AnAr'Cn
1: Shuf AnAr'
2: Shuf AnAr'+2Cn(8,9)
3: Shuf AnAr'Cn
KL divergence
AnAr' = 0.255 ***
AnAr'+2Cn(8,9) = 0.12
AnAr'Cn = 0.084 *
KL comparison tests
1−2 : p−val = 0.166
1−3 : p−val = 0.0111
2−3 : p−val = 0.24

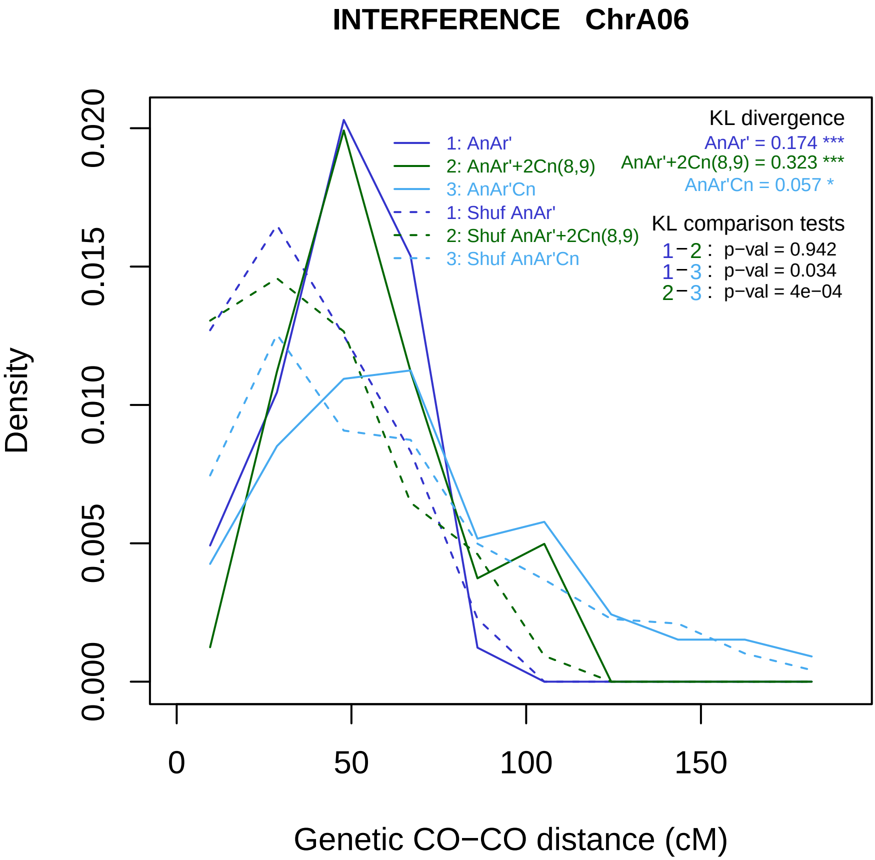
INTERFERENCE ChrA06
1: AnAr'
2: AnAr'+2Cn(8,9)
3: AnAr'Cn
1: Shuf AnAr'
2: Shuf AnAr'+2Cn(8,9)
3: Shuf AnAr'Cn
KL divergence
AnAr' = 0.174 ***
AnAr'+2Cn(8,9) = 0.323 ***
AnAr'Cn = 0.057 *
KL comparison tests
1−2 : p−val = 0.942
1−3 : p−val = 0.034
2−3 : p−val = 4e−04
Density
0.000
0.005
0.010
0.015
0.020
0
50
100
150
Genetic CO−CO distance (cM)

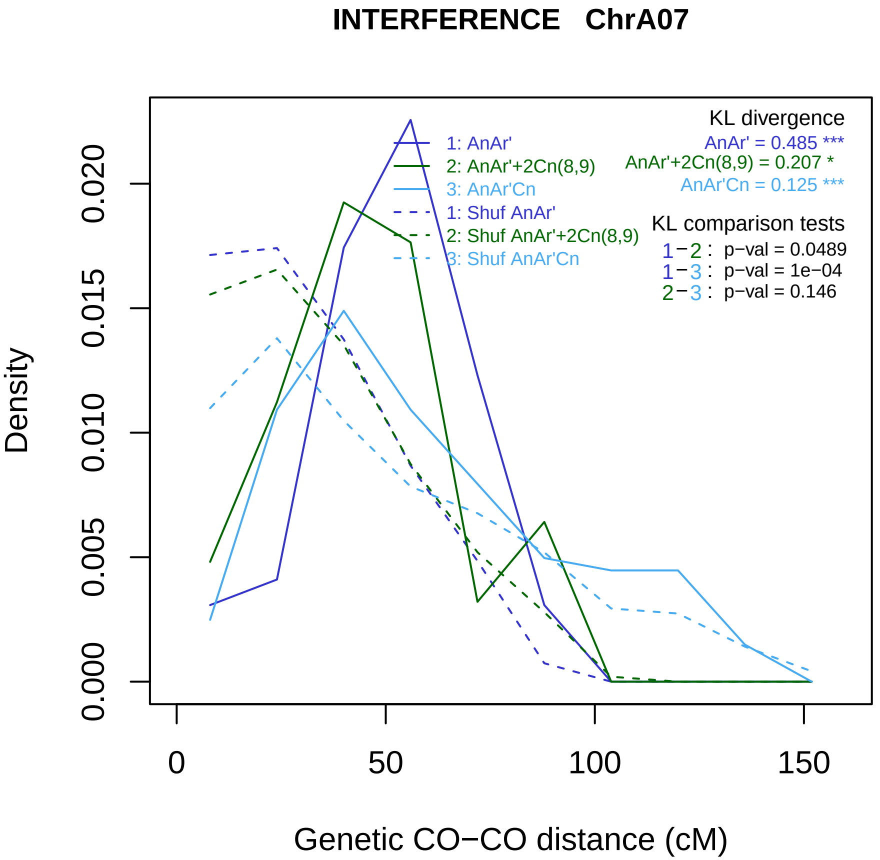

INTERFERENCE ChrA07
KL divergence
AnAr' = 0.485 ***
AnAr'+2Cn(8,9) = 0.207 *
AnAr'Cn = 0.125 ***
1: AnAr'
2: AnAr'+2Cn(8,9)
3: AnAr'Cn
1: Shuf AnAr'
2: Shuf AnAr'+2Cn(8,9)
3: Shuf AnAr'Cn
KL comparison tests
1−2 : p−val = 0.0489
1−3 : p−val = 1e−04
2−3 : p−val = 0.146
Density
0.000
0.005
0.010
0.015
0.020
0
50
100
150
Genetic CO−CO distance (cM)

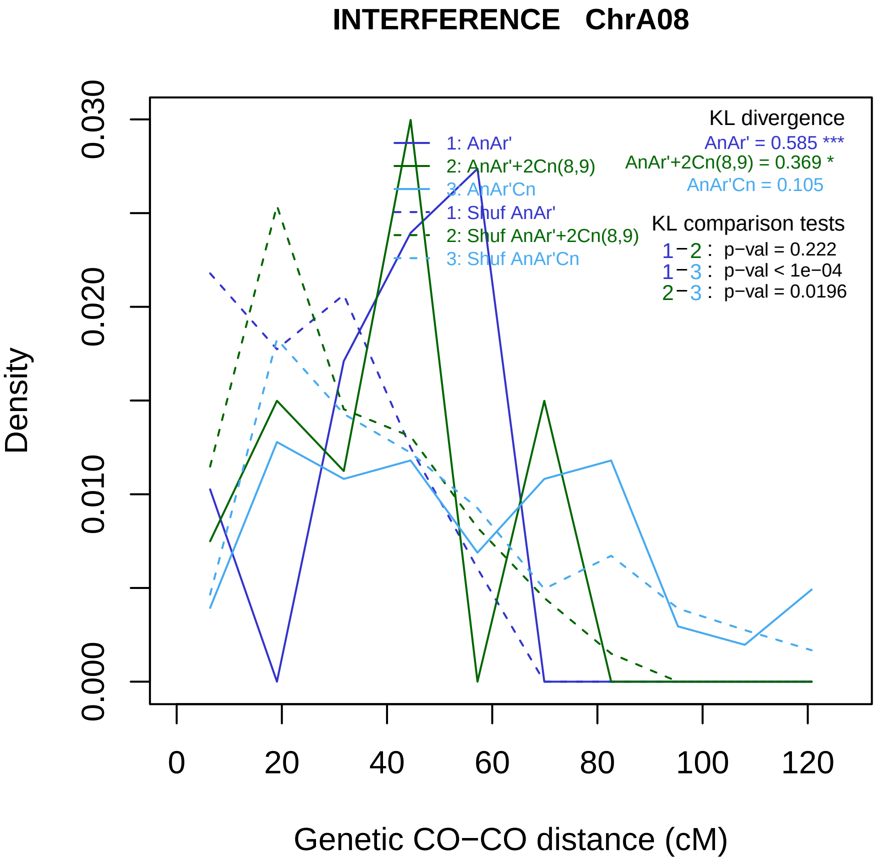
INTERFERENCE ChrA08
Density
0.000
0.010
0.020
0.030
0
20
40
60
80
100
120
Genetic CO−CO distance (cM)
1: AnAr'
2: AnAr'+2Cn(8,9)
3: AnAr'Cn
1: Shuf AnAr'
2: Shuf AnAr'+2Cn(8,9)
3: Shuf AnAr'Cn
KL divergence
AnAr' = 0.585 ***
AnAr'+2Cn(8,9) = 0.369 *
AnAr'Cn = 0.105
KL comparison tests
1−2 : p−val = 0.222
1−3 : p−val < 1e−04
2−3 : p−val = 0.0196

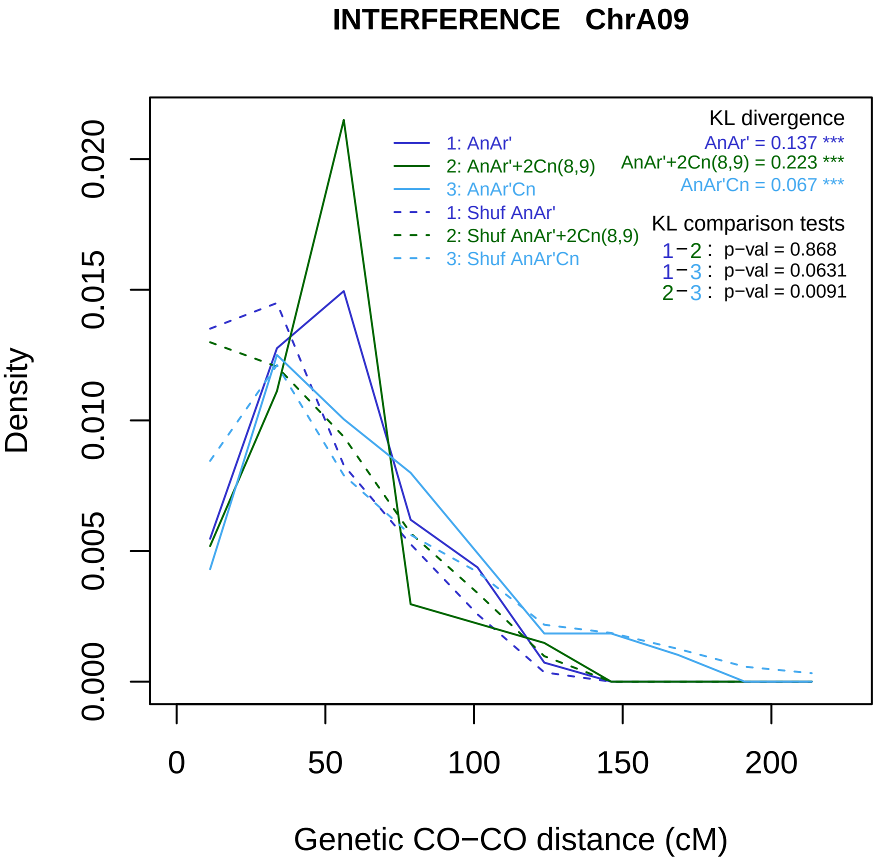

INTERFERENCE ChrA09
Density
Genetic CO−CO distance (cM)
0.000
0.005
0.010
0.015
0.020
0
50
100
150
200
1: AnAr'
2: AnAr'+2Cn(8,9)
3: AnAr'Cn
1: Shuf AnAr'
2: Shuf AnAr'+2Cn(8,9)
3: Shuf AnAr'Cn
KL divergence
AnAr' = 0.137 ***
AnAr'+2Cn(8,9) = 0.223 ***
AnAr'Cn = 0.067 ***
KL comparison tests
1−2 : p−val = 0.868
1−3 : p−val = 0.0631
2−3 : p−val = 0.0091

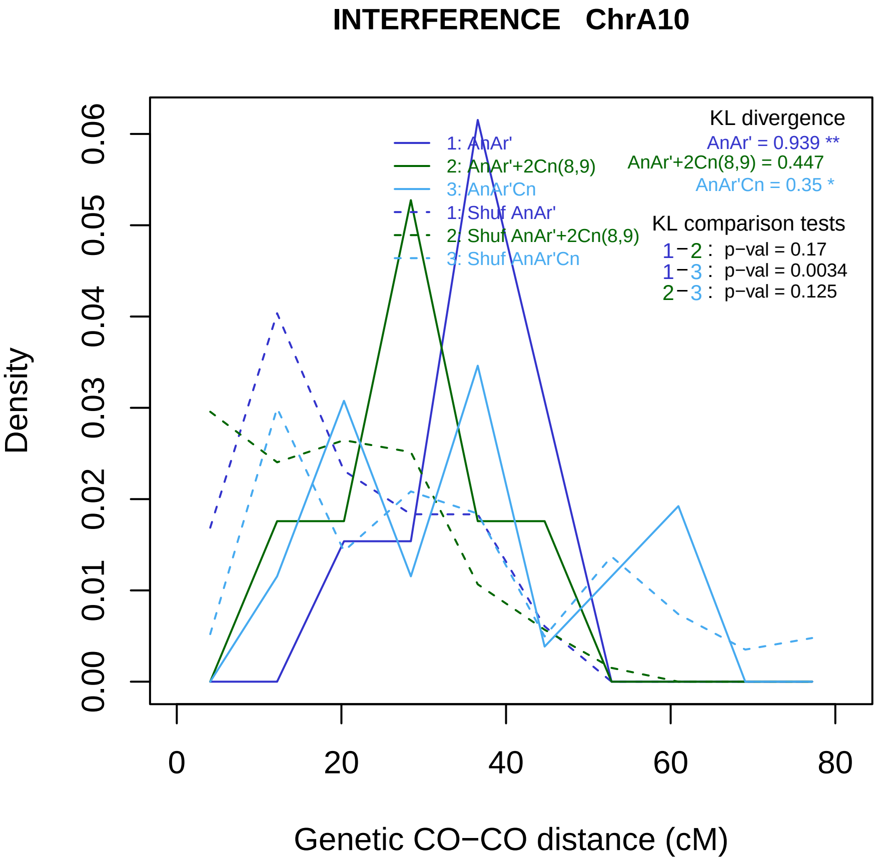
INTERFERENCE ChrA10
Density
0.00
0.01
0.02
0.03
0.04
0.05
0.06
0
20
40
60
80
Genetic CO−CO distance (cM)
1: AnAr'
2: AnAr'+2Cn(8,9)
3: AnAr'Cn
1: Shuf AnAr'
2: Shuf AnAr'+2Cn(8,9)
3: Shuf AnAr'Cn
KL divergence
AnAr' = 0.939 **
AnAr'+2Cn(8,9) = 0.447
AnAr'Cn = 0.35 *
KL comparison tests
1−2 : p−val = 0.17
1−3 : p−val = 0.0034
2−3 : p−val = 0.125

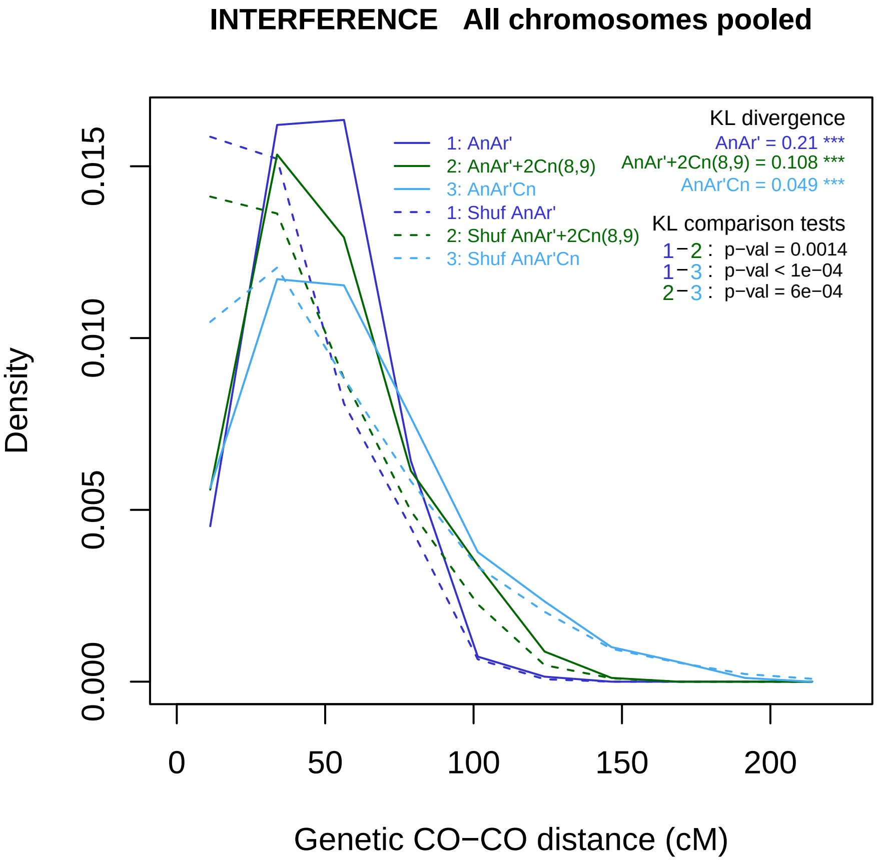

INTERFERENCE All chromosomes pooled
Density
0.000
0.005
0.010
0.015
0
50
100
150
200
Genetic CO−CO distance (cM)
1: AnAr'
2: AnAr'+2Cn(8,9)
3: AnAr'Cn
1: Shuf AnAr'
2: Shuf AnAr'+2Cn(8,9)
3: Shuf AnAr'Cn
KL divergence
AnAr' = 0.21 ***
AnAr'+2Cn(8,9) = 0.108 ***
AnAr'Cn = 0.049 ***
KL comparison tests
1−2 : p−val = 0.0014
1−3 : p−val < 1e−04
2−3 : p−val = 6e−04

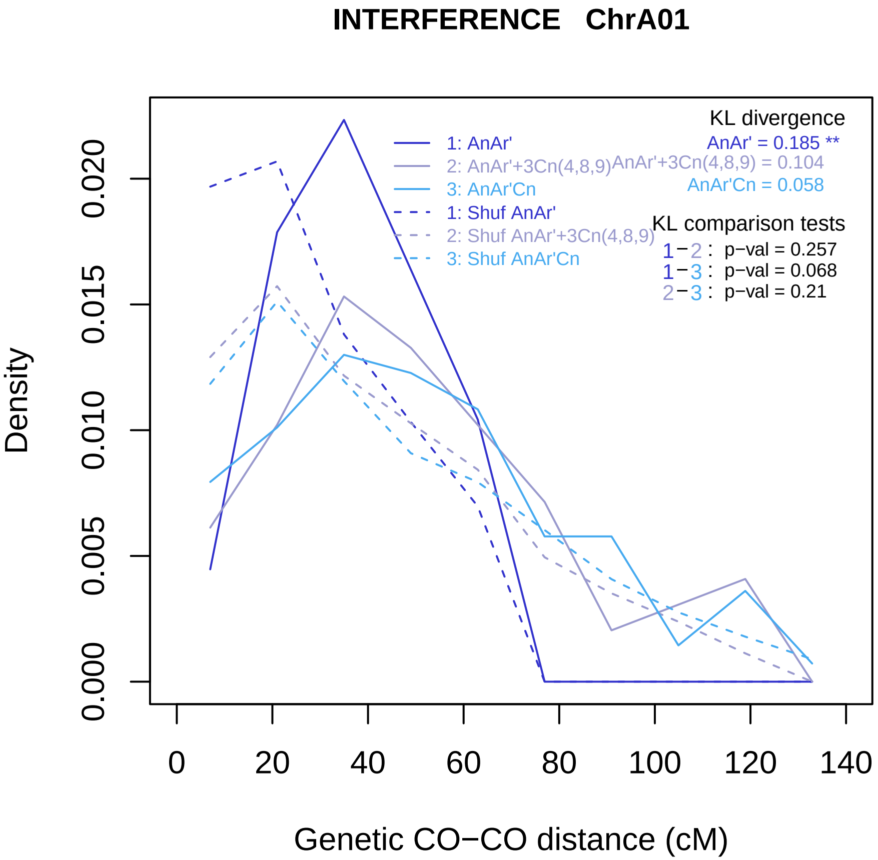

INTERFERENCE ChrA01
KL divergence
AnAr' = 0.185 **
AnAr'+3Cn(4,8,9) = 0.104
AnAr'Cn = 0.058
1: AnAr'
2: AnAr'+3Cn(4,8,9)
3: AnAr'Cn
1: Shuf AnAr'
2: Shuf AnAr'+3Cn(4,8,9)
3: Shuf AnAr'Cn
KL comparison tests
1−2 : p−val = 0.257
1−3 : p−val = 0.068
2−3 : p−val = 0.21
Density
0.000
0.005
0.010
0.015
0.020
0
20
40
60
80
100
120
140
Genetic CO−CO distance (cM)

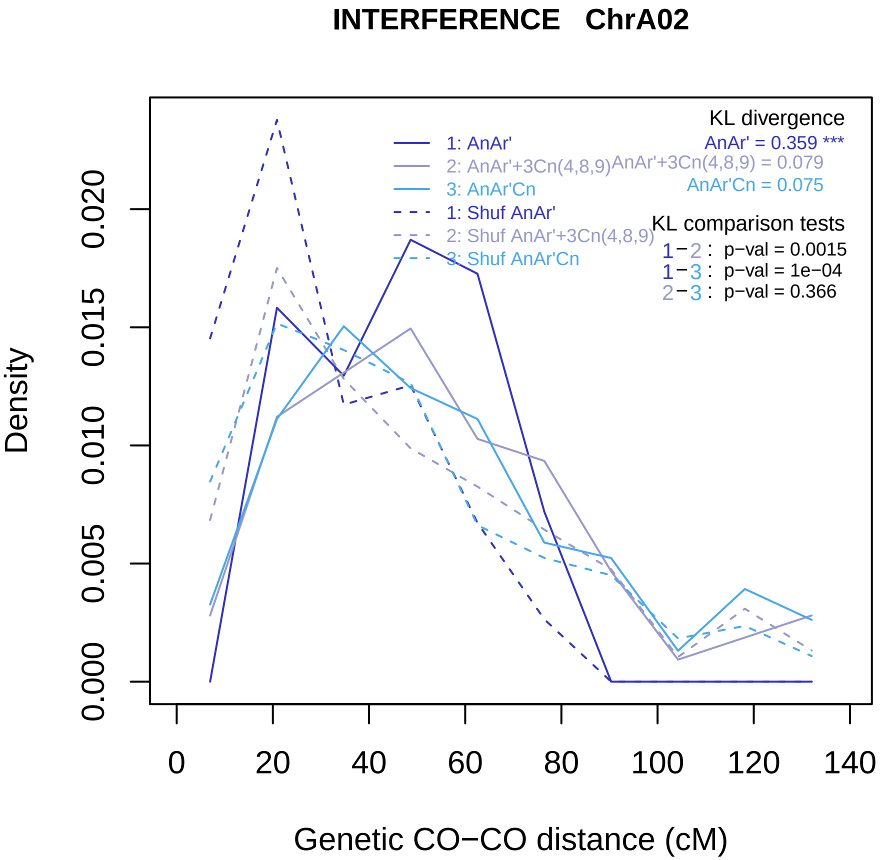

INTERFERENCE ChrA02
KL divergence
AnAr' = 0.359 ***
AnAr'+3Cn(4,8,9) = 0.079
AnAr'Cn = 0.075
1: AnAr'
2: AnAr'+3Cn(4,8,9)
3: AnAr'Cn
1: Shuf AnAr'
2: Shuf AnAr'+3Cn(4,8,9)
3: Shuf AnAr'Cn
KL comparison tests
1−2 : p−val = 0.0015
1−3 : p−val = 1e−04
2−3 : p−val = 0.366
Density
0.000
0.005
0.010
0.015
0.020
0
20
40
60
80
100
120
140
Genetic CO−CO distance (cM)

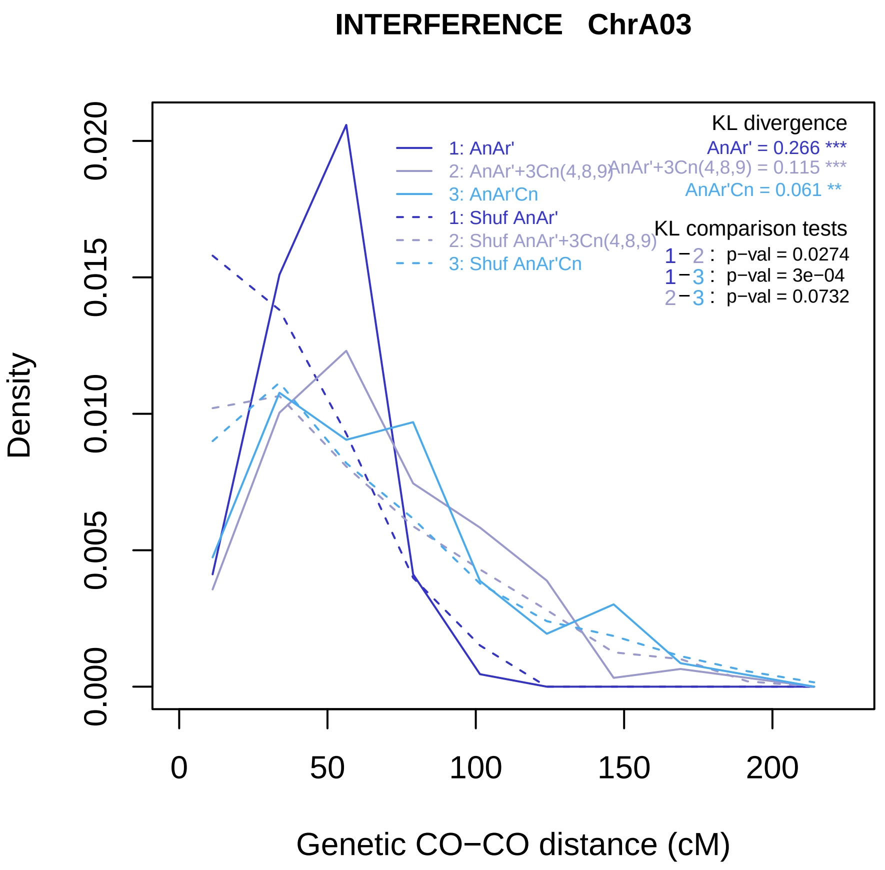

INTERFERENCE ChrA03
Density
Genetic CO−CO distance (cM)
0.000
0.005
0.010
0.015
0.020
0
50
100
150
200
1: AnAr'
2: AnAr'+3Cn(4,8,9)
3: AnAr'Cn
1: Shuf AnAr'
2: Shuf AnAr'+3Cn(4,8,9)
3: Shuf AnAr'Cn
KL divergence
AnAr' = 0.266 ***
AnAr'+3Cn(4,8,9) = 0.115 ***
AnAr'Cn = 0.061 **
KL comparison tests
1−2 : p−val = 0.0274
1−3 : p−val = 3e−04
2−3 : p−val = 0.0732

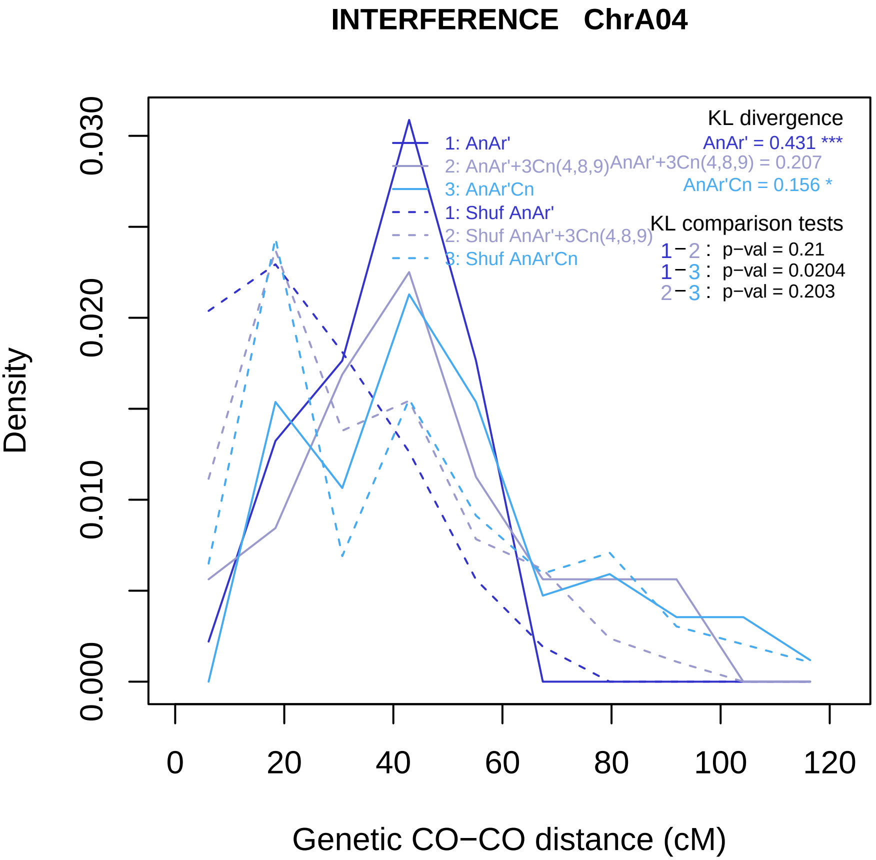

INTERFERENCE ChrA04
KL divergence
AnAr' = 0.431 ***
AnAr'+3Cn(4,8,9) = 0.207
AnAr'Cn = 0.156 *
1: AnAr'
2: AnAr'+3Cn(4,8,9)
3: AnAr'Cn
1: Shuf AnAr'
2: Shuf AnAr'+3Cn(4,8,9)
3: Shuf AnAr'Cn
KL comparison tests
1−2 : p−val = 0.21
1−3 : p−val = 0.0204
2−3 : p−val = 0.203
Density
0.000
0.010
0.020
0.030
0
20
40
60
80
100
120
Genetic CO−CO distance (cM)

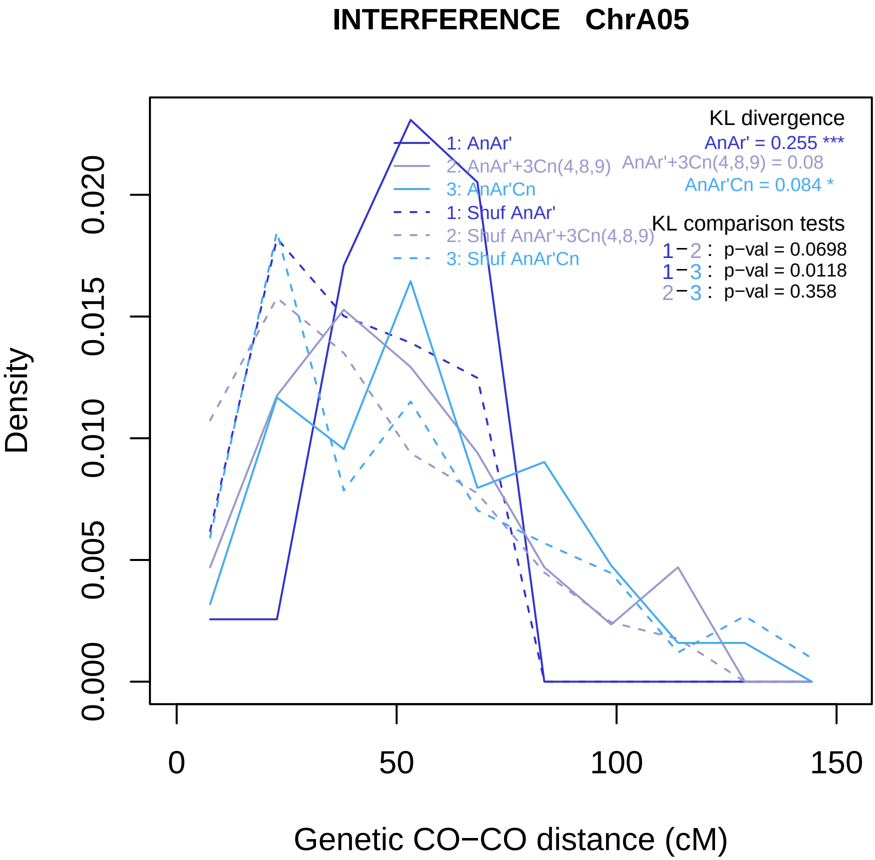
INTERFERENCE ChrA05
Density
0.000
0.005
0.010
0.015
0.020
0
50
100
150
Genetic CO−CO distance (cM)
1: AnAr'
2: AnAr'+3Cn(4,8,9)
3: AnAr'Cn
1: Shuf AnAr'
2: Shuf AnAr'+3Cn(4,8,9)
3: Shuf AnAr'Cn
KL divergence
AnAr' = 0.255 ***
AnAr'+3Cn(4,8,9) = 0.08
AnAr'Cn = 0.084 *
KL comparison tests
1−2 : p−val = 0.0698
1−3 : p−val = 0.0118
2−3 : p−val = 0.358

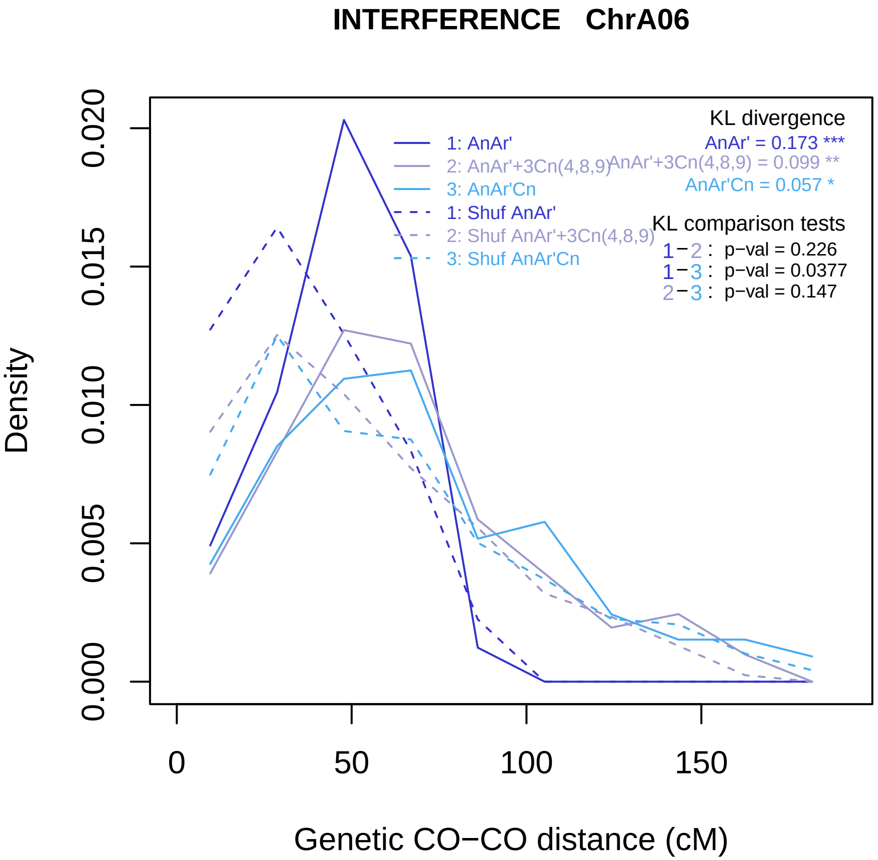

INTERFERENCE ChrA06
Density
0.000
0.005
0.010
0.015
0.020
0
50
100
150
Genetic CO−CO distance (cM)
1: AnAr'
2: AnAr'+3Cn(4,8,9)
3: AnAr'Cn
1: Shuf AnAr'
2: Shuf AnAr'+3Cn(4,8,9)
3: Shuf AnAr'Cn
KL divergence
AnAr' = 0.173 ***
AnAr'+3Cn(4,8,9) = 0.099 **
AnAr'Cn = 0.057 *
KL comparison tests
1−2 : p−val = 0.226
1−3 : p−val = 0.0377
2−3 : p−val = 0.147

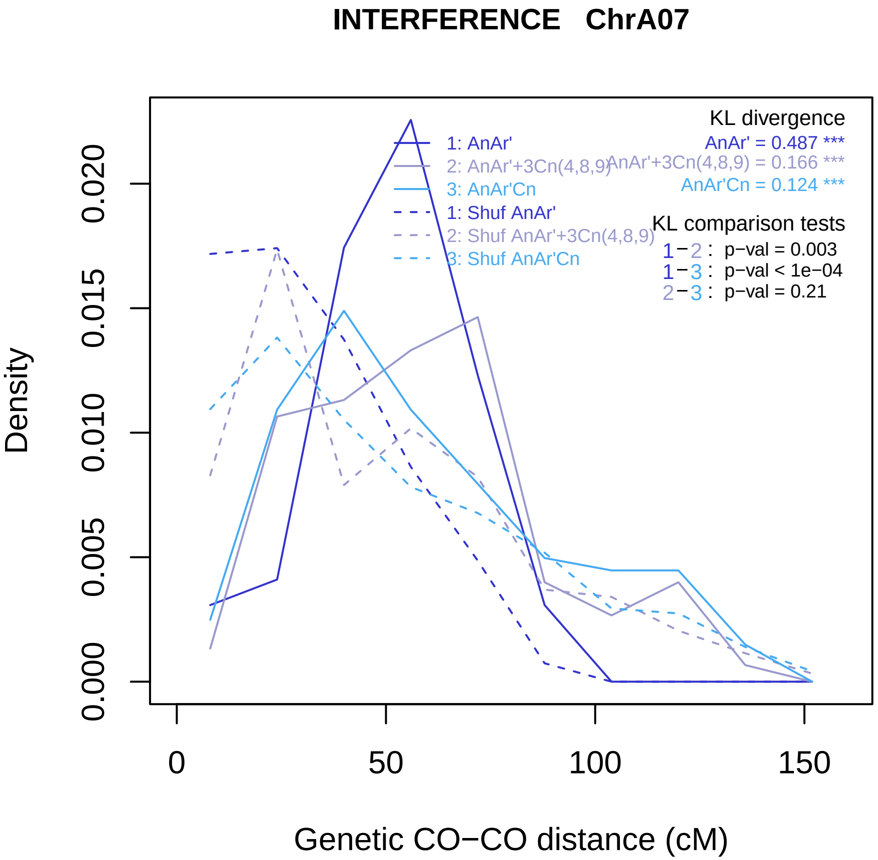

INTERFERENCE ChrA07
KL divergence
AnAr' = 0.487 ***
AnAr'+3Cn(4,8,9) = 0.166 ***
AnAr'Cn = 0.124 ***
1: AnAr'
2: AnAr'+3Cn(4,8,9)
3: AnAr'Cn
1: Shuf AnAr'
2: Shuf AnAr'+3Cn(4,8,9)
3: Shuf AnAr'Cn
KL comparison tests
1−2 : p−val = 0.003
1−3 : p−val < 1e−04
2−3 : p−val = 0.21
Density
0.000
0.005
0.010
0.015
0.020
0
50
100
150
Genetic CO−CO distance (cM)

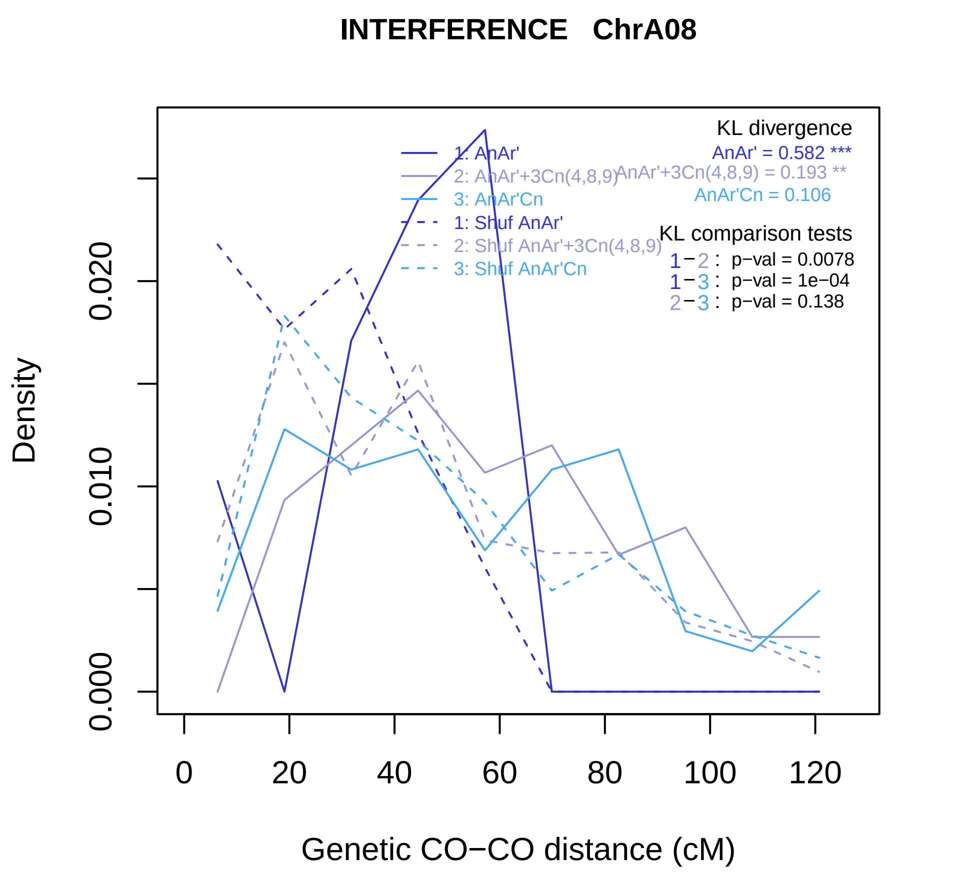
INTERFERENCE ChrA08
KL divergence
AnAr' = 0.582 ***
AnAr'+3Cn(4,8,9) = 0.193 **
AnAr'Cn = 0.106
1: AnAr'
2: AnAr'+3Cn(4,8,9)
3: AnAr'Cn
1: Shuf AnAr'
2: Shuf AnAr'+3Cn(4,8,9)
3: Shuf AnAr'Cn
KL comparison tests
1−2 : p−val = 0.0078
1−3 : p−val = 1e−04
2−3 : p−val = 0.138
Density
0.000
0.010
0.020
0
20
40
60
80
100
120
Genetic CO−CO distance (cM)

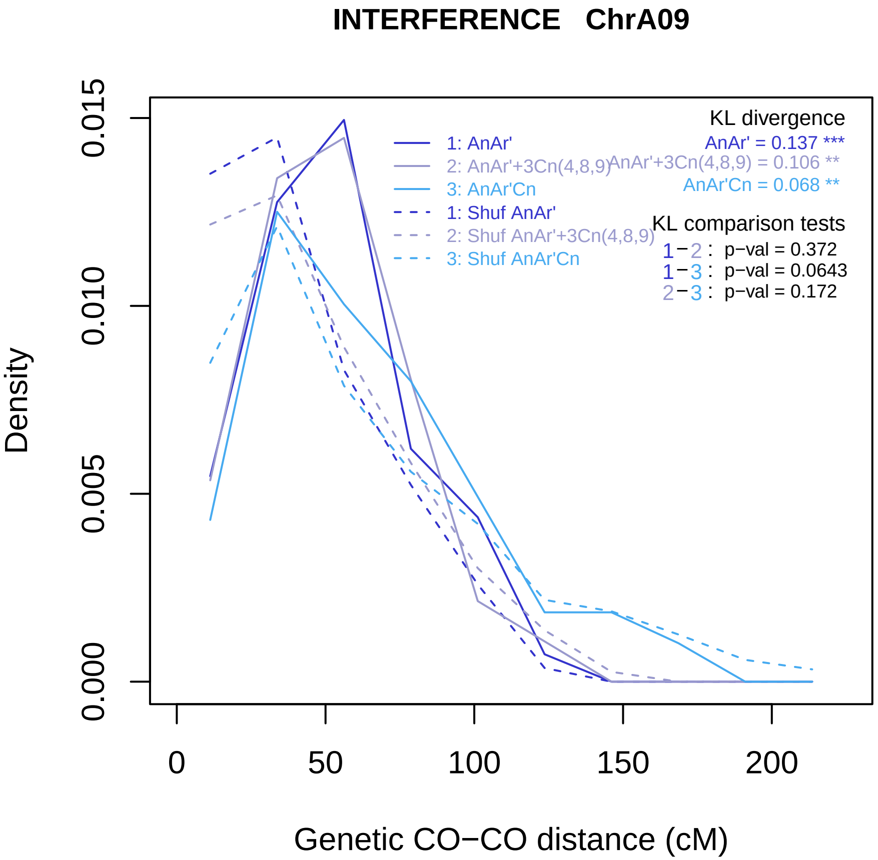

INTERFERENCE ChrA09
Density
0.000
0.005
0.010
0.015
0
50
100
150
200
Genetic CO−CO distance (cM)
1: AnAr'
2: AnAr'+3Cn(4,8,9)
3: AnAr'Cn
1: Shuf AnAr'
2: Shuf AnAr'+3Cn(4,8,9)
3: Shuf AnAr'Cn
KL divergence
AnAr' = 0.137 ***
AnAr'+3Cn(4,8,9) = 0.106 **
AnAr'Cn = 0.068 **
KL comparison tests
1−2 : p−val = 0.372
1−3 : p−val = 0.0643
2−3 : p−val = 0.172

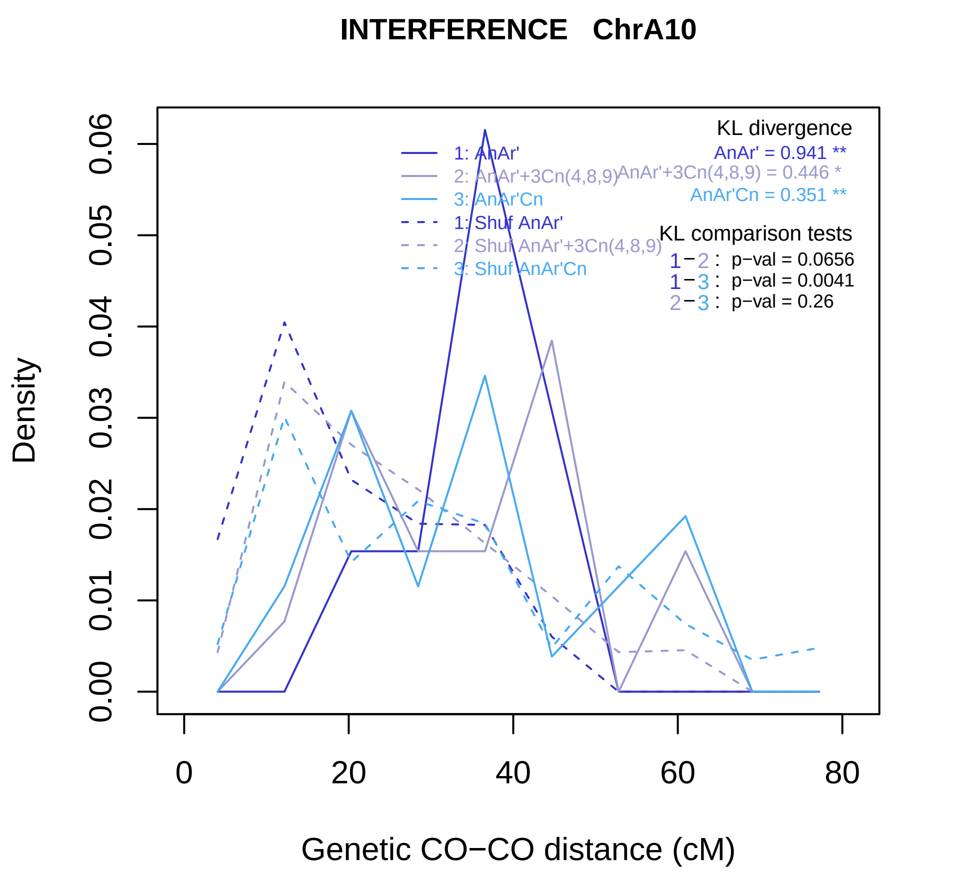
INTERFERENCE ChrA10
KL divergence
AnAr' = 0.941 **
AnAr'+3Cn(4,8,9) = 0.446 *
AnAr'Cn = 0.351 **
1: AnAr'
2: AnAr'+3Cn(4,8,9)
3: AnAr'Cn
1: Shuf AnAr'
2: Shuf AnAr'+3Cn(4,8,9)
3: Shuf AnAr'Cn
KL comparison tests
1−2 : p−val = 0.0656
1−3 : p−val = 0.0041
2−3 : p−val = 0.26
Density
0.00
0.01
0.02
0.03
0.04
0.05
0.06
0
20
40
60
80
Genetic CO−CO distance (cM)

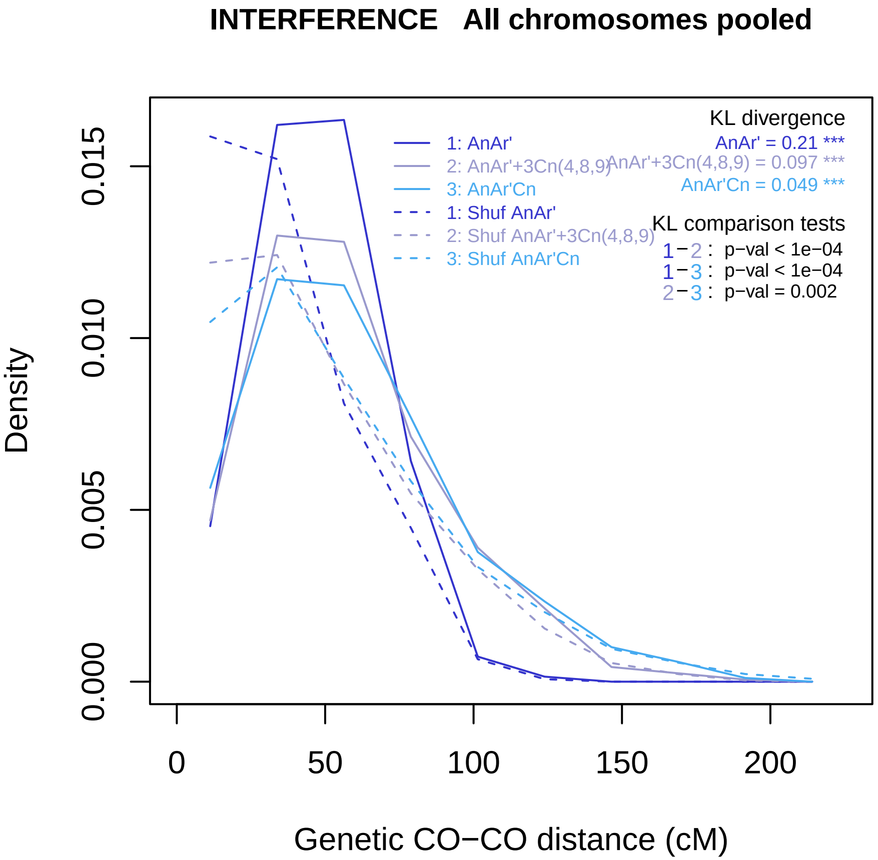
INTERFERENCE All chromosomes pooled
Density
Genetic CO−CO distance (cM)
0.000
0.005
0.010
0.015
0
50
100
150
200
1: AnAr'
2: AnAr'+3Cn(4,8,9)
3: AnAr'Cn
1: Shuf AnAr'
2: Shuf AnAr'+3Cn(4,8,9)
3: Shuf AnAr'Cn
KL divergence
AnAr' = 0.21 ***
AnAr'+3Cn(4,8,9) = 0.097 ***
AnAr'Cn = 0.049 ***
KL comparison tests
1−2 : p−val < 1e−04
1−3 : p−val < 1e−04
2−3 : p−val = 0.002
