## Supplemental Figure 7 for "Genomic divergence shaped the genetic regulation of meiotic homologous recombination in *Brassica* allopolyploids"

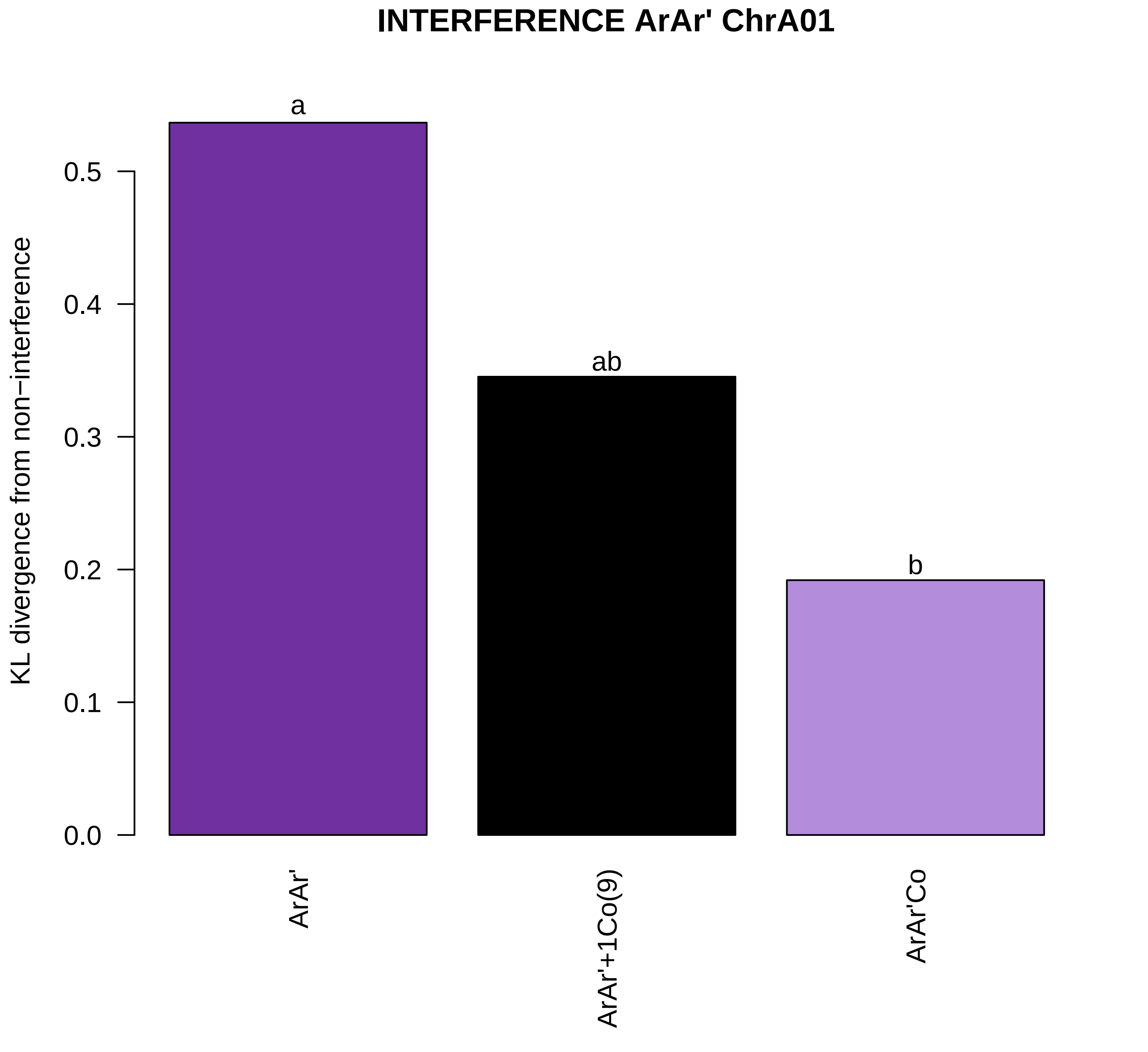
INTERFERENCE ArAr' ChrA01
KL divergence from non−interference
0.0
0.1
0.2
0.3
0.4
0.5
a
ab
b
ArAr'
ArAr'+1Co(9)
ArAr'Co

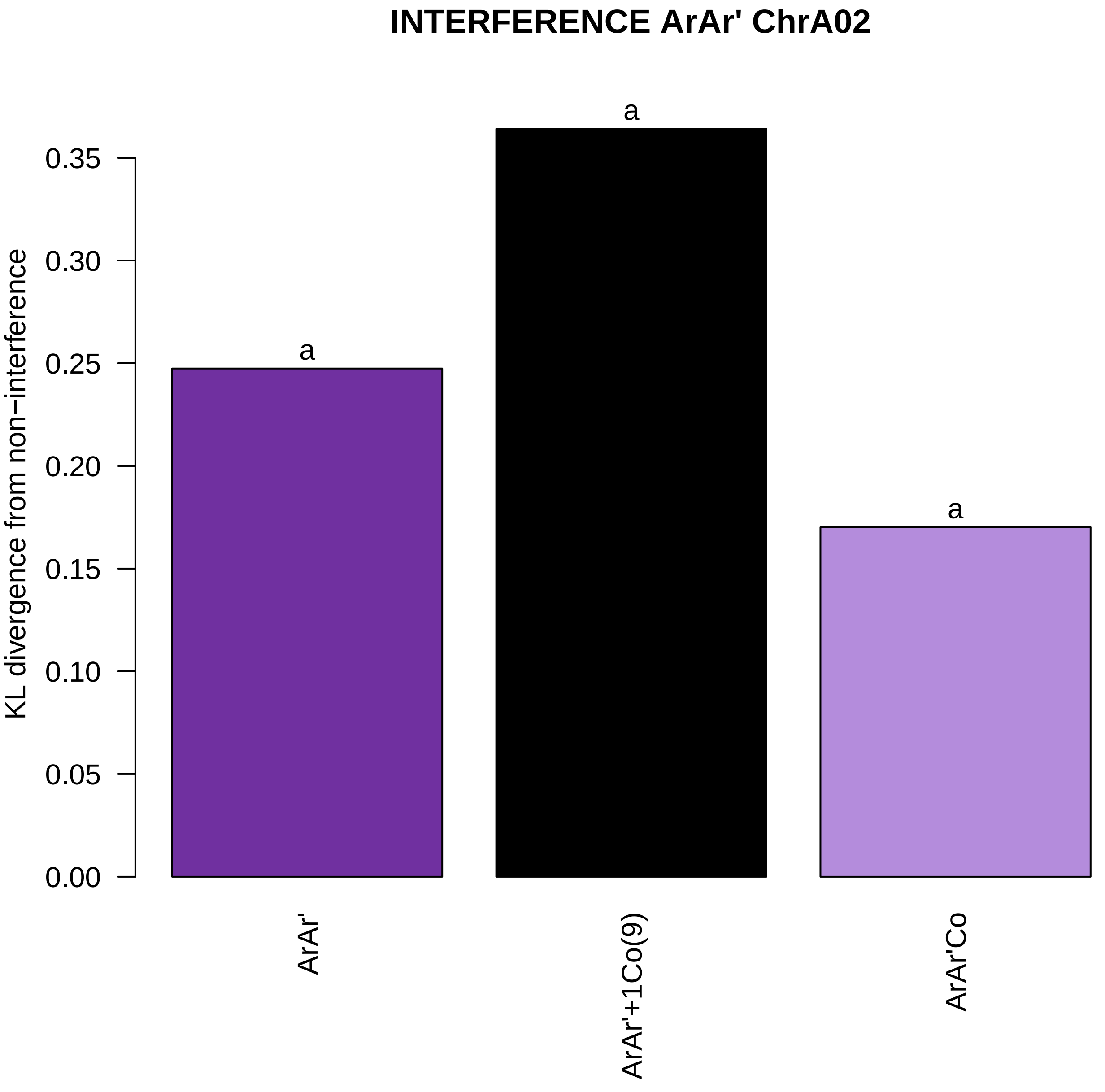

INTERFERENCE ArAr' ChrA02
KL divergence from non−interference
0.00
0.05
0.10
0.15
0.20
0.25
0.30
0.35
a
a
a
ArAr'
ArAr'+1Co(9)
ArAr'Co

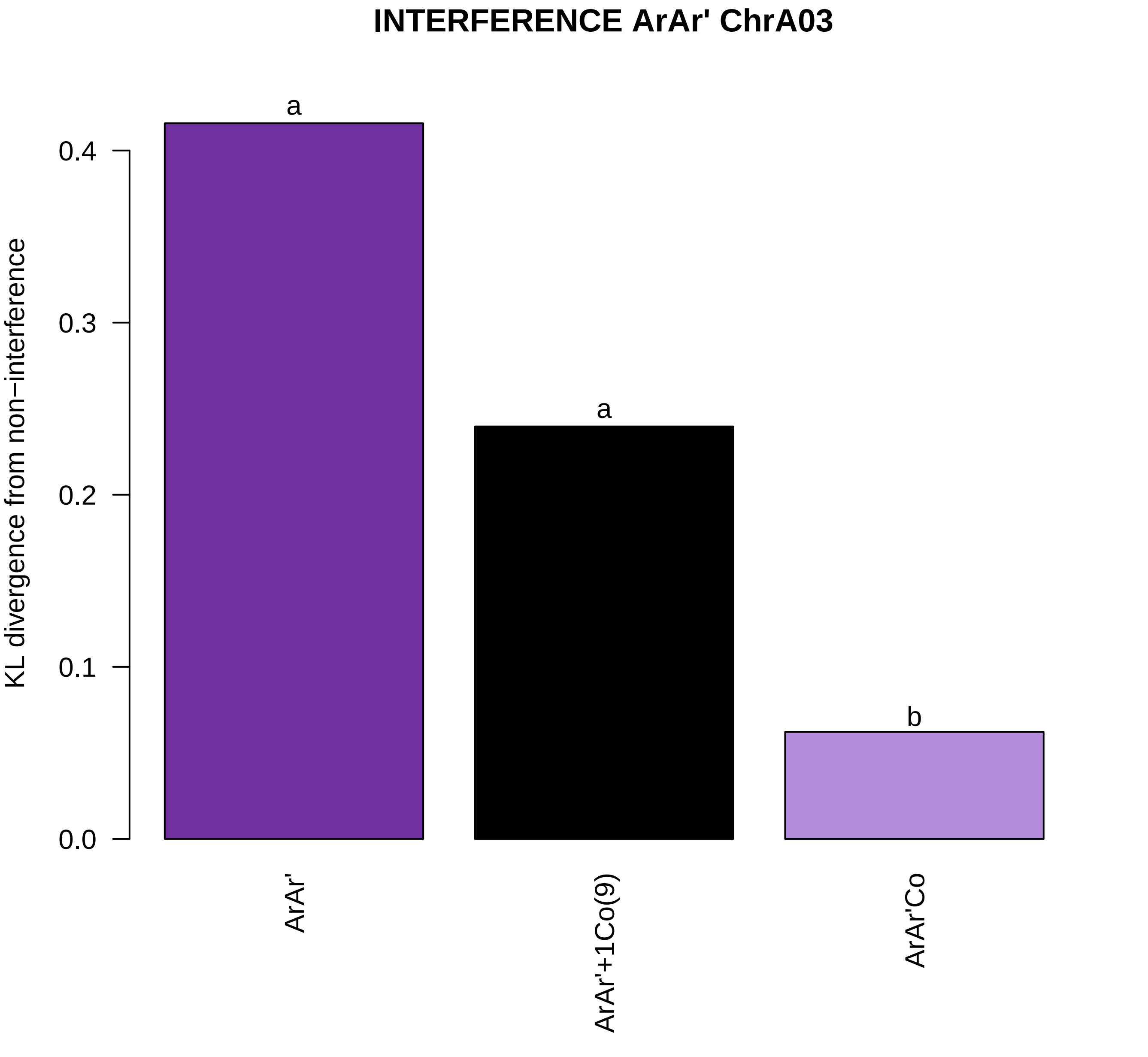
INTERFERENCE ArAr' ChrA03
KL divergence from non−interference
0.0
0.1
0.2
0.3
0.4
a
a
b
ArAr'
ArAr'+1Co(9)
ArAr'Co

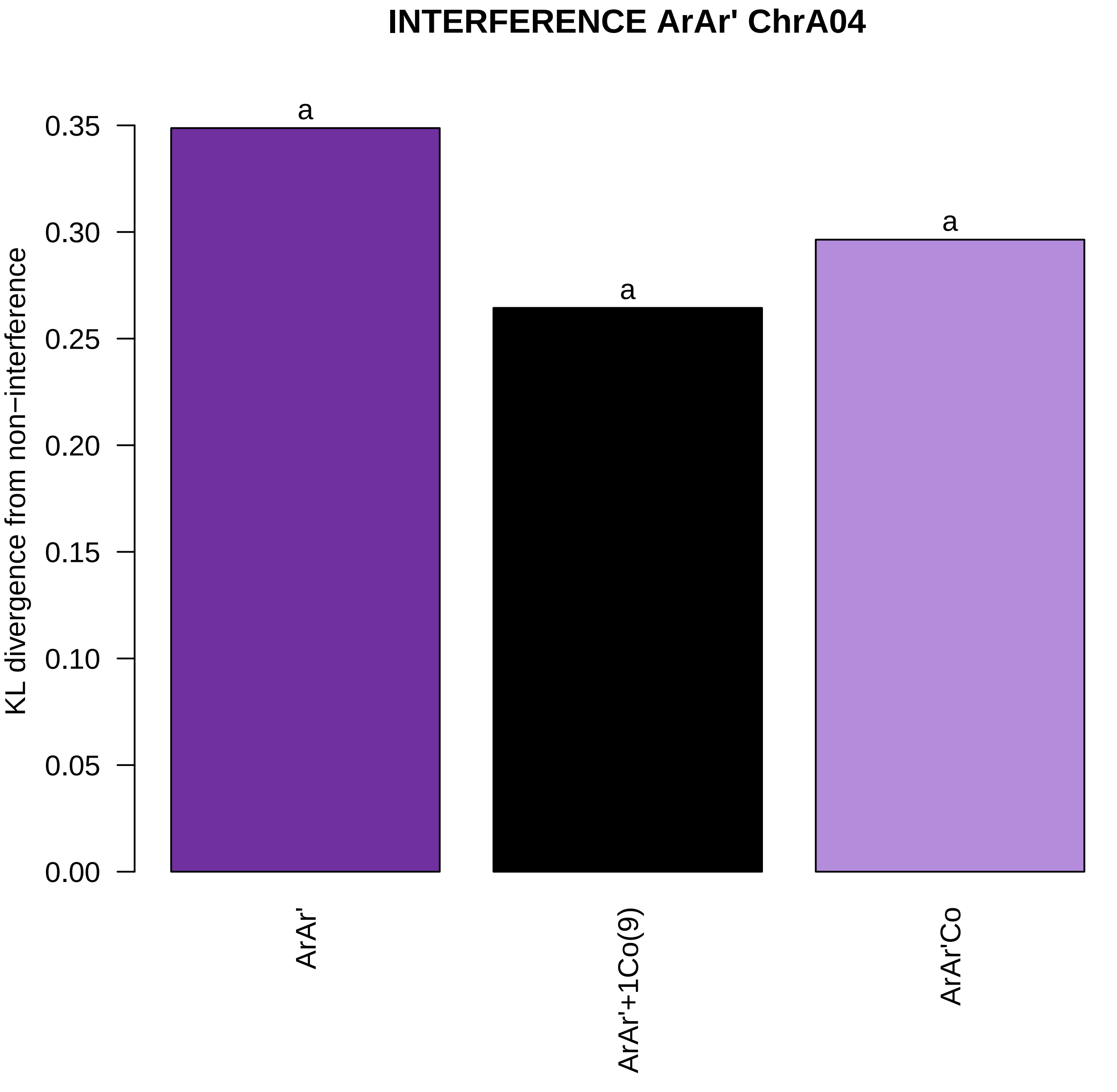
INTERFERENCE ArAr' ChrA04
KL divergence from non−interference
0.35
0.30
0.25
0.20
0.15
0.10
0.05
0.00
a
a
a
ArAr'
ArAr'+1Co(9)
ArAr'Co

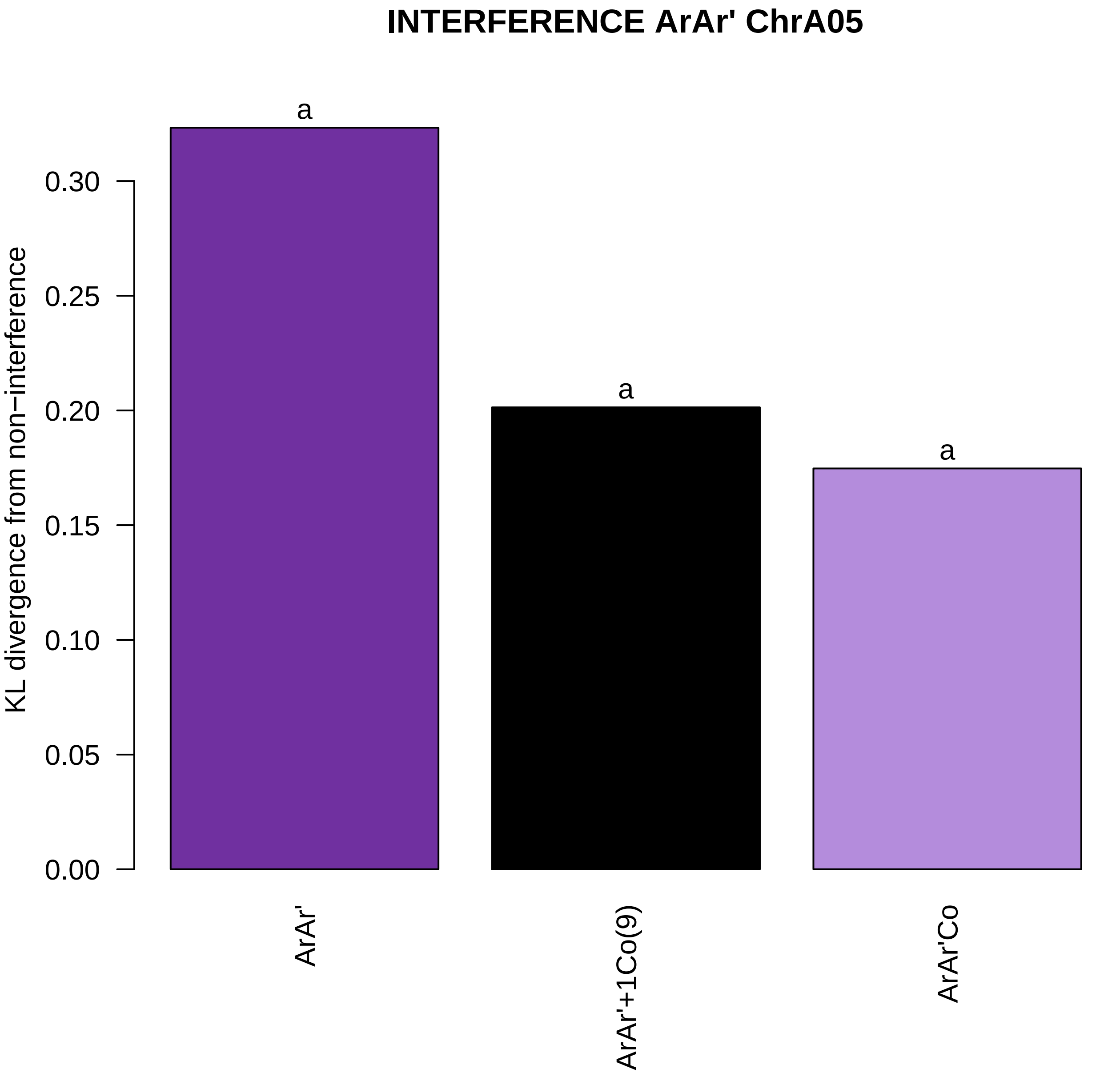
INTERFERENCE ArAr' ChrA05
KL divergence from non−interference
0.00
0.05
0.10
0.15
0.20
0.25
0.30
a
a
a
ArAr'
ArAr'+1Co(9)
ArAr'Co

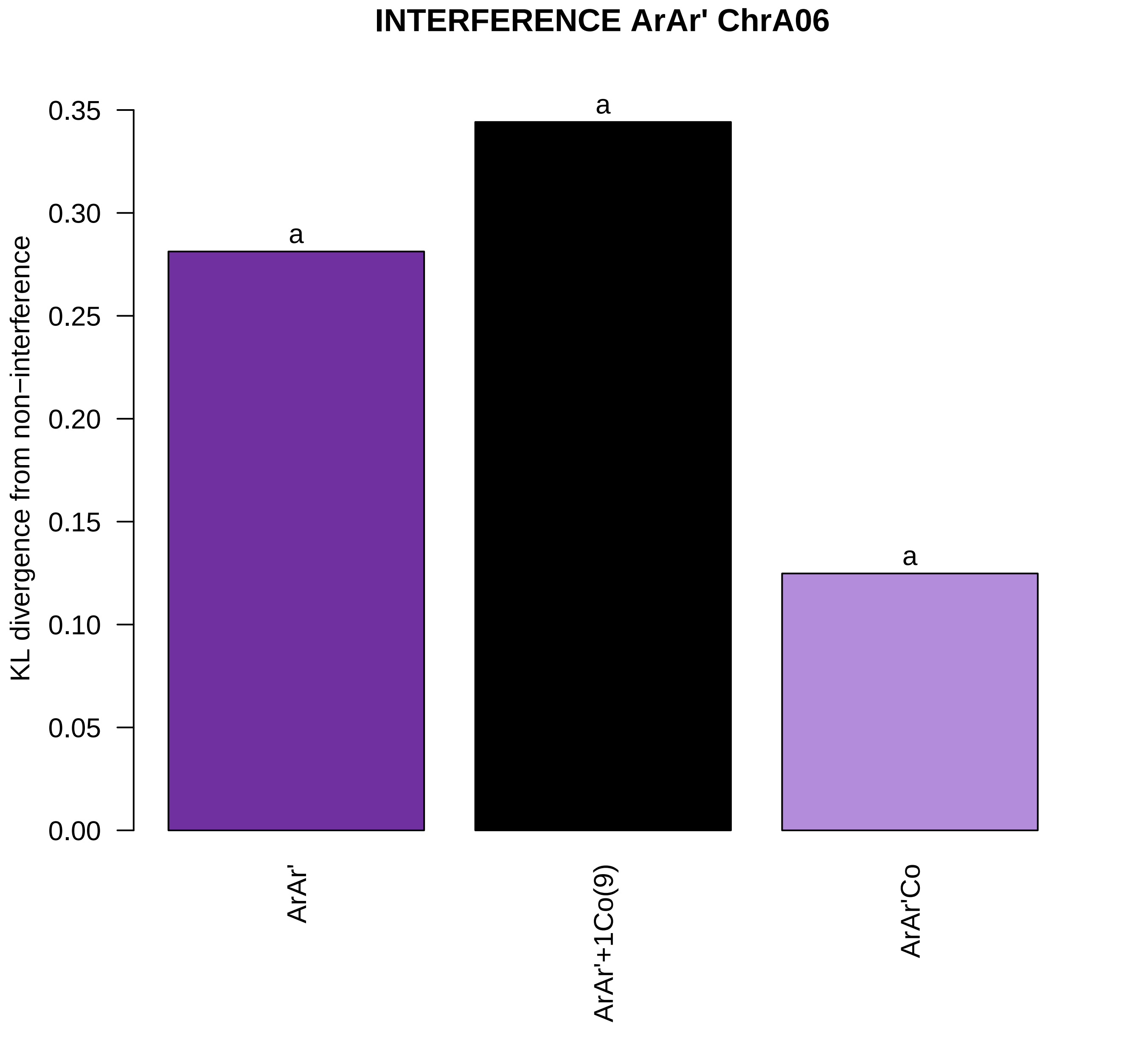
INTERFERENCE ArAr' ChrA06
KL divergence from non−interference
0.35
0.30
0.25
0.20
0.15
0.10
0.05
0.00
a
a
a
ArAr'
ArAr'+1Co(9)
ArAr'Co

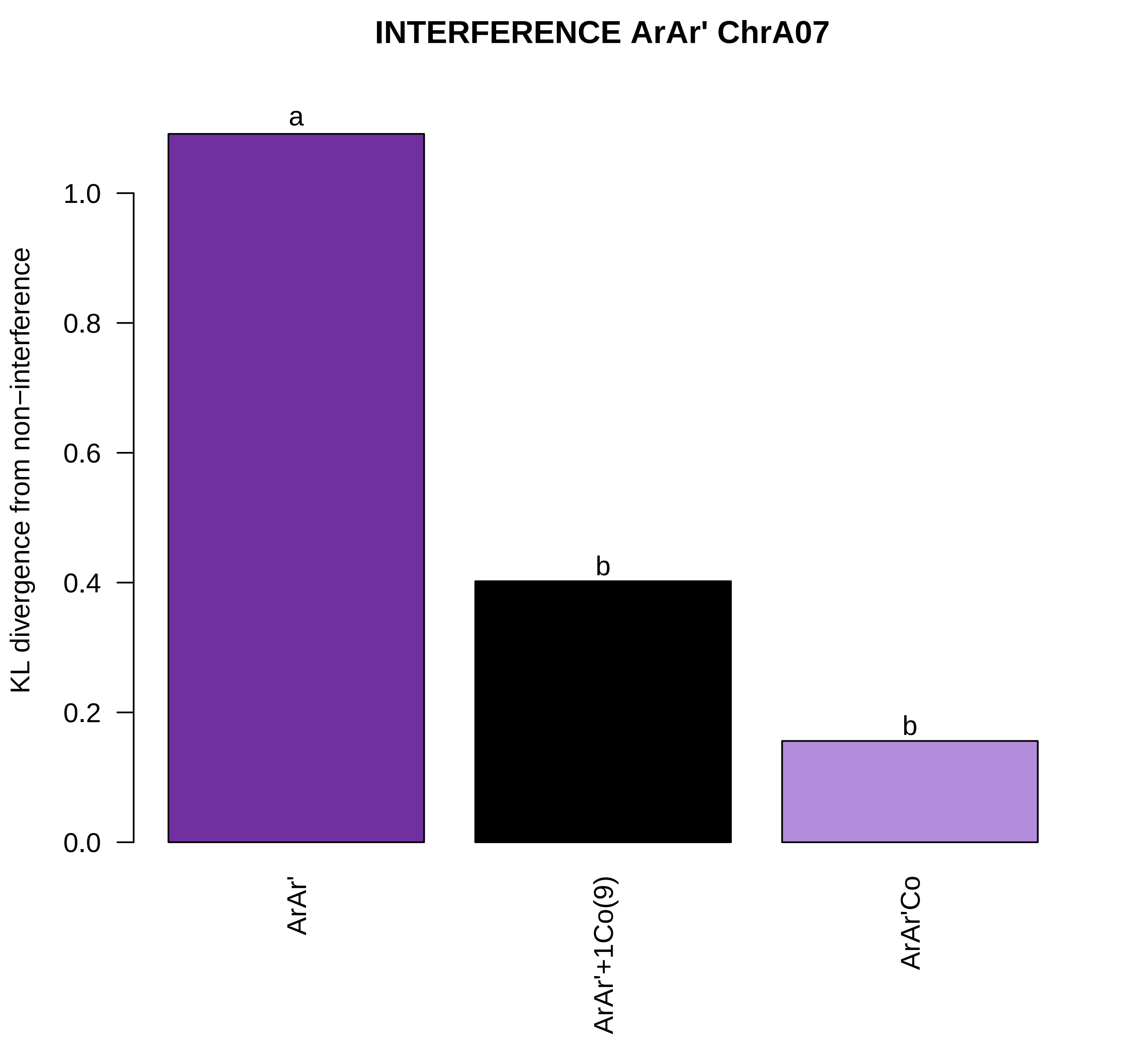
INTERFERENCE ArAr' ChrA07
KL divergence from non−interference
1.0
0.8
0.6
0.4
0.2
0.0
a
b
b
ArAr'
ArAr'+1Co(9)
ArAr'Co

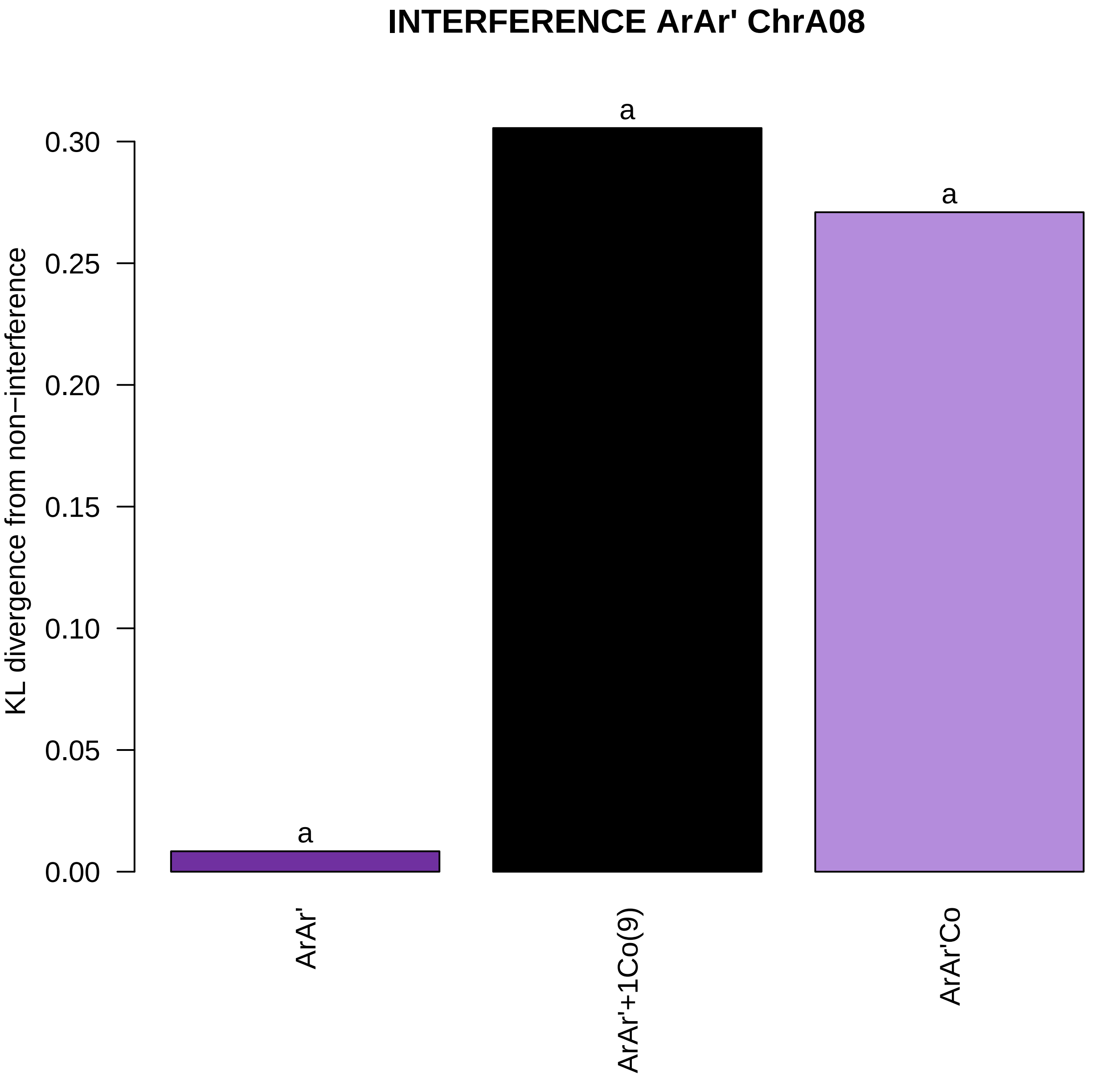
INTERFERENCE ArAr' ChrA08
KL divergence from non−interference
0.30
0.25
0.20
0.15
0.10
0.05
0.00
a
a
a
ArAr'
ArAr'+1Co(9)
ArAr'Co

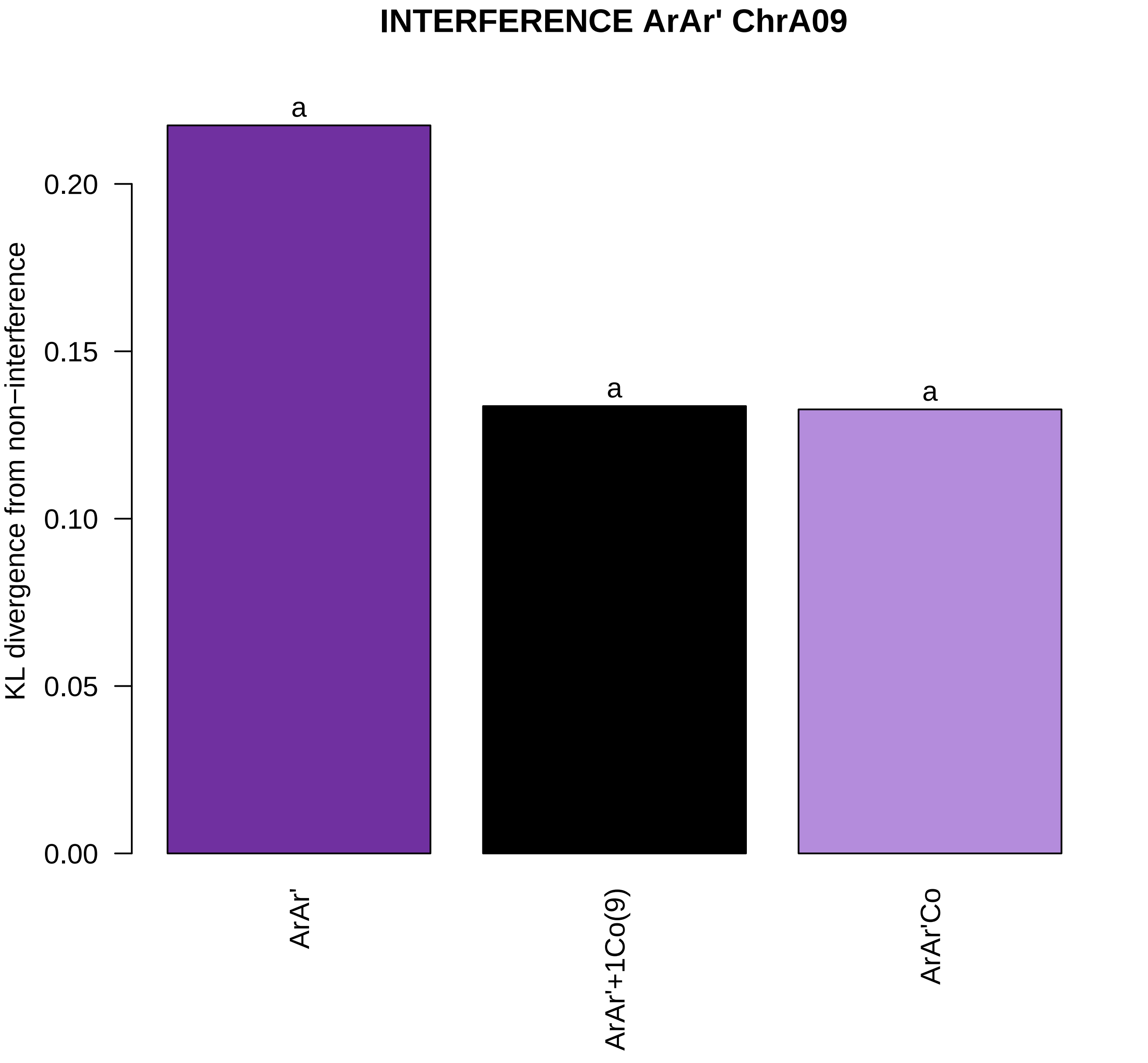
INTERFERENCE ArAr' ChrA09
KL divergence from non−interference
0.00
0.05
0.10
0.15
0.20
a
a
a
ArAr'
ArAr'+1Co(9)
ArAr'Co

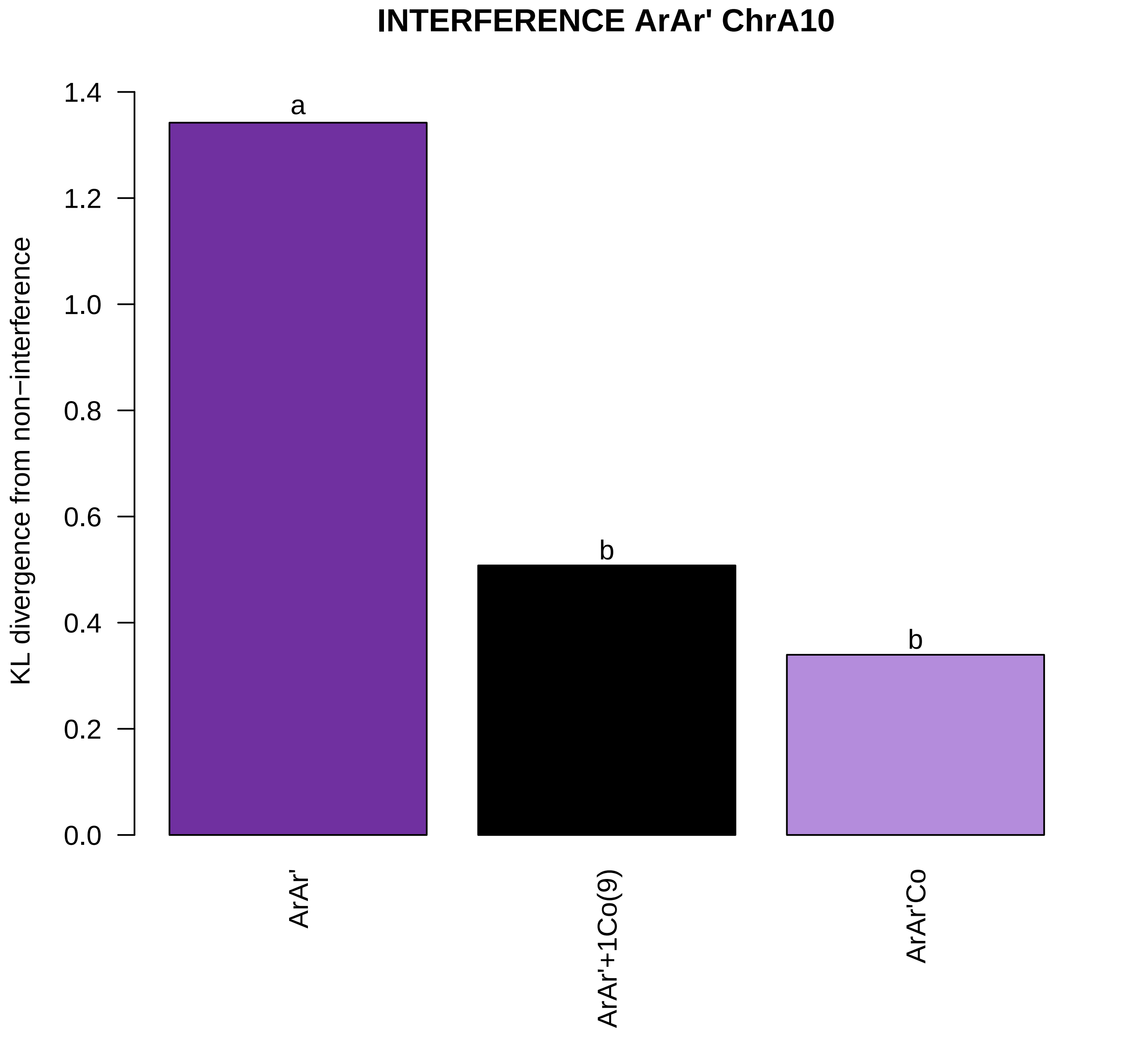
INTERFERENCE ArAr' ChrA10
KL divergence from non−interference
0.0
0.2
0.4
0.6
0.8
1.0
1.2
1.4
a
b
b
ArAr'
ArAr'+1Co(9)
ArAr'Co

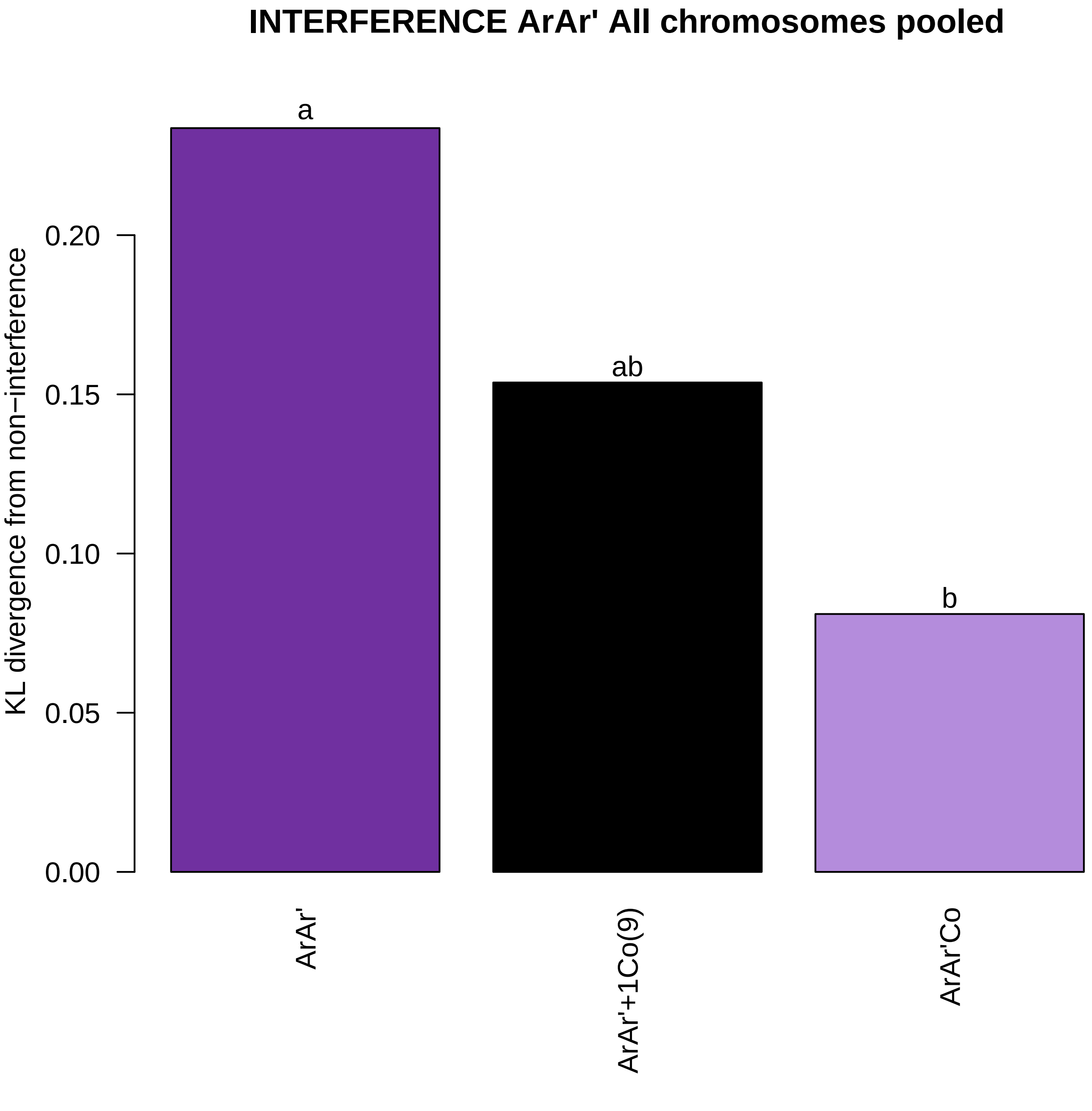
INTERFERENCE ArAr' All chromosomes pooled
KL divergence from non−interference
0.00
0.05
0.10
0.15
0.20
a
ab
b
ArAr'
ArAr'+1Co(9)
ArAr'Co

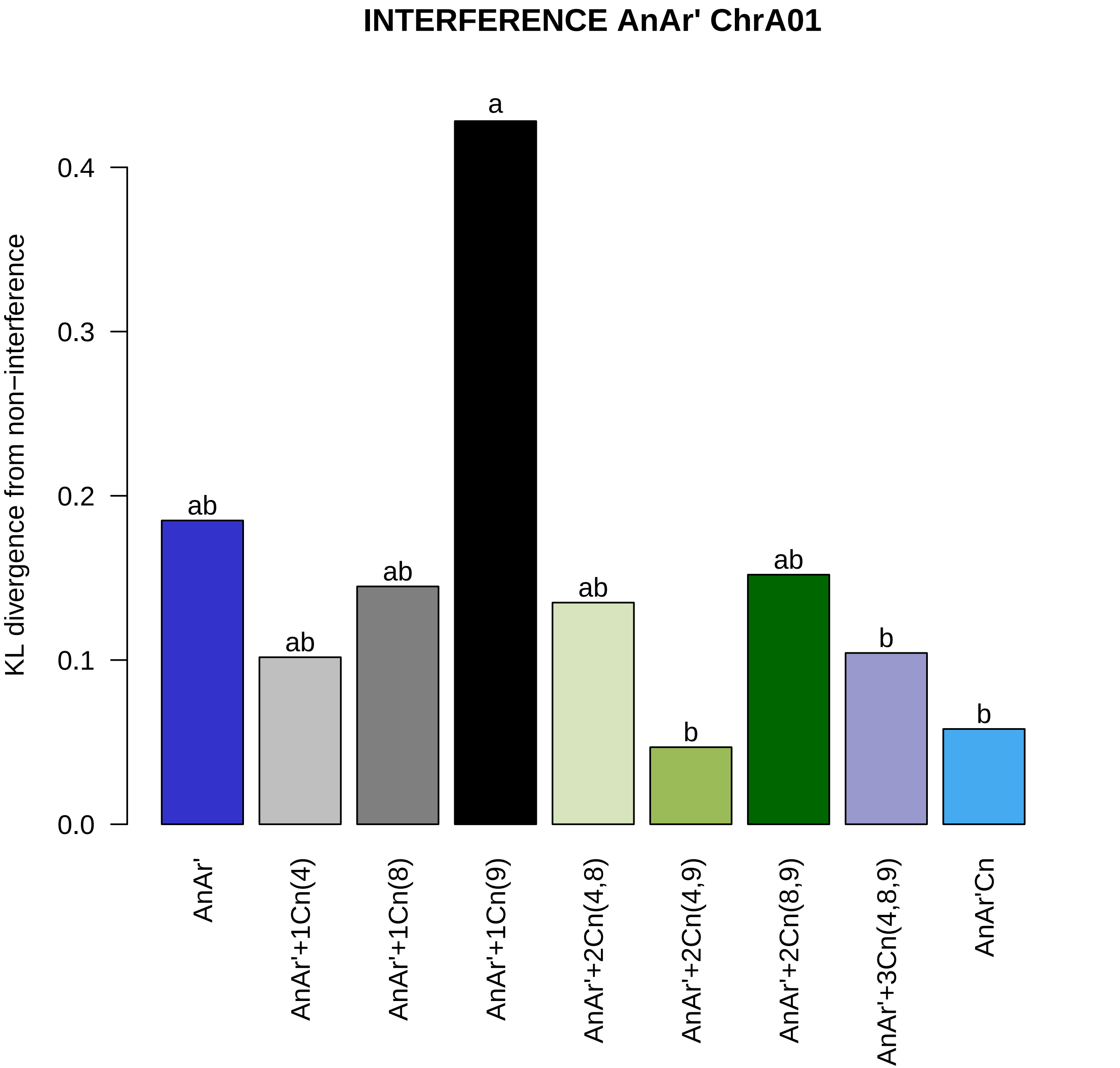

INTERFERENCE AnAr' ChrA01
KL divergence from non−interference
0.0
0.1
0.2
0.3
0.4
ab
ab
ab
a
ab
b
ab
b
b
AnAr'
AnAr'+1Cn(4)
AnAr'+1Cn(8)
AnAr'+1Cn(9)
AnAr'+2Cn(4,8)
AnAr'+2Cn(4,9)
AnAr'+2Cn(8,9)
AnAr'+3Cn(4,8,9)
AnAr'Cn

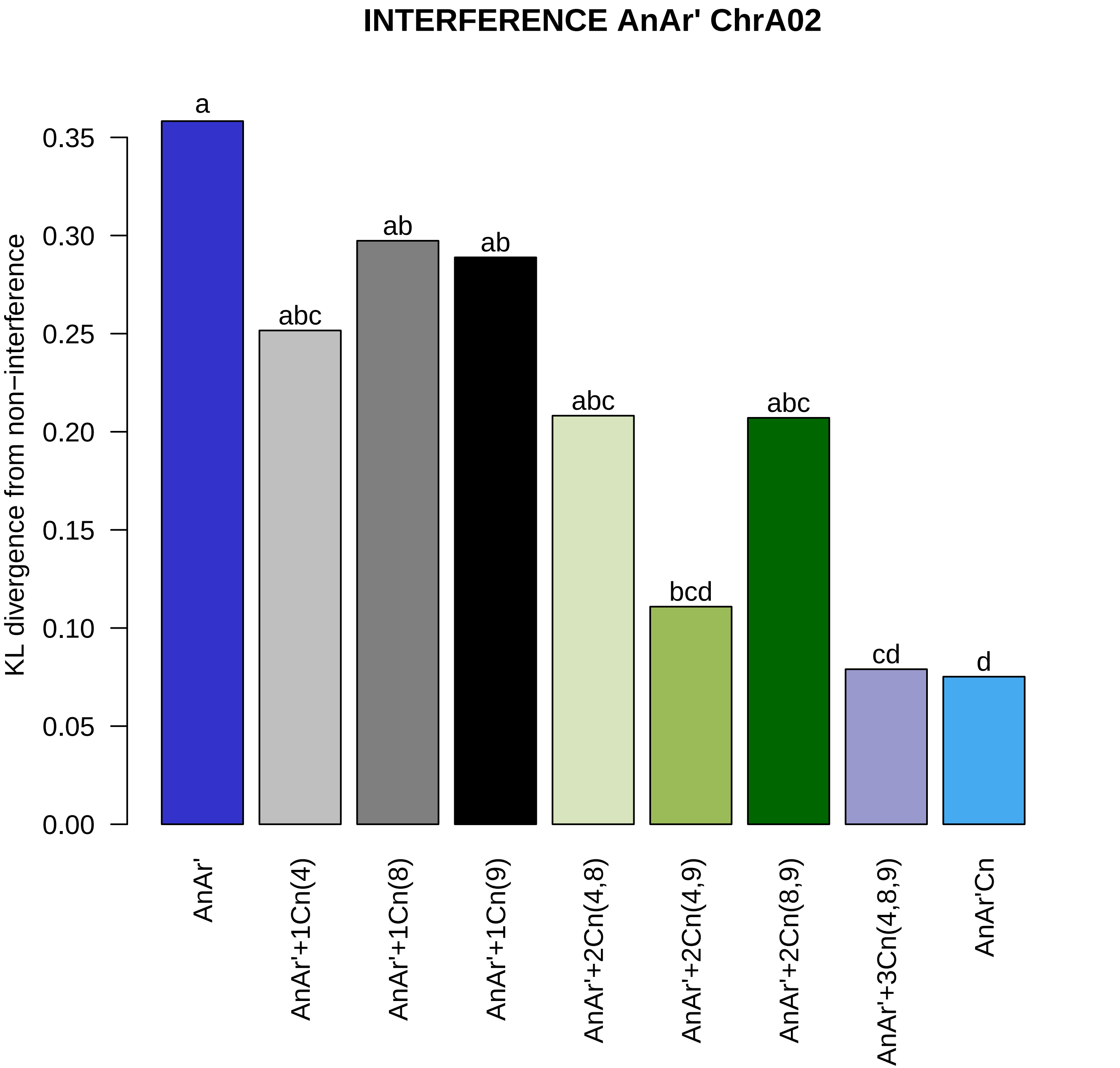

INTERFERENCE AnAr' ChrA02
KL divergence from non−interference
0.00
0.05
0.10
0.15
0.20
0.25
0.30
0.35
a
abc
ab
ab
abc
bcd
abc
cd
d
AnAr'
AnAr'+1Cn(4)
AnAr'+1Cn(8)
AnAr'+1Cn(9)
AnAr'+2Cn(4,8)
AnAr'+2Cn(4,9)
AnAr'+2Cn(8,9)
AnAr'+3Cn(4,8,9)
AnAr'Cn

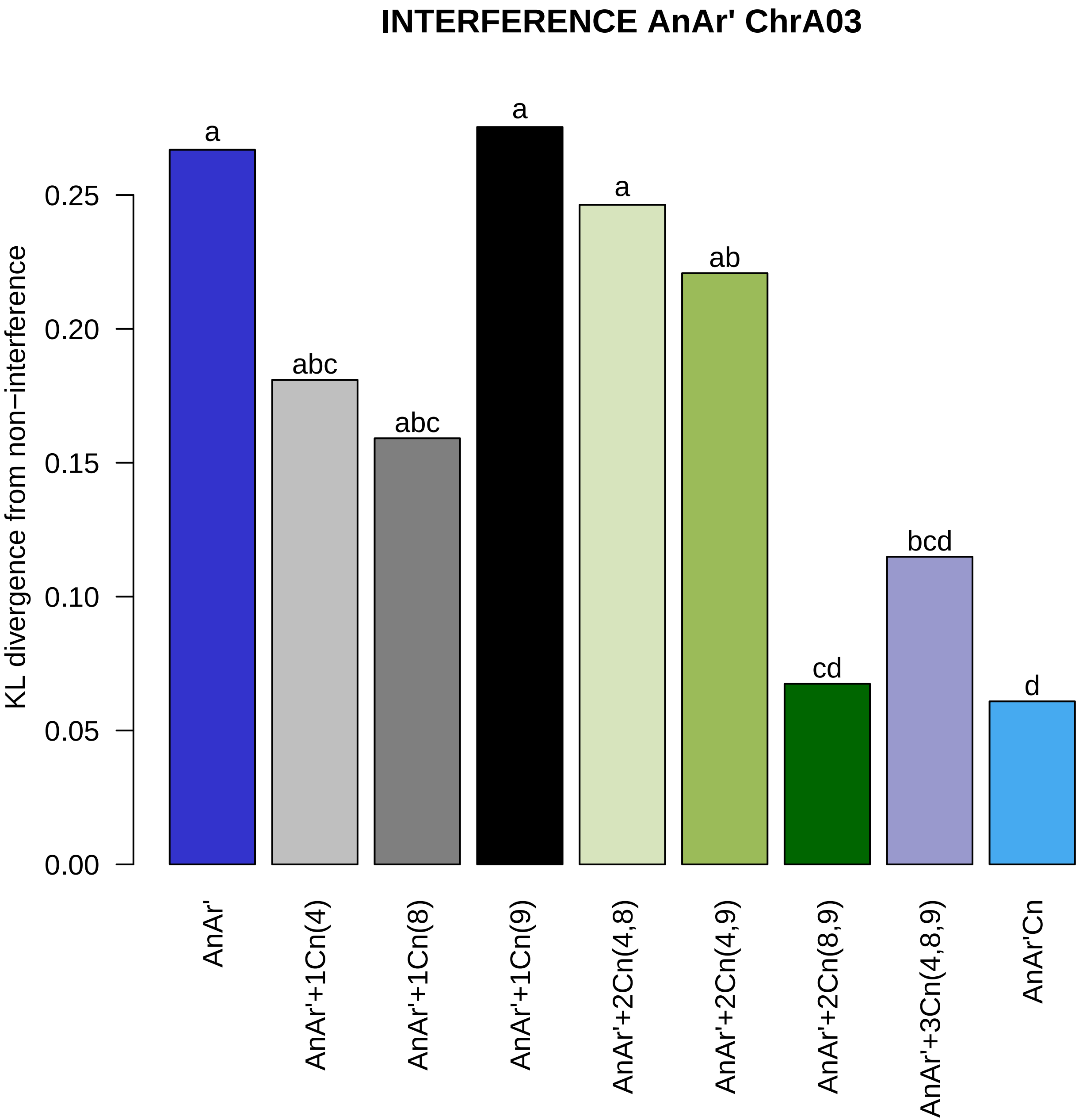
INTERFERENCE AnAr' ChrA03
KL divergence from non−interference
0.00
0.05
0.10
0.15
0.20
0.25
a
abc
abc
a
a
ab
cd
bcd
d
AnAr'
AnAr'+1Cn(4)
AnAr'+1Cn(8)
AnAr'+1Cn(9)
AnAr'+2Cn(4,8)
AnAr'+2Cn(4,9)
AnAr'+2Cn(8,9)
AnAr'+3Cn(4,8,9)
AnAr'Cn

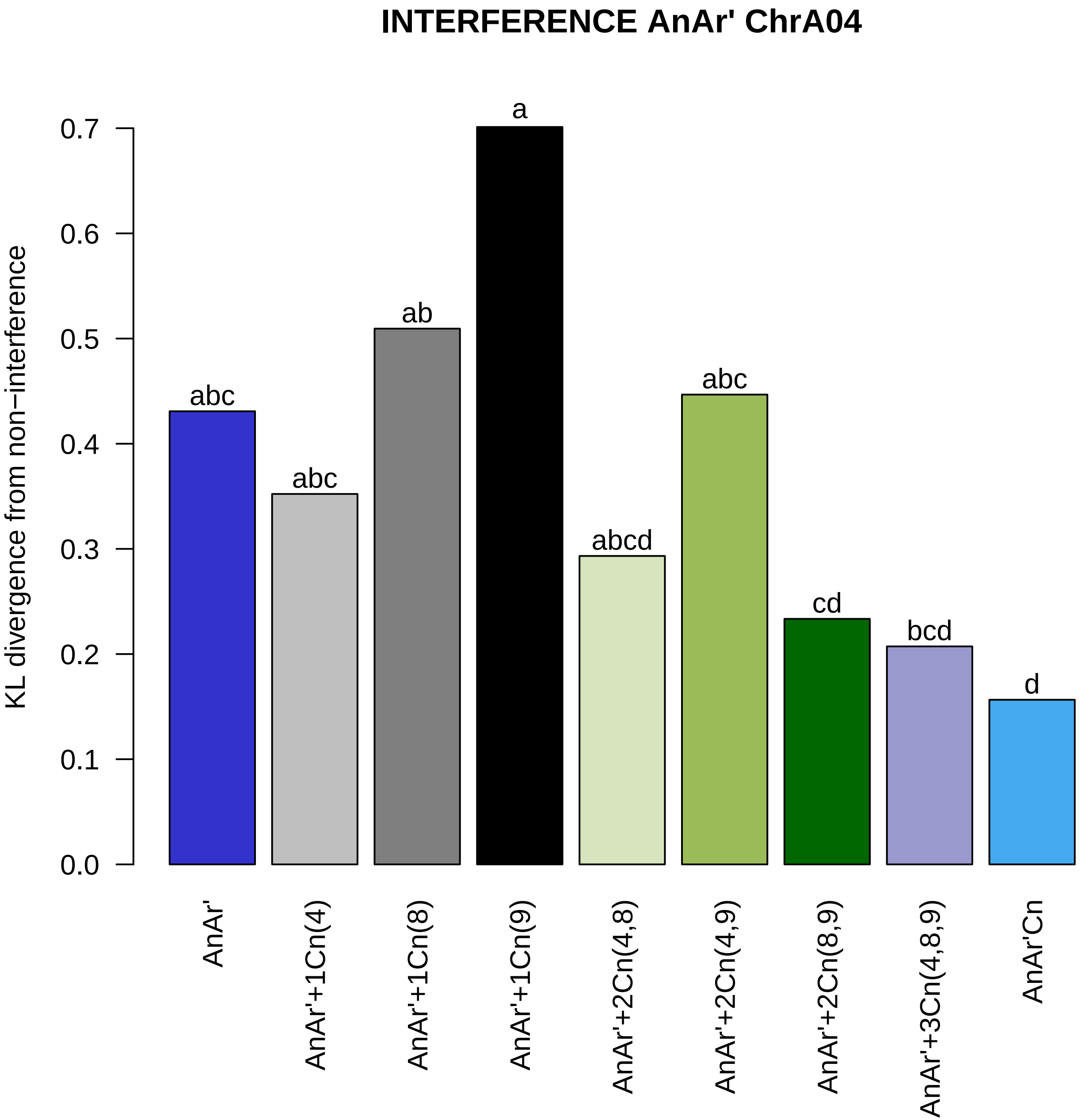
INTERFERENCE AnAr' ChrA04
KL divergence from non−interference
0.0
0.1
0.2
0.3
0.4
0.5
0.6
0.7
abc
abc
ab
a
abcd
abc
cd
bcd
d
AnAr'
AnAr'+1Cn(4)
AnAr'+1Cn(8)
AnAr'+1Cn(9)
AnAr'+2Cn(4,8)
AnAr'+2Cn(4,9)
AnAr'+2Cn(8,9)
AnAr'+3Cn(4,8,9)
AnAr'Cn

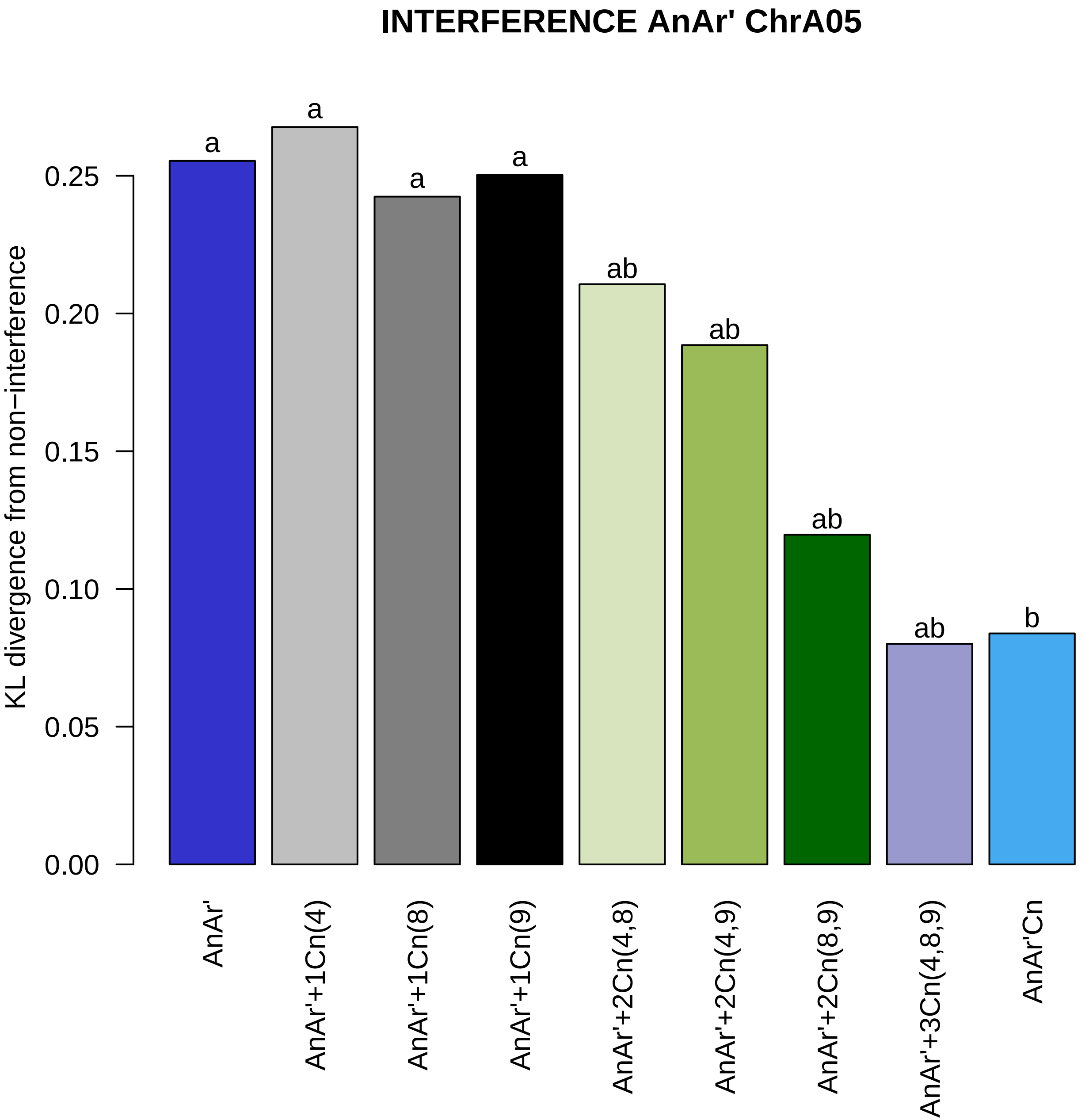
INTERFERENCE AnAr' ChrA05
KL divergence from non−interference
0.00
0.05
0.10
0.15
0.20
0.25
a
a
a
a
ab
ab
ab
ab
b
AnAr'
AnAr'+1Cn(4)
AnAr'+1Cn(8)
AnAr'+1Cn(9)
AnAr'+2Cn(4,8)
AnAr'+2Cn(4,9)
AnAr'+2Cn(8,9)
AnAr'+3Cn(4,8,9)
AnAr'Cn

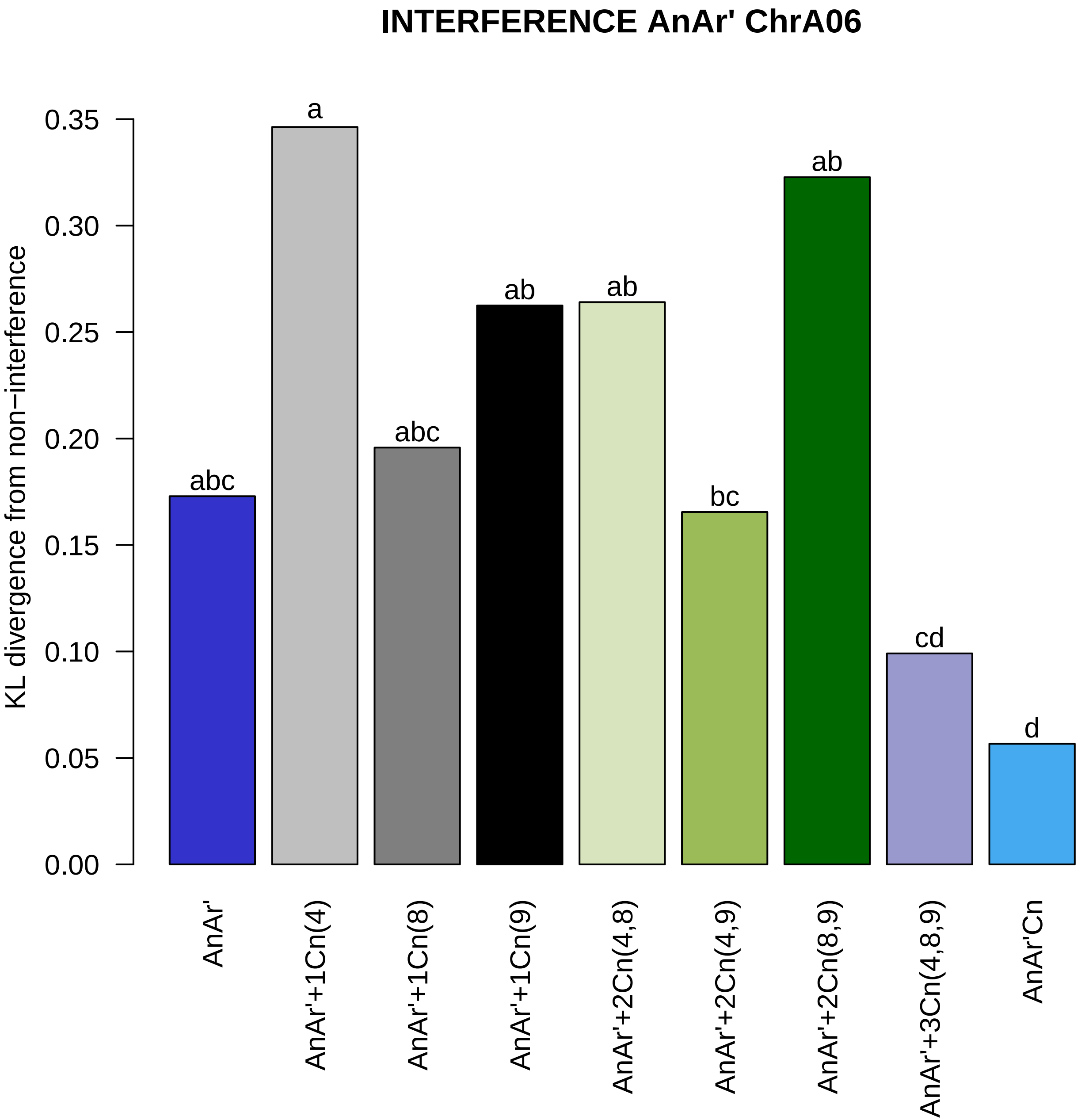
INTERFERENCE AnAr' ChrA06
KL divergence from non−interference
0.00
0.05
0.10
0.15
0.20
0.25
0.30
0.35
abc
a
abc
ab
ab
bc
ab
cd
d
AnAr'
AnAr'+1Cn(4)
AnAr'+1Cn(8)
AnAr'+1Cn(9)
AnAr'+2Cn(4,8)
AnAr'+2Cn(4,9)
AnAr'+2Cn(8,9)
AnAr'+3Cn(4,8,9)
AnAr'Cn

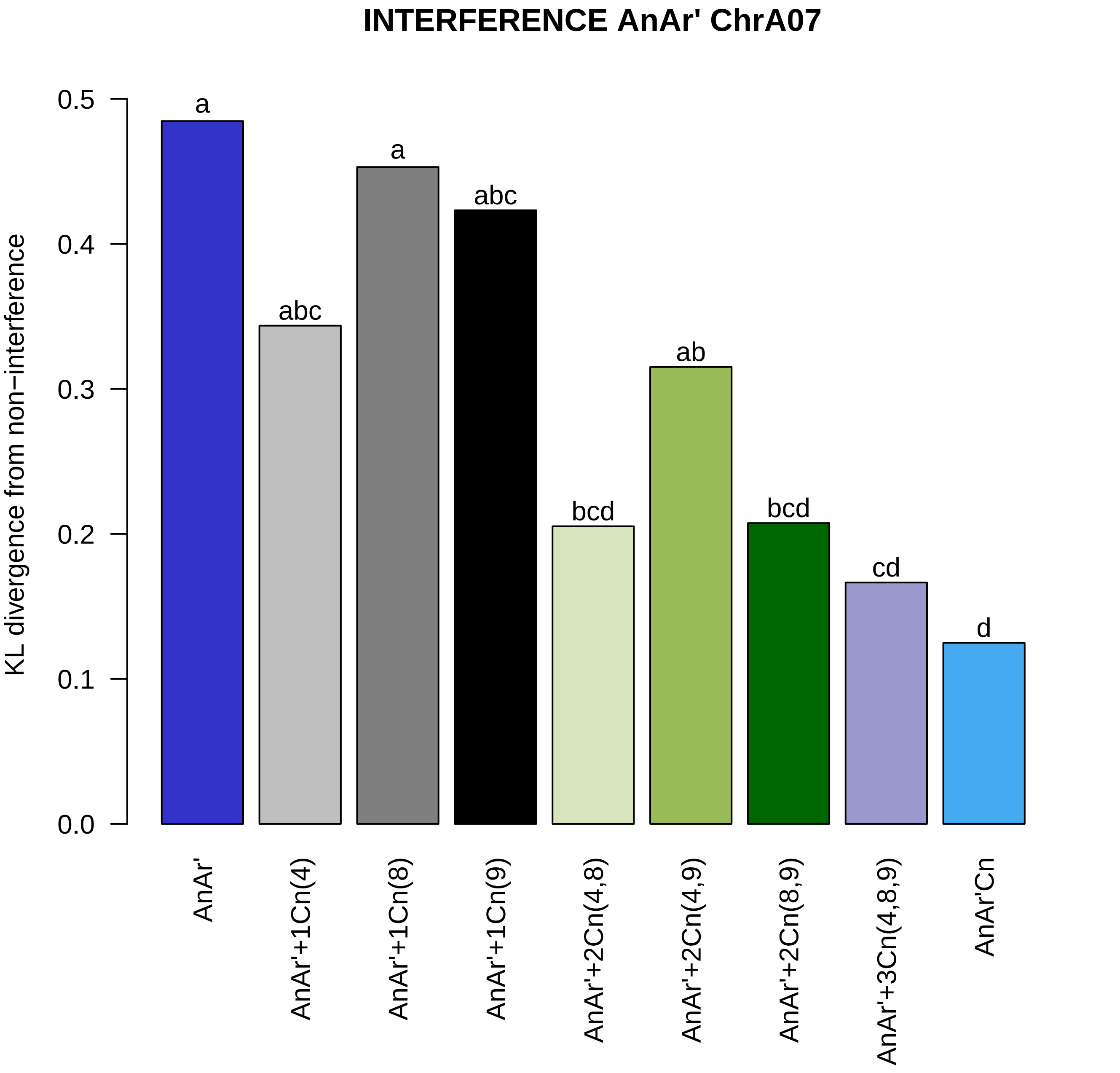
INTERFERENCE AnAr' ChrA07
KL divergence from non−interference
0.0
0.1
0.2
0.3
0.4
0.5
a
abc
a
abc
bcd
ab
bcd
cd
d
AnAr'
AnAr'+1Cn(4)
AnAr'+1Cn(8)
AnAr'+1Cn(9)
AnAr'+2Cn(4,8)
AnAr'+2Cn(4,9)
AnAr'+2Cn(8,9)
AnAr'+3Cn(4,8,9)
AnAr'Cn

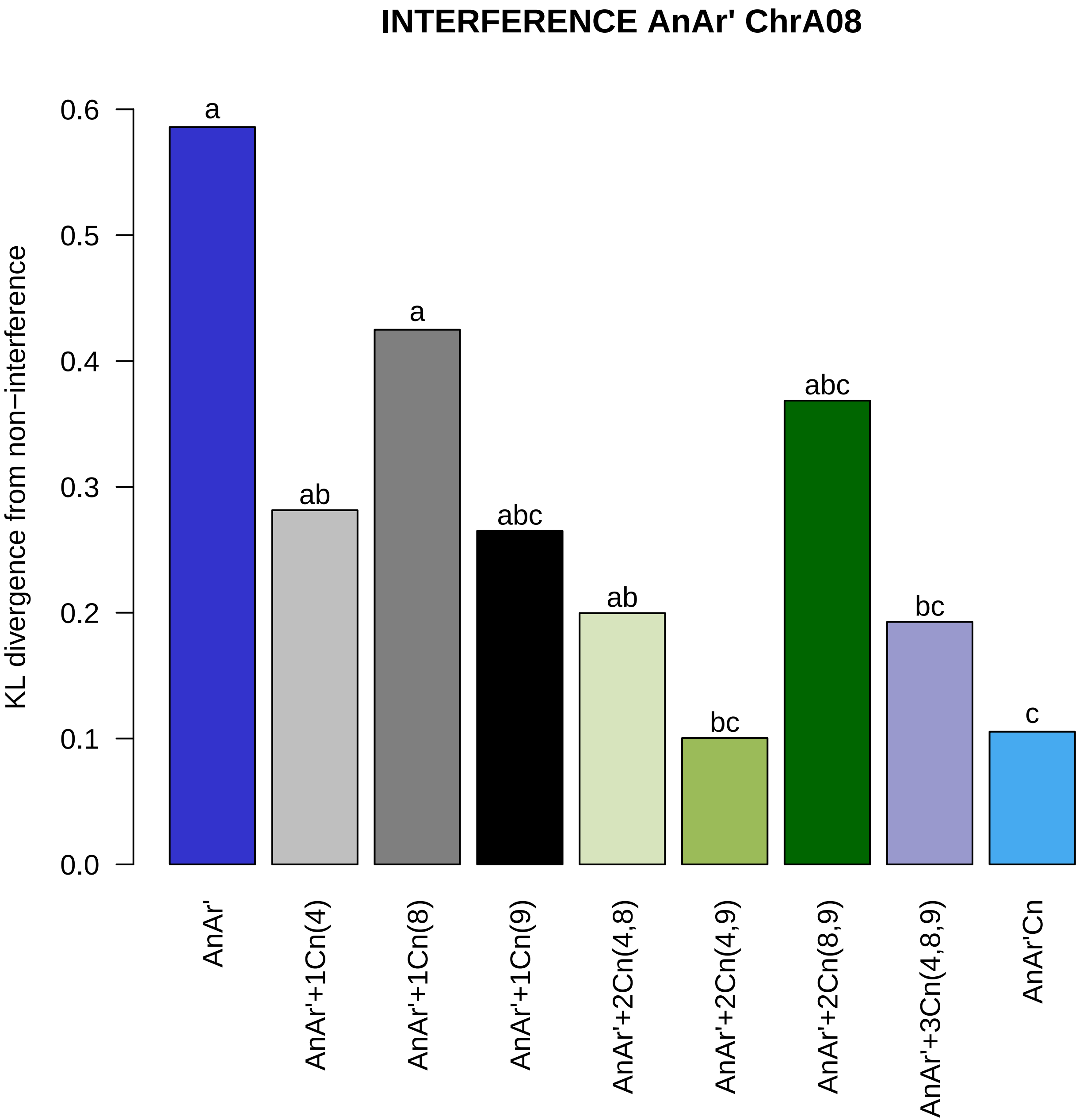

INTERFERENCE AnAr' ChrA08
KL divergence from non−interference
0.0
0.1
0.2
0.3
0.4
0.5
0.6
a
ab
a
abc
ab
bc
abc
bc
c
AnAr'
AnAr'+1Cn(4)
AnAr'+1Cn(8)
AnAr'+1Cn(9)
AnAr'+2Cn(4,8)
AnAr'+2Cn(4,9)
AnAr'+2Cn(8,9)
AnAr'+3Cn(4,8,9)
AnAr'Cn

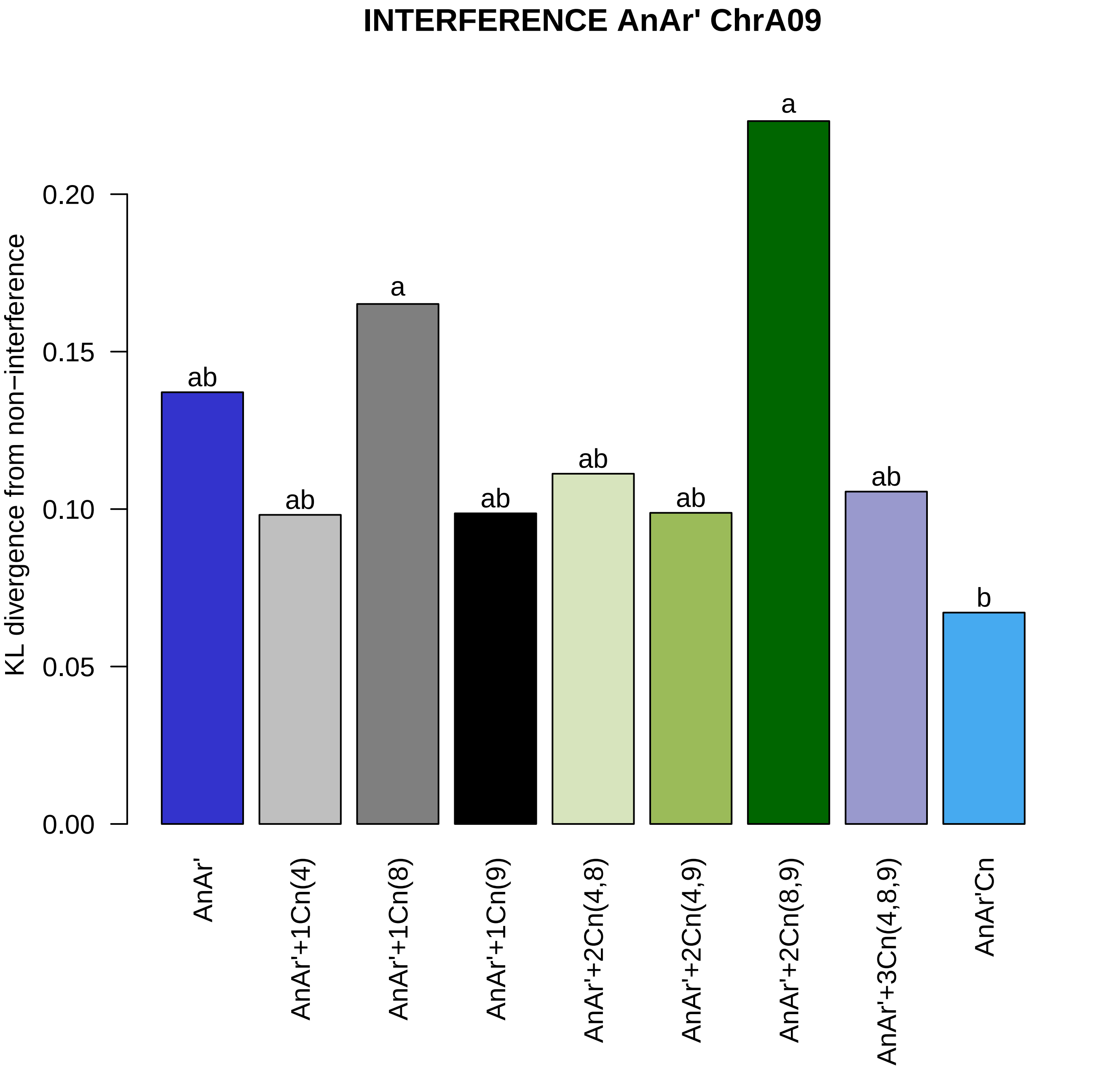
INTERFERENCE AnAr' ChrA09
KL divergence from non−interference
0.00
0.05
0.10
0.15
0.20
ab
ab
a
ab
ab
ab
a
ab
b
AnAr'
AnAr'+1Cn(4)
AnAr'+1Cn(8)
AnAr'+1Cn(9)
AnAr'+2Cn(4,8)
AnAr'+2Cn(4,9)
AnAr'+2Cn(8,9)
AnAr'+3Cn(4,8,9)
AnAr'Cn

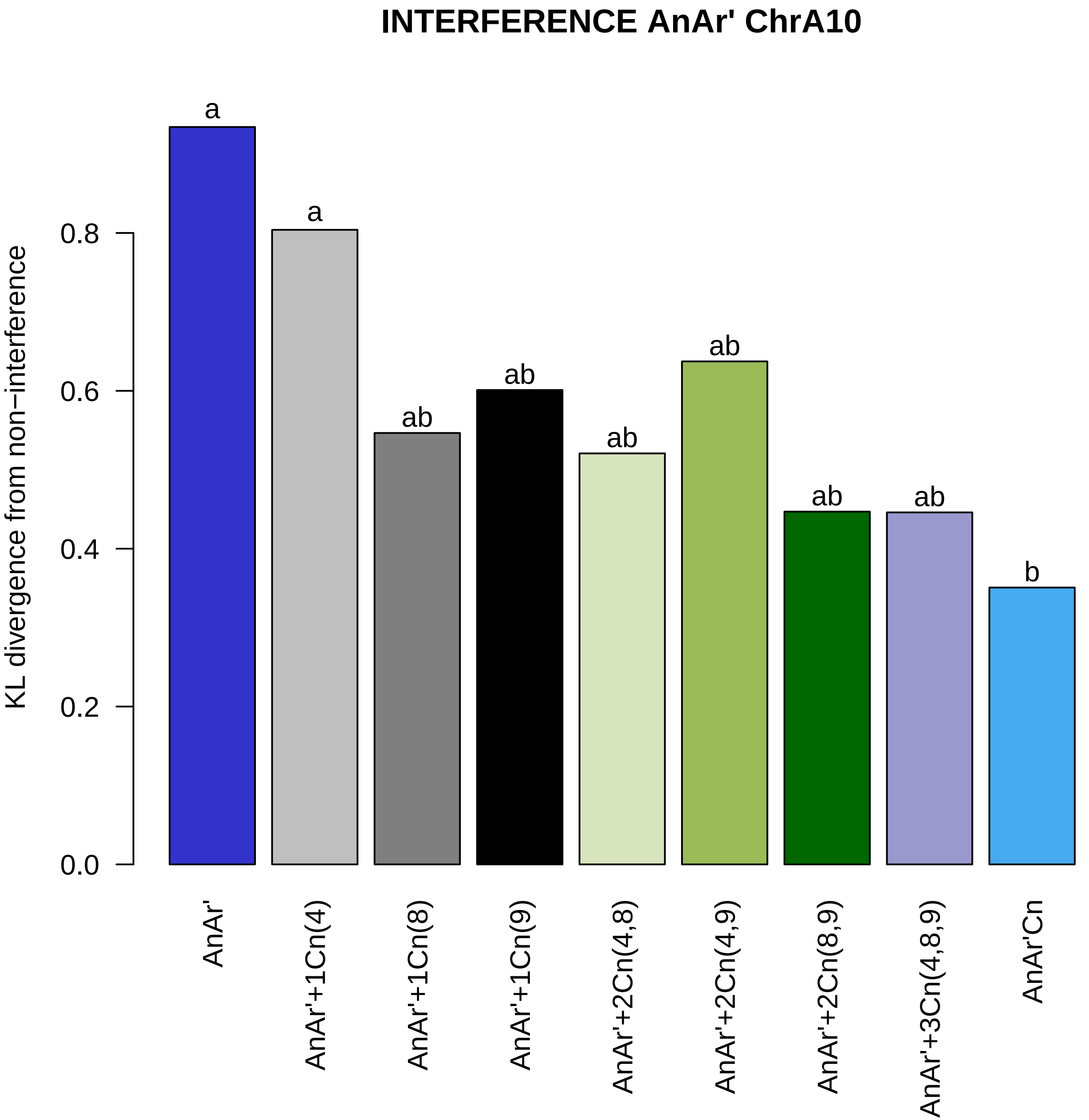
INTERFERENCE AnAr' ChrA10
KL divergence from non−interference
0.0
0.2
0.4
0.6
0.8
a
a
ab
ab
ab
ab
ab
ab
b
AnAr'
AnAr'+1Cn(4)
AnAr'+1Cn(8)
AnAr'+1Cn(9)
AnAr'+2Cn(4,8)
AnAr'+2Cn(4,9)
AnAr'+2Cn(8,9)
AnAr'+3Cn(4,8,9)
AnAr'Cn

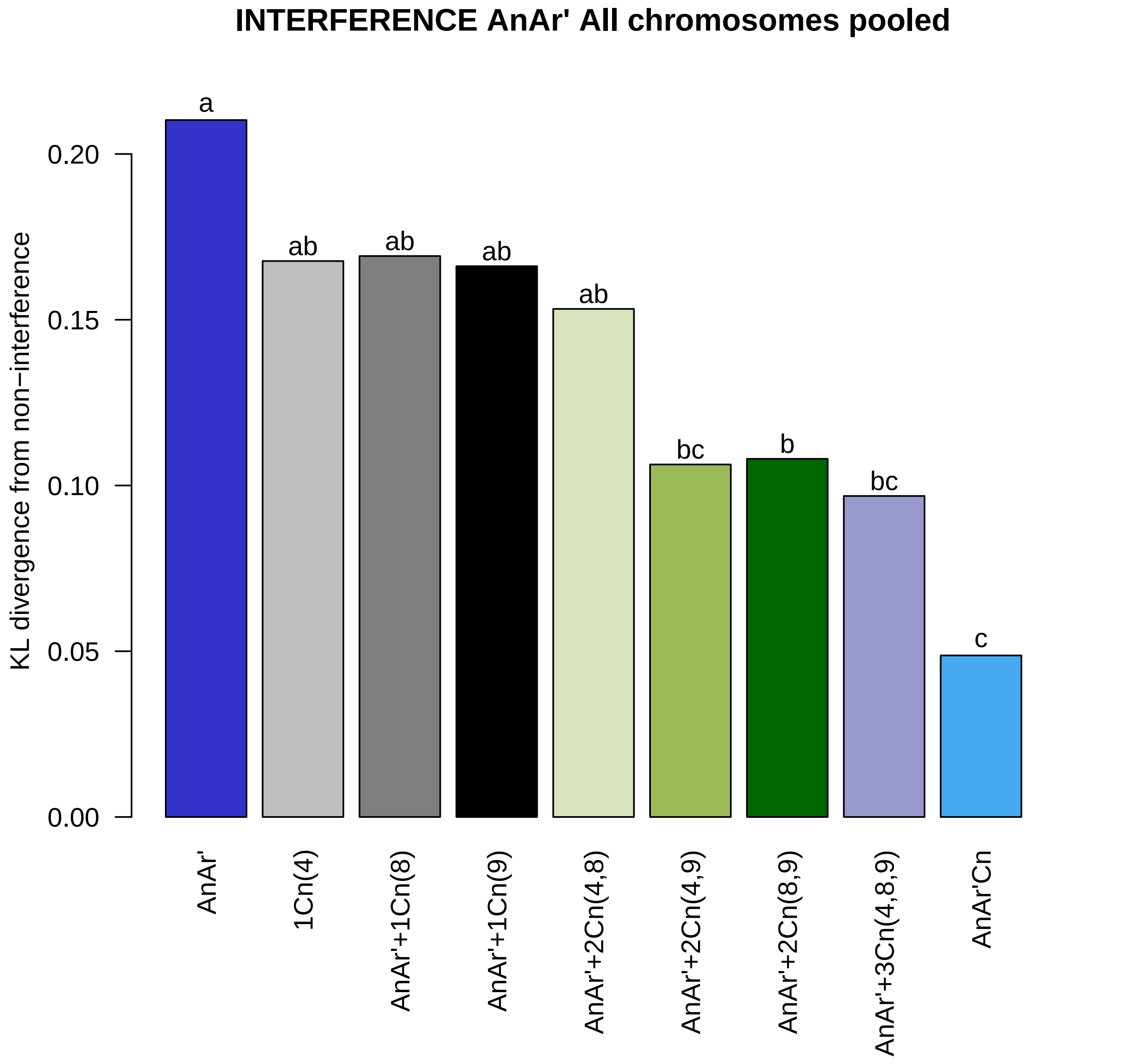

INTERFERENCE AnAr' All chromosomes pooled
KL divergence from non−interference
0.00
0.05
0.10
0.15
0.20
a
ab
ab
ab
ab
bc
b
bc
c
AnAr'
1Cn(4)
AnAr'+1Cn(8)
AnAr'+1Cn(9)
AnAr'+2Cn(4,8)
AnAr'+2Cn(4,9)
AnAr'+2Cn(8,9)
AnAr'+3Cn(4,8,9)
AnAr'Cn
